# Exceeding expectations: the genomic basis of nitrogen utilization efficiency and integrated trait plasticity as avenues to improve nutrient stress tolerance in cultivated sunflower (*Helianthus annuus* L.)

**DOI:** 10.1101/2022.08.28.505579

**Authors:** Andries A. Temme, Kelly L. Kerr, Kristen M. Nolting, Emily L. Dittmar, Rishi R. Masalia, Alexander Bucksch, John M. Burke, Lisa A. Donovan

## Abstract

Maintaining crop productivity is a challenge as population growth, climate change, and increasing fertilizer costs necessitate expanding crop production to poorer lands whilst reducing inputs. Enhancing crops’ nutrient use efficiency is thus an important goal, but requires a better understanding of related traits and their genetic basis.

We investigated variation in low nutrient stress tolerance in a diverse panel of cultivated sunflower genotypes grown under high and low nutrient conditions, assessing relative growth rate (RGR) as performance. We assessed variation in traits related to nitrogen utilization efficiency (NUtE), mass allocation, and leaf elemental content.

Across genotypes, nutrient limitation reduced RGR. Moreover, higher vigor (higher control RGR) was associated with a greater absolute decrease under stress. Given this trade-off, we focused on nutrient stress tolerance independent from vigor. This tolerance metric correlated with the change in NUtE, plasticity for a suite of morphological traits, and leaf element content. Genome-wide association analyses revealed regions associated with variation and plasticity in multiple traits, including two key regions with ostensibly additive effects on NUtE change.

Our results demonstrate potential avenues for improving sunflower nutrient stress tolerance independent from vigor and highlight specific traits and genomic regions that could play a role in enhancing tolerance.

**Highlight:** Genetic associations and trait correlations show that, in cultivated sunflower, selection for increased nitrogen utilization efficiency and plasticity in key traits is a promising avenue for increasing nutrient stress tolerance.

## Introduction

Rising population levels and climate change are increasing pressures on our global agricultural system to realize higher productivity on marginal lands (Ramankutty *et al*., 2018). Additionally, demand for oilseed crops is projected to increase 90% by 2050 as compared to 2007 (Tilman *et al*., 2011; Alexandratos and Bruinsma, 2012; Ramankutty *et al*., 2018). To meet this challenge, the utility of increased fertilizer inputs is hampered by increasing cost of fertilizer and large negative impacts of fertilizer on the environment (Vitousek *et al*., 1997; Robertson and Vitousek, 2009). A more sustainable strategy is to improve the nutrient use efficiency of crops, i.e. increase productivity per unit of available nutrients (Xu *et al*., 2012). However, maintaining or even expanding productivity under low nutrient availability is a challenge.

Prior work on cultivated sunflower, a major oilseed crop, has revealed genotypic variation in tolerance to salt, flooding, and drought stress (Masalia *et al*., 2018; Gao *et al*., 2019; Temme *et al*., 2020). Improvements in cultivated sunflower nutrient uptake and use efficiency can be key in meeting demand while maintaining or even reducing inputs. Prior work on sunflower nutrient stress response in a small number of genotypes showed a remarkable consistency in the performance ranking of genotypes across a range of nutrient levels (i.e., high performing genotypes tend to exhibit high performance across conditions, with lower performing genotypes exhibiting low performance across conditions; (Bowsher *et al*., 2017). Here, we more fully explore the effects of nutrient stress in cultivated sunflower and seek to identify key traits and genomic regions with a potential for improving nutrient stress tolerance.

Evaluating stress tolerance in the context of vigor (growth/performance in benign conditions) is critical for improving resource use efficiency without associated decreases in productivity. Given the potential for a trade-off between vigor and the response to stress (e.g., (Temme *et al*., 2020), relating tolerance directly to stress response runs the risk of confounding tolerance with low vigor. Rather, by taking this negative relationship into consideration, we can score genotypes on being more or less tolerant than expected based on their vigor. Thus, tolerance is defined as performing better than would be expected based on this trade-off. This “expectation-deviation tolerance” (ExDev-tolerance) metric makes it possible to isolate those traits or genomic regions that are directly involved in stress response independent of those underlying high performance (Temme *et al*., 2020). However, this raises the question of whether there are trade-offs (sensu (Agrawal, 2020) in growth and development under stressful vs. benign conditions and whether different suites of traits are associated with performance and tolerance.

Much progress has been made in understanding how nitrogen (N) use efficiency (growth per unit of available N) and its components, nitrogen uptake efficiency (efficiency of gathering N from soil) and nitrogen utilization efficiency (NUtE, growth per unit acquired N) can be improved (Han *et al*., 2015; Tegeder and Masclaux-Daubresse, 2018; Swarbreck *et al*., 2019). Realizing improved NUtE could involve efficient (re)distribution of N (and phosphorus) to upper parts of the canopy, altering mass allocation, and adjusting leaf mass per area and/or having more efficient photosynthetic machinery (Lammerts van Bueren and Struik, 2017). Understanding how these trait changes relate to ExDev-tolerance and NUtE is a potential avenue for improving tolerance independent from vigor.

Increased growth and development rate has been heavily selected for during domestication and crop improvement (Milla *et al*., 2018). In improving plant traits for specific environments the multivariate nature of trait covariation should be considered. Traits tend to covary due to a complex web of interactions (Poorter *et al*., 2013, 2019, 2021). Identifying independent axes of trait variation can help focus improvement efforts to enhance tolerance independent from vigor. Improving overall nitrogen use efficiency will require integrating physiology and breeding to target the many interrelated (integrated) traits related to nitrogen uptake efficiency and NUtE.

Due to the multivariate nature of trait covariation, breeding efforts to select on particular traits can have unintended effects on other traits (Svensson *et al*., 2021; Chebib and Guillaume, 2021). Due to pleiotropy or close linkage among traits, selection on one trait can affect others. Previous work on cultivated sunflower has shown a genomic landscape of trait colocalization where certain genomic regions are associated with a range of diverse traits (Masalia *et al*., 2018; Temme *et al*., 2020). By studying this landscape of trait colocalization we can identify those regions with minimal effects on other traits in order to decouple and adjust this network of trait covariation. Thus, developing breeding strategies aimed at improving yields under a range of environmental conditions is facilitated by an understanding of the genomic regions underlying trait variation.

To determine the effects of low nutrient stress on key traits and genomic regions linked to improving nutrient stress tolerance in cultivated sunflower, we asked the following questions:

1. What is the relationship between vigor (growth under benign conditions) and the decline in performance in response to low nutrient stress?
2. What is the relationship between nitrogen utilization efficiency (NUtE) and nutrient stress tolerance independent of vigor (ExDev-tolerance)?
3. What is the effect of nutrient stress on traits potentially related to nutrient stress tolerance (e.g. morphology and leaf elemental content)?
4. Which suites of trait variation and/or trait plasticity relate to ExDev-tolerance and vigor?
5. Can we identify shared and unique genomic regions associated with trait variation across a range of traits?

## Methods

We grew a subset of 261 (out of 287) genotypes of the Sunflower Association Mapping (SAM) population (Mandel *et al*., 2011, 2013), which captures ca. 95% of the allelic diversity in cultivated sunflower. The SAM population includes both heterotic groups (i.e., male [RHA] and female [HA] lines) as well as both major market types (i.e., oil and non-oil [confectionery] lines). The SAM population has been used extensively for GWAS studies (Masalia *et al*., 2018; Gao *et al*., 2019; Temme *et al*., 2020; Stahlhut *et al*., 2021) because of the substantial genetic/trait diversity contained within the population, relevant commercial uses, and the availability of whole genome re-sequencing data for the entire population.

In spring of 2016, we grew individuals of 261 SAM genotypes in a randomized block experimental design with 2 treatments and 4 replicates in the Botany greenhouses at the University of Georgia (Athens, GA, USA). Seeds were germinated in sand and transplanted to pots 7 days after sowing. Individuals were grown in 7.6 liter pots filled with a 3:1 mixture of sand and turface (Turface Athletics, PROFILE Products, LLC, Buffalo Grove, IL) to improve water holding capacity. Two individuals per genotype were randomly assigned to each of four greenhouse bays resulting in 2,088 experimental plants. In each greenhouse bay, the two individuals of each genotype received either 80 g or 8 g of Osmocote plus (15-12-9 NPK; ScottsMiracle-Gro, Marysville, OH) slow release fertilizer to simulate favorable nutrient conditions and a broad spectrum nutrient deficiency. We supplemented with calcium using 5 ml gypsum (Performance Minerals Corporation, Birmingham, AL) and 5 ml lime powder (Austinville Limestone, Austinville, VA) per pot because prior experience has shown that calcium limitation results in developmental abnormalities under greenhouse conditions. Plants were watered daily, switching to twice daily when pots started to dry out due to temperature and plant size to prevent water stress.

Individual plants were tagged when they reached floral initiation (budding) (R1 stage) and harvested when they reached R2 stage, when the peduncle (flower stalk) had elongated to the point at which the primary bud was >1 cm above the nearest leaves (Schneiter and Miller, 1981). The process of harvesting plants at a specified developmental stage allowed us to identify the effect of nutrient stress on flowering time and early flower development while minimizing pot size constraints on biomass.

At harvest, plants were measured for height (to the nearest 0.5 cm from the base of the stem to the top of the stem), stem diameter (using calipers halfway between the soil and cotyledons), and chlorophyll content index (MC-100, Apogee Instruments, Inc., Logan, UT). Plant biomass was separated into the most recent fully expanded leaf (MRFEL), all other leaves, stem and branches (including bud[s]), and root. The MRFEL is an easy to define specific leaf in sunflower that standardizes leaf selection to a fully developed leaf toward the top of the plant. Generally, this leaf is in the upper 10% of the plant with the number of underdeveloped leaves higher up on the stem varying by genotype. Roots were stored in a chilled environment and washed in order of harvest. Images were taken of the MRFEL at 300dpi and of a single lateral root (near the soil surface but < 2 mm diameter) at 600dpi with a flatbed scanner (Canon CanoScan LiDE120) for use in determining leaf mass per area (LMA) and specific root length (SRL). MRFEL scans were measured for area using imageJ (Schneider *et al*., 2012). Root scans were analyzed for SRL using RhizoVision Explorer v2.0.3 (Seethepalli *et al*., 2021) to calculate total length, median root diameter, and branching frequency of the root sample. Leaf mass per area (LMA) was calculated by dividing the MRFEL weight (without petiole) by its measured area. Specific root length was calculated by dividing measured root length by the mass of the sampled root.

After oven drying at 60°C for at least 48 hrs, dried samples were stored until weighing. Prior to weighing, all samples were redried at 60°C for at least 2 hrs. After drying, lateral roots were separated from the taproot (up to the point at which the taproot and lateral roots had similar widths), and the primary and axillary buds were separated from the stem. The resulting biomass samples weighed separately for each individual were: the MRFEL, remaining leaves, stem, primary bud, axillary buds, taproot, lateral root, and SRL root sample. Biomass fractions including: root mass fraction (RMF), leaf mass fraction (LMF), and stem mass fraction (SMF), were calculated by dividing component parts by the total summed individual plant weight. Root mass fraction was further divided into tap and fine root mass as fractions of the whole plant weight and root biomass.

After weighing, the MRFEL samples (without petiole) were pooled per genotype and treatment. Samples were coarse ground using a Wiley Mill (Thomas Scientific, Swedesboro, NJ) and the resulting powder was homogenized. A 2 ml sample of leaf powder was then transferred to an Eppendorf tube and ground to a fine powder using a metal bead in a Tissuelyzer (Qiagen, Germantown, MD). The finely powdered leaf tissue was then sent to Midwest Laboratories (Omaha, NE) for ICP-MS analysis to determine the amounts of phosphorous (P), potassium (K), calcium (Ca), sodium (Na), sulfur (S), iron (Fe), zinc (Zn), copper (Cu), magnesium (Mg), manganese (Mn), and boron (B), and via the Dumas method, nitrogen (N), hereafter collectively referred to as elemental traits.

We assessed relative growth rate (RGR) as a performance metric of the plant (Hoffmann and Poorter, 2002). RGR (g_plant_ g _plant_ ^-1^ day^-1^) was calculated as

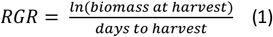

Formula (1) assumes exponential growth during the experiment and uniform size of the seedlings at transplanting to account for temporal differences in reaching R2 stage under stress and non-limiting conditions.

Nitrogen utilization efficiency (NUtE; g_plant_ g_nitrogen_ ^-1^ day^-1^) was subsequently estimated as

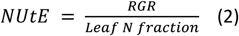

This formula for NUtE assumes that leaf nitrogen content of the MRFEL is a good estimate for whole plant nitrogen content.

Tolerance to nutrient stress was defined by explicitly taking an expected negative relationship between RGR in control conditions and the decrease in RGR under stressed conditions into account. We did this by fitting a linear slope between genotypes RGR in control and their difference in RGR under nutrient limited conditions. The residual of each genotype from this fitted line was then taken as its ExDev-tolerance (Temme *et al*., 2020).

Results were analyzed using R (v4.1.2, R Core Team, 2021) in RStudio (RStudio Team, 2021). For all traits excluding leaf element content (due to lack of replication after pooling), we obtained genotype estimated marginal means (using emmeans v1.7 (Lenth 2021) from a mixed model analysis (using LME4 v1.1-27.1 (Bates *et al*., 2015) taking genotype and treatment as fixed factors and greenhouse bay as a random factor. Wald’s Chi-square test was used to test the significance of the main effects of genotype, treatment, and their interaction. Type 3 Sums of Squares were calculated (using car v3.0-12 (Fox and Weisberg, 2011)) given the presence of a significant interaction between genotype and treatment for many traits. For the pooled leaf biomass, two-sample t-tests were run to test for an effect of treatment on element content across genotypes.

To calculate trait plasticity per genotype, we took the difference in natural log transformed values in control and stressed conditions. This proportional (can be converted to Δ% vs. control via e^Δln(trait)^ −1) metric of plasticity has the additional benefit of being viewpoint agnostic. Only the sign of the natural log difference changes when it is viewed from the control vs. stressed perspective. By using the natural log difference, a halving or doubling in trait value has the same magnitude of plasticity. We determined the level of correlation between mean trait values per genotype separately for trait values under control vs. nutrient limited conditions and for trait plasticities using Spearman correlation in *corrr* v0.4.3 (Kuhn *et al*., 2020).

Given the level of expected covariation among traits, we explored major axes of variation within and across environments using principal component analysis (PCA) using *prcomp* on scaled and centered trait values. To avoid biomass differences masking apparent responses between treatments, and to be able to relate major axes of trait variation and trait plasticity to performance, we selected a set of traits deemed putatively size-independent (i.e., not directly reflecting any aspect of plant biomass) traits. These included chlorophyll content, fine root allocation (mass fraction and root fraction), LMA, LMF, RMF, SMF, and tap root allocation (mass fraction and root fraction). In addition to major axes of variation between treatments, we determined major axes of variation among genotypes within treatments and in trait plasticity for both size-independent traits and element content.

Trait heritability was estimated as both broad (*H*^2^) and narrow sense heritability (*h*^2^). Broad sense heritability was calculated per treatment on those traits with multiple replicates per genotype by fitting a mixed effects model with genotype as a random effect and greenhouse bay as a fixed effect. Subsequently, we calculated *H*^2^ by dividing the genotypic variance by the sum of genotypic variance and residual variance divided by the number of replicates (*H*^2^ =V_g_ /(V_g_ +(V_e_ /n)). Narrow sense heritability was calculated using the R package *heritability* (Kruijer *et al*., 2015) that combines trait data (at either individual and genotypic mean level) with genotypic relatedness (based on pairwise genetic distance calculated using GEMMA (Zhou and Stephens, 2014)). This approach allowed us to estimate *h*^2^ for traits within each treatment and the plasticity between them.

Genome-wide association (GWA) analyses were carried out following Temme et al. (Temme *et al*., 2020). Briefly, a collection of ca. 1.4M high quality SNPs with minor allele frequency > 5% and heterozygosity < 10% (Todesco *et al*., 2020) were clustered into haplotypic blocks based on linkage disequilibrium estimated as *D*’ (Gabriel *et al*., 2002) using PLINK v1.9 (Chang *et al*., 2015). This resulted in 9,179 singleton SNPs and 20,652 co-inherited, multi SNP haplotypic blocks across all 17 chromosomes that were used for the association analyses. Due to possible misordering of SNPs, these numbers are likely an overestimate since “true” haplotypic blocks can be broken up by misplaced SNPs. GWA analyses were then carried out using GEMMA (Zhou and Stephens, 2014) on the full 1.4M snp set with our significance threshold (**α** = 0.05) being adjusted for the number of observed haplotypic blocks (i.e., 0.05/20,652). When different traits had significant SNPs within the same haplotypic block (even if they were not the same SNPs) they were considered to colocalize to the same genomic region. Suggestive SNPs were defined as being in the top 0.01% of all SNPs (without meeting our significance threshold) for a trait and in a region that was significant for at least one other trait. All GWA analyses and visualizations were performed using our custom sunflower GWA pipeline (https://github.com/aatemme/Sunflower-GWAS-v2).

To connect observed instances of trait colocalization with pairwise trait-trait correlation values, we counted the number of significant and suggestive overlaps between all possible pairs of traits in each treatment and their plasticity and related these counts to observed correlation coefficients. Because pairs of traits frequently shared zero regions, we fitted a negative binomial model with |absolute| correlation coefficient as the predictor and the number of shared regions as the dependent variable using *glm*.*nb* from MASS v7.3-54 (Venables and Ripley).

## Results

### More vigorous genotypes experience a greater effect of nutrient limitation

Nutrient limitation generally had a significant impact on growth and development with strong differences between genotypes (Table 1, Fig. S1 for all individual traits). Under low nutrient stress, plants had a reduced developmental rate, delaying the onset of budding (R1 stage) by a median increase in days to reach R1 stage of 11.5%. Bud development was further slowed, resulting in a median increase in days to R2 of 16% (Fig. 1a inset), though genotypes differed widely from a 7% reduction to an 84% increase in time to reach this stage.

**Table 1.** Effect of nutrient stress on sunflower traits. Traits measured in our cultivated sunflower diversity panel with their median value (based on genotype means) and range (in parentheses) when grown under control (i.e., high nutrient) and low nutrient (10% of control) conditions, as well as an estimate of their plasticity (trait adjustment) between treatments. Plasticity was calculated as the difference in natural log transformed values (ln(low nutrients)-ln(control)) but converted here to Δ% vs. control (via e^Δln(trait)^ − 1) for ease of interpretation. Stars indicate significance (Wald’s Chi-square of genotype (G), treatment (T), and their interaction (GxT)). ns: non-significant, *: *P* < 0.05, **: *P* < 0.01, ***: *P* < 0.001.

| Traits | Control | Low nutrients | Plasticity ($\Delta\%$) | T | G | GxT |
| --- | --- | --- | --- | --- | --- | --- |
| **"Size" traits** |  |  |  |  |  |  |
| RGR ($g_{plant} g_{plant}^{-1} day^{-1}$) | 0.09 (0.04 - 0.11) | 0.06 (0.03 - 0.08) | -31.1 (-49.64 - 6.4) | *** | *** | ns |
| Plant mass (g) | 14.95 (2.43 - 61.08) | 8.82 (1.93 - 26.84) | -42.6 (-75.04 - 68.18) | *** | *** | ns |
| Leaf mass (g) | 8.42 (1.57 - 28.11) | 3.42 (1.13 - 7.52) | -60.3 (-81.82 - -3.91) | *** | *** | *** |
| Stem mass (g) | 4.06 (0.36 - 25.81) | 3.53 (0.26 - 16.3) | -14.1 (-71.32 - 272.18) | *** | *** | *** |
| Reproductive tissue mass (g) | 0.17 (0.06 - 0.53) | 0.16 (0.08 - 0.34) | 0 (-56.47 - 137.41) | *** | *** | *** |
| Tap root mass (g) | 1.03 (0.16 - 3.49) | 0.51 (0.12 - 1.98) | -49.2 (-81.25 - 56.18) | *** | *** | ns |
| Lateral root mass (g) | 1.27 (0.23 - 6.26) | 1.14 (0.23 - 4.85) | -14.6 (-67.73 - 407.03) | *** | *** | ns |
| All root mass (g) | 2.26 (0.39 - 8.67) | 1.68 (0.41 - 5.68) | -32.4 (-69.78 - 280.73) | *** | *** | ns |
| All shoot mass (g) | 12.48 (2.04 - 54.07) | 7.03 (1.52 - 22.91) | -45 (-76.27 - 54.39) | *** | *** | *** |
| Height (cm) | 51.94 (17.37 - 118.5) | 62 (15.75 - 126.12) | 14.7 (-39.68 - 152.27) | *** | *** | *** |
| Stem diameter (mm) | 11.87 (6.37 - 19.62) | 8.86 (5.44 - 13.46) | -25.6 (-52.98 - 17.63) | *** | *** | *** |
| Lateral root diameter (mm) | 0.44 (0.3 - 0.67) | 0.41 (0.28 - 0.55) | -8.1 (-37.08 - 42.05) | *** | *** | ns |
| **Development traits** |  |  |  |  |  |  |
| Days to R1 stage | 26 (18.5 - 38.75) | 28.77 (19.43 - 44.5) | 11.5 (-15.55 - 65.52) | *** | *** | *** |
| Days to R2 stage | 30 (19.75 - 42.5) | 35 (19.5 - 49.75) | 16 (-7.09 - 84.26) | *** | *** | *** |
| Days between R1 and R2 | 4.25 (1 - 11) | 6.25 (1 - 13.75) | 41.2 (-30.56 - 200) | *** | *** | *** |
| **Size-independent traits** |  |  |  |  |  |  |
| LMF ($g_{leaves} g_{plant}^{-1}$) | 0.57 (0.47 - 0.69) | 0.39 (0.25 - 0.61) | -31 (-51.53 - -4.63) | *** | *** | *** |
| SMF ($g_{stem} g_{plant}^{-1}$) | 0.26 (0.12 - 0.41) | 0.4 (0.14 - 0.63) | 48.7 (-7.29 - 120.6) | *** | *** | *** |
| RMF ($g_{roots} g_{plant}^{-1}$) | 0.15 (0.09 - 0.23) | 0.19 (0.11 - 0.28) | 21.4 (-28.45 - 132.45) | *** | *** | ns |
| Root shoot ratio ($g_{roots} g_{shoot}^{-1}$) | 0.18 (0.1 - 0.55) | 0.23 (0.13 - 0.57) | 27.6 (-54.05 - 222.89) | *** | *** | ns |
| Tap root MF ($g_{tap root} g_{plant}^{-1}$) | 0.07 (0.03 - 0.15) | 0.06 (0.03 - 0.13) | -13.3 (-58.65 - 57.62) | *** | *** | ns |
| Fine root MF ($g_{fine root} g_{plant}^{-1}$) | 0.08 (0.04 - 0.15) | 0.13 (0.07 - 0.22) | 49.2 (-15.57 - 255.51) | *** | *** | * |
| Tap root RF ($g_{tap root} g_{roots}^{-1}$) | 0.46 (0.21 - 0.74) | 0.32 (0.14 - 0.6) | -27.4 (-65.36 - 17.29) | *** | *** | * |
| Fine root RF ($g_{fine root} g_{roots}^{-1}$) | 0.54 (0.26 - 0.79) | 0.68 (0.4 - 0.86) | 22.8 (-11.76 - 91.94) | *** | *** | * |
| LMA ($g m^{-2}$) | 36.1 (26.05 - 59.43) | 37.17 (26.05 - 57.2) | 2 (-22.84 - 41.97) | *** | *** | *** |
| Leaf Area Ratio ($m^2 g_{plant}^{-1}$) | 0.02 (0.01 - 0.03) | 0.01 (0 - 0.02) | -32.9 (-64.51 - 15.19) | *** | *** | ** |
| Specific Stem Length ($cm g^{-1}$) | 14.72 (4.7 - 49.39) | 18.22 (6.72 - 62.47) | 30.4 (-61.34 - 194.28) | *** | *** | *** |
| Specific Root Length ($m g^{-1}$) | 9.64 (4.71 - 19.45) | 10.55 (5.98 - 20.29) | 9 (-45.05 - 207.29) | *** | *** | ns |
| Root branching frequency ($mm^{-1}$) | 0.47 (0.27 - 0.74) | 0.5 (0.3 - 0.75) | 5 (-45.71 - 110.83) | *** | *** | * |
| Chlorophyll content (index) | 22.39 (12.39 - 36.85) | 11.01 (6.95 - 18.23) | -50.2 (-73.6 - -26.55) | *** | *** | *** |

**Figure 1.**
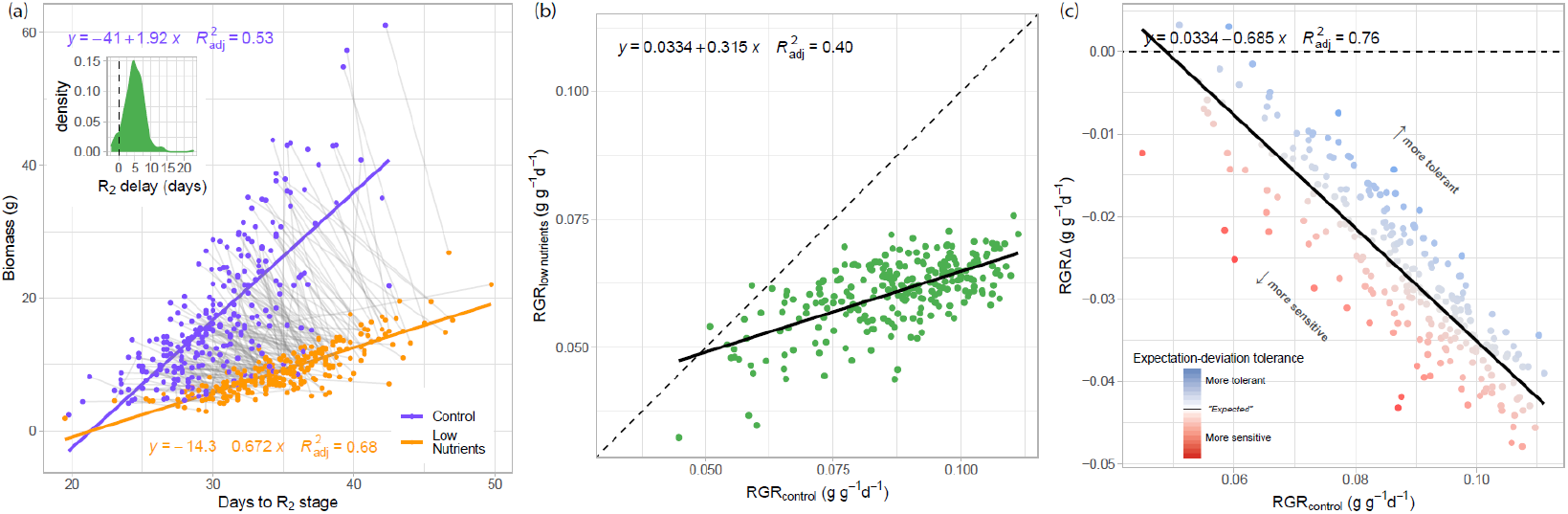
Effect of nutrient limitation on development time, relative growth rate (RGR) and the effect of stress. Plants under low nutrient stress had reduced biomass, RGR, and took longer to reach the R2 stage **(a)** Relationship between days to reach R2 (early budding) stage and biomass at that time under control conditions (purple) and nutrient limitation treatment (orange). Genotypes (colored dots) are connected between treatments with gray lines. Generally nutrient limitation delayed R2 stage. Inset panel shows the distribution of the delay in reaching the R2 stage. **(b)** Relationship between RGR under control conditions and RGR under nutrient limitation across all genotypes (green dots). The dotted line shows the 1:1 relationship. **(c)** Relative growth rate (RGR) in control vs the difference in RGR in nutrient limited conditions. Genotypes colored by their residual from the fitted line. This ExDev-tolerance metric shows more sensitive (red) and more tolerant (blue) genotypes. Equations describe the best-fit line with the R^2^ _adj_ of that model.

Despite the overall increase in time to reach R2, biomass was reduced by a median of 47% at R2 (Table 1, Fig. 1a). As genotypes developed, within both control and stressed conditions, those that took longer to reach R2 tended to have accumulated more biomass, though this effect was diminished under nutrient stress (Fig. 1a). This slower development time and reduced biomass accumulation due to low nutrient availability resulted in reduced relative growth rate (RGR) (Table 1, Fig. 1b). Genotypes with a higher RGR in control conditions (i.e., higher vigor) tended to have a higher RGR under low nutrient stress. Differences in the treatment effect on RGR across genotypes did not rise to the level of significance as indicated by the lack of GxT interaction on RGR (Table 1). This lack of a significant interaction could be the result of substantial within genotype variation. However, when investigated across genotype means, our results do suggest that genotypes with overall higher vigor showed larger differences in growth between control and nutrient stress, indicating a possible trade-off between vigor and the effect of stress (Fig 1b, 1c).

Our results show that when viewed at the mean genotype level, genotypes with higher RGR under control conditions (i.e., more vigorous genotypes) had a greater reduction in RGR under low nutrients (*P* < 0.001, Fig. 1c), relating tolerance to only the difference in RGR due to stress runs the risk of confounding tolerance with low vigor. By fitting a linear relationship between the difference in RGR and and vigor for each genotype, performance relative to the overall expectation can be estimated as the residual from this fitted line. We define this residual as the ExDev-tolerance of each genotype, allowing us to score them as being more or less tolerant than expected based on their vigor.

### Expectation-deviation tolerance is positively correlated with the change in nitrogen utilization efficiency

Surprisingly, genotypes with a higher leaf nitrogen content at harvest (R2 stage) had a lower RGR, both under control and low nutrient conditions (Fig 2a inset). Given that RGR and leaf nitrogen content both changed in response to low nutrient availability, we used growth rate per unit leaf nitrogen as a metric for nitrogen utilization efficiency (NUtE). Genotypes with the highest NUtE in control tended to remain the highest under nutrient stress (Fig. 2a). Moreover, nearly all genotypes increased NUtE, indicated by positive ΔNUtE, though the extent of this varied among genotypes (Fig. 2b). Strikingly, there was a strong positive relationship between the increase in NUtE and ExDev-tolerance (Fig 2c). Genotypes that exhibited greater than expected increases in NUtE (independent of their actual NUtE), tended to have a higher ExDev-tolerance (i.e., a smaller reduction in RGR than expected) suggesting a possible role for NUtE in low nutrient tolerance.

**Figure 2.**
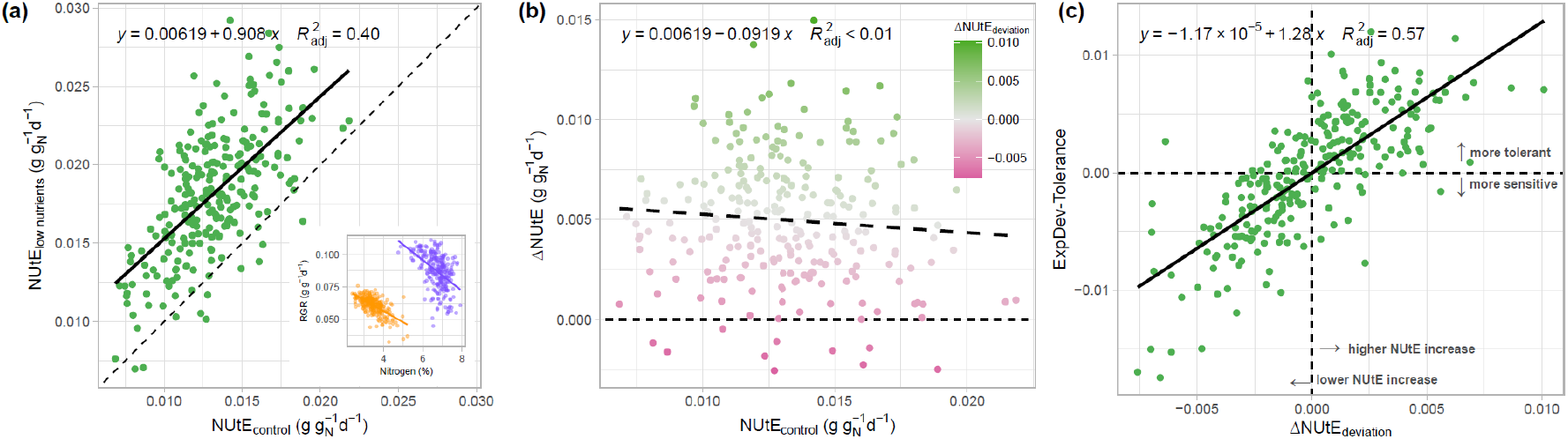
Nitrogen utilization efficiency (NUtE) and its relation to nutrient stress tolerance. **(a)** Relationship between NUtE [estimated as growth rate per unit leaf nitrogen (g_biomass_ g_N_ ^-1^ d^-1^)] in control conditions and NUtE in low limited conditions. The dotted line shows the 1:1 relationship. Inset shows the relationship between leaf N content and RGR, used to calculate NUtE (control: purple, low nutrients: orange) **(b)** Relationship between NUtE in control conditions and the change in NUtE from control to low nutrient conditions. **(c)** Relationship between the deviation from the mean NUtE increase and ExDev-tolerance.

### Nutrient stress has large impacts on diverse plant traits and leaf elemental content

In the low nutrient treatment, more biomass was allocated to roots leading to median 21.4% increased root mass fraction (RMF) as compared to control; this was primarily driven by an increase in fine root mass fraction. In addition to roots, more biomass was allocated to stem tissue as well (Table 1). Low nutrient stress also resulted in alterations in plant morphology where the MRFEL became smaller and thicker or denser (i.e., increased leaf mass per area), roots became thinner or less dense (i.e., increased specific root length), and stems became thinner despite an increase in height (i.e., increased specific stem length). Chlorophyll content of the leaf was drastically reduced, median −50%, tracking the large effect of low nutrient availability on leaf elemental composition (Table 1, Figure S1).

With the exception of sodium and manganese, nutrient stress generally had large and significant effects on leaf element content (Table 2, Fig. S1). Leaf nitrogen decreased by 49%, phosphorus by 36%, and potassium by 15%. Calcium content increased dramatically with a 123% increase under low nutrient conditions vs. control, although it should be noted that we supplied additional calcium as gypsum and lime and as such calcium was abundantly available in both treatments. Additionally, magnesium content increased by 32%. However, while the concentration of calcium and magnesium increased, the total amount in the leaf remained largely the same for calcium and was lower for magnesium (Fig. S2) due to a concomitant decrease in leaf mass.

**Table 2.**
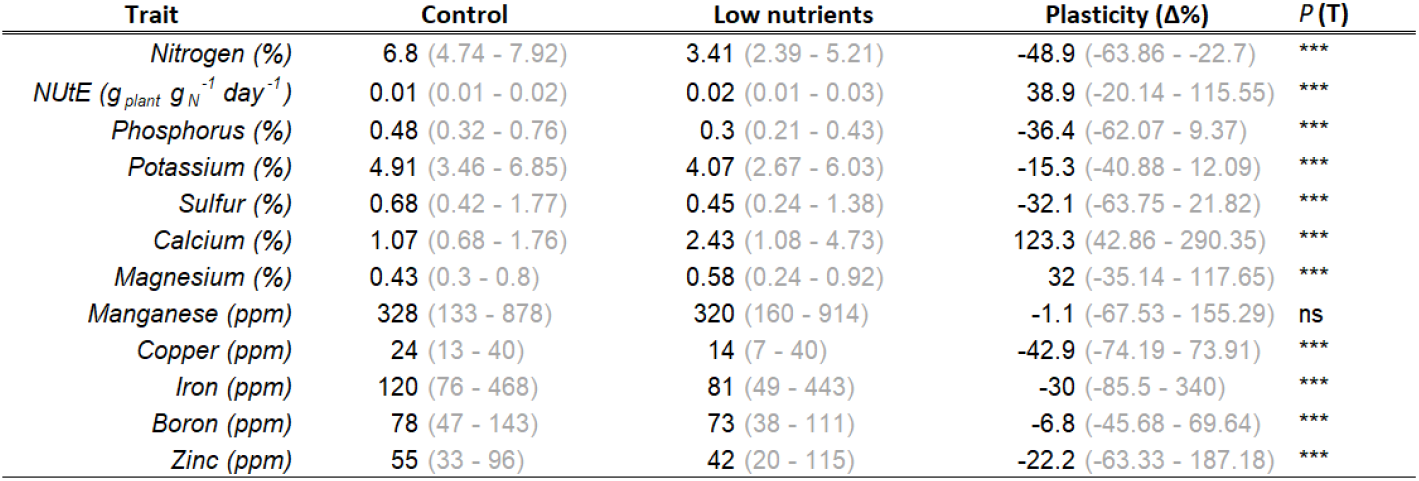
Effect of nutrient stress on sunflower leaf elemental content. Leaf element content of the most recent fully expanded leaf measured in bulked samples on our sunflower diversity panel. Values shown are the median content across genotypes (with range in parentheses) when grown under control (i.e., high nutrient) and nutrient limited (10% of high nutrients) conditions, as well as the plasticity (as Δ%) between them. Plasticity was calculated as the difference in natural log transformed values (ln(low nutrients)-ln(control)) but converted here to Δ% vs. control (via e^Δln(trait)^ − 1) for ease of interpretation. Element content is shown on a mass basis (g g^-1^). Since individual leaves per genotype were bulked, stars indicate significance (based on a *t*-test) of nutrient limitation treatment. ns: non-significant, *: *P* < 0.05, **: *P* 0.01, ***: *P* < 0.001.

### Major axes of trait variation/plasticity correlate with expectation-deviation tolerance and vigor

To investigate trait relationships in the context of nutrient stress tolerance, and to determine if they differ from traits related to high vigor, we sought to determine the correspondence between tolerance, performance, and variation in broad multivariate suites of traits. Across both treatments, >60% of the variation in size-independent traits (Fig 3a) and >60% of the variation in leaf elemental content (Fig 3b) could be captured in the first two principal components. Differences between treatments largely followed the first principal component with a shift towards below ground resource allocation (i.e., decreased chlorophyll, LMF, and LAR, increased RMF and investment in fine roots). Leaf element content was highly correlated between treatments with reduced levels of N, P, K, S, and Cu coupled with increased levels of Ca and Mg under low nutrient availability.

**Figure 3.**
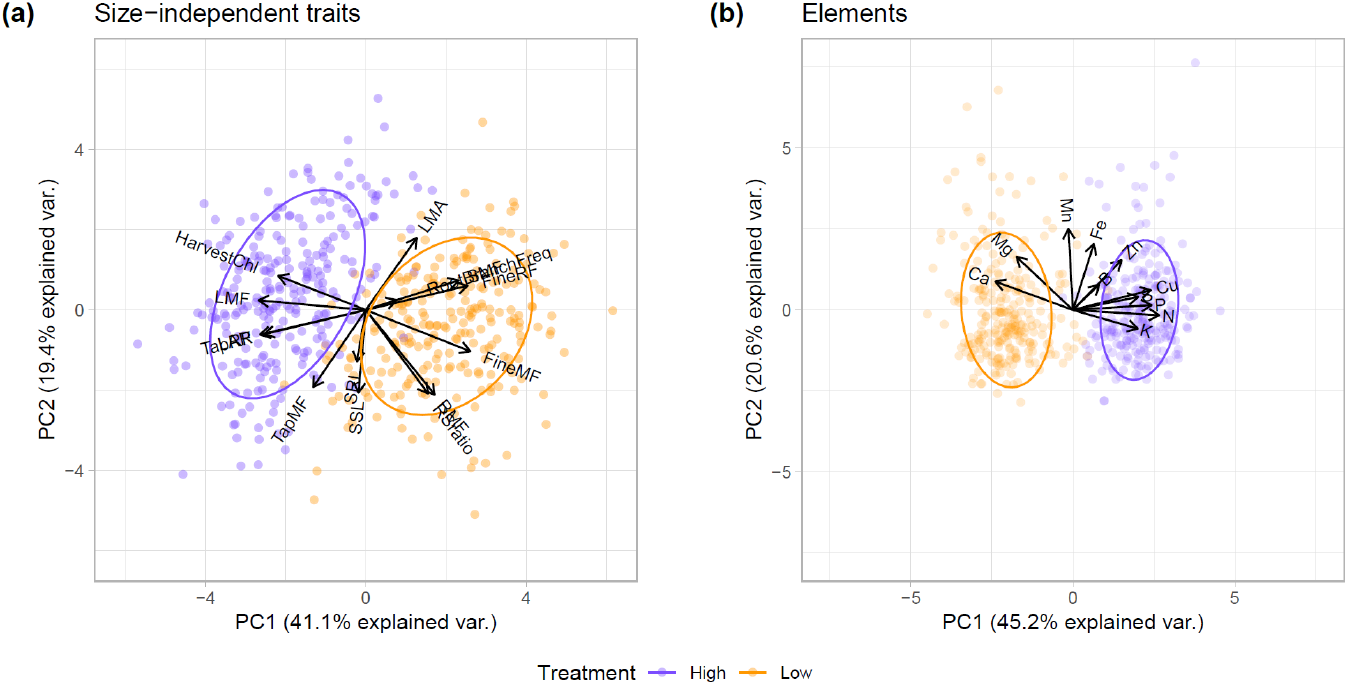
Multivariate trait shifts in response to low nutrient availability in sunflower. Principal component analysis (PCA) reveals correlated trait adjustments to low nutrient stress for **(a)** putatively size independent traits (chlorophyll content, fine root allocation [mass fraction and root fraction], LMA, LMF, RMF, SMF, tap root allocation [mass fraction and root fraction]), SRL, and root branching frequency, and **(b**) leaf elemental traits (N, P, K, S, Ca, Mg, Mn, Cu, Fe, B, Zn) in control (high nutrient) treatment (purple) and low nutrient treatment (orange).

To connect trait variation to differences in performance and tolerance, we adopted a within-treatment and plasticity-between-treatments approach. We related major axes of trait variation in (putatively) size-independent traits and leaf elemental traits under control and low nutrient conditions, along with trait plasticity between treatments to RGR and ExDev-tolerance. We found that the first principal component of size-independent traits under control and low nutrient conditions was closely correlated with RGR in those environments (Table 3). ExDev-tolerance was significantly correlated with the first two principal components of the plasticity in size independent traits, but the relationship with PC1 was more robust (Table 3, Fig 4b). Multivariate leaf elemental content was likewise correlated with RGR as well as ExDev-tolerance. More specifically, variation in PC2 of leaf elemental content was associated with RGR and, to a lesser extent, ExDev-tolerance (Fig 4d).

**Table 3.** Relationship between sunflower relative growth rate and ExDev-tolerance with correlated suites of size-independent and leaf ionomic traits. *R*^2^ and significance of the ordinary least-squares regression of the PC1 and PC2 values of the putatively size-independent traits (chlorophyll content, fine root allocation [mass fraction and root fraction], LMA, LMF, RMF, SMF, tap root allocation [mass fraction and root fraction]) and elemental traits (B, Ca, Cu, Fe, Mg, Mn, N, P, K, S, Zn) under control (high nutrients) and low nutrient (10% of control) conditions, as well as the plasticity in trait values between treatments. Highlighted in bold are the model fits RGR and ExDev-tolerance against the principal components with the highest explanatory power for vigor and tolerance for each trait. *: *P* < 0.05, **: *P* < 0.01, ***: *P* < 0.001.

| Trait suite | Treatment | PCA | RGR | RGR Low | ExDev- |
| --- | --- | --- | --- | --- | --- |
|  |  |  | Control | Nutrients | tolerance |
| Size-independent | Control | PC1 | **0.51\*\*\*** | 0.2*** |  |
|  |  | PC2 |  | 0.02* |  |
|  | Low nutrients | PC1 | 0.33*** | **0.45\*\*\*** | 0.14*** |
|  |  | PC2 |  |  |  |
|  | Plasticity | PC1 |  | 0.11*** | **0.23\*\*\*** |
|  |  | PC2 | 0.06*** |  | 0.02* |
| Elements | Control | PC1 | 0.07*** |  |  |
|  |  | PC2 | 0.15*** | 0.08*** |  |
|  | Low nutrients | PC1 | 0.05*** | 0.03** |  |
|  |  | PC2 | 0.22*** | **0.32\*\*\*** | **0.12\*\*\*** |
|  | Plasticity | PC1 |  | 0.02* |  |
|  |  | PC2 | 0.07*** | 0.16*** | 0.09*** |

**Figure 4.**
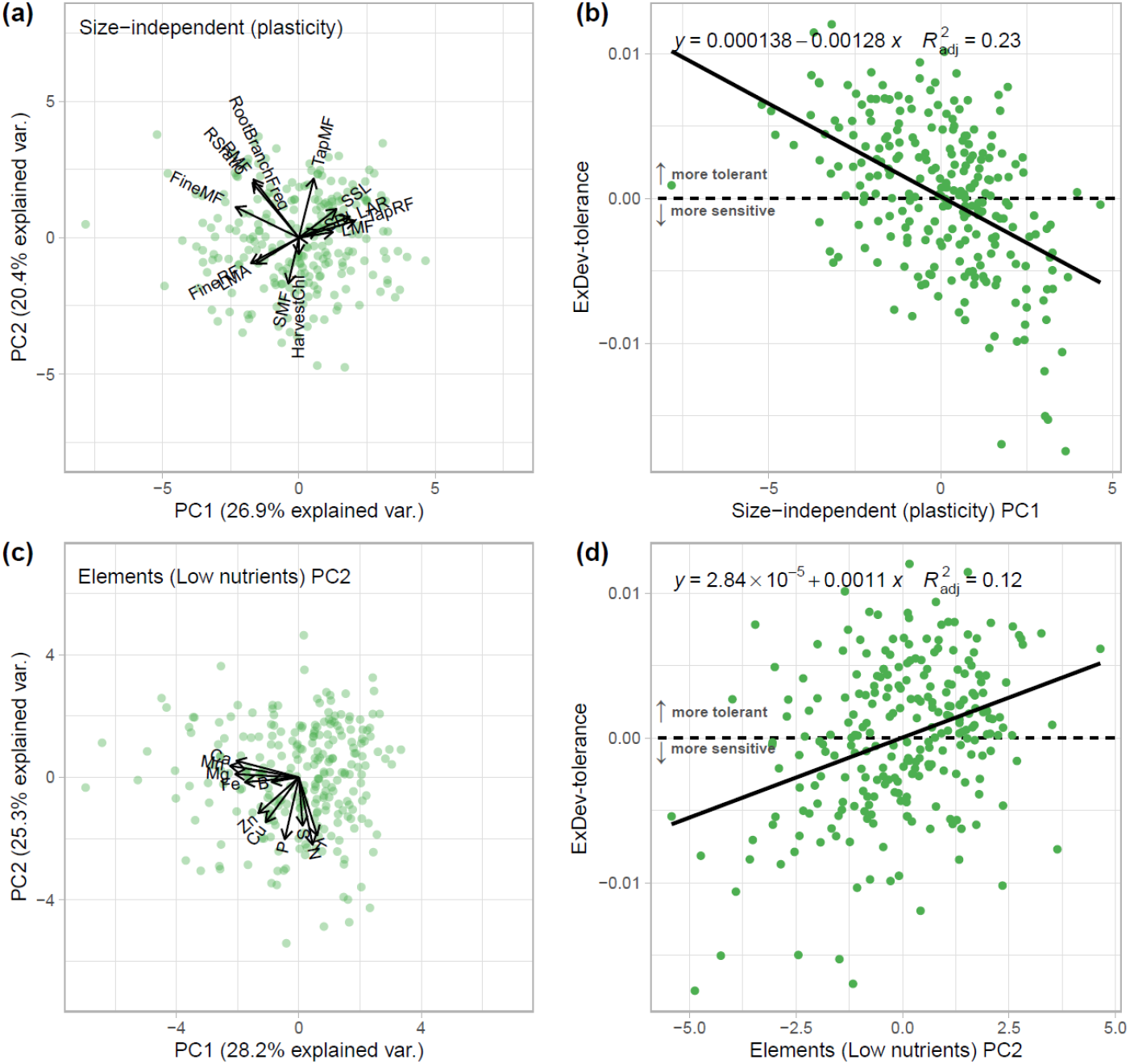
Major axes of trait plasticity and leaf elemental content are correlated with expectation-deviation tolerance. **(a)** Principal component analysis (PCA) of the plasticity in (putatively) size-independent traits in response to nutrient limitation. **(b)** Relationship between genotypes (green points) loading on PC1 of the plasticity in size-independent traits and genotypic nutrient stress ExDev-tolerance. **(c)** PCA of leaf elemental content under low nutrient availability. **(d)** Relationship between genotypes (green points) loading on PC2 of leaf elemental content in low nutrients and genotypes nutrient stress tolerance.

Genotypes with a suite of size-independent trait adjustments related to a greater decrease in LAR and a greater increase in the fraction of biomass allocated to fine roots (at the expense of tap root) were more tolerant (Table S1, Fig. 4a-b). In terms of elemental content, genotypes that had higher leaf N, P, and K content had higher ExDev-tolerance (Table S1, Fig. 4c-d). For a full picture of trait loadings onto principal components, see Table S1.

### Multiple genomic regions associated with trait variation and plasticity to nutrient stress

Variation of all measured traits (Table 4) in control and low nutrients, and trait plasticity between treatments, could be associated with 215 unique, putatively independent genomic regions (based on linkage disequilibrium; Fig. S5). On average, we found 2.3 significantly associated regions per trait (Table 4), though this number varied widely and tended to be lower for trait plasticity across treatments as compared to trait values within treatments. Our finding of > 20 distinct regions for sulfur content in close proximity to each other (Fig. S4) may reflect localized misordering in the genome assembly; in reality, these regions may very well be grouped together. Thus, our finding of 215 independent regions should be viewed as an overestimate, likely there are fewer major effect regions involved in variation in these traits, though there are likely other regions with effects below the level of detectability as well.

**Table 4.**
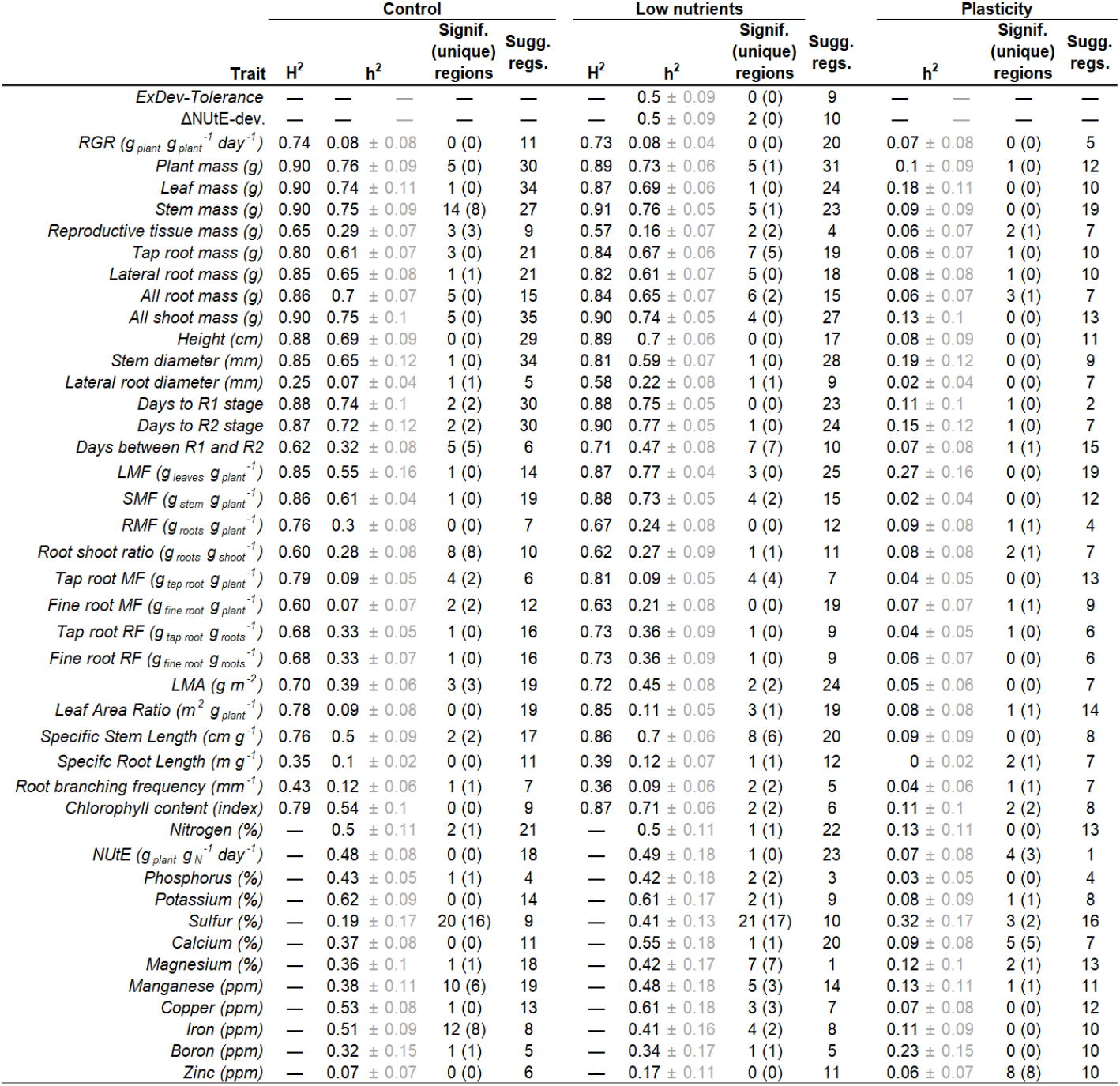
Genomic associations and trait heritability at control, low nutrients, and plasticity. Broad sense (H^2^) and narrow sense (h^2^) heritability of all measured traits at control conditions, under low nutrient stress and the plasticity between treatments. For traits where only genotype means were available only h^2^ could be calculated. Additionally the number of significant (Signif.) genomic regions per trait (with those unique to that trait/treatment combination in parentheses) and the number of suggestive regions (Sugg. Regs.), SNPs in that region in the top 0.01% of association strength and significant for at least one other trait.

In terms of trait colocalization, 51 regions (across 13 of the 17 chromosomes, Fig. S7) were significantly associated with multiple trait/environment combinations, though this number rises to 178 regions if we include suggestive associations (i.e., regions having SNPs in the top 0.01% of *P*-values and significant for at least one other trait) (Table S3, Fig. S6). The region associating with variation in the largest number of traits was region 17-09, which was significantly associated with 9 traits (related to aspects of root mass, S & Mn content, and development time) and a diverse set of 54 suggestive traits. Unfortunately, this region was quite large (>50Mb) and contained 699 genes, making it difficult to distinguish between the pleiotropic effects of a single locus and close linkage of multiple functional variants. In contrast, region 03-05 was significantly associated with eight biomass-related traits and contained only seven genes. Unfortunately only three of the seven genes had know or putative functions (*Putative EH domain, EF-hand domain pair protein, Putative protein kinase TKL-CTR1-DRK-2 family, Putative Late embryogenesis abundant protein, LEA_2 subgroup*), highlighting the difficulty of determining candidate genes from GWAS results alone.

As one might expect, at the genomic level, we saw a reflection of the correlated nature of variation in traits (Fig. 5). Strongly positively or negatively correlated traits tended to share a greater number of genomic regions with significant and/or suggestive SNPs associated with those traits. Indeed, in both treatments we found a significant relationship between bivariate trait correlation strength and the number of shared genomic regions associating with those traits. While the shape of this relationship was comparable between control (high nutrients) and low nutrient treatment, highly correlated trait plasticities tended to share a lower number of genomic regions. (Fig. 5d)

**Figure 5.**
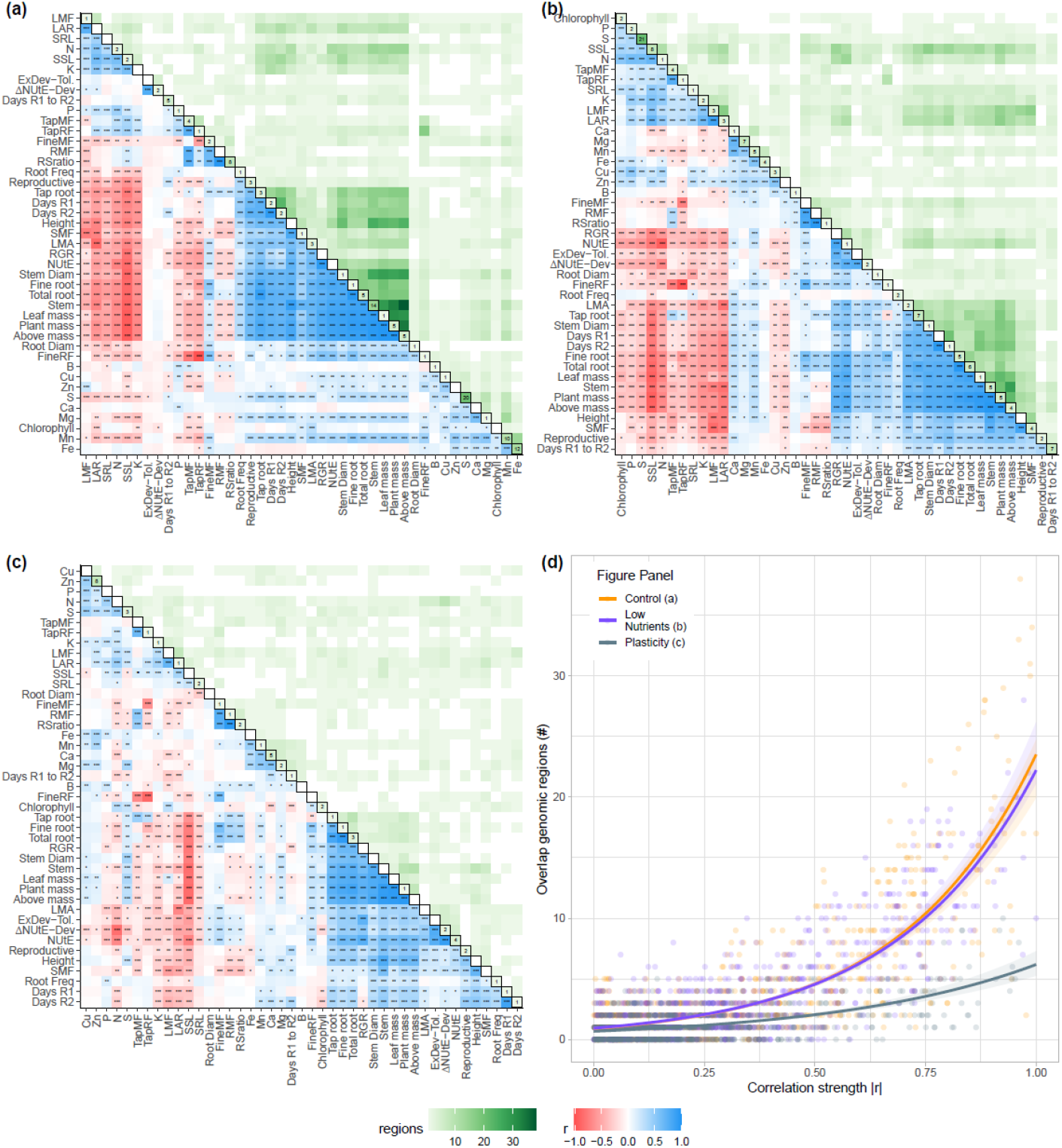
Sunflower trait correlations and genomic colocalization. **(a)** Spearman correlation matrix of phenotypic and elemental traits in our cultivated sunflower diversity panel under control (control conditions; lower diagonal) and the number of overlapping genomic regions (at the suggestive level, upper diagonal). The total number of significant regions listed per trait on the diagonal. Traits are ordered by hierarchical clustering in each panel so that closely correlated traits are close together. **(b)** Correlation of traits and overlapping genomic regions under low nutrient conditions. **(c)** Correlation and overlap in genomic regions for trait plasticity values between treatments (ln(control)-ln(stress)). **(d)** Logistic regression of absolute pairwise trait-trait correlation coefficient and the number of overlapping genomic regions. Correlation values range from −1 (red) to 1 (Blue). Stars in tiles indicate the significance of correlations. *: *P* < 0.05, **: *P* < 0.01, ***: *P* < 0.001

We found no significant genomic regions directly underlying nutrient stress tolerance (i.e., having a lower than expected, based on vigor, reduction in RGR). However, we did find eight regions with suggestive associations for tolerance that were significant for key traits identified by our principal component analyses (e.g., root mass allocation and nitrogen utilization efficiency [NUtE]; Table 3, Fig. 5). For the genotypic deviations from expected (average) NUtE increase (Fig. 3b), a key trait involved in tolerance in this study (Fig. 3c), we found two significant genomic regions (Fig. 6a). These regions on chromosome 1 (01-01) and chromosome 17 (17-13) contained 176 genes and 8 genes respectively.

**Figure 6.**
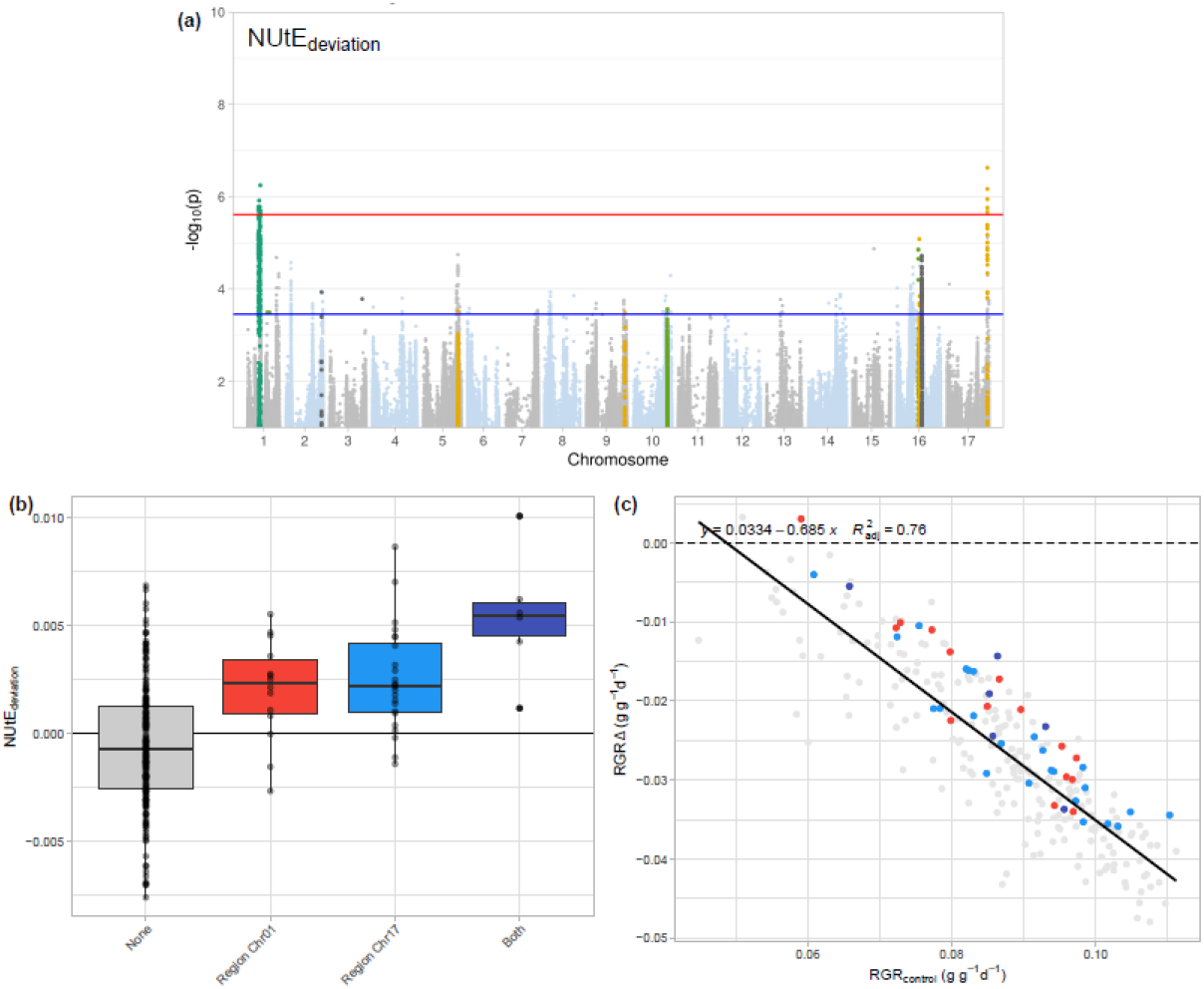
Regions associated with the change in NUtE and their suggestive effect on ExpDev-tolerance. **(a)** Manhattan plot of the deviation from expected increase in NUtE across genotypes. Two haplotypic regions, 01-01 and 17-29, are significantly associated with this trait (SNPs above the red line). Several other regions show suggestive associations that colocalize with other traits (highlighted SNPs above the blue line). **(b)** Deviation in NUtE increase under low nutrient conditions (negative values = smaller increase than expected, positive values = larger increase than expected) and minor allele status for regions 01-01 and 17-29 (the most significant SNP within each haplotypic region was selected as the tag SNP for that region). Genotypes with the minor allele in either region tend to be more tolerant with an ostensibly additive effect for the few genotypes that have both minor alleles. **(c)** Visualization of the genotypes carrying the minor allele for regions 01-01 and 17-29 (colors correspond to those used in panel [b]) and where they fall on a plot of expectation-deviation tolerance.

Focusing on these latter two regions, we found that genotypes that carry the minor allele in both regions (generally homozygous due to our filtering) tended to have a higher NUtE increase under low nutrient conditions as compared to those carrying the major allele (Fig. 6a). Interestingly, the six genotypes with the minor alleles of both regions had (on average) an even further improved NUtE suggesting an additive effect for these regions (Fig. 6b). While it should be noted that these regions were not significantly associated with ExDev-tolerance, when plotted on the relationship between RGR in control and the decline in RGR (similar to Fig. 1c), it can be seen that the genotypes that carry the minor allele for either or both regions tend towards being more tolerant than would be expected based on their vigor (Fig. 6c).

## Discussion

Improving our understanding of crop nutrient stress tolerance will aid in developing varieties capable of high performance on marginal lands and/or with reduced inputs (Good *et al*., 2004). Here, we sought to determine the traits and genomic regions involved in low nutrient stress tolerance in cultivated sunflower, one the world’s most important oilseed crops.

### The greater the vigor the harder the fall

Similar to prior findings for salt stress (Tran *et al*., 2020; Temme *et al*., 2020), plants with higher vigor (high RGR) under control conditions tended to have the best overall performance (i.e., higher RGR) under nutrient limited conditions. However, these same genotypes tended to exhibit a greater reduction in RGR under nutrient limited conditions. Thus, genotypes with high performance under nutrient limitation vs. those with a low effect of nutrient limitation on performance represent largely non-overlapping sets, making it difficult to identify ‘tolerant’ on the basis of performance *per se*. To address this challenge, our definition of tolerance accounts for the negative relationship between vigor and the decrease in RGR due to nutrient limitation. In doing so, genotypes are scored by their deviation from their expected reduction in growth, given their vigor. This expectation-deviation tolerance (ExDev-tolerance) exhibited moderate heritability across genotypes, showing the potential for improving genotypes by focusing on this tolerance metric, though we did not identify any specific genomic regions associated with this trait,

Low nutrient stress slowed sunflower’s rate of development, with nutrient limitation exhibiting a particularly strong effect on the number of days to reach R2 (early bud development). More specifically, we found an average 11.5% increase in the time to reach R1 (onset of bud development) and a 40% increase in the time to reach the following R2 (Table 1). In an agricultural setting, such a slowdown in development could result in plants running into unfavorable environmental conditions (e.g., later season heat, drought) (Kazan and Lyons, 2016). Interestingly, this slowdown in development was highly heterogeneous across genotypes, with a small fraction exhibiting more rapid development (up to 15.5% shorter time to R1) while others extended their development time by as much as 65.5%. The low heritability of genotypic plasticity in developmental time and the lack of overlap in genomic regions associated with developmental rate traits in both environments suggests difficulty in optimizing a cultivar for a range of environments. However, given the high heritability of development time (Table 4) within environments, it may be possible to optimize cultivars for particular environments.

### Higher nitrogen utilization efficiency is associated with greater expectation deviation tolerance

Relating growth to nitrogen uptake is of particular interest due to the essential role nitrogen plays in multiple physiological processes, including the conversion of CO2 into biomass. Improving a plant’s capacity to acquire and use nitrogen more efficiently may be the key to improving performance in poor nutrient conditions (Han *et al*., 2015; Tegeder and Masclaux-Daubresse, 2018; Swarbreck *et al*., 2019). Similar to results from ryegrass (Zhao *et al*., 2020), sunflower was found to generally increase nitrogen utilization efficiency under nutrient limitation (Fig 2a). Moreover, contrary to RGR, the magnitude of this increase was unrelated to NUtE under benign conditions. However, we did find substantial genotypic variation in the magnitude of this NUtE increase (Fig 2b).

While our measures of growth (RGR) and NUtE are not wholly independent, relating our measure of ExDev-tolerance to genotypes change in NUtE highlights the importance of NUtE in determining the response to low nutrient stress (Xu *et al*., 2012; Han *et al*., 2015). Indeed, genotypes that exhibit greater increases in NUtE under stress tend to exhibit greater ExDev-tolerance. Thus, improving NUtE could be a strategy for improving nutrient tolerance in sunflower without reducting vigor. For NUtE in control and nutrient stressed conditions and genotypes deviation from the overall NUtE increase, we found moderate narrow sense heritability and several genomic regions were associated with variation in NUtE under low nutrient stress and plasticity in NUtE (Table 4). Interestingly, for the deviation in NUtE, we found two regions with ostensibly additive effects (Fig 6), showing the potential for trait optimization via selection.

### Vigor and tolerance are correlated with distinct multivariate suites of traits and trait plasticity

Similar to other species (Weih *et al*., 2018; Meyer *et al*., 2019), low nutrient stress leads to a host of trait changes in cultivated sunflower, with broad multivariate changes in trait expression across environments (Fig 3). Over 50% of the variation in size-independent traits and leaf elemental content across control and low nutrient stress were captured by the first two principal components in each case. Similarly, variation in traits within treatments and plasticity between treatments, could be simplified in a limited number of major axes of variation (Fig S3). These strong multivariate axes of covariation illustrate the difficulty in isolating changes in individual traits to produce novel trait combinations (Walsh and Blows, 2009).

With strong covariation among trait variation within treatments as well as plasticity between them, relating single traits to tolerance would be a gross oversimplification. Rather, we related the major axes of variation in size-independent traits and leaf element content to expectation-deviation tolerance and RGR. In both, control treatment and low nutrient stress treatment RGR was correlated with a suite of traits related to carbon uptake (LAR, LMF, SSL). ExDev-tolerance was, however, correlated with a suite of trait plasticities with genotypes exhibiting a greater decrease in LAR and a greater increase in the fraction of biomass allocated to fine roots at the expense of tap root being more tolerant overall (Table S1, Fig. 4a-b). Results from cotton show this same connection between root system architecture and nitrate uptake efficiency (Iqbal *et al*., 2020). Moreover, comparable to our findings for salt stress (Temme *et al*., 2020), traits related to ED-tolerance differ from those related to vigor indicating that in principle it could be possible to combine high tolerance with high vigor.

### Multiple genomic regions impact trait variation and plasticity

A large number of genomic regions were involved in trait variation under control and low nutrient conditions. Interestingly, similar to findings in maize we find additional genomic regions associated with trait plasticity (Kusmec *et al*., 2017; Gage *et al*., 2017), often distinct from those involved in variation in either treatment (Fig S4). The often large size of regions identified, containing many genes, makes functional inferences at the level of individual genes difficult (Table S4). However, some interesting trends and key genes could be found in the regions of interest.

For developmental rate, days to reach R2 (i.e., bud formation) was significantly associated with region 04_01 under control conditions. This region of two genes contains a “Transcription factor interactor and regulator AUX-IAA family” gene, suggesting a specific mechanism related to auxin transport involved in development rate (Sauer *et al*., 2013). Belowground, region 08_14, significant for plasticity in RMF and containing 14 genes, included a “Transcription factor MYB-HB-like family” gene. This family of transcription factors is known to be involved in several processes including response to stress and development (Ambawat *et al*., 2013). Root morphology, as reflected in root branching frequency, was associated with two regions (02_03 and 02_06) under low nutrient conditions. These regions contained 51 and 35 genes respectively with both harboring a putative gene in the RLK/Pelle family of kinases. This family of kinases is involved in a host of processes including development (Gish and Clark, 2011). The *Putative protein kinase RLK-Pelle-CrRLK1L-1 family* in region 02_06 is of a type linked to cell expansion (Nissen *et al*., 2016), suggesting a mechanism for altered branching frequency. Two key genomic regions on chromosomes 1 and 17 could be directly linked to NUtE increases with ostensibly additive effects. Region 17-14 (8 genes) contained a “*LAZY1*” gene, which is known to be involved in auxin transport (Dong *et al*., 2013). However, extreme care should be taken in overinterpreting these results as far more genes of unknown function are also contained in these significant regions and due to LD any of these could also be a causal variant.

In comparing overlaps in associated genomic regions among traits we found that traits that tended to covary more strongly tended to share a higher number of genomic regions associated with those traits (Fig 5d). In interpreting this result care should be taken since some of these correlations could be due to the fact that trait pairs could be compound traits that share underlying physiological processes or have mathematical dependencies. This connection between phenotypes/traits, also called trait integration or the study of an organism’s multivariate phenotype is an ongoing avenue of research (Pigliucci and Preston, 2004; Messier *et al*., 2017). Experiments have shown that trait adjustments to the environment rarely happen in isolation, and that through understanding a genotype’s integrated phenotype, responses to the environments can be better understood (Richards *et al*., 2006). Due to pleiotropy, either based in close linkage or shared genetic pathways, selecting on one trait or trait plasticity has the potential to impact many others (Wagner and Zhang, 2011; Svensson *et al*., 2021; Mural *et al*., 2021).

## Conclusion

Different abiotic stresses (e.g., drought, salinity, or low nutrient stress) exhibit different modes of action and tolerance to these stresses is thus conferred by different sets of traits. However, here, in cultivated sunflower, we find overall responses to low nutrient stress that reflect results from prior work on salinity stress (Temme *et al*., 2020). Across genotypes, those with the highest vigor (i.e., growth in benign conditions) tend to remain the best performers under low nutrient stress. However, these same high vigor genotypes also suffer the most under low nutrient conditions (i.e., they exhibit the greatest decrease in RGR). In defining tolerance, we therefore took this vigor/stress effect relationship into account. Genotypes with a smaller decrease than expected based on their vigor are thus viewed as being more stress tolerant, and vice versa. For nutrient stress, we found that this expectation-deviation tolerance metric was positively correlated with nitrogen utilization efficiency. More specifically, those genotypes that exhibited an above average increase in NUtE are those that have a high expectation-deviation tolerance. In addition to NUtE, we found that ExDev-tolerance was to a suite of multivariate trait plasticities where genotypes that exhibit a greater decrease in LAR, a greater increase in the fraction of biomass allocated to fine roots, and less to tap root were more tolerant of low nutrient conditions. Numerous genomic regions were found to be associated with trait variation and plasticity. While we found many regions associated with variation in multiple traits, unique regions for traits were found as well. Thus, while there are generally more regions involved in variation in multiple traits, observed instances of genomic regions affecting only individual traits in this experiment leaves open the possibility of genetically decoupling certain trait combinations in the interest of exploring novel phenotypic space. Genotypic variation in ExDev-tolerance, a close tie between ExDev-tolerance and NUtE, multivariate suites of traits correlated with ExDev-tolerance, and a host of potential genomic targets show the potential enhancing low nutrient stress tolerance in cultivated sunflower.

## Acknowledgements

We thank K. Bettinger, M. Boyd, K. Tarner, UGA greenhouse staff, the sunflower undergraduate army, and numerous past and present members of the Donovan and Burke labs for their help during the experiment. Special thanks to the reviewers for their valuable feedback. This work was supported by a grant from the NSF Plant Genome Research Program (IOS-1444522) to JMB and LAD.

## Supplementary materials

Table S1 **Trait loadings on PC axes (companion to table 3, figure 4)** Loadings (fraction of variance in trait explained by specific principal component) based on trait variation in our cultivated sunflower diversity panel for the first and second PCs under control and low nutrient stressed conditions as well as the plasticity in trait values between treatments. Highlighted are the top three size-independent traits/elements per PC with the rank of the loading in parentheses.

Table S2 **Significant and suggestive traits per genomic region**. List of significant genomic regions and associated significant/suggestive traits. Genomic regions of interest along with the number of significant (GEMMA GWAS algorithm Wald test −log10(P)>5.6) and suggestive traits (SNPs in top 0.1% of *P*-values for trait and significant for at least one other trait) and number of genes in each region. Regions significant or suggestive for tolerance are bolded in the list of traits.

Table S4 **Genes found in significant regions** As detailed in the manuscript text, gene names were extracted from the annotation of the HA412-HOv2 assembly. All gene products putative

Figure S1 **Detailed per trait response to nutrient limitation**. One multipanel per trait showing (a) density distribution of trait values in control (red) and low nutrients (blue). (b) Boxplots of trait values in control (red) and low nutrients (blue). Individual are shown as black circles and connected by black lines between treatments. (c) Trait value in control versus trait value in low nutrients. Genotypes are plotted as green circles with the 1:1 relationship (equal value in control and low nutrients) shown as a dotted line. (d) Trait value in control versus the plasticity in trait value. Plasticity was calculated as the difference in natural log transformed values (control-salt) but converted here to Δ% vs. control (via e^Δln(trait)^ −1) for ease of interpretation. Trait names and units as in table 1 and table 2.

Figure S2 **Whole leaf calcium and magnesium amount**. (a, c) density plot of total leaf calcium and magnesium content (multiplying leaf weight by concentration) under control and low nutrient treatment. (b, d) Boxplots of trait values in control (red) and low nutrients (blue). Individual are shown as black circles and connected by black lines between treatments.

Figure S3 **All PCAs (companion to table 3)**. Principal component analysis of variation in putatively size-independent traits (chlorophyll content, fine root allocation [mass fraction and root fraction], LMA, LMF, RMF, SMF, tap root allocation [mass fraction and root fraction]) and elemental traits (B, Ca, Cu, Fe, Mg, Mn, N, P, K, S, Zn) under control (high nutrients) and low nutrient (10% of control) conditions, as well as the plasticity in trait values between treatments.

Figure S4 **All Manhattans**. Per-trait Manhattan plots are shown for trait values under control and low nutrient stressed conditions, as well as the plasticity between treatments (listed as log difference). SNPs above the red line are significant after multiple comparison correction, SNPs above the blue line are in the top 0.1% of P-values. SNPs are colored by the “significant” haplotype regions per chromosome as displayed in Fig. S7.

Figure S5 **LD plots per chromosome** Linkage disequilibrium (estimated as R^2^) between all SNPs that are significant for at least one trait within and between both treatments. Colored bar notes the haplotype blocks SNPs belong to based on the haplotype map presented in Fig. S7. Block membership is indicated for the calculation based on the genome-wide collection of SNPs (“genome”) and for the re-calculation based on just the significant SNPs (“significant”).

Figure S6 **Trait colocalization per chromosome** Traits in (a) control, (b) low nutrient stressed, and their (c) plasticity (marked as “logdiff”) are clustered based on hierarchical clustering of their bivariate (spearman) correlations. Traits grouped more closely together are more highly correlated. If a linkage block contained a significant SNP for a trait, all other traits were assessed to determine if any had a significant (green) or suggestive (i.e., top 0.1% of all SNPs; gray) in the same block. Plus/minus symbols in the tiles indicate the sign of ββ (effect of minor allele on trait). Linkage blocks are ordered by chromosome with black lines separating them. Sizes of the linkage blocks vary both in physical size as well as gene number.

Figure S7 **Significant regions on genome haplotype block map** Haplotype blocks on the 17 individual sunflower genomes. Blocks are colored from the first SNP to the last SNP within each block. Adjacent blocks are colored in an alternating fashion repeating the same 4 colors. Gray marks along the x-axis note regions with significant trait associations..

