## Supplementary material for "Exceeding expectations: the genomic basis of nitrogen utilization efficiency and integrated trait plasticity as avenues to improve nutrient stress tolerance in cultivated sunflower (*Helianthus annuus* L.)": Suplementary figure

Figure S1 **Detailed per trait response to nutrient limitation.** One multipanel per trait showing (a) density distribution of trait values in control (red) and low nutrients (blue). (b) Boxplots of trait values in control (red) and low nutrients (blue). Individual are shown as black circles and connected by black lines between treatments. (c) Trait value in control versus trait value in low nutrients. Genotypes are plotted as green circles with the 1:1 relationship (equal value in control and low nutrients) shown as a dotted line. (d) Trait value in control versus the plasticity in trait value. Plasticity was calculated as the difference in natural log transformed values (control-salt) but converted here to Δ% vs. control (via $e^{\Delta \ln(\text{trait})}$ - 1) for ease of interpretation. Trait names and units as in table 1 and table 2.

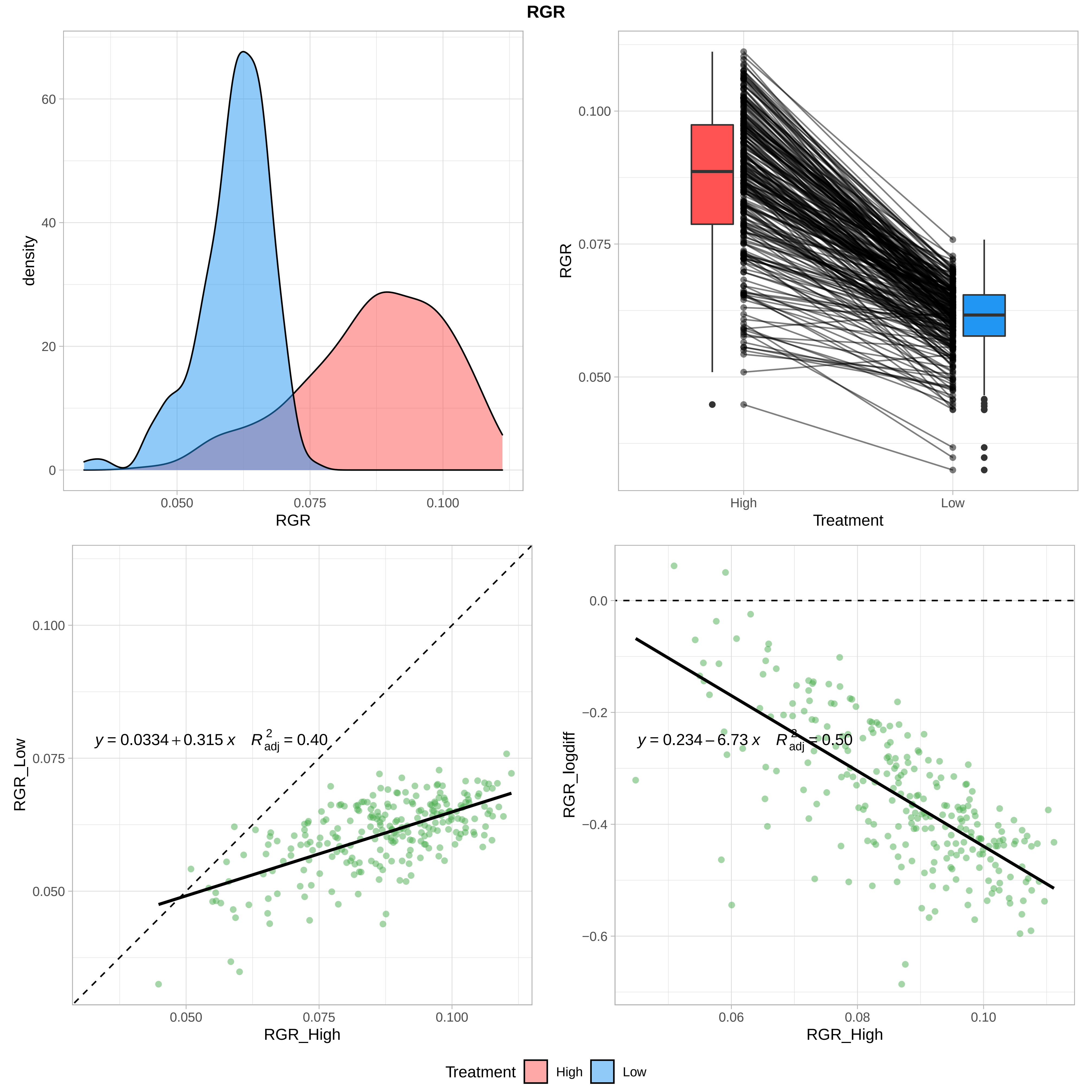
RGR
density
RGR
0.050
0.075
0.100
0
20
40
60
RGR
Treatment
High
Low
0.050
0.075
0.100
y = 0.0334 + 0.315 x    R²adj = 0.40
RGR_Low
RGR_High
y = 0.234 − 6.73 x    R²adj = 0.50
RGR_logdiff
0.0
−0.2
−0.4
−0.6
0.06
0.08
0.10
Treatment
High
Low

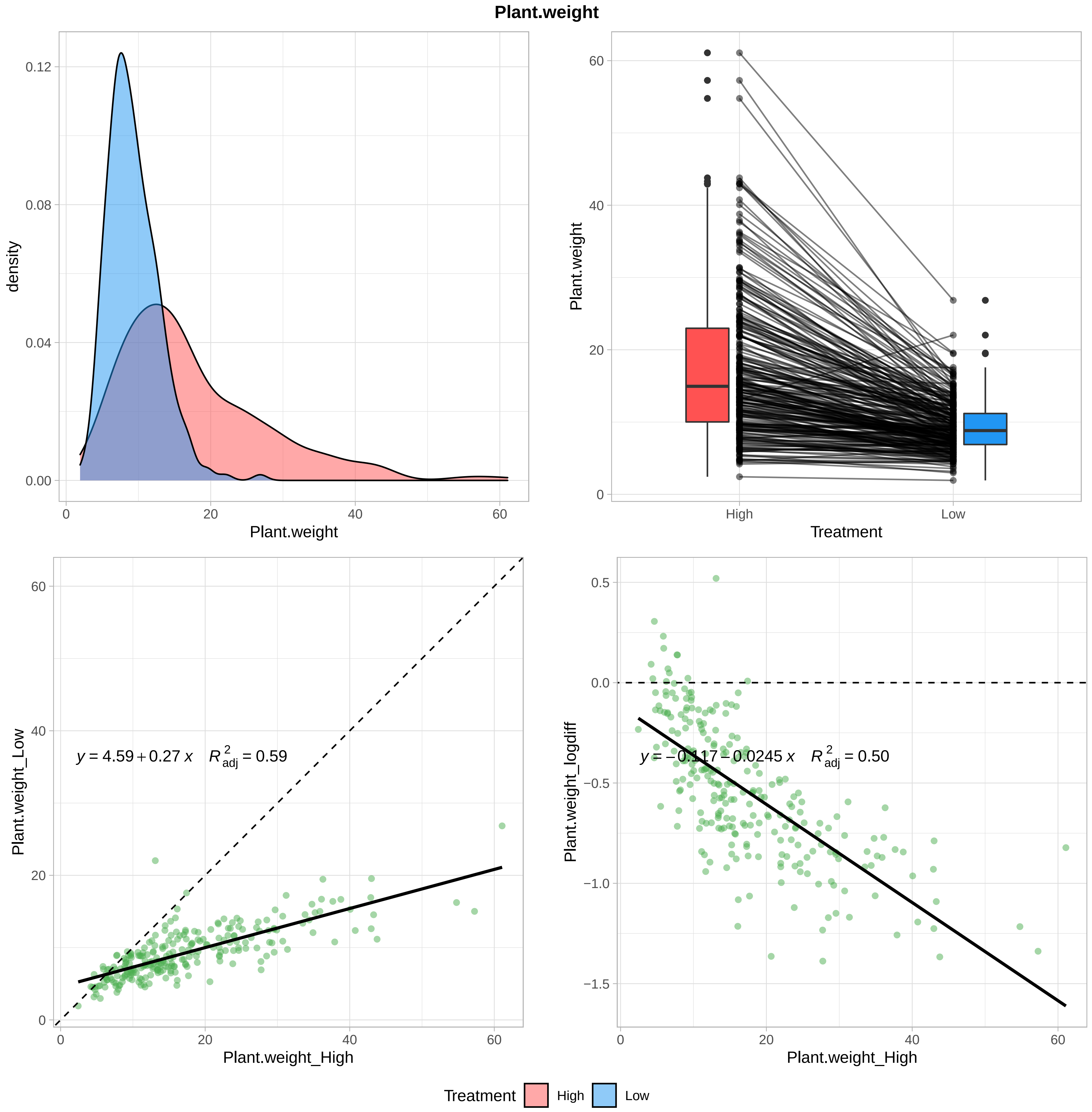

Plant.weight
density
Plant.weight
Plant.weight
Treatment
High
Low
Plant.weight_Low
$y = 4.59 + 0.27\,x \quad R^2_{adj} = 0.59$
Plant.weight_High
Plant.weight_logdiff
$y = -0.117 - 0.0245\,x \quad R^2_{adj} = 0.50$
Plant.weight_High
Treatment
High
Low

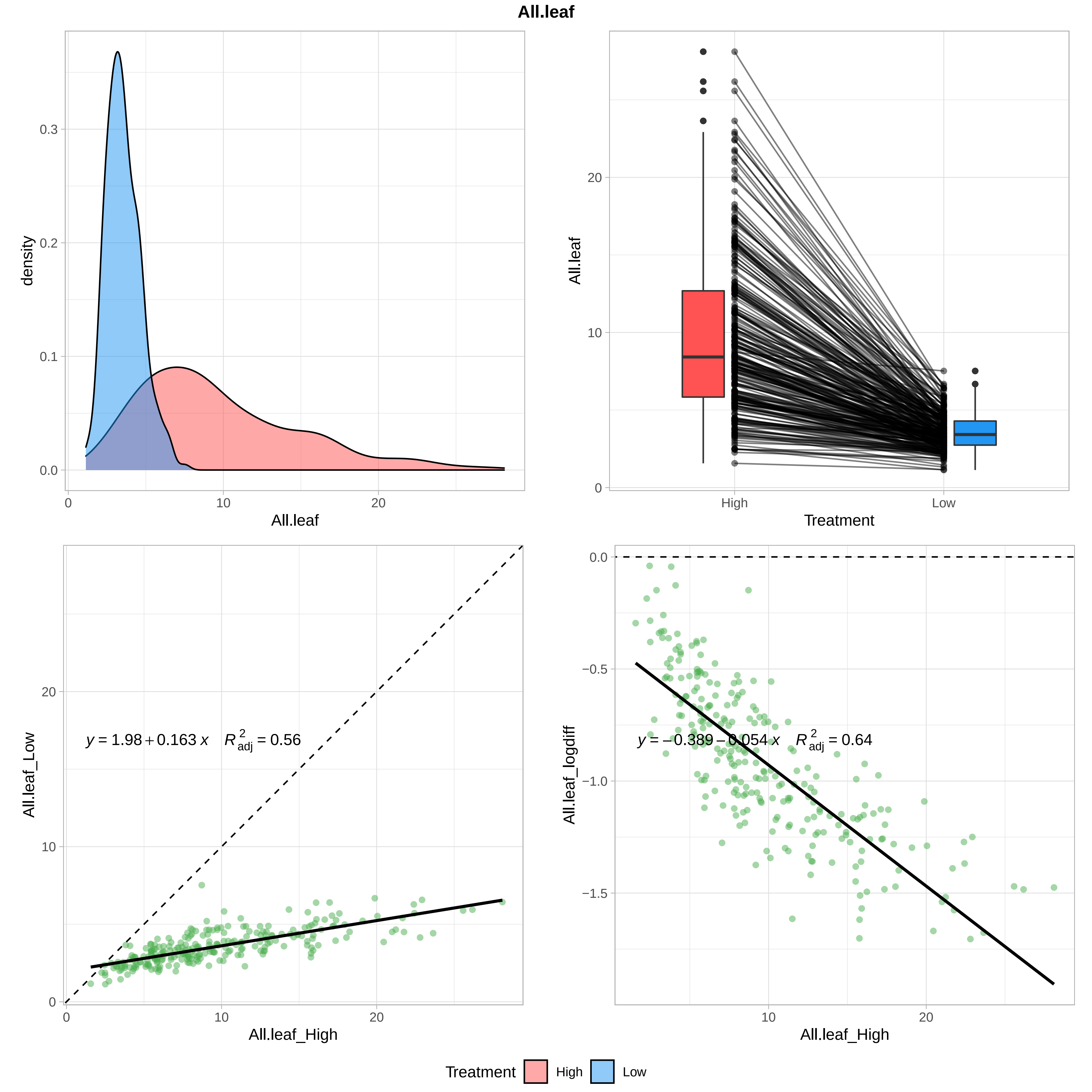
All.leaf
density
0.0
0.1
0.2
0.3
0
10
20
All.leaf
All.leaf
0
10
20
High
Low
Treatment
All.leaf_Low
0
10
20
y = 1.98 + 0.163 x  R2adj = 0.56
All.leaf_High
All.leaf_logdiff
0.0
-0.5
-1.0
-1.5
y = −0.389 − 0.054 x  R2adj = 0.64
All.leaf_High
Treatment
High
Low

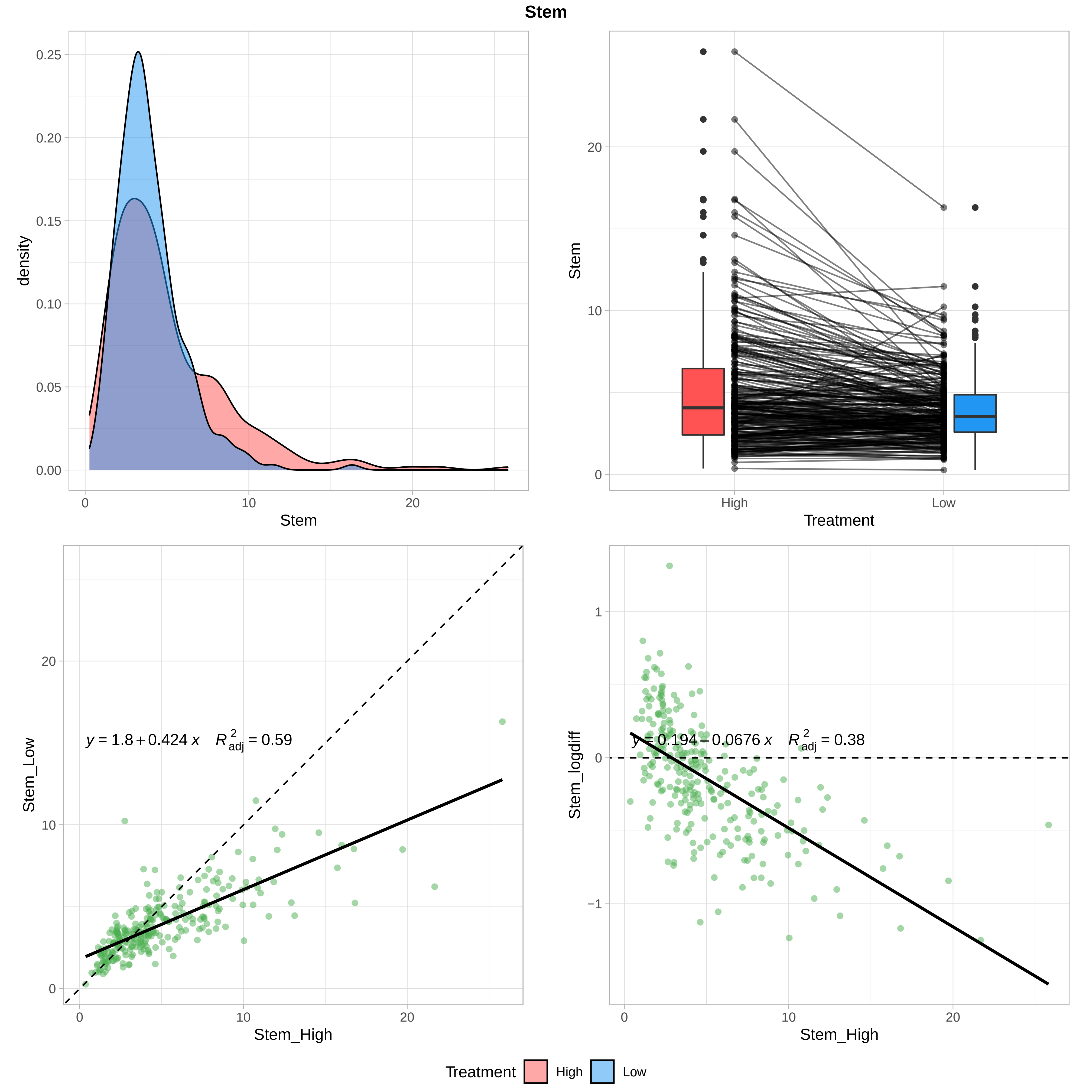

Stem
density
Stem
Treatment
High
Low
Stem_Low
Stem_High
$y = 1.8 + 0.424\,x \quad R^2_{adj} = 0.59$
Stem_logdiff
$y = 0.194 - 0.0676\,x \quad R^2_{adj} = 0.38$
Treatment High Low

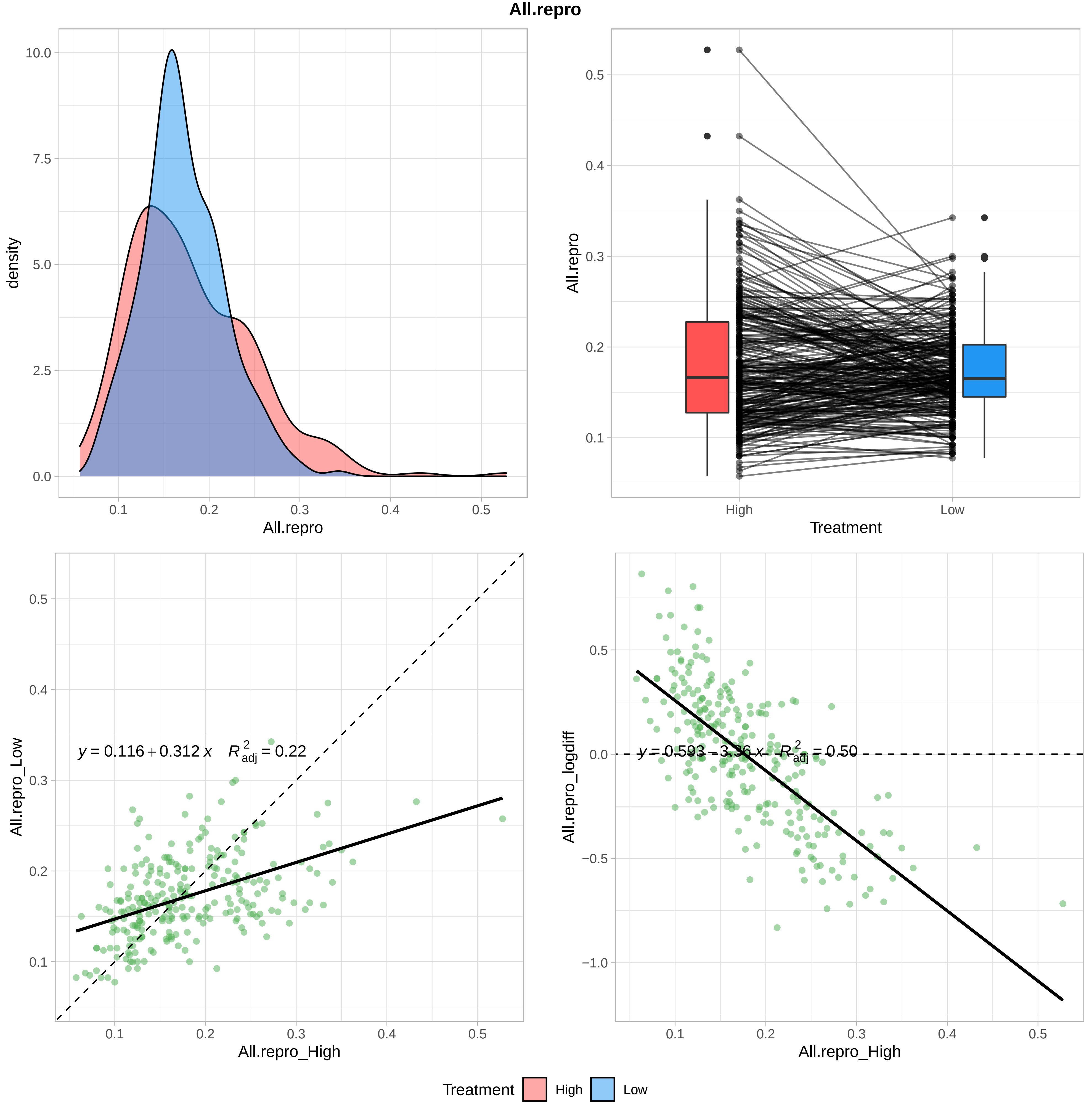
All.repro
density
All.repro
10.0
7.5
5.0
2.5
0.0
0.1
0.2
0.3
0.4
0.5
All.repro
Treatment
High
Low
0.5
0.4
0.3
0.2
0.1
All.repro_Low
All.repro_High
y = 0.116 + 0.312 x    R2adj = 0.22
All.repro_logdiff
y = 0.593 − 3.36 x    R2adj = 0.50
−1.0
−0.5
Treatment
High
Low

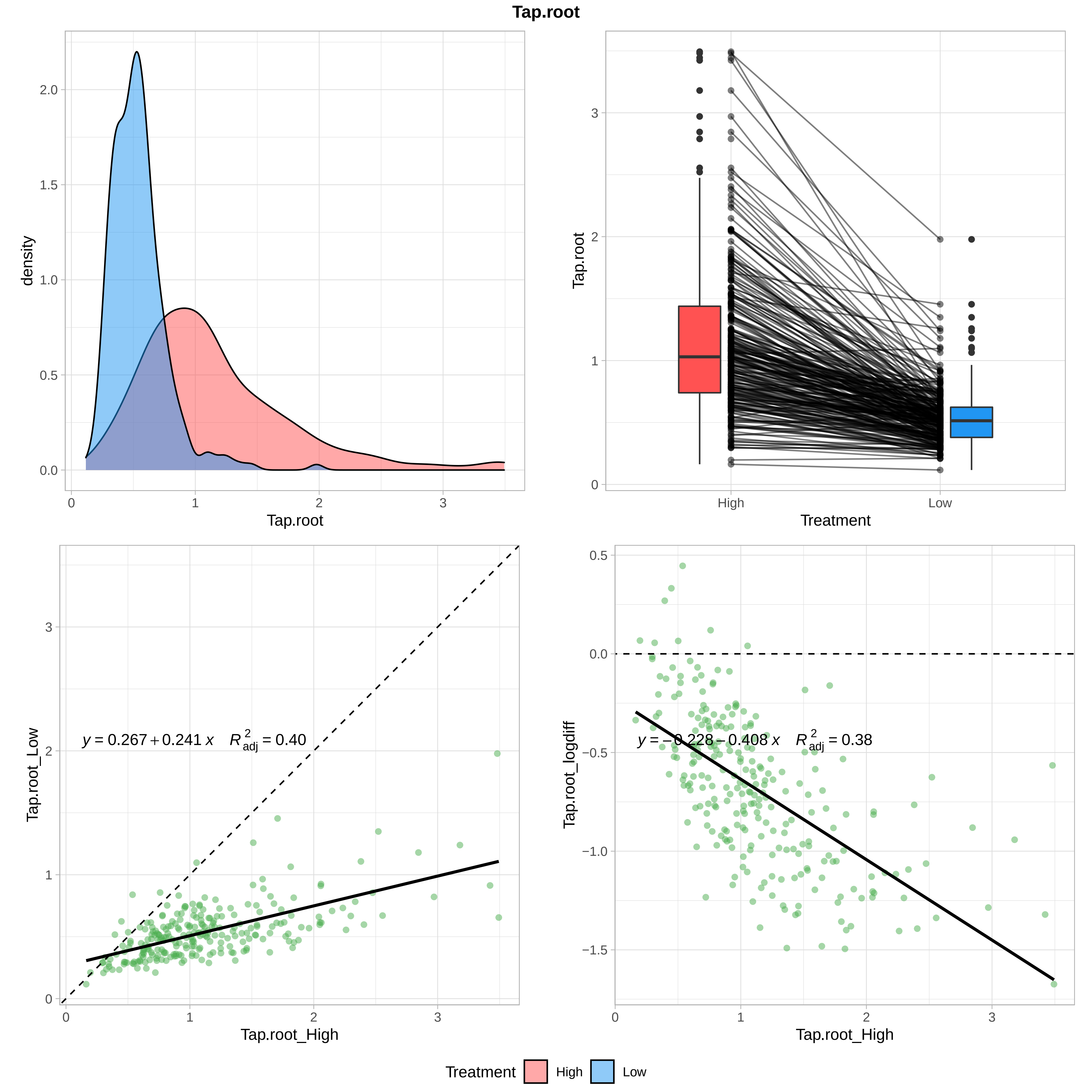
Tap.root
density
Tap.root
Tap.root
High
Low
Treatment
$y = 0.267 + 0.241\ x \quad R^2_{adj} = 0.40$
Tap.root_Low
Tap.root_High
$y = -0.228 - 0.408\ x \quad R^2_{adj} = 0.38$
Tap.root_logdiff
Tap.root_High
Treatment
High
Low

**All.fine**

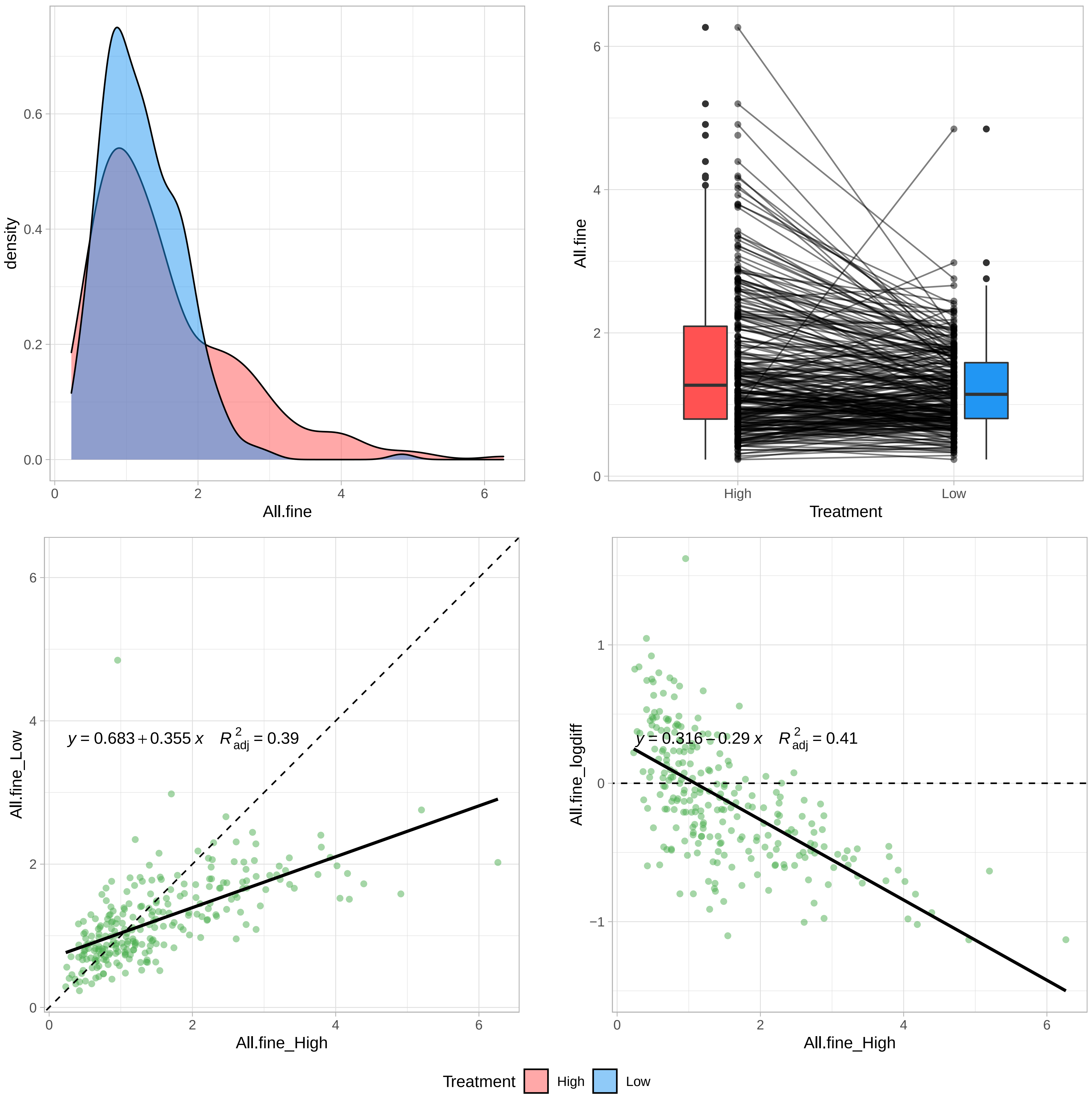

density
All.fine
All.fine
Treatment
High
Low
y = 0.683 + 0.355 x   R²adj = 0.39
All.fine_Low
All.fine_High
y = 0.316 − 0.29 x   R²adj = 0.41
All.fine_logdiff
All.fine_High
Treatment
High
Low

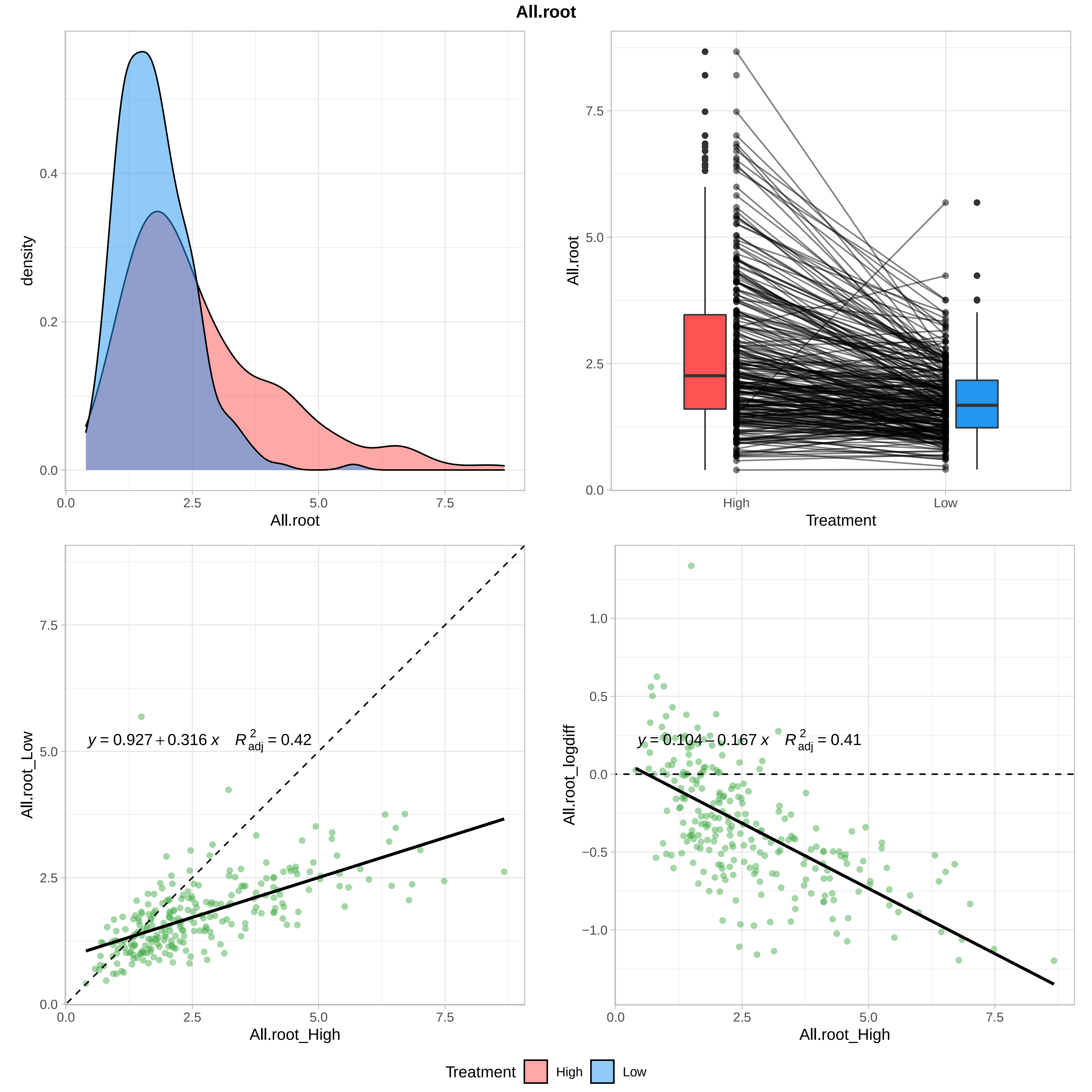

All.root
density
0.0
0.2
0.4
0.0
2.5
5.0
7.5
All.root
7.5
5.0
2.5
0.0
High
Low
Treatment
All.root_Low
All.root_High
y = 0.927 + 0.316 x   R2adj = 0.42
All.root_logdiff
1.0
0.5
0.0
−0.5
−1.0
y = 0.104 − 0.167 x   R2adj = 0.41
Treatment
High
Low

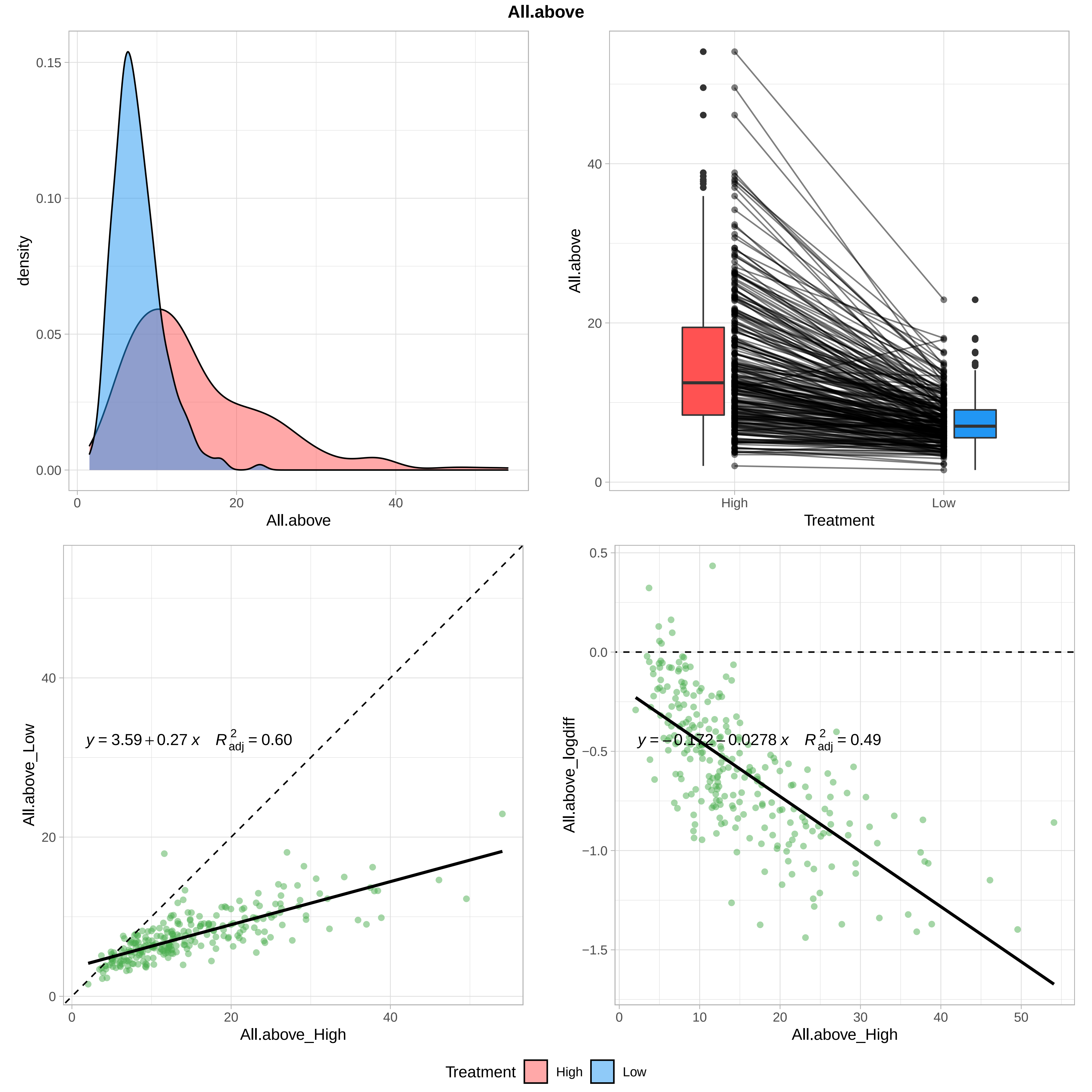

All.above
density
All.above
All.above
Treatment
High
Low
All.above_Low
All.above_High
$y = 3.59 + 0.27\,x \quad R^2_{adj} = 0.60$
All.above_logdiff
All.above_High
$y = -0.172 - 0.0278\,x \quad R^2_{adj} = 0.49$
Treatment
High
Low

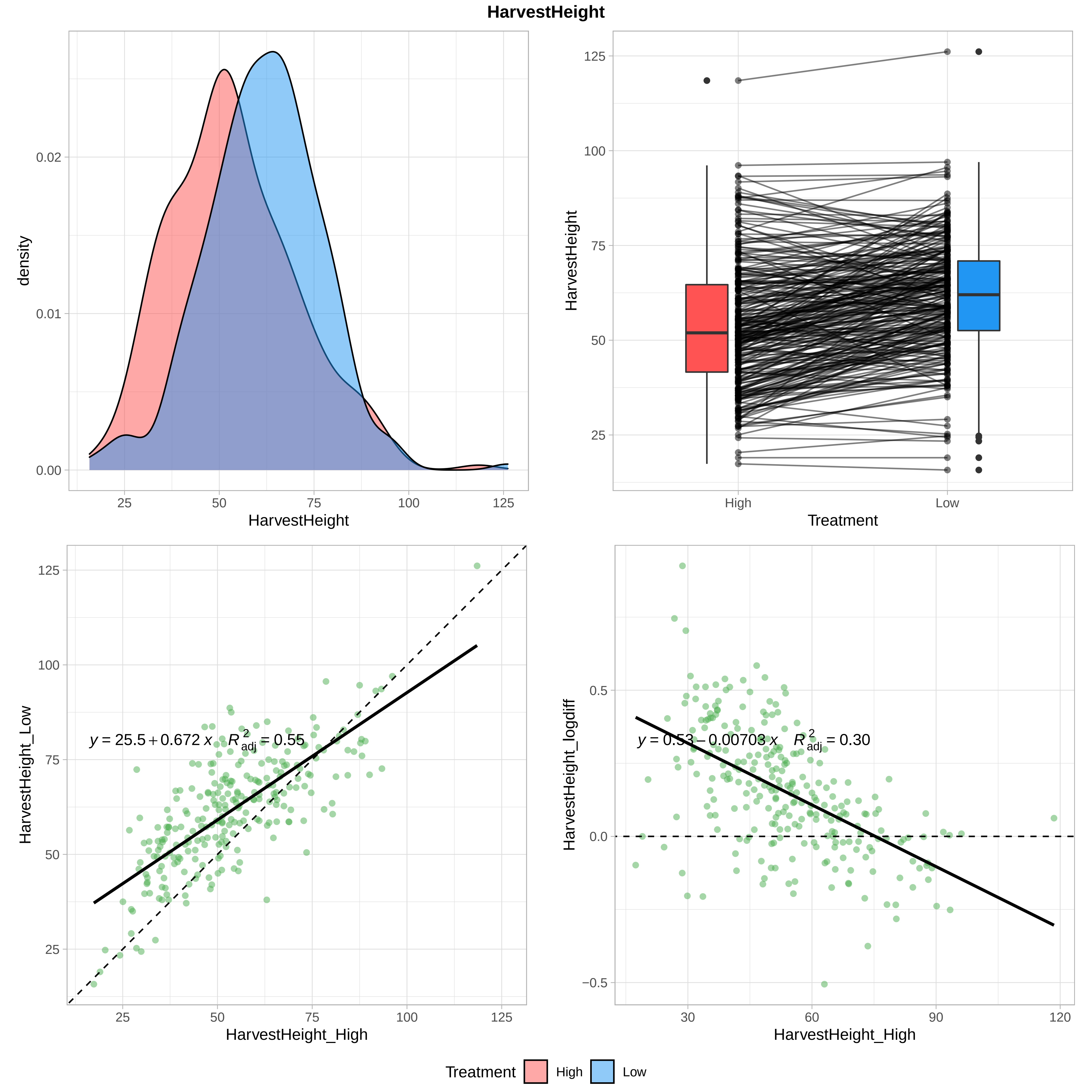

HarvestHeight
density
HarvestHeight
Treatment
High
Low
HarvestHeight_Low
HarvestHeight_High
y = 25.5 + 0.672 x   R²adj = 0.55
HarvestHeight_logdiff
y = 0.53 − 0.00703 x   R²adj = 0.30
Treatment
High
Low

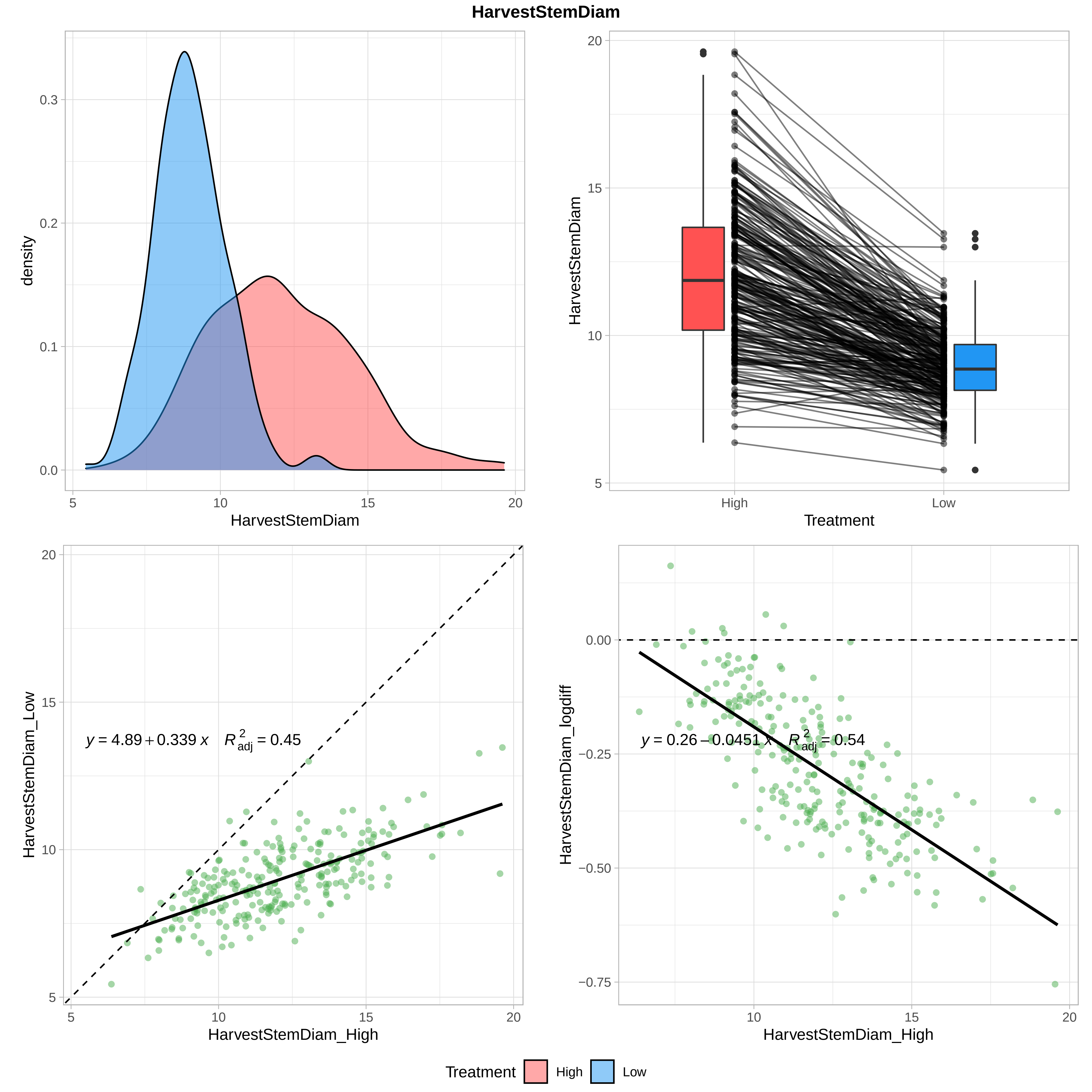

HarvestStemDiam
density
HarvestStemDiam
HarvestStemDiam
Treatment
High
Low
HarvestStemDiam_Low
HarvestStemDiam_High
y = 4.89 + 0.339 x   R²adj = 0.45
HarvestStemDiam_logdiff
HarvestStemDiam_High
y = 0.26 − 0.0451 x   R²adj = 0.54
Treatment
High
Low

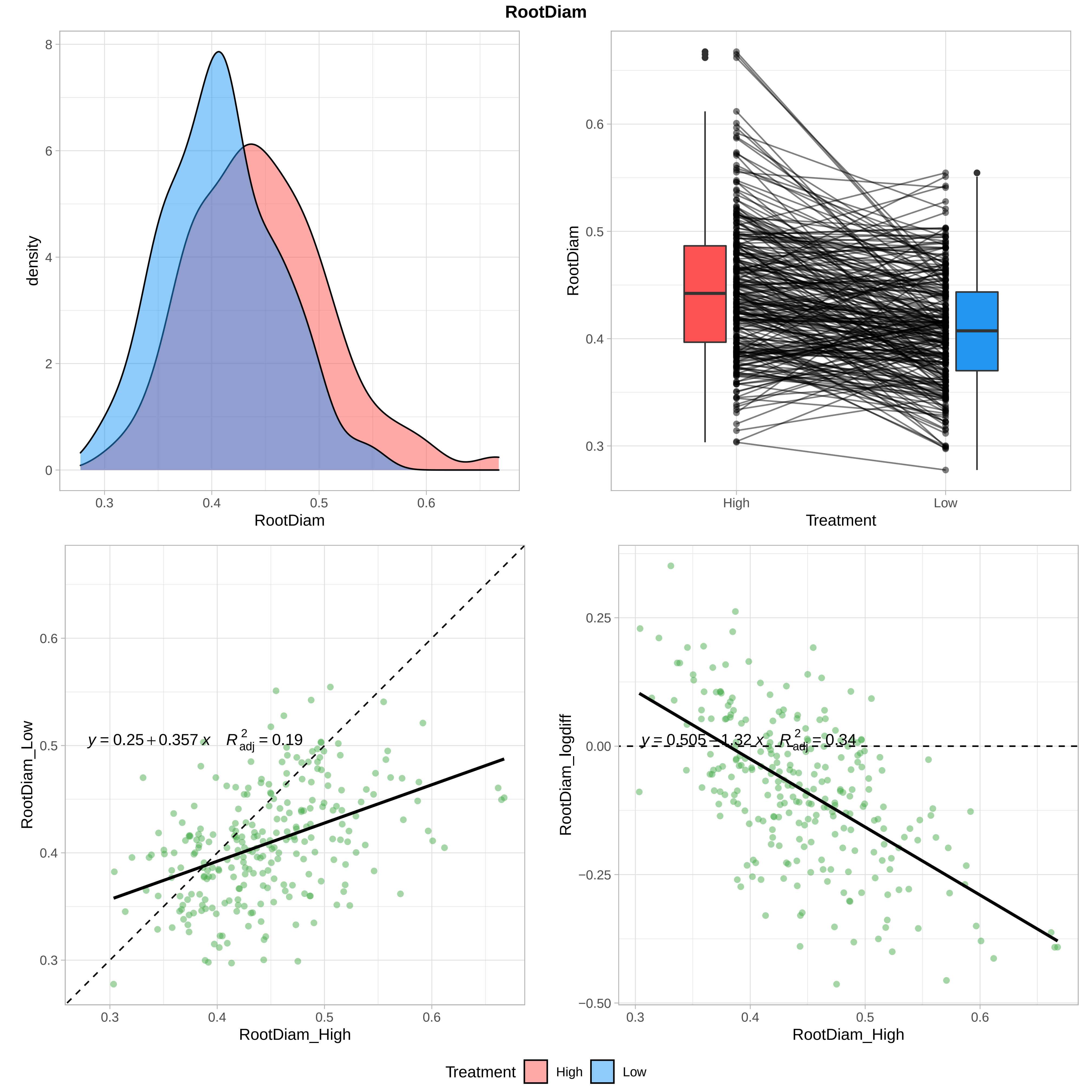
RootDiam
density
RootDiam
Treatment
High
Low
$y = 0.25 + 0.357\,x \quad R^2_{adj} = 0.19$
RootDiam_Low
RootDiam_High
$y = 0.505 - 1.32\,x \quad R^2_{adj} = 0.34$
RootDiam_logdiff
Treatment High Low

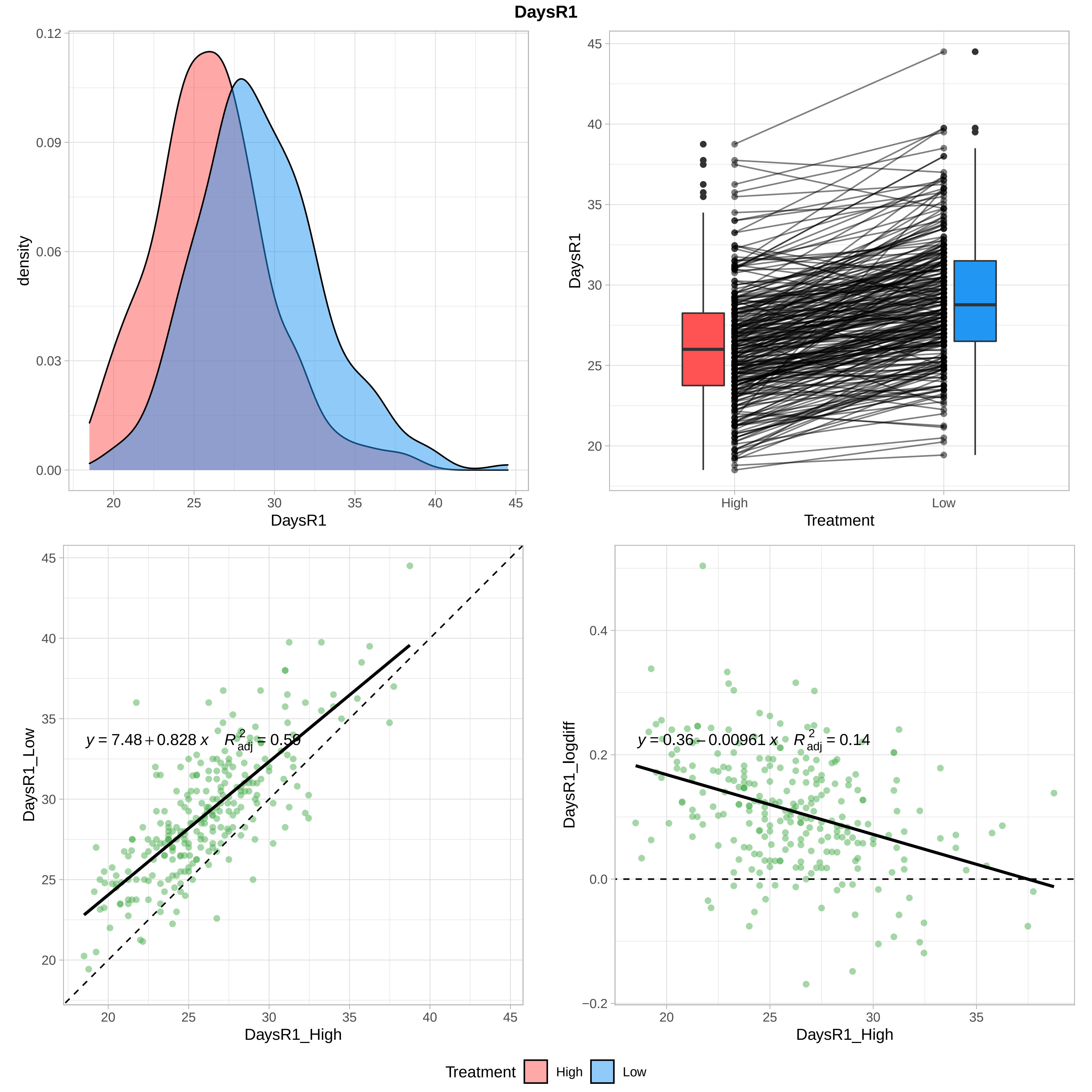

DaysR1
density
DaysR1
Treatment
High
Low
$y = 7.48 + 0.828\,x \quad R^2_{adj} = 0.59$
DaysR1_Low
DaysR1_High
$y = 0.36 - 0.00961\,x \quad R^2_{adj} = 0.14$
DaysR1_logdiff
DaysR1_High
Treatment High Low

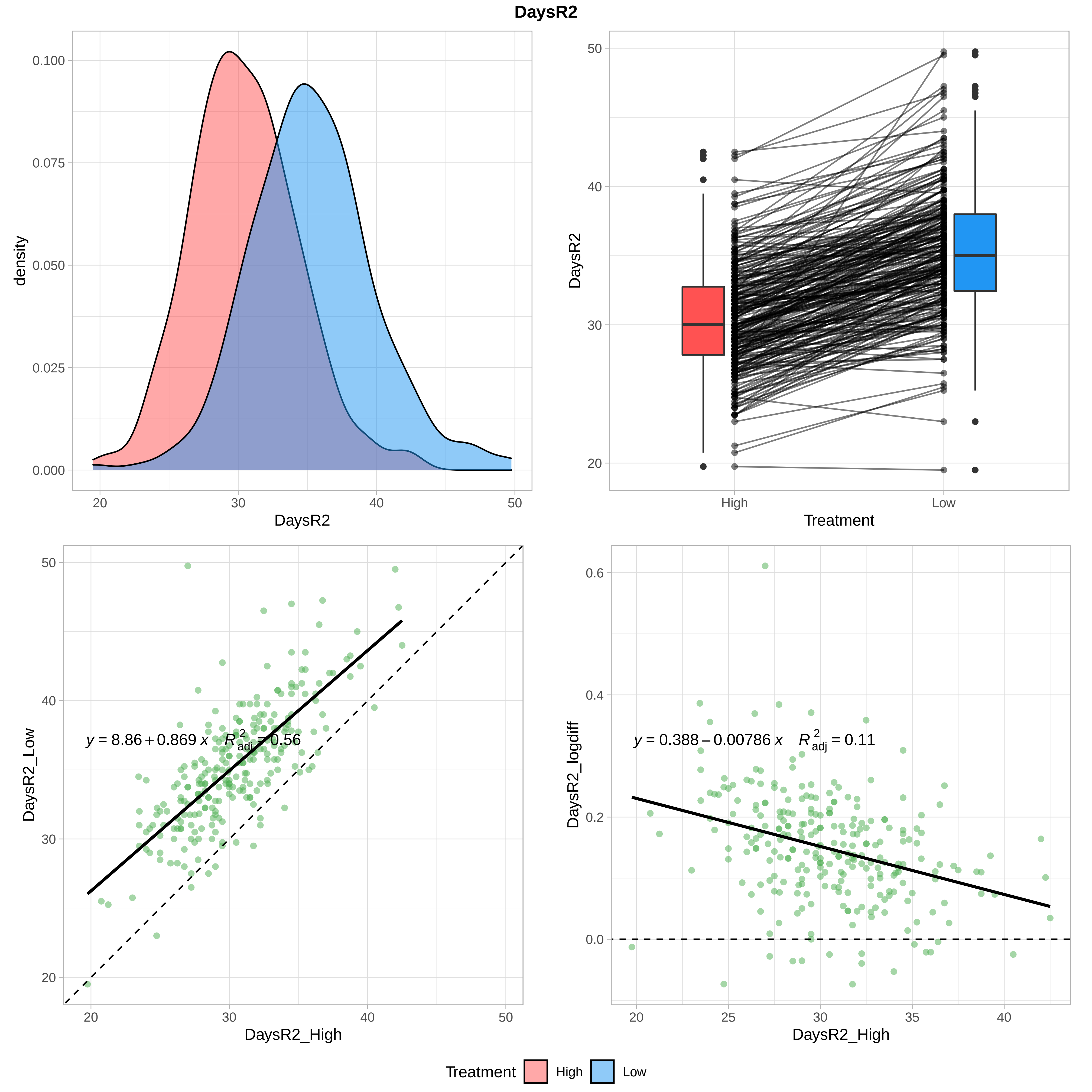
DaysR2
density
DaysR2
Treatment
DaysR2_Low
DaysR2_High
y = 8.86 + 0.869 x   R²adj = 0.56
DaysR2_logdiff
DaysR2_High
y = 0.388 − 0.00786 x   R²adj = 0.11
High
Low
Treatment
High
Low

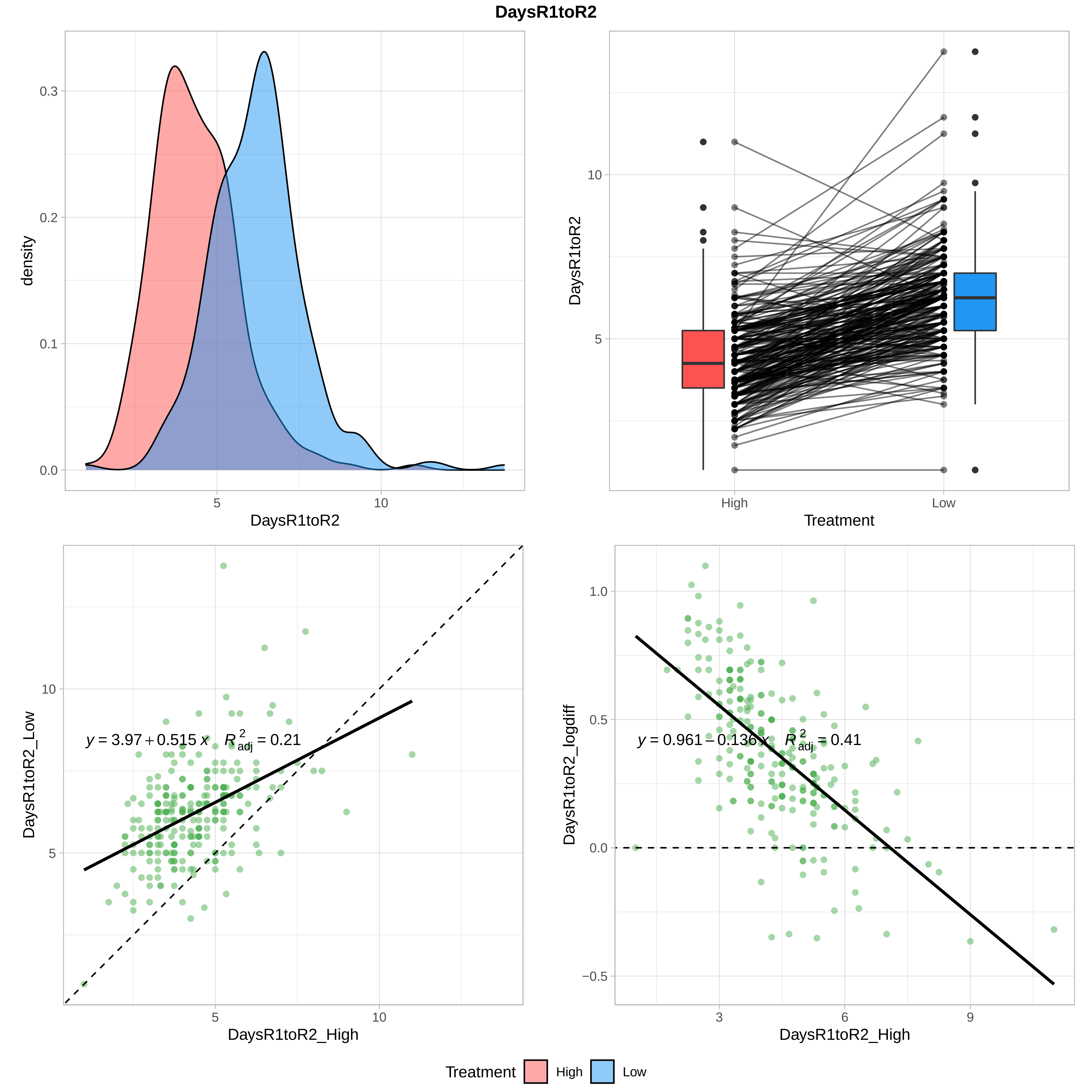
DaysR1toR2
density
0.3
0.2
0.1
0.0
5
10
DaysR1toR2
DaysR1toR2
10
5
High
Low
Treatment
DaysR1toR2_Low
$y = 3.97 + 0.515\,x \quad R^2_{adj} = 0.21$
10
5
5
10
DaysR1toR2_High
DaysR1toR2_logdiff
$y = 0.961 - 0.136\,x \quad R^2_{adj} = 0.41$
1.0
0.5
0.0
−0.5
3
6
9
DaysR1toR2_High
Treatment
High
Low

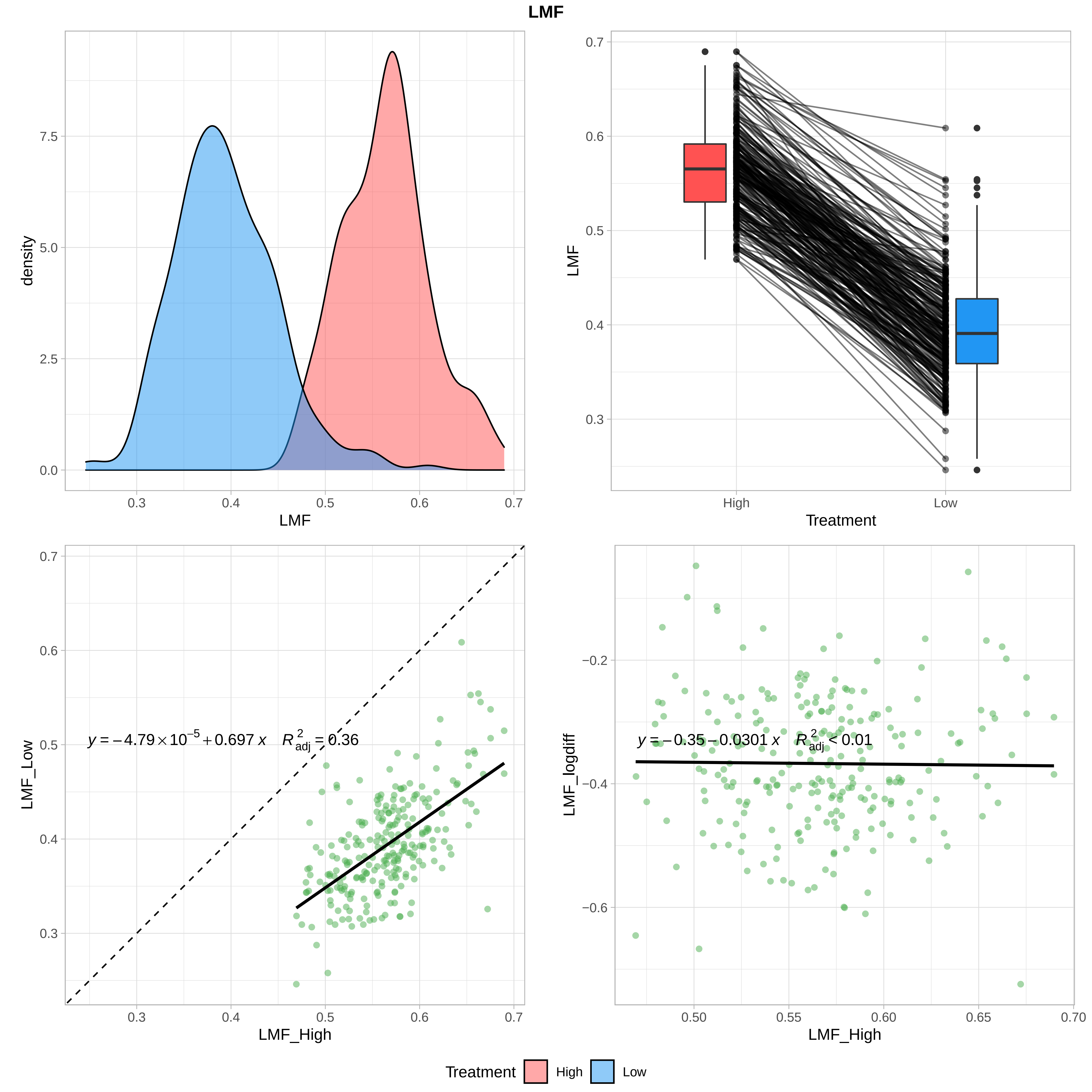
LMF
density
7.5
5.0
2.5
0.0
0.3
0.4
0.5
0.6
0.7
LMF
0.7
0.6
0.5
0.4
0.3
LMF
High
Low
Treatment
0.7
0.6
0.5
0.4
0.3
LMF_Low
y = −4.79 × 10^−5 + 0.697 x   R²adj = 0.36
0.3
0.4
0.5
0.6
0.7
LMF_High
−0.2
−0.4
−0.6
LMF_logdiff
y = −0.35 − 0.0301 x   R²adj < 0.01
0.50
0.55
0.60
0.65
0.70
LMF_High
Treatment
High
Low

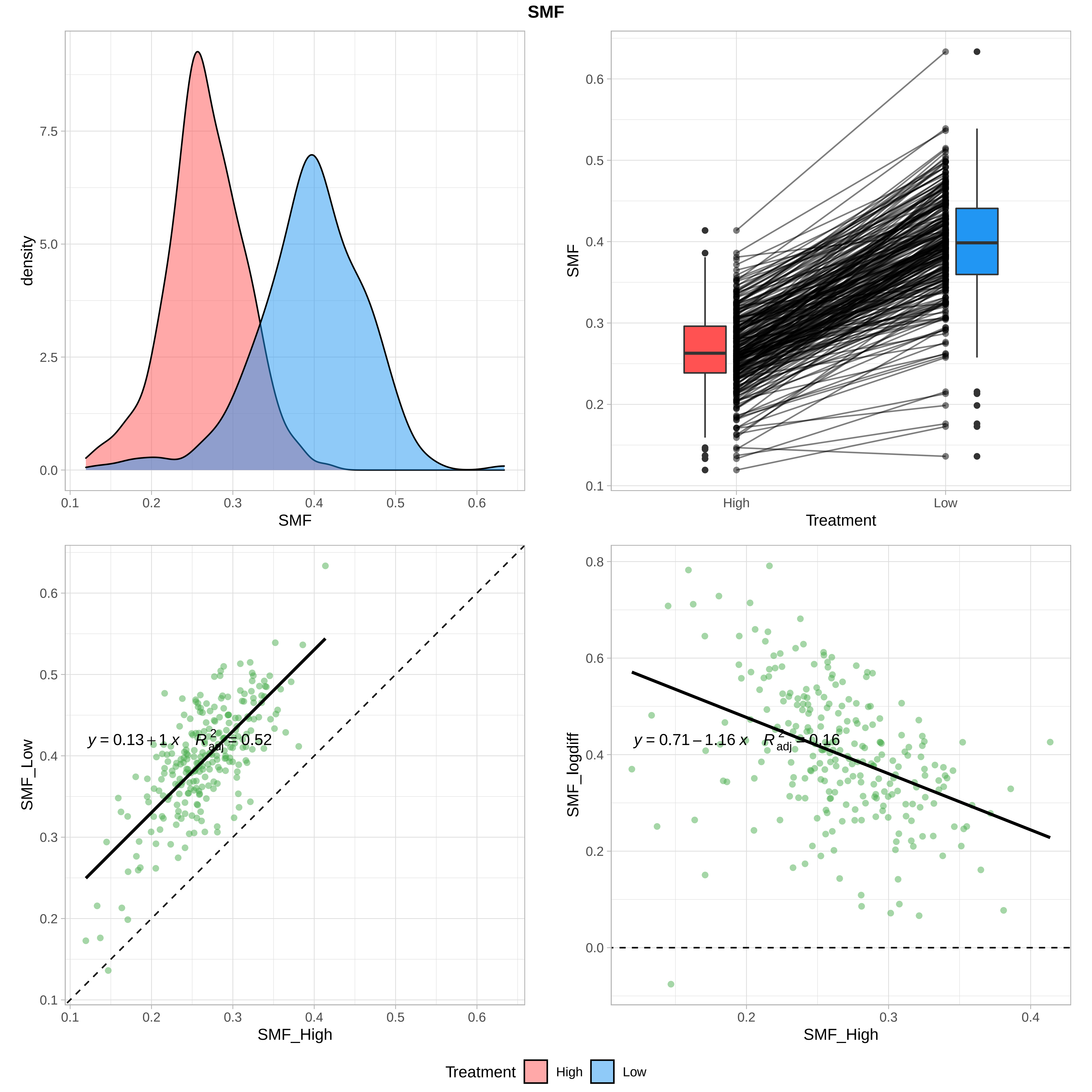

SMF
density
SMF
SMF
Treatment
High
Low
y = 0.13 + 1 x   $R^2_{adj}$ = 0.52
SMF_Low
SMF_High
y = 0.71 − 1.16 x   $R^2_{adj}$ = 0.16
SMF_logdiff
SMF_High
Treatment High Low

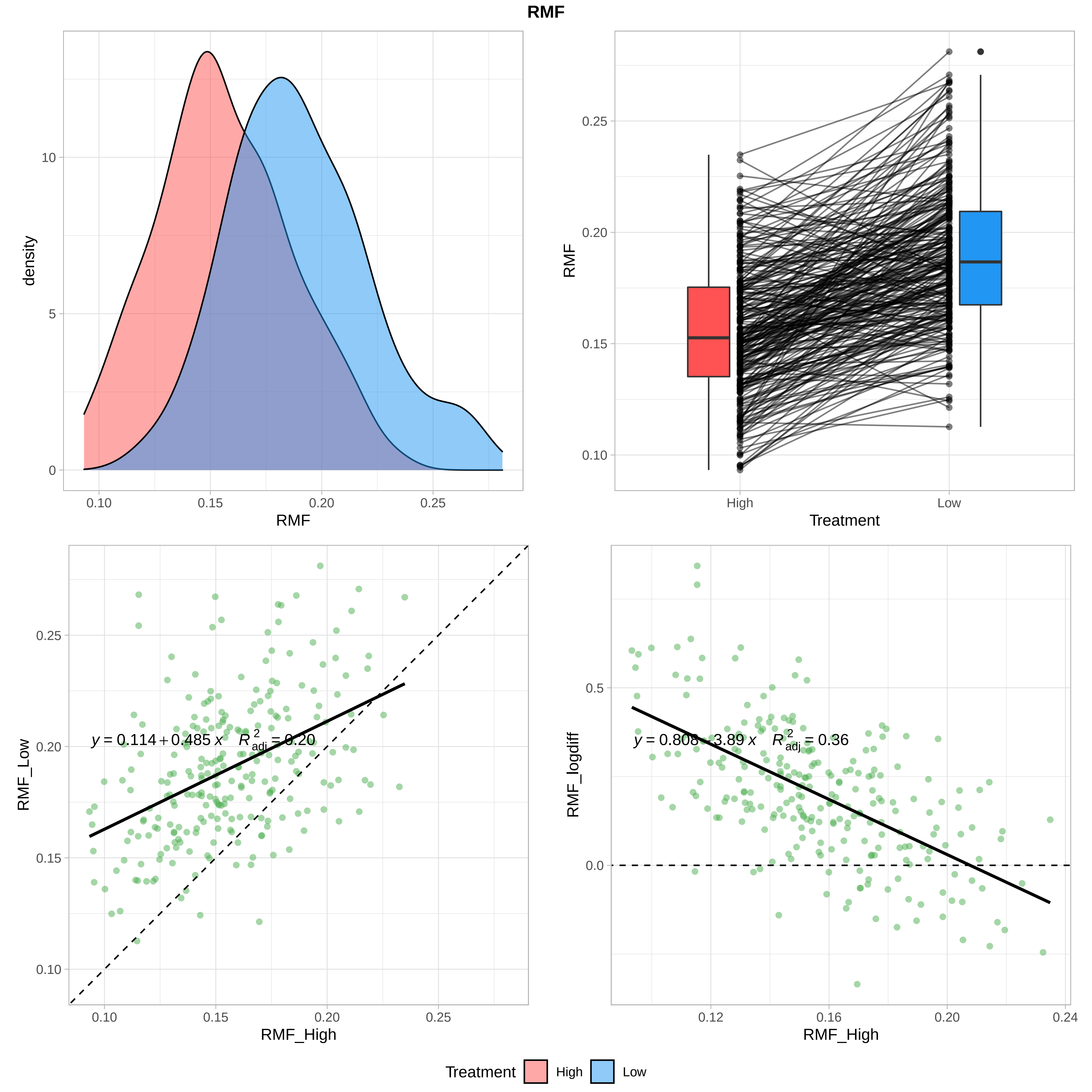
RMF
density
RMF
Treatment
RMF_Low
RMF_High
$y = 0.114 + 0.485\,x \quad R^2_{adj} = 0.20$
RMF_logdiff
$y = 0.808 - 3.89\,x \quad R^2_{adj} = 0.36$
Treatment High Low

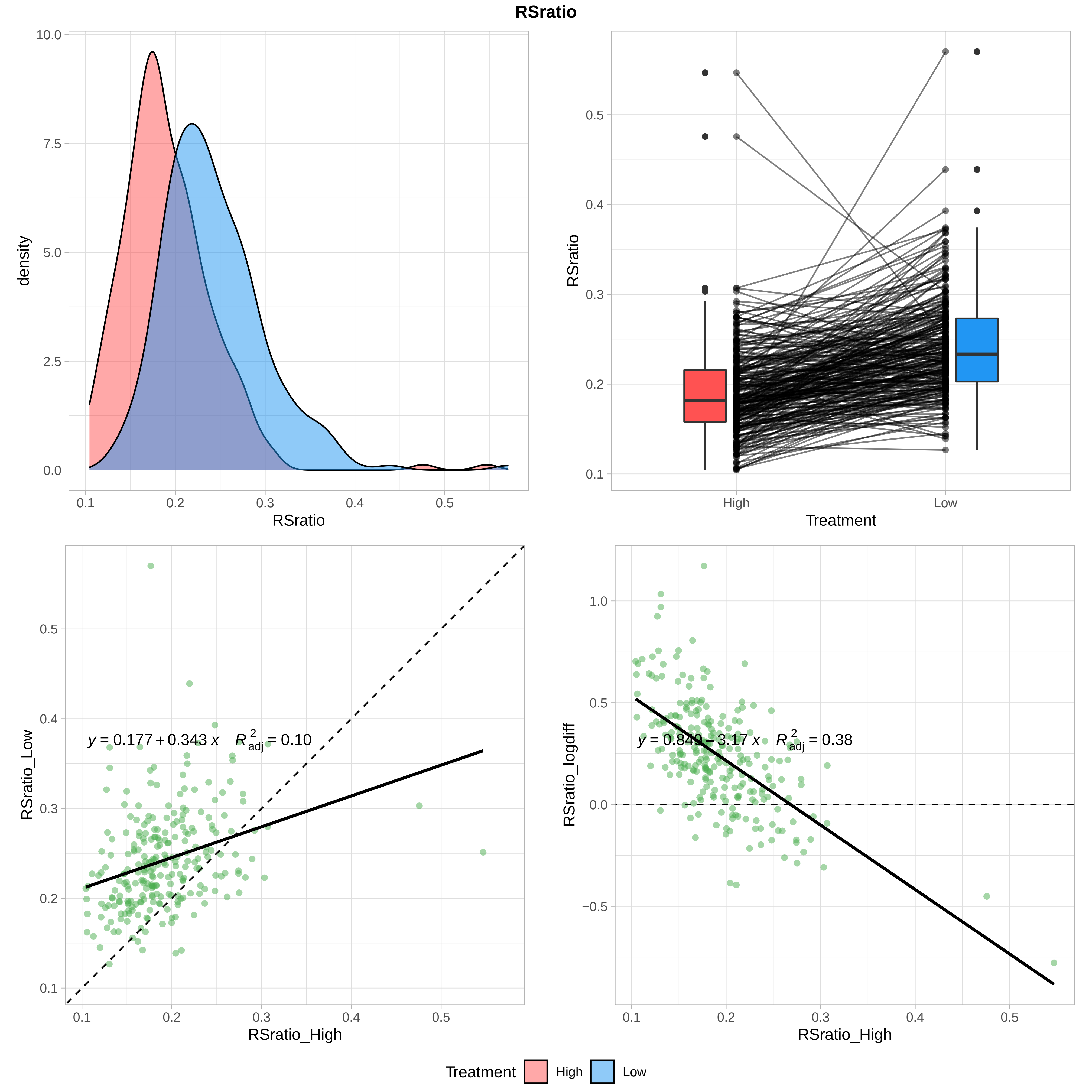
RSratio
density
RSratio
Treatment
High
Low
$y = 0.177 + 0.343\,x \quad R^2_{adj} = 0.10$
RSratio_Low
RSratio_High
$y = 0.849 - 3.17\,x \quad R^2_{adj} = 0.38$
RSratio_logdiff
RSratio_High
Treatment High Low

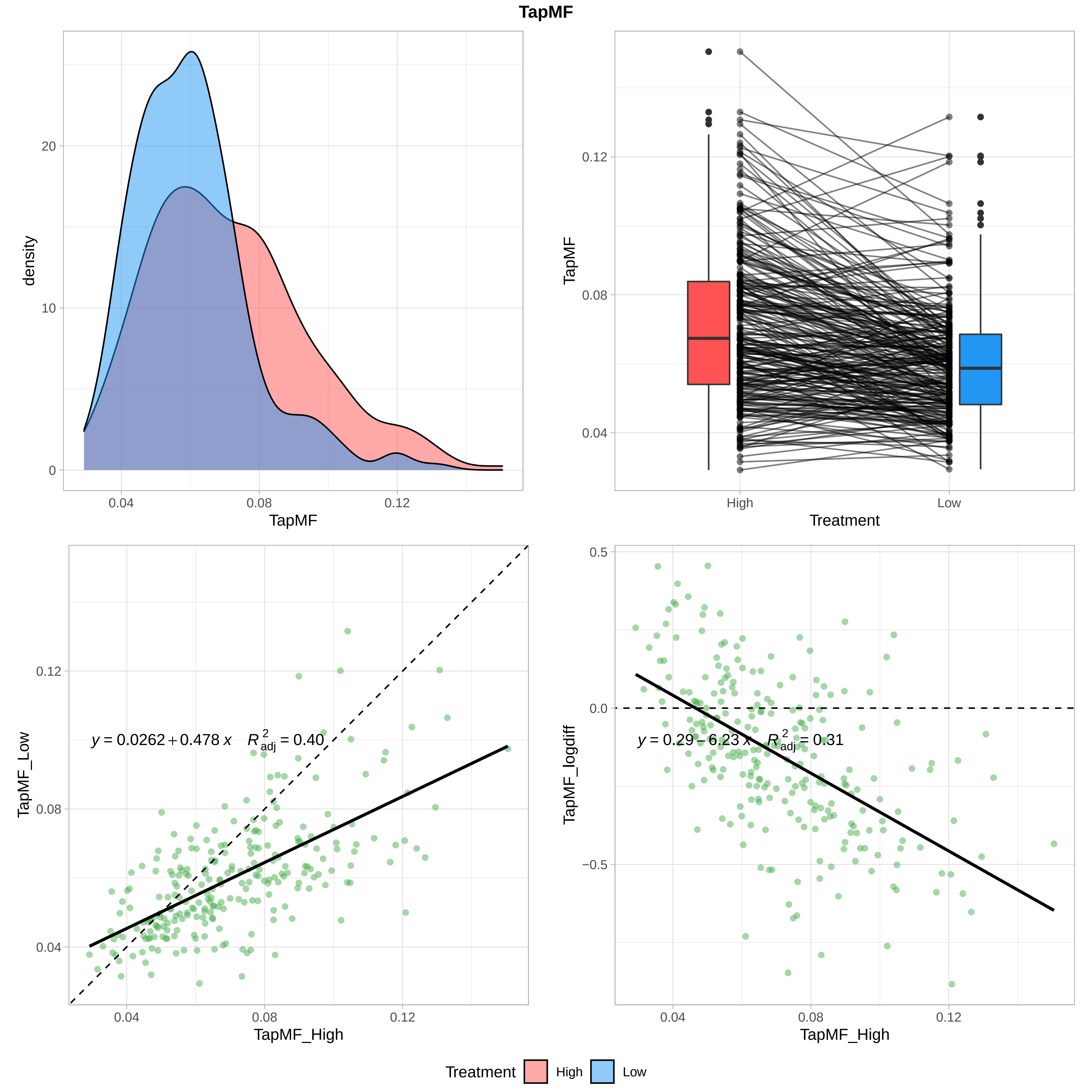

TapMF
density
TapMF
0
10
20
0.04
0.08
0.12
TapMF
Treatment
High
Low
0.04
0.08
0.12
TapMF_Low
TapMF_High
y = 0.0262 + 0.478 x   R²adj = 0.40
TapMF_logdiff
y = 0.29 − 6.23 x   R²adj = 0.31
0.5
0.0
−0.5
Treatment
High
Low

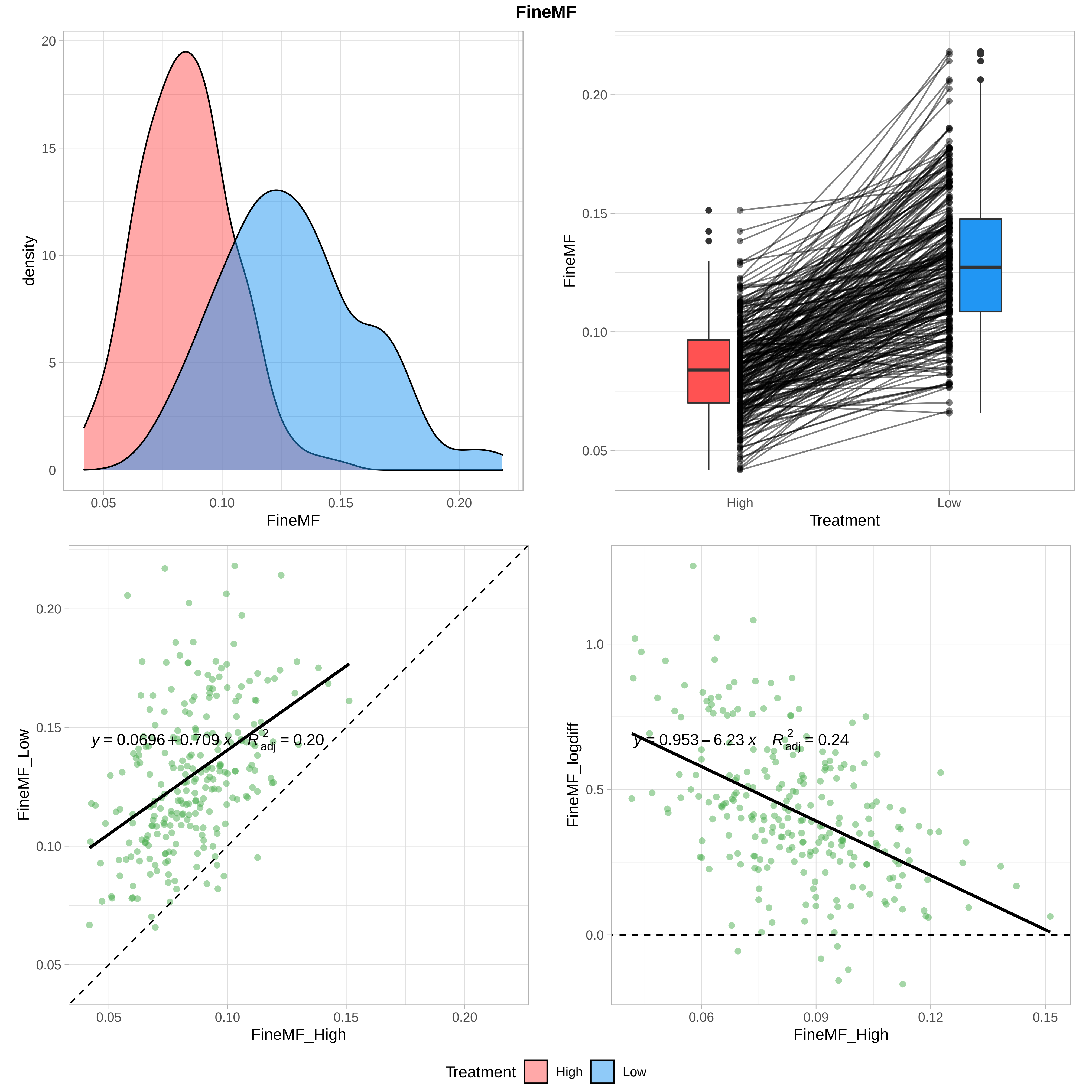

FineMF
density
FineMF
Treatment
High
Low
FineMF_Low
FineMF_High
FineMF_logdiff
$y = 0.0696 + 0.709\,x \quad R^2_{adj} = 0.20$
$y = 0.953 - 6.23\,x \quad R^2_{adj} = 0.24$
Treatment
High
Low

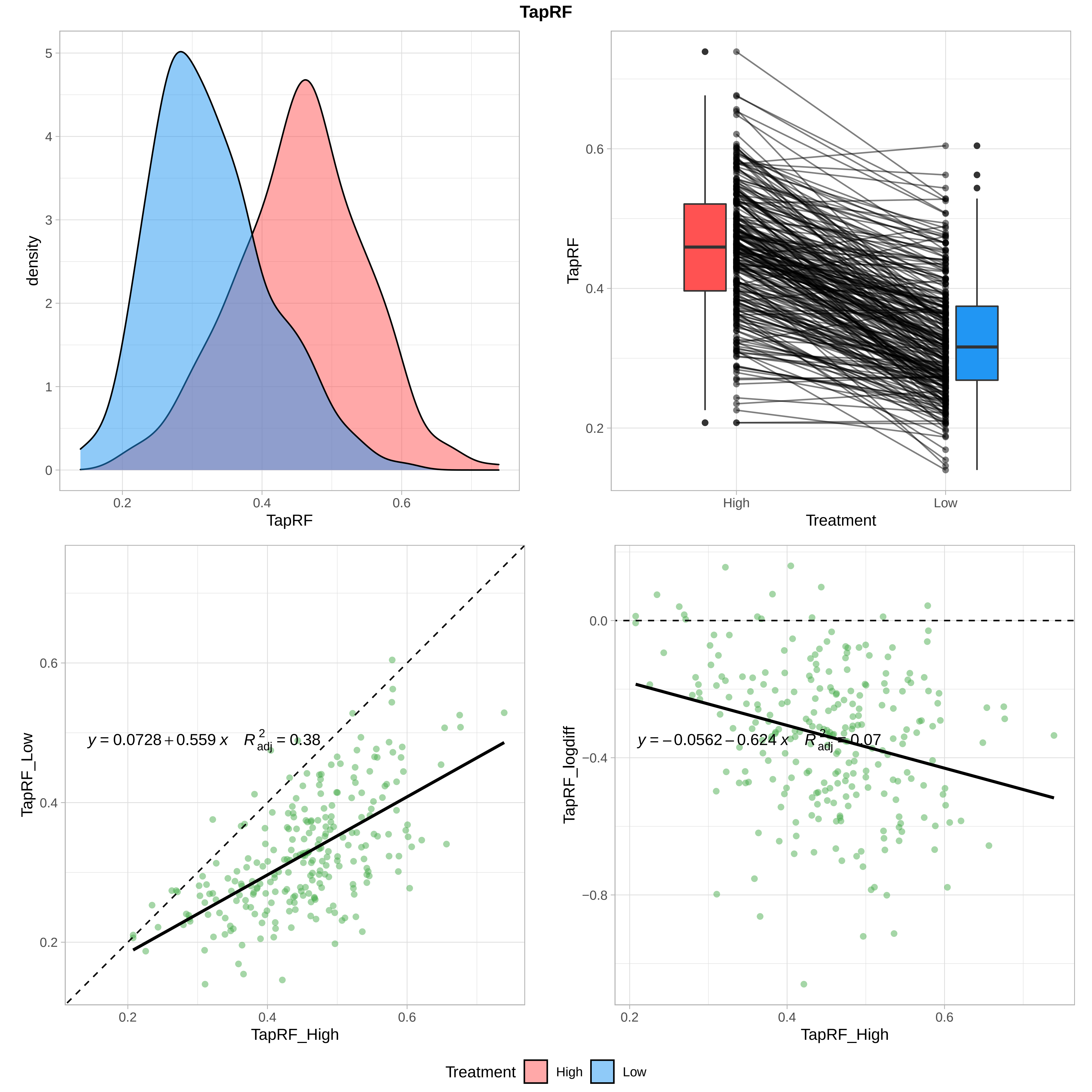

TapRF
density
TapRF
TapRF
Treatment
High
Low
TapRF_Low
TapRF_High
y = 0.0728 + 0.559 x   R²adj = 0.38
TapRF_logdiff
TapRF_High
y = −0.0562 − 0.624 x   R²adj = 0.07
Treatment
High
Low

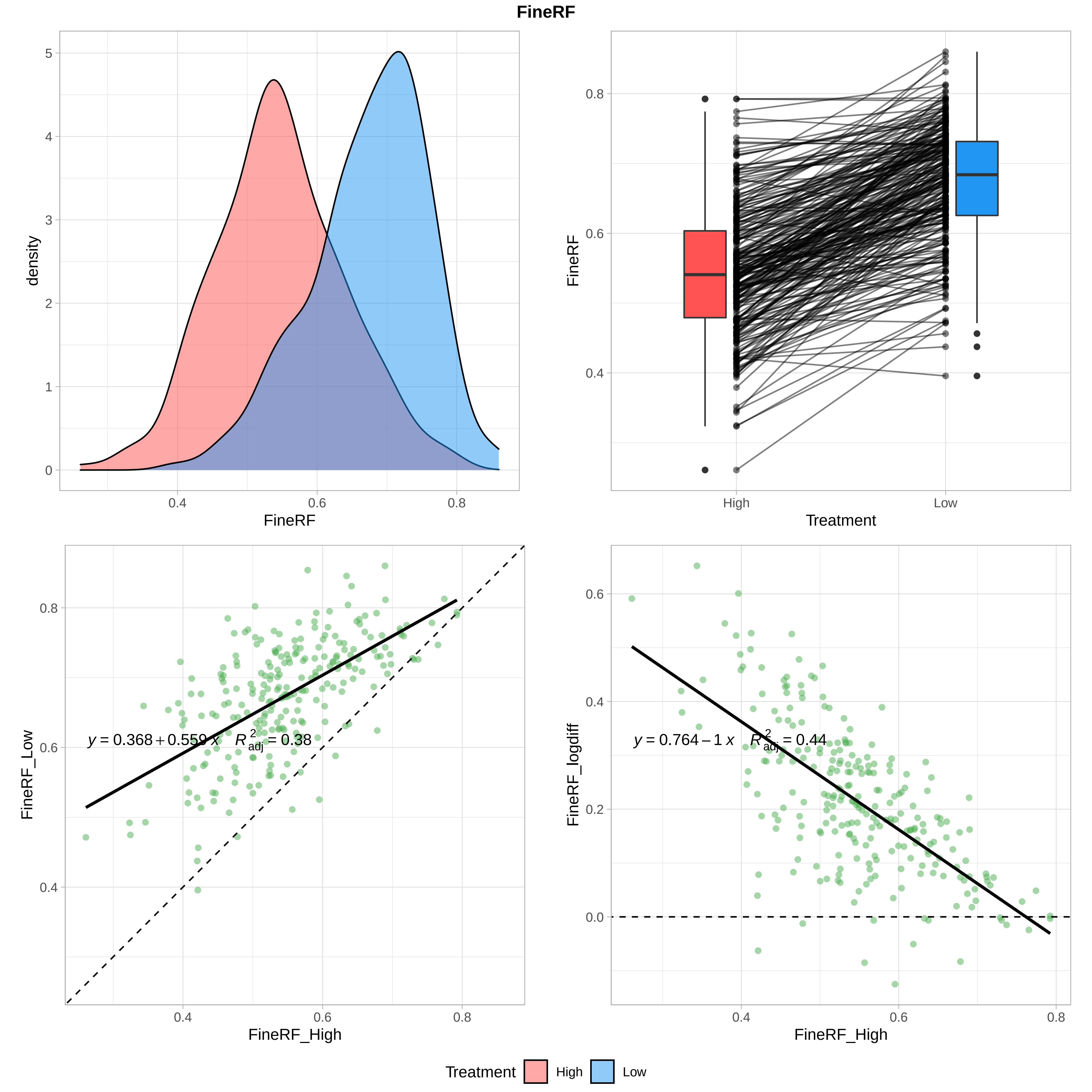

FineRF
density
FineRF
Treatment
High
Low
FineRF_Low
FineRF_High
$y = 0.368 + 0.559\,x \quad R^2_{adj} = 0.38$
FineRF_logdiff
$y = 0.764 - 1\,x \quad R^2_{adj} = 0.44$
Treatment High Low

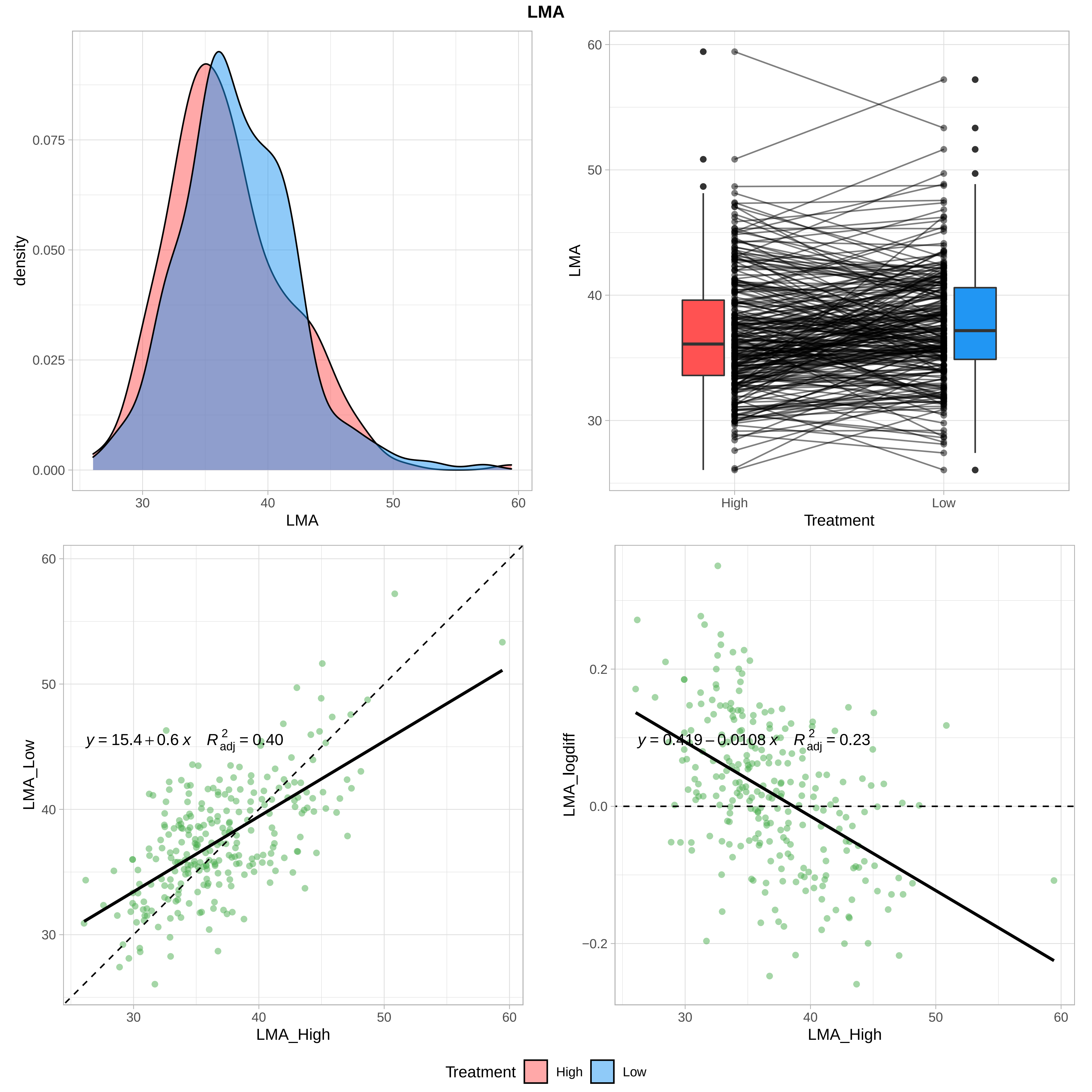
LMA
density
0.000
0.025
0.050
0.075
30
40
50
60
LMA
LMA
Treatment
High
Low
LMA_Low
LMA_High
y = 15.4 + 0.6 x  R²adj = 0.40
LMA_logdiff
y = 0.419 − 0.0108 x  R²adj = 0.23
−0.2
0.0
0.2
Treatment
High
Low

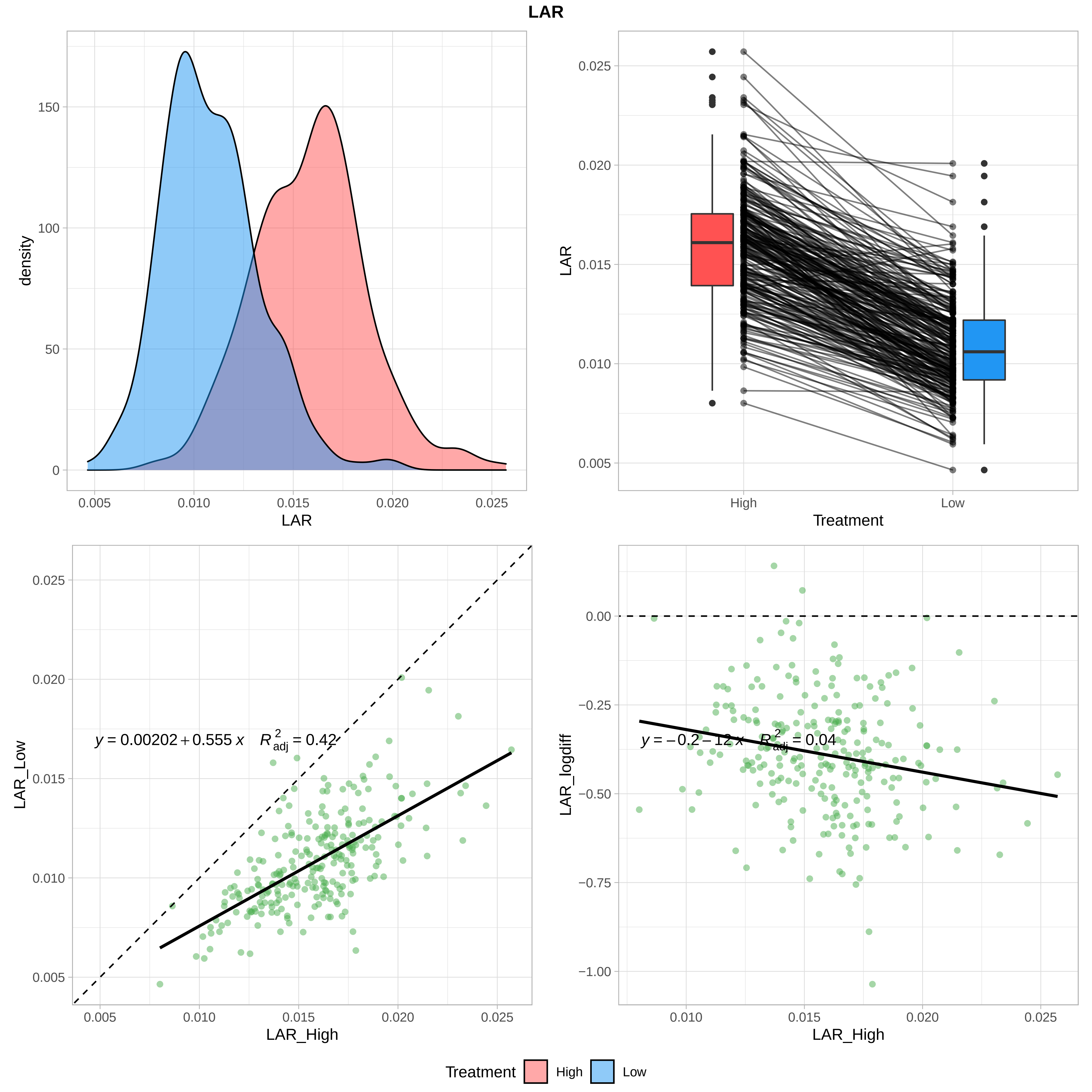

LAR
density
LAR
LAR
Treatment
High
Low
LAR_Low
LAR_High
$y = 0.00202 + 0.555\,x \quad R^2_{adj} = 0.42$
LAR_logdiff
LAR_High
$y = -0.2 - 12\,x \quad R^2_{adj} = 0.04$
Treatment
High
Low

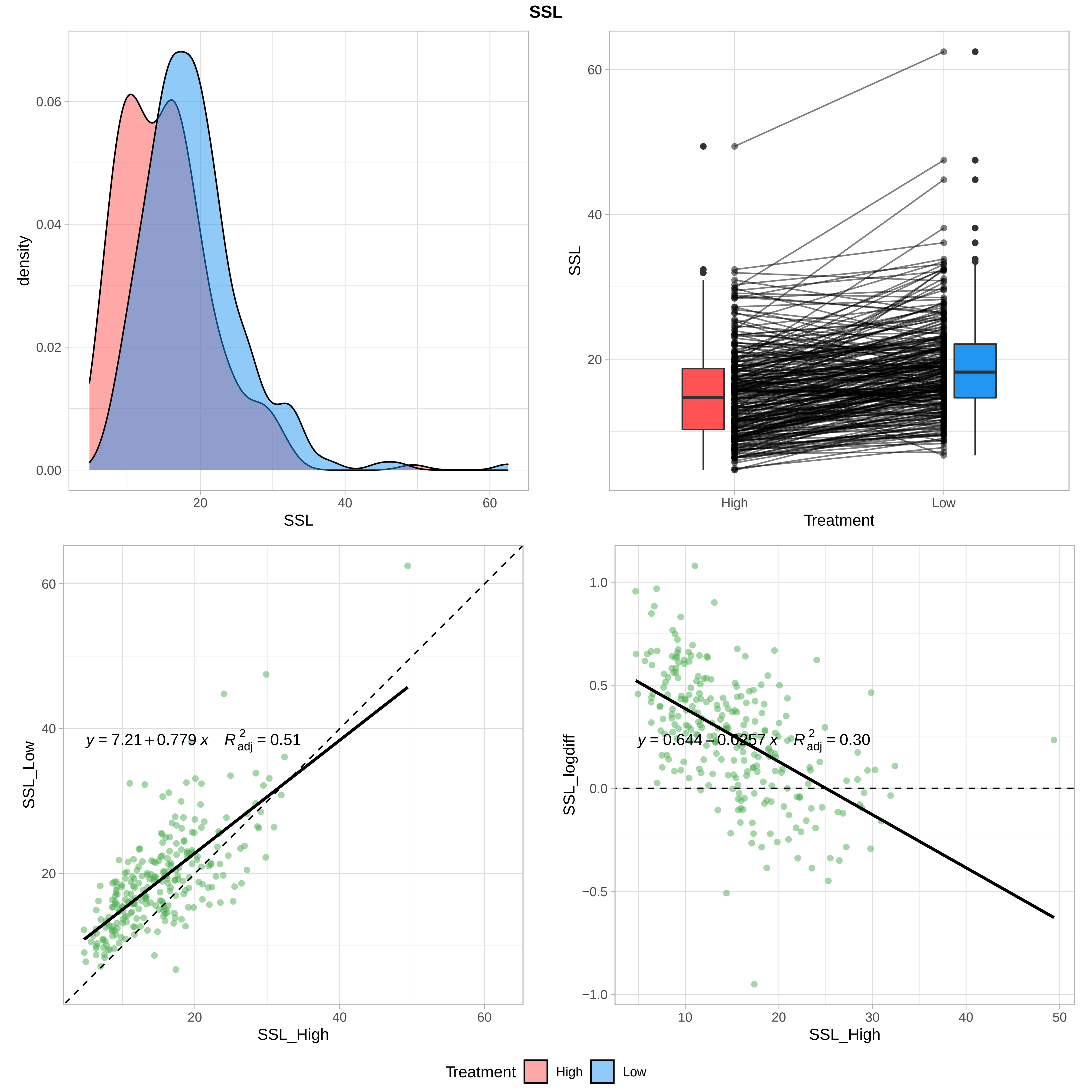

SSL
0.06
0.04
0.02
0.00
density
20
40
60
SSL
60
40
20
SSL
High
Low
Treatment
60
40
20
SSL_Low
y = 7.21 + 0.779 x   R²adj = 0.51
SSL_High
1.0
0.5
0.0
−0.5
−1.0
SSL_logdiff
y = 0.644 − 0.0257 x   R²adj = 0.30
10
30
50
SSL_High
Treatment
High
Low

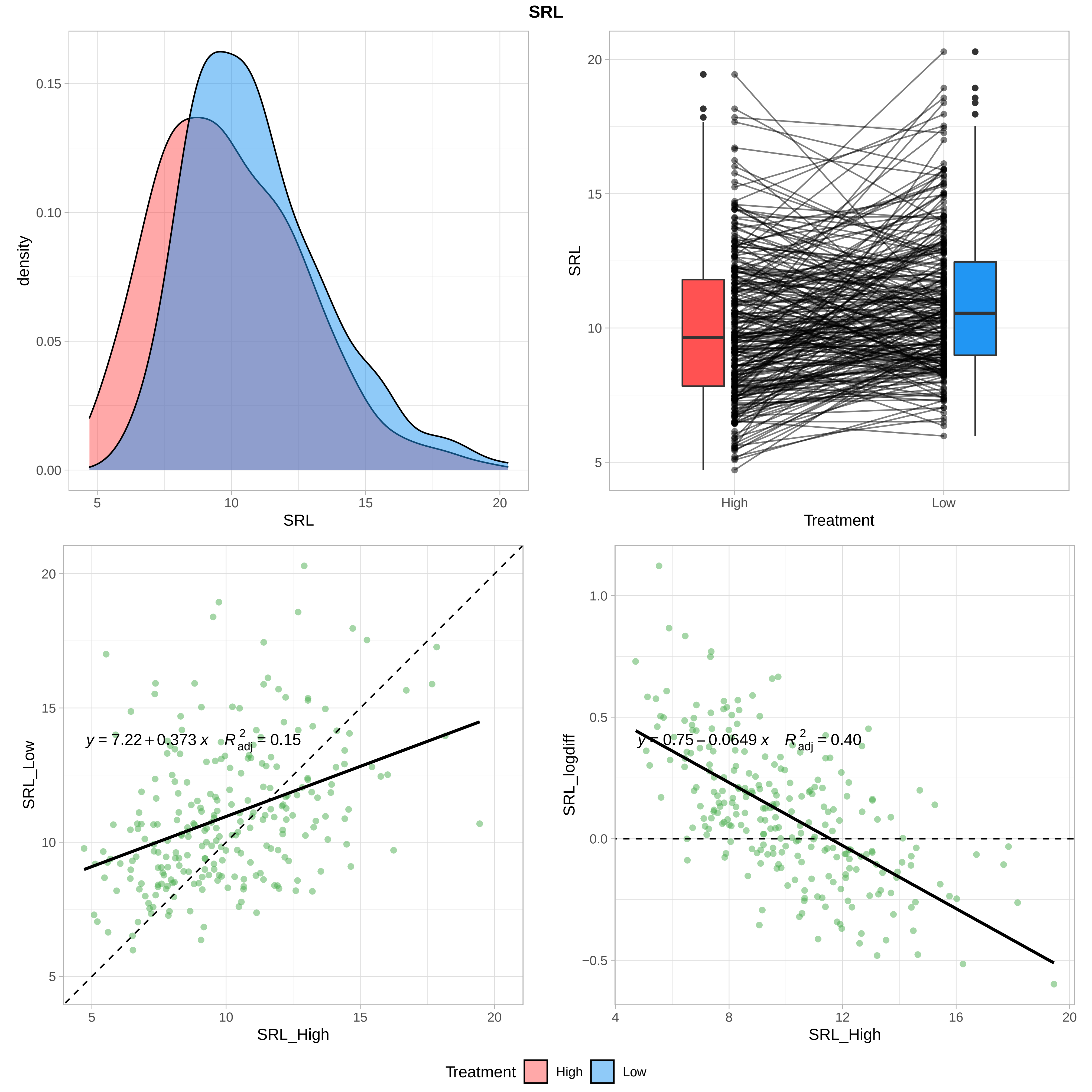

SRL
density
0.00
0.05
0.10
0.15
5
10
15
20
SRL
SRL
High
Low
Treatment
5
10
15
20
SRL_Low
SRL_High
y = 7.22 + 0.373 x   R²adj = 0.15
SRL_logdiff
y = 0.75 − 0.0649 x   R²adj = 0.40
−0.5
0.0
0.5
1.0
4
8
12
16
20
Treatment
High
Low

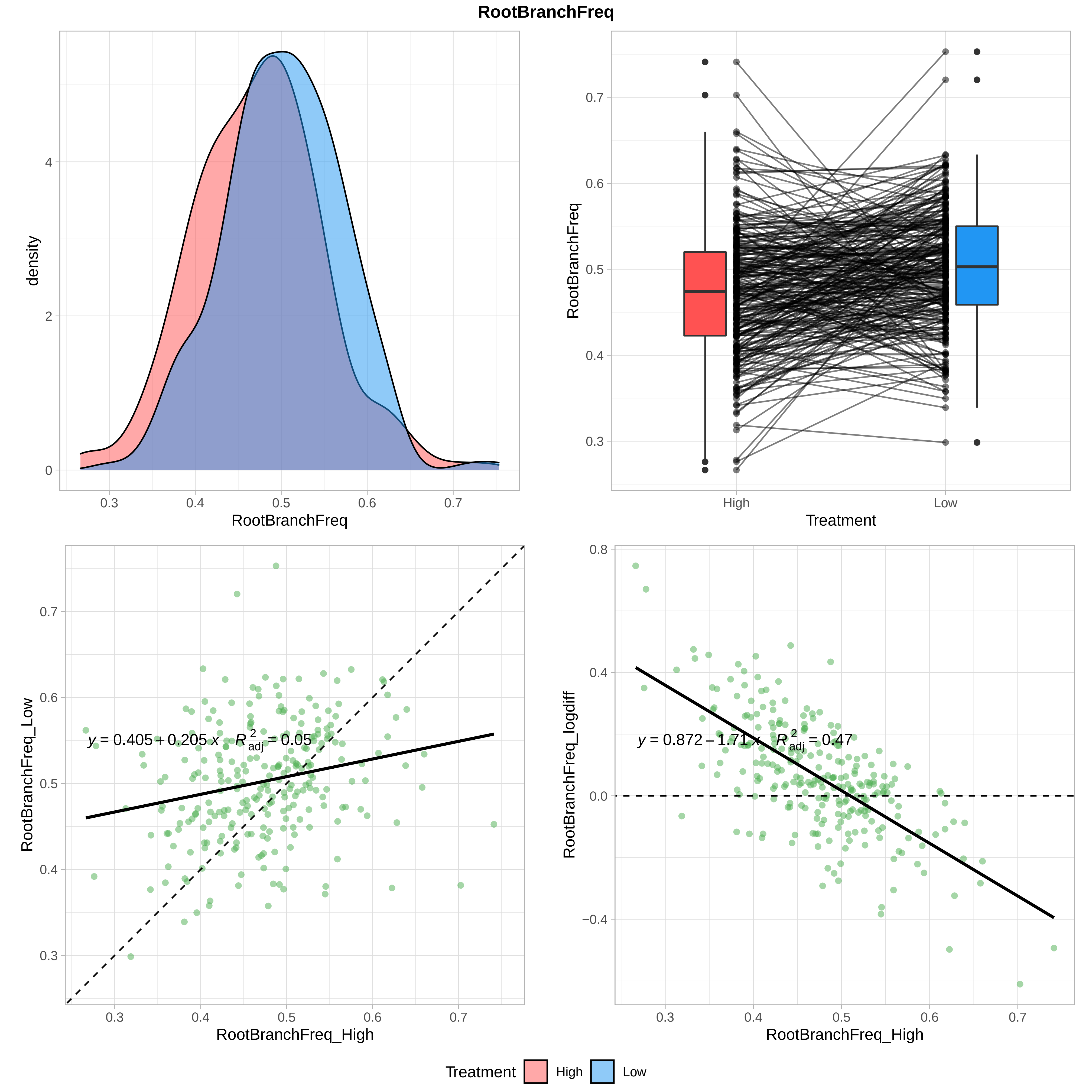
RootBranchFreq
density
RootBranchFreq
RootBranchFreq
Treatment
High
Low
RootBranchFreq_Low
RootBranchFreq_High
y = 0.405 + 0.205 x   R²adj = 0.05
RootBranchFreq_logdiff
RootBranchFreq_High
y = 0.872 − 1.71 x   R²adj = 0.47
Treatment
High
Low

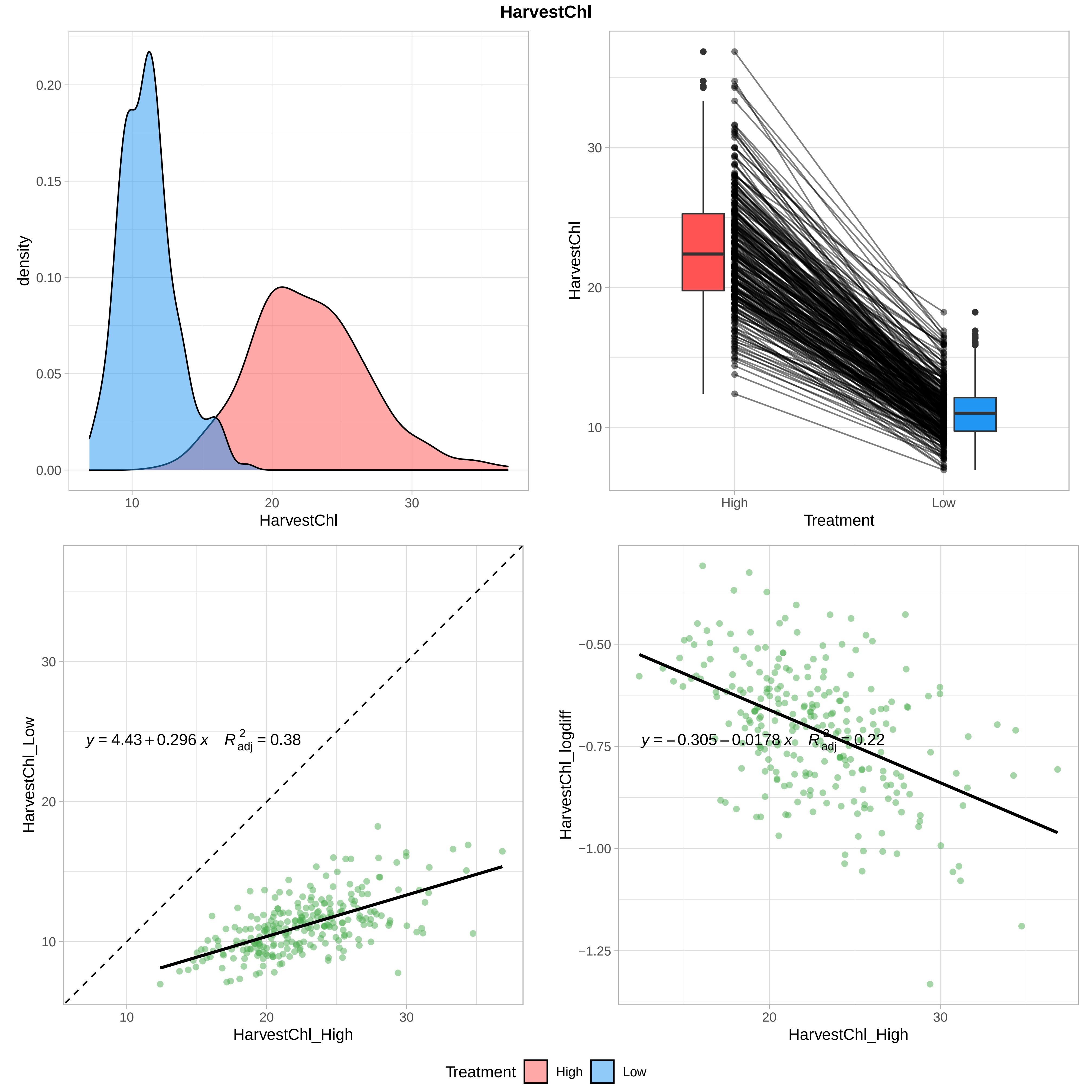
HarvestChl
density
0.00
0.05
0.10
0.15
0.20
10
20
30
HarvestChl
HarvestChl
10
20
30
High
Low
Treatment
HarvestChl_Low
$y = 4.43 + 0.296\,x \quad R^2_{adj} = 0.38$
HarvestChl_High
HarvestChl_logdiff
−0.50
−0.75
−1.00
−1.25
$y = -0.305 - 0.0178\,x \quad R^2_{adj} = 0.22$
Treatment
High
Low

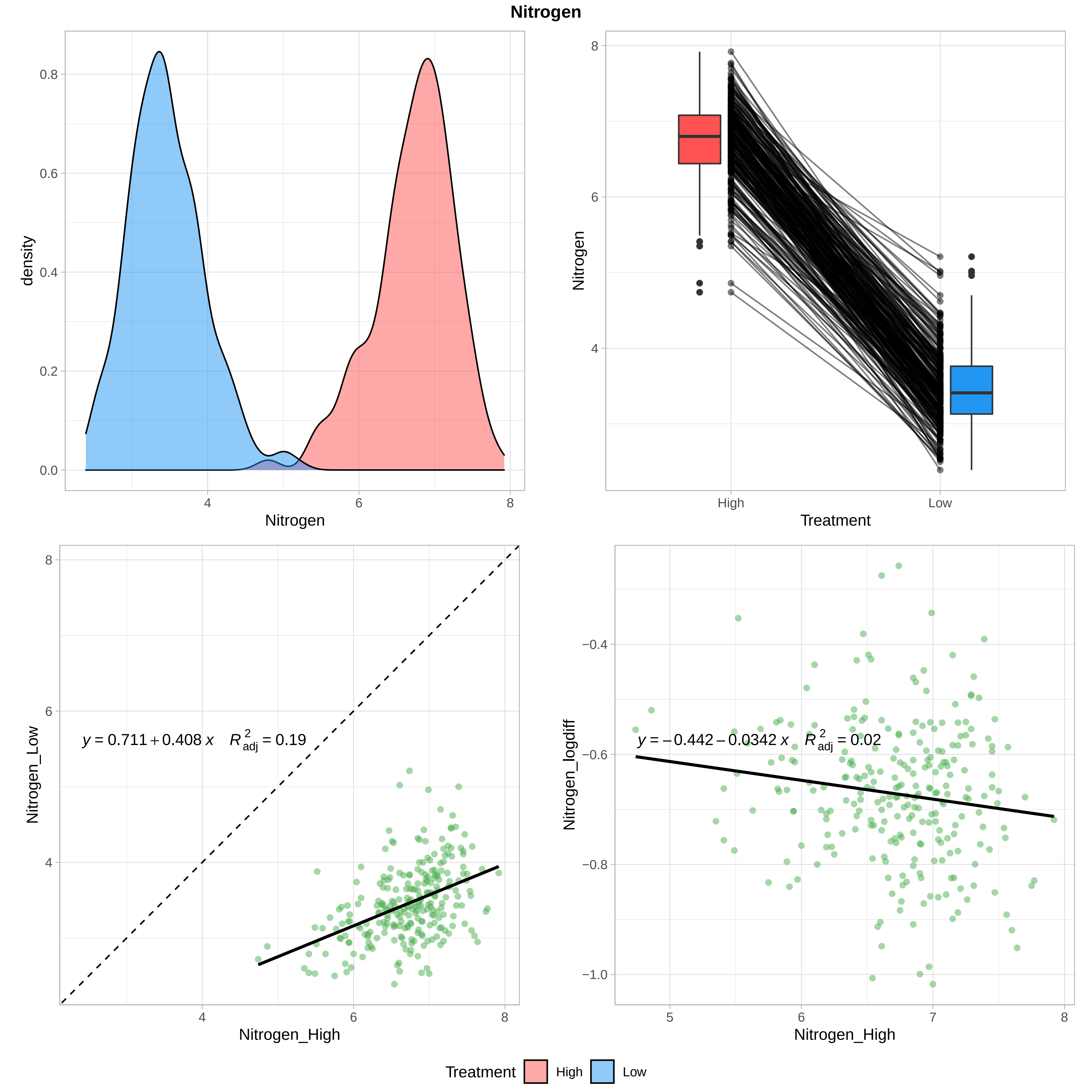
Nitrogen
density
0.0
0.2
0.4
0.6
0.8
4
6
8
Nitrogen
8
6
4
High
Low
Treatment
y = 0.711 + 0.408 x   R²adj = 0.19
Nitrogen_Low
Nitrogen_High
y = −0.442 − 0.0342 x   R²adj = 0.02
Nitrogen_logdiff
−0.4
−0.6
−0.8
−1.0
5
7
Treatment
High
Low

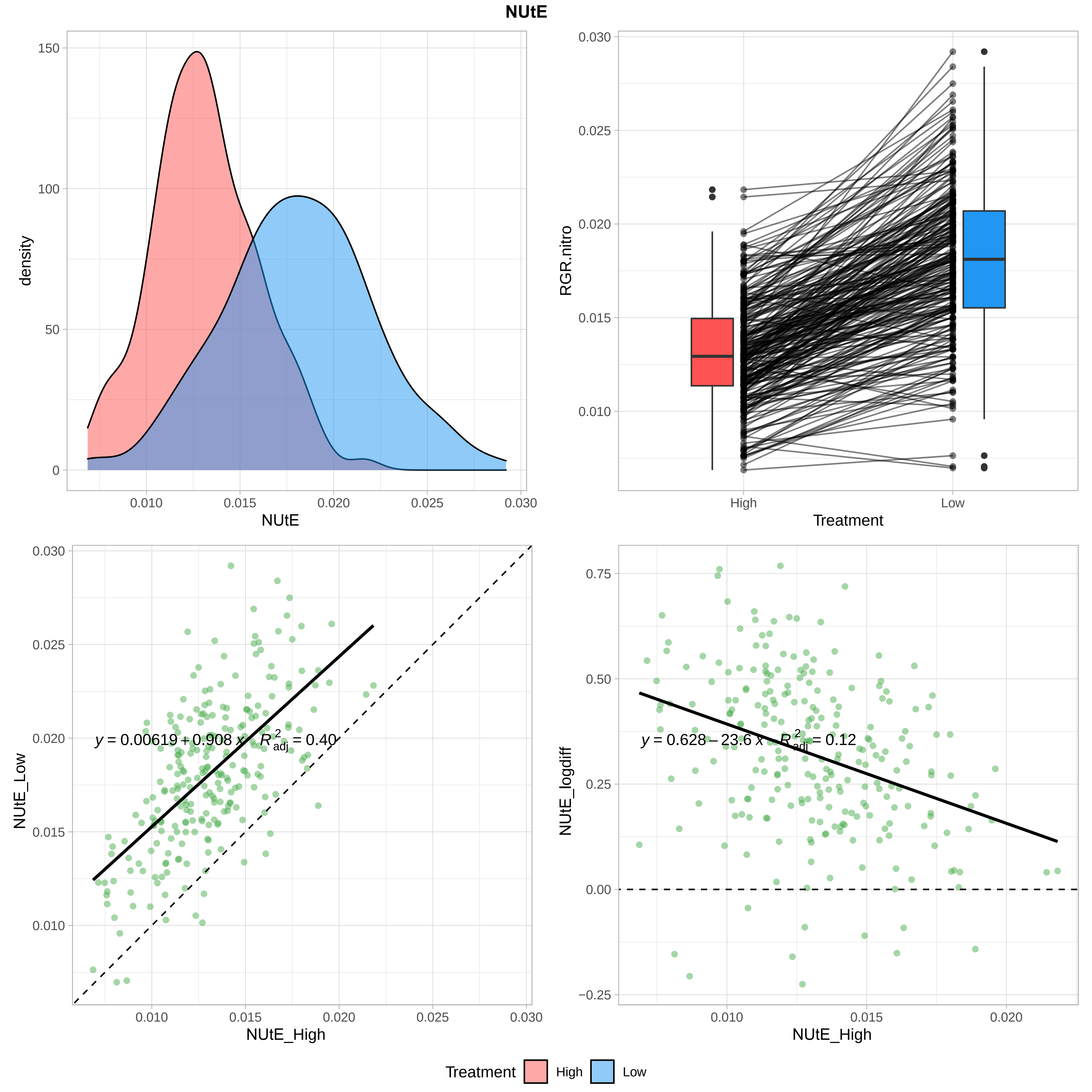

NUtE
density
NUtE
RGR.nitro
Treatment
High
Low
NUtE_Low
NUtE_High
y = 0.00619 + 0.908 x   R²adj = 0.40
NUtE_logdiff
y = 0.628 − 23.6 x   R²adj = 0.12
Treatment High Low

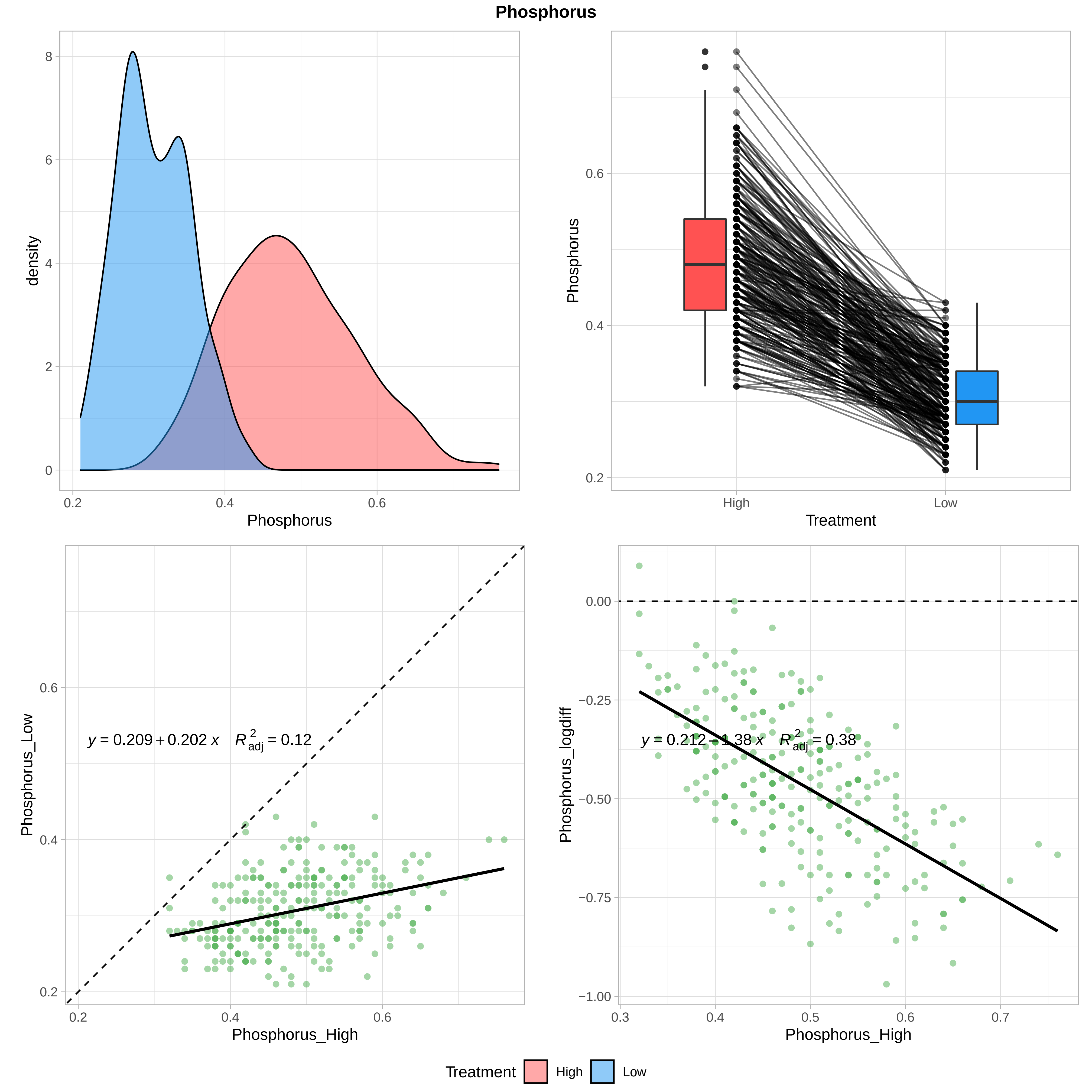
Phosphorus
density
Phosphorus
Treatment
High
Low
Phosphorus_Low
Phosphorus_High
$y = 0.209 + 0.202\,x \quad R^2_{adj} = 0.12$
Phosphorus_logdiff
$y = 0.212 - 1.38\,x \quad R^2_{adj} = 0.38$
Treatment High Low

### Potassium

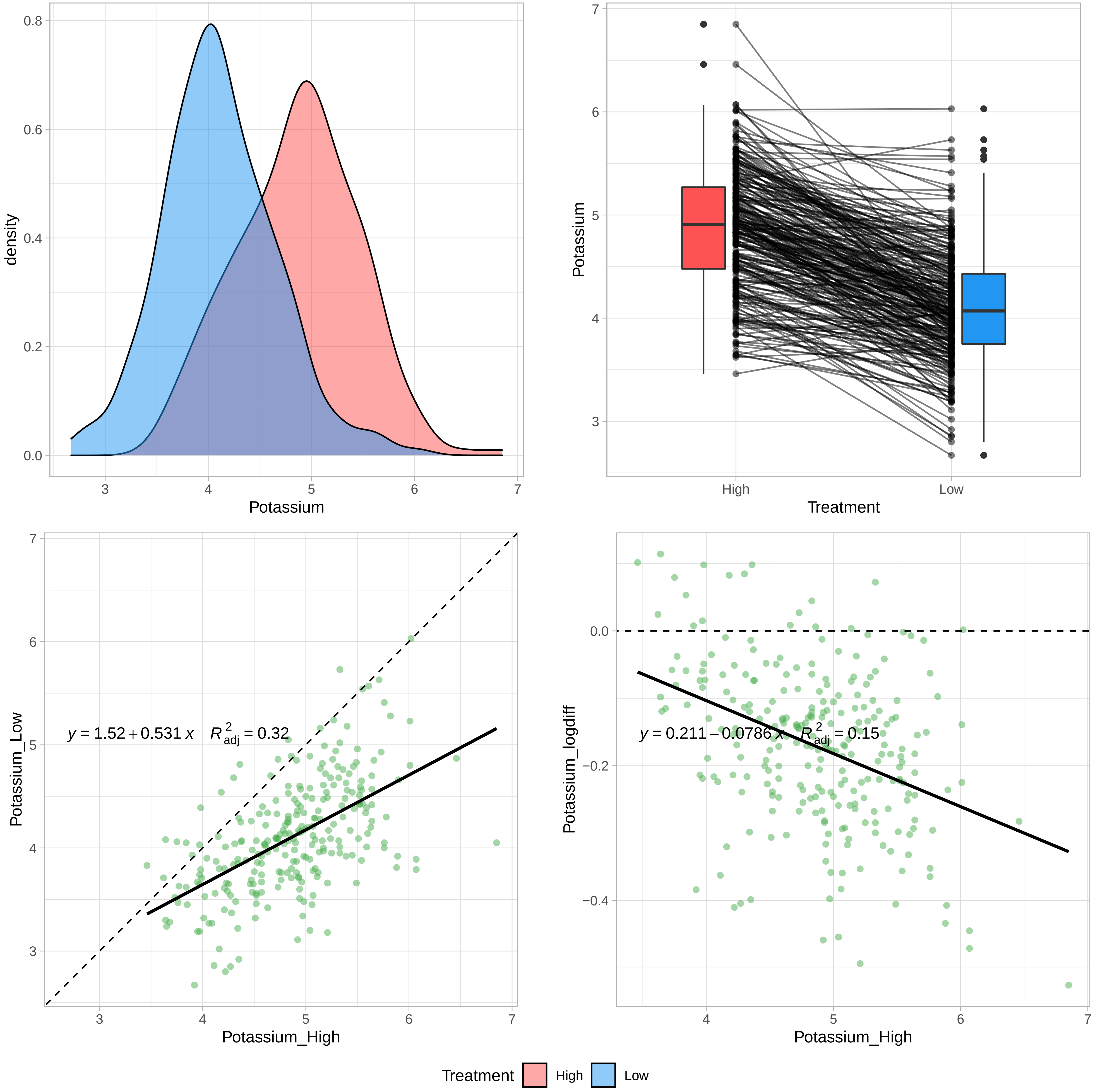
density
Potassium
Treatment
High
Low
Potassium_Low
Potassium_High
y = 1.52 + 0.531 x    R2adj = 0.32
Potassium_logdiff
y = 0.211 − 0.0786 x    R2adj = 0.15
Treatment High Low

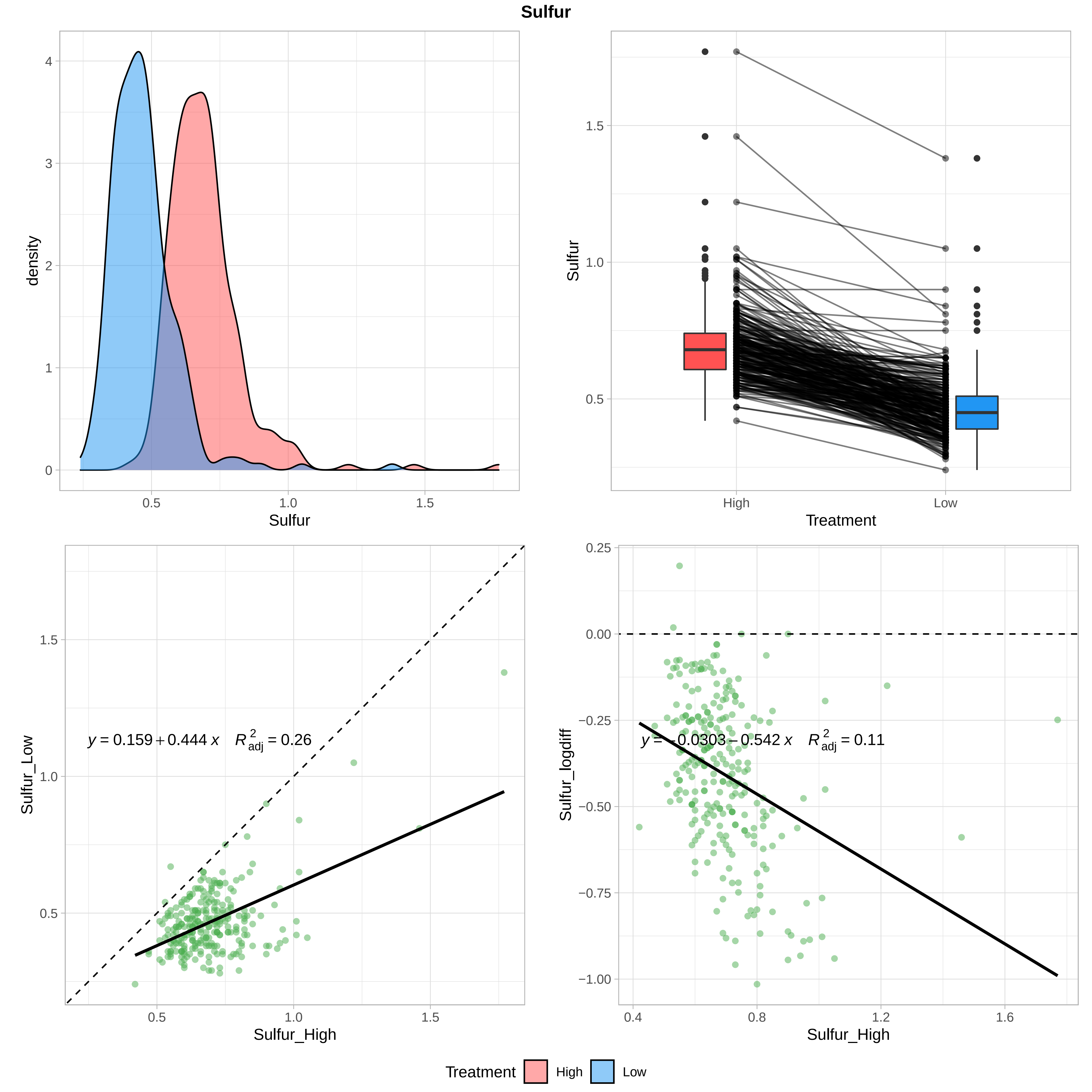
Sulfur
density
Sulfur
Treatment
High
Low
Sulfur_Low
Sulfur_High
$y = 0.159 + 0.444\,x \quad R^2_{adj} = 0.26$
Sulfur_logdiff
$y = -0.0303 - 0.542\,x \quad R^2_{adj} = 0.11$
Treatment High Low

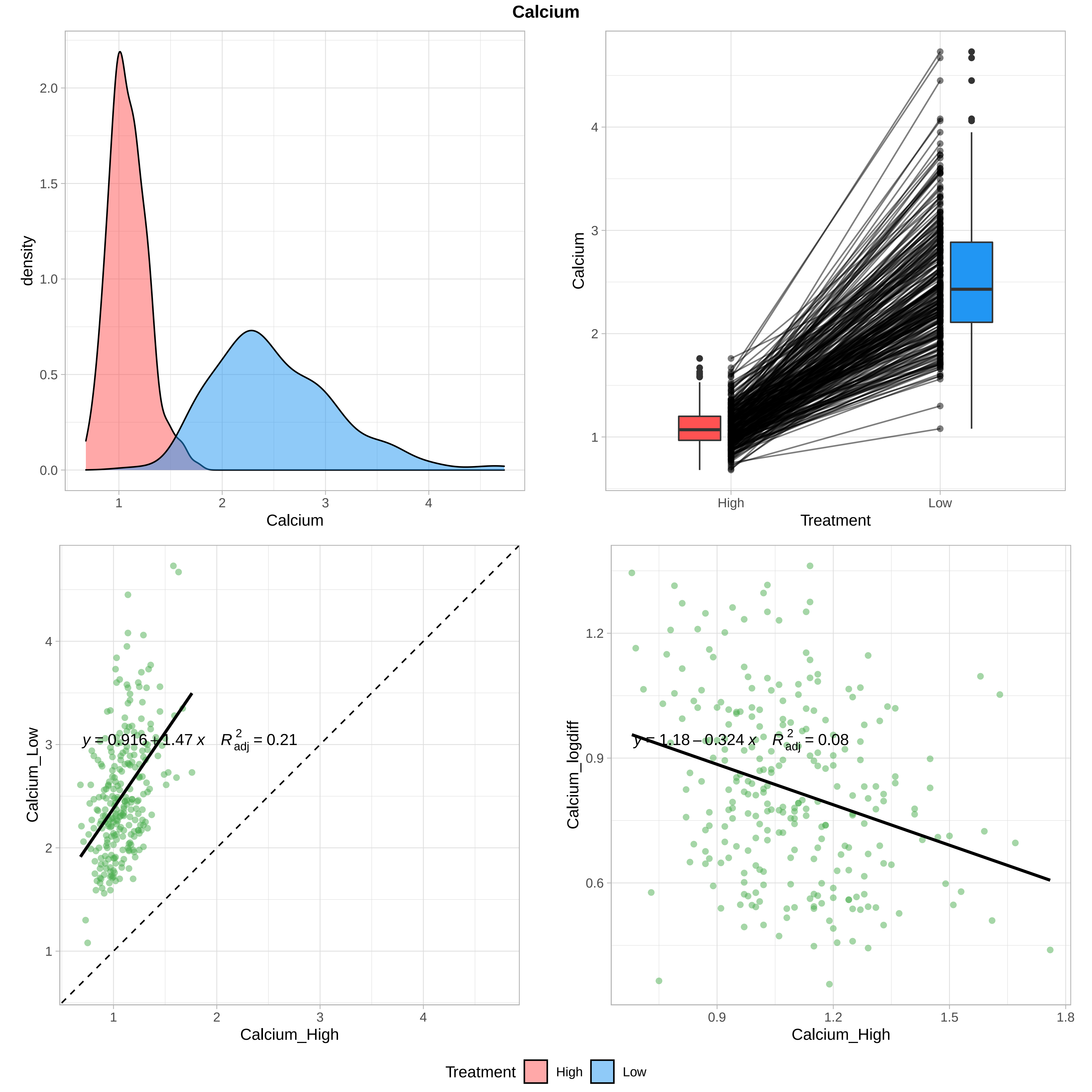

Calcium
density
Calcium
Treatment
High
Low
Calcium_High
Calcium_Low
$y = 0.916 + 1.47\,x \quad R^2_{adj} = 0.21$
Calcium_logdiff
$y = 1.18 - 0.324\,x \quad R^2_{adj} = 0.08$
Treatment High Low

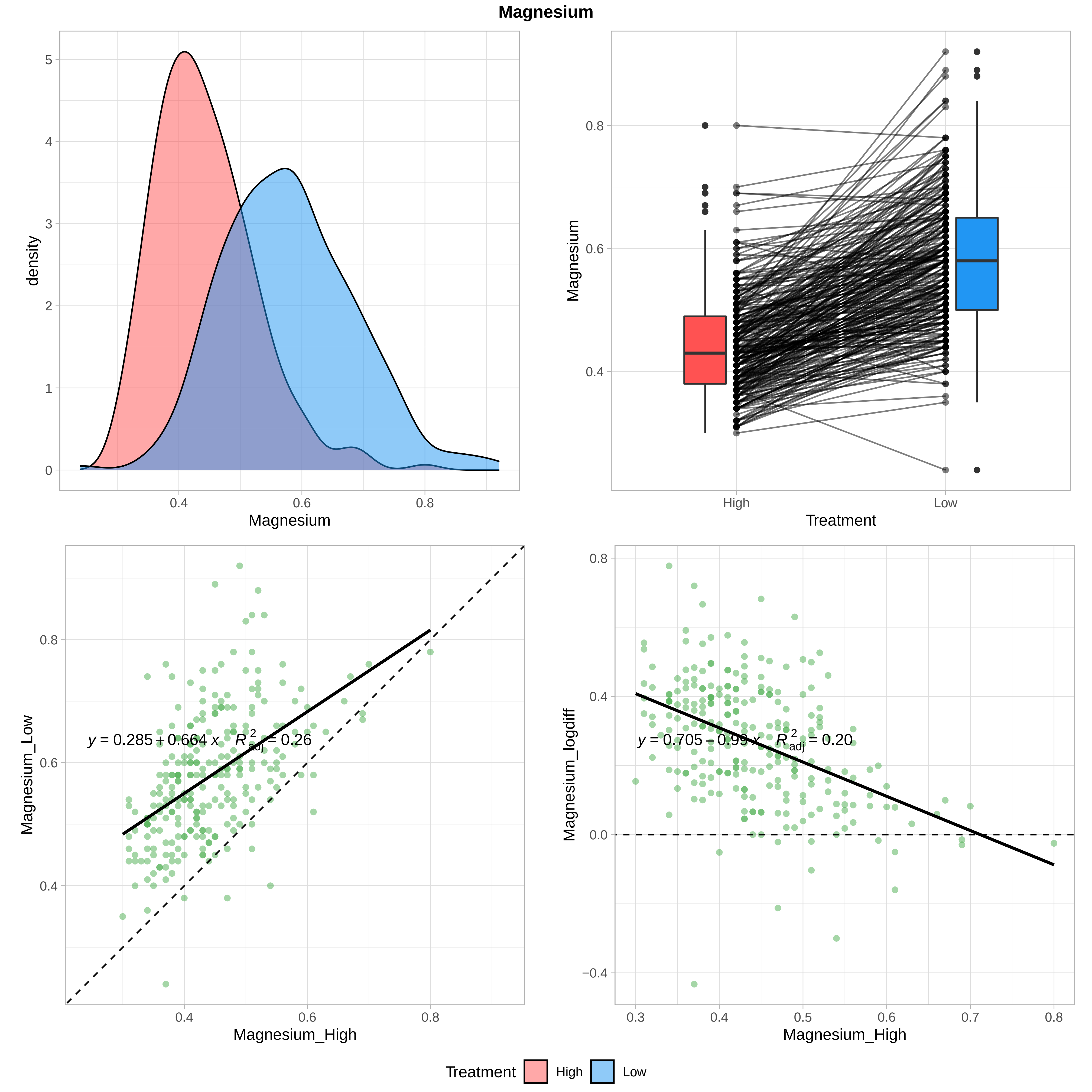

Magnesium
density
Magnesium
0.4
0.6
0.8
0
1
2
3
4
5
Magnesium
Treatment
High
Low
0.4
0.6
0.8
Magnesium_Low
Magnesium_High
y = 0.285 + 0.664 x   R²adj = 0.26
Magnesium_logdiff
y = 0.705 − 0.99 x   R²adj = 0.20
0.3
0.5
0.7
−0.4
0.0
Treatment
High
Low

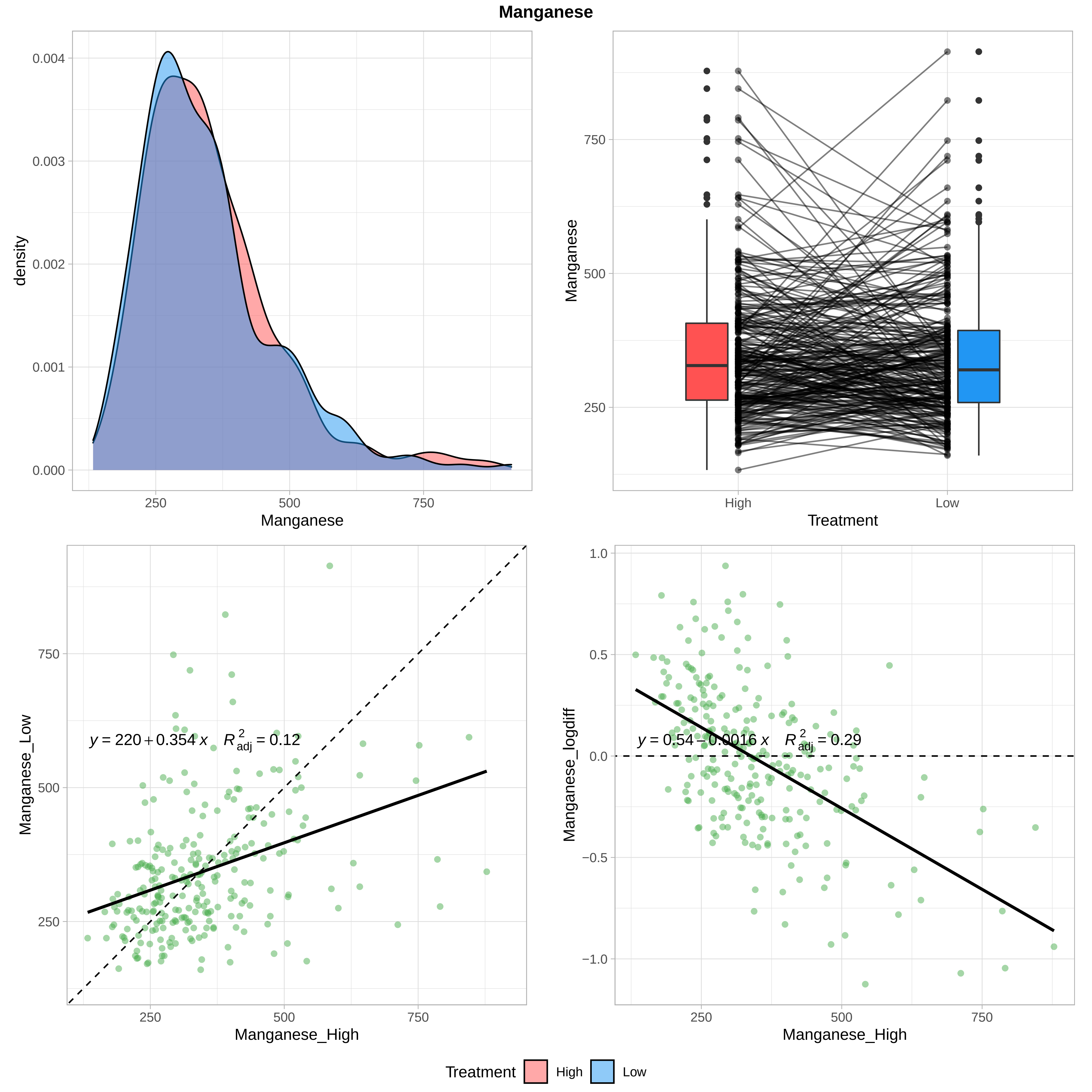
Manganese
density
0.000
0.001
0.002
0.003
0.004
250
500
750
Manganese
Manganese
250
500
750
High
Low
Treatment
Manganese_Low
y = 220 + 0.354 x  R²adj = 0.12
250
500
750
Manganese_High
Manganese_logdiff
y = 0.54 − 0.0016 x  R²adj = 0.29
1.0
0.5
0.0
−0.5
−1.0
250
500
750
Manganese_High
Treatment
High
Low

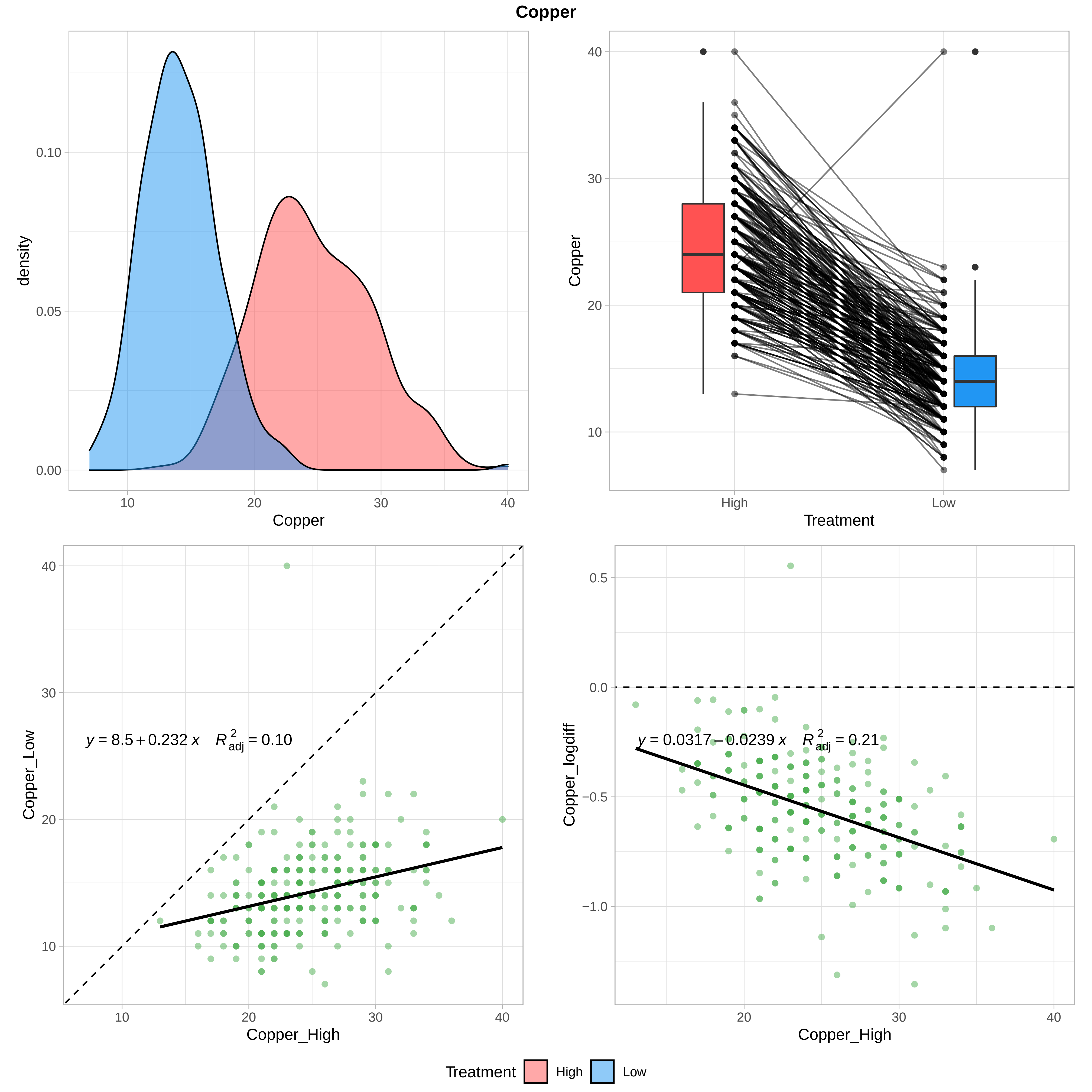

Copper
density
0.10
0.05
0.00
10
20
30
40
Copper
Copper
40
30
20
10
High
Low
Treatment
Copper_Low
y = 8.5 + 0.232 x   R²adj = 0.10
Copper_High
Copper_logdiff
0.5
0.0
−0.5
−1.0
y = 0.0317 − 0.0239 x   R²adj = 0.21
Copper_High
Treatment
High
Low

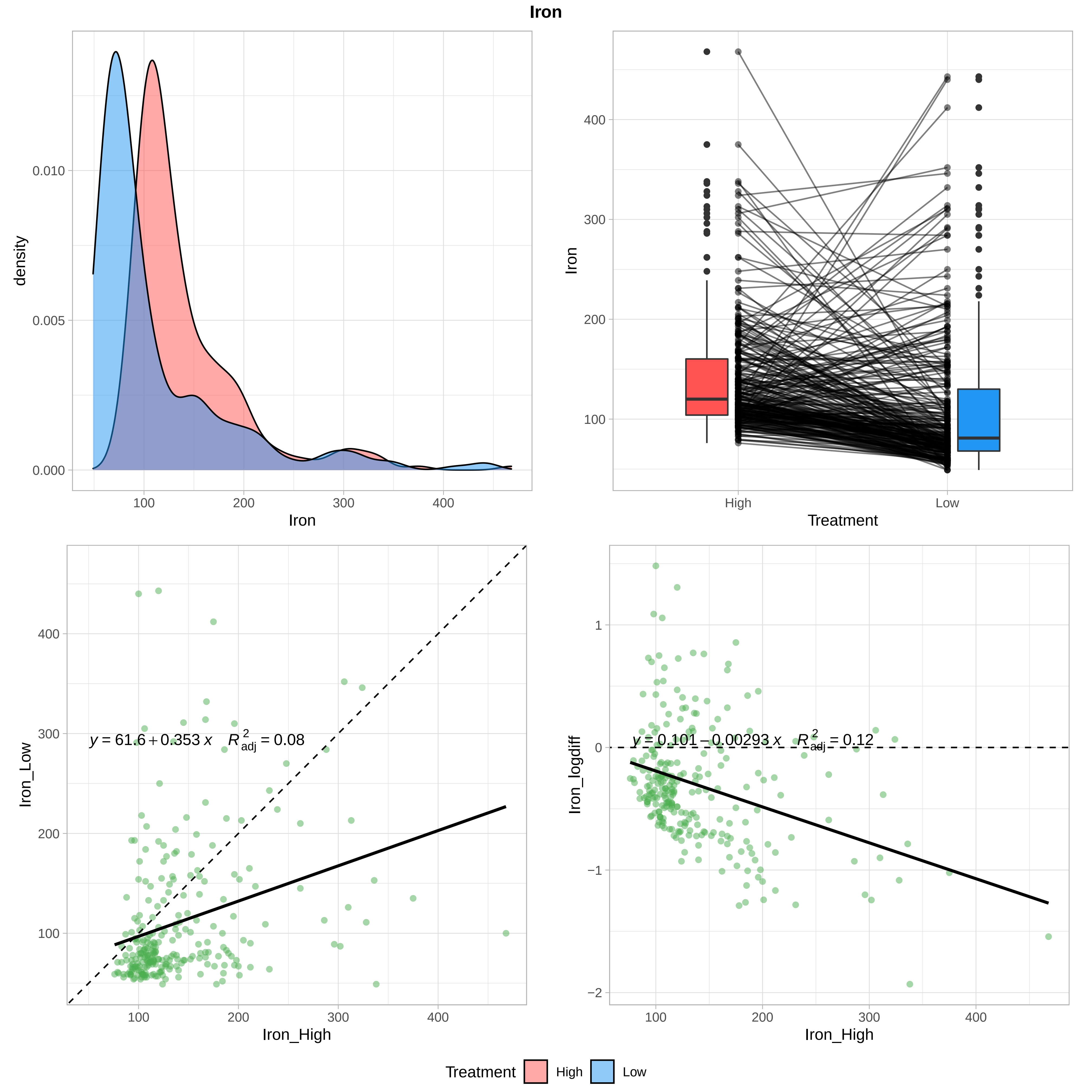
Iron
density
0.010
0.005
0.000
100
200
300
400
Iron
400
300
200
100
High
Low
Treatment
y = 61.6 + 0.353 x   R²adj = 0.08
Iron_Low
Iron_High
y = 0.101 − 0.00293 x   R²adj = 0.12
Iron_logdiff
1
0
−1
−2
Treatment
High
Low

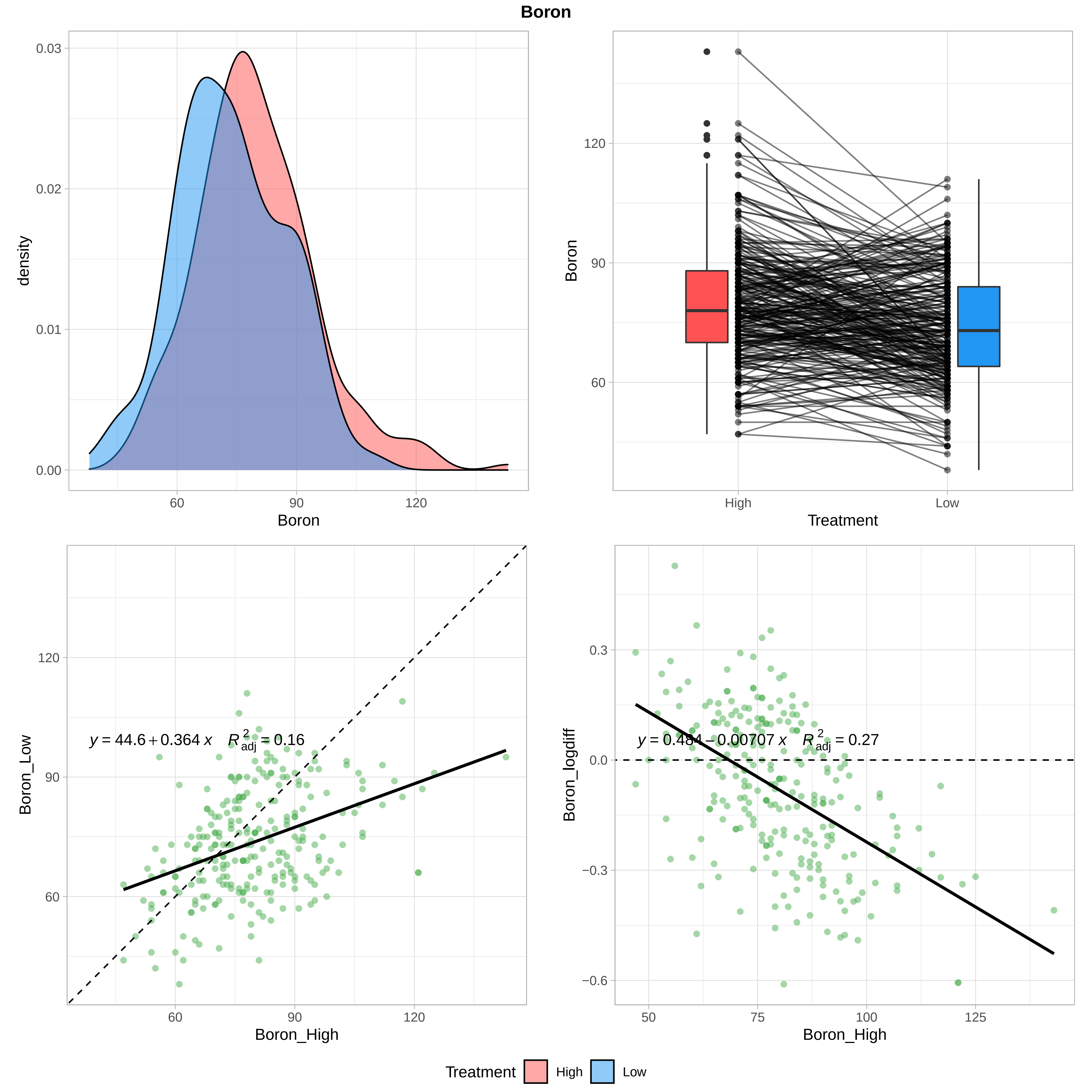

Boron
density
0.03
0.02
0.01
0.00
60
90
120
Boron
Boron
120
90
60
High
Low
Treatment
y = 44.6 + 0.364 x    R²adj = 0.16
Boron_Low
Boron_High
y = 0.484 − 0.00707 x    R²adj = 0.27
Boron_logdiff
0.3
0.0
−0.3
−0.6
50
75
100
125
Treatment
High
Low

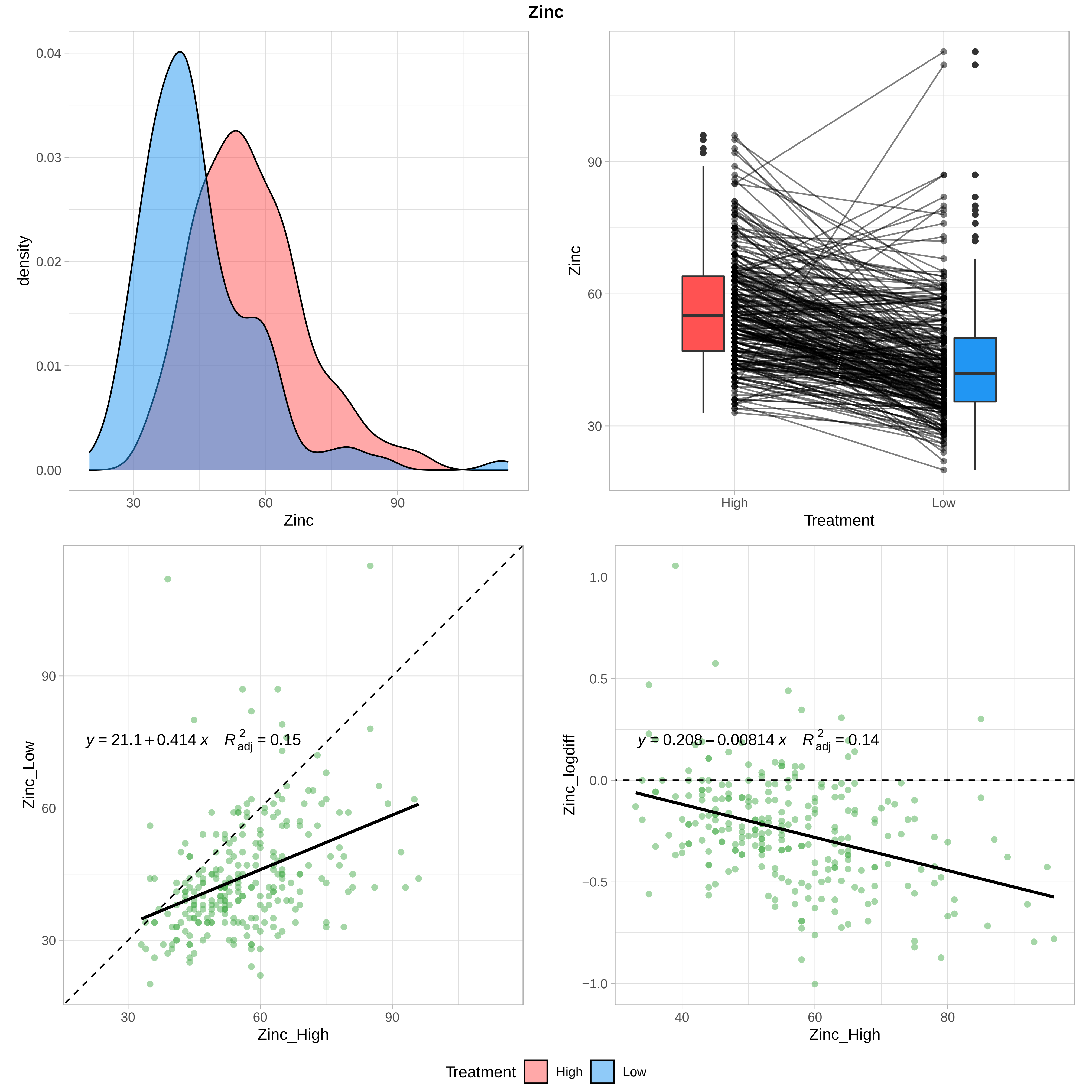

Zinc
density
0.04
0.03
0.02
0.01
0.00
30
60
90
Zinc
Zinc
90
60
30
High
Low
Treatment
Zinc_Low
90
60
30
$y = 21.1 + 0.414\,x \quad R^2_{adj} = 0.15$
Zinc_High
Zinc_logdiff
1.0
0.5
0.0
−0.5
−1.0
$y = 0.208 - 0.00814\,x \quad R^2_{adj} = 0.14$
40
60
80
Zinc_High
Treatment
High
Low

Figure S2 **Whole leaf calcium and magnesium amount.** (a, c) density plot of total leaf calcium and magnesium content (multiplying leaf weight by concentration) under control and low nutrient treatment. (b, d) Boxplots of trait values in control (red) and low nutrients (blue). Individual are shown as black circles and connected by black lines between treatments.

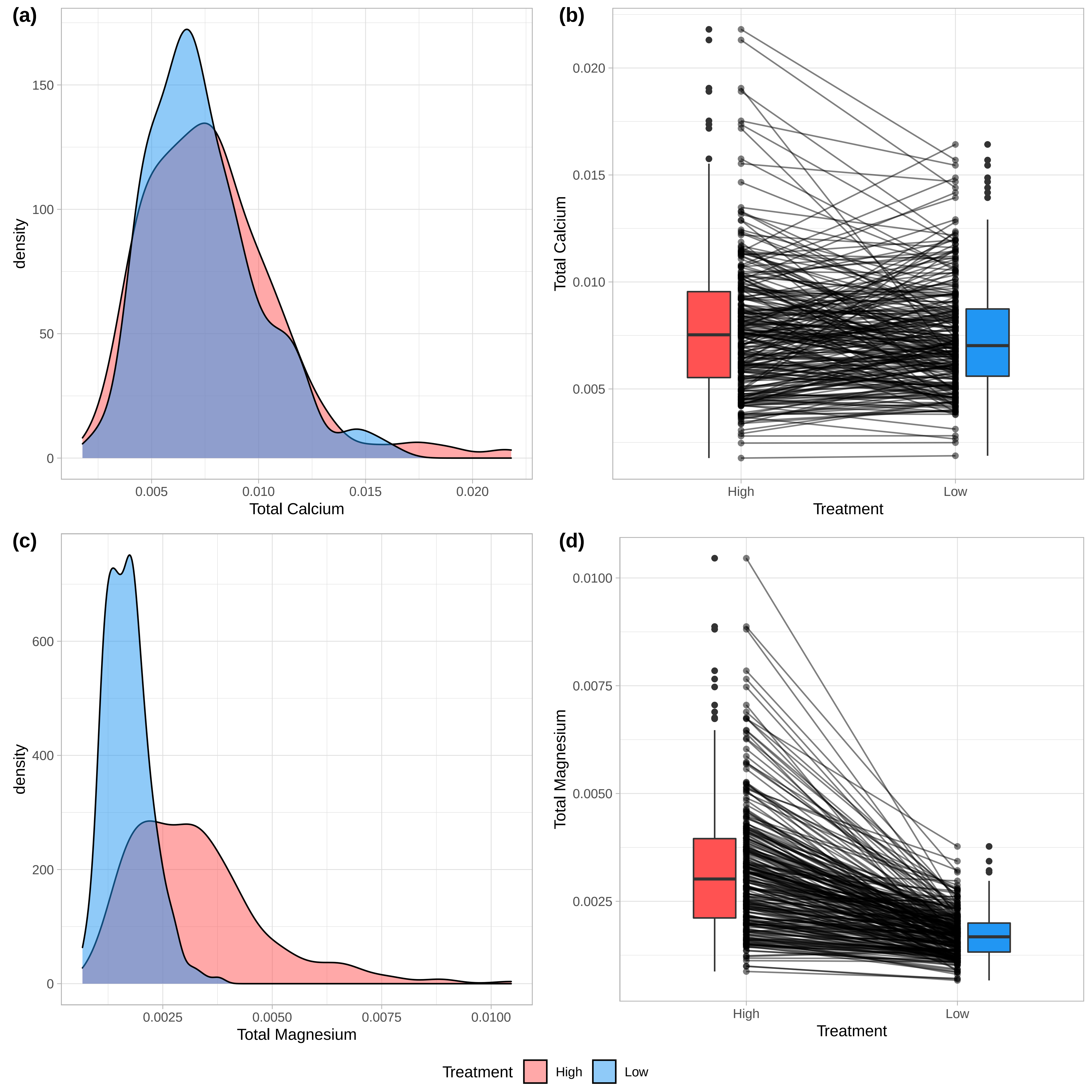

(a)
density
Total Calcium
0
50
100
150
0.005
0.010
0.015
0.020
(b)
Total Calcium
0.005
0.010
0.015
0.020
High
Low
Treatment
(c)
density
Total Magnesium
0
200
400
600
0.0025
0.0050
0.0075
0.0100
(d)
Total Magnesium
0.0025
0.0050
0.0075
0.0100
High
Low
Treatment
Treatment
High
Low

Figure S3 **All PCAs (companion to table 3).** Principal component analysis of variation in putatively size-independent traits (chlorophyll content, fine root allocation [mass fraction and root fraction], LMA, LMF, RMF, SMF, tap root allocation [mass fraction and root fraction]) and elemental traits (B, Ca, Cu, Fe, Mg, Mn, N, P, K, S, Zn) under control (high nutrients) and low nutrient (10% of control) conditions, as well as the plasticity in trait values between treatments.

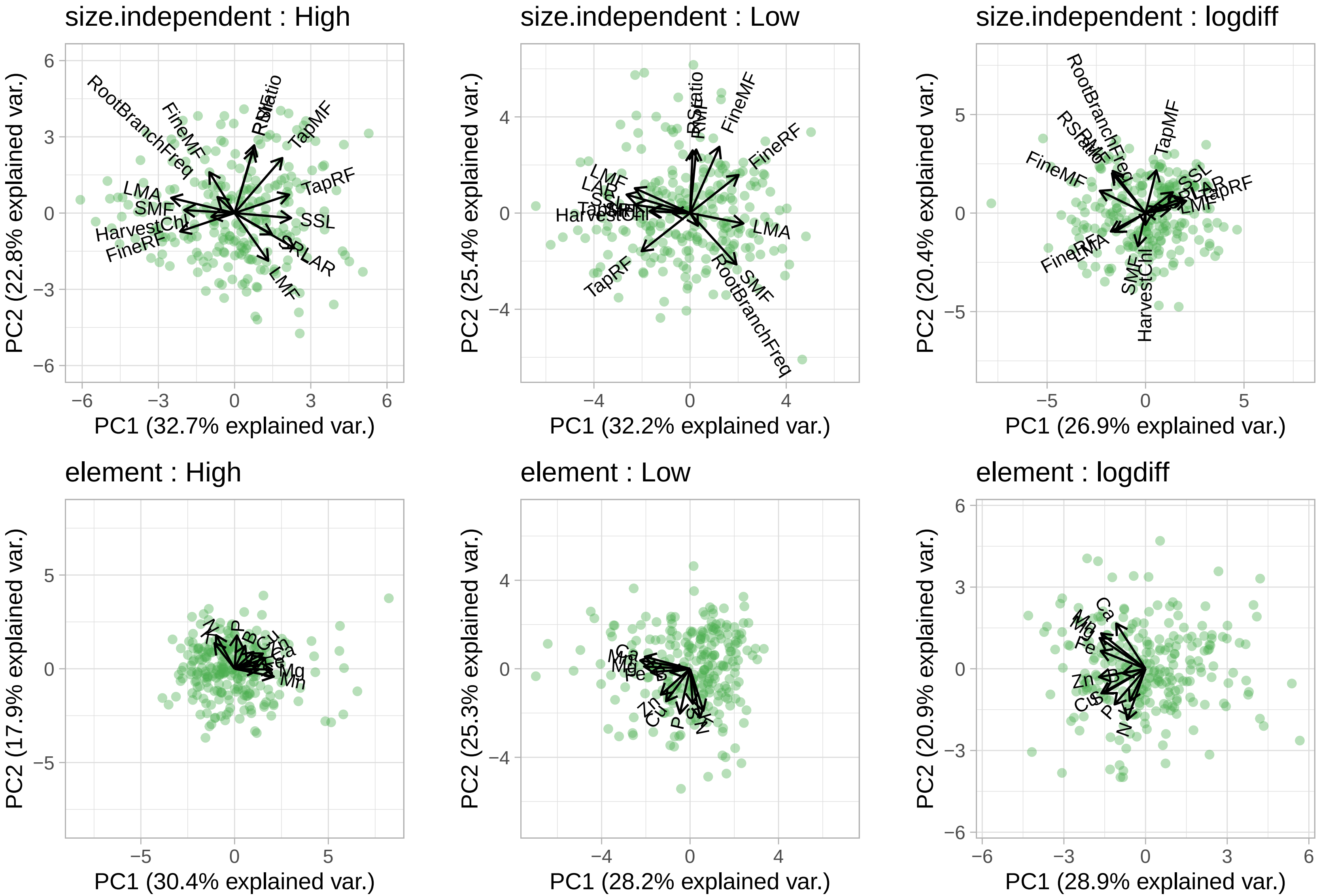
size.independent : High
PC1 (32.7% explained var.)
PC2 (22.8% explained var.)
size.independent : Low
PC1 (32.2% explained var.)
PC2 (25.4% explained var.)
size.independent : logdiff
PC1 (26.9% explained var.)
PC2 (20.4% explained var.)
element : High
PC1 (30.4% explained var.)
PC2 (17.9% explained var.)
element : Low
PC1 (28.2% explained var.)
PC2 (25.3% explained var.)
element : logdiff
PC1 (28.9% explained var.)
PC2 (20.9% explained var.)
RootBranchFreq
FineMF
FineRF
TapMF
TapRF
LMA
SMF
LMF
SSL
SRLAR
LAR
HarvestChl
RSRatio
RMF
Ca
Mn
Mg
Fe
Zn
Cu
B
P
S
N
K

Figure S4 **All Manhattans**. Per-trait Manhattan plots are shown for trait values under control and low nutrient stressed conditions, as well as the plasticity between treatments (listed as log difference). SNPs above the red line are significant after multiple comparison correction, SNPs above the blue line are in the top 0.1% of P-values. SNPs are colored by the “significant” haplotype regions per chromosome as displayed in Fig. S7.

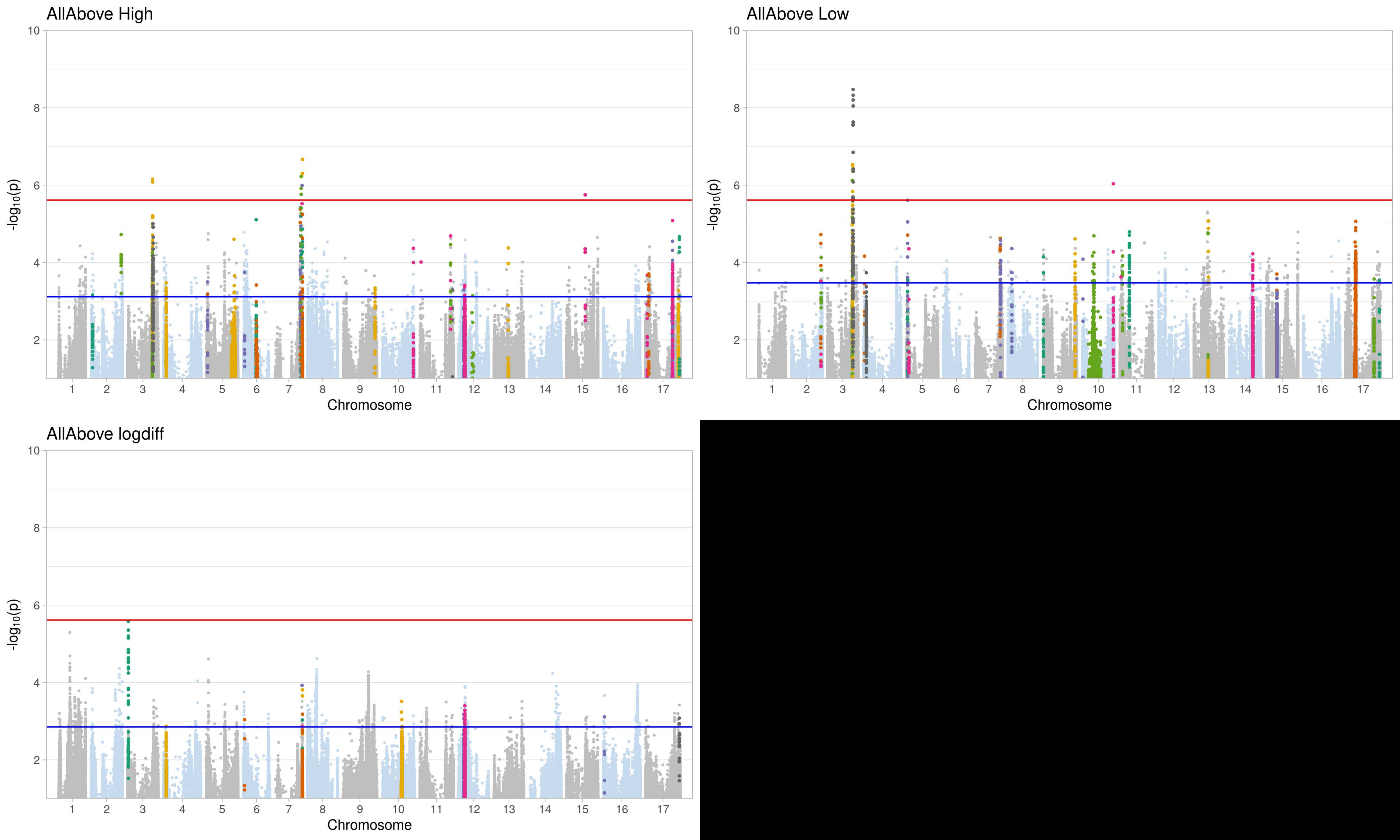
AllAbove High
-log10(p)
Chromosome
AllAbove Low
-log10(p)
Chromosome
AllAbove logdiff
-log10(p)
Chromosome

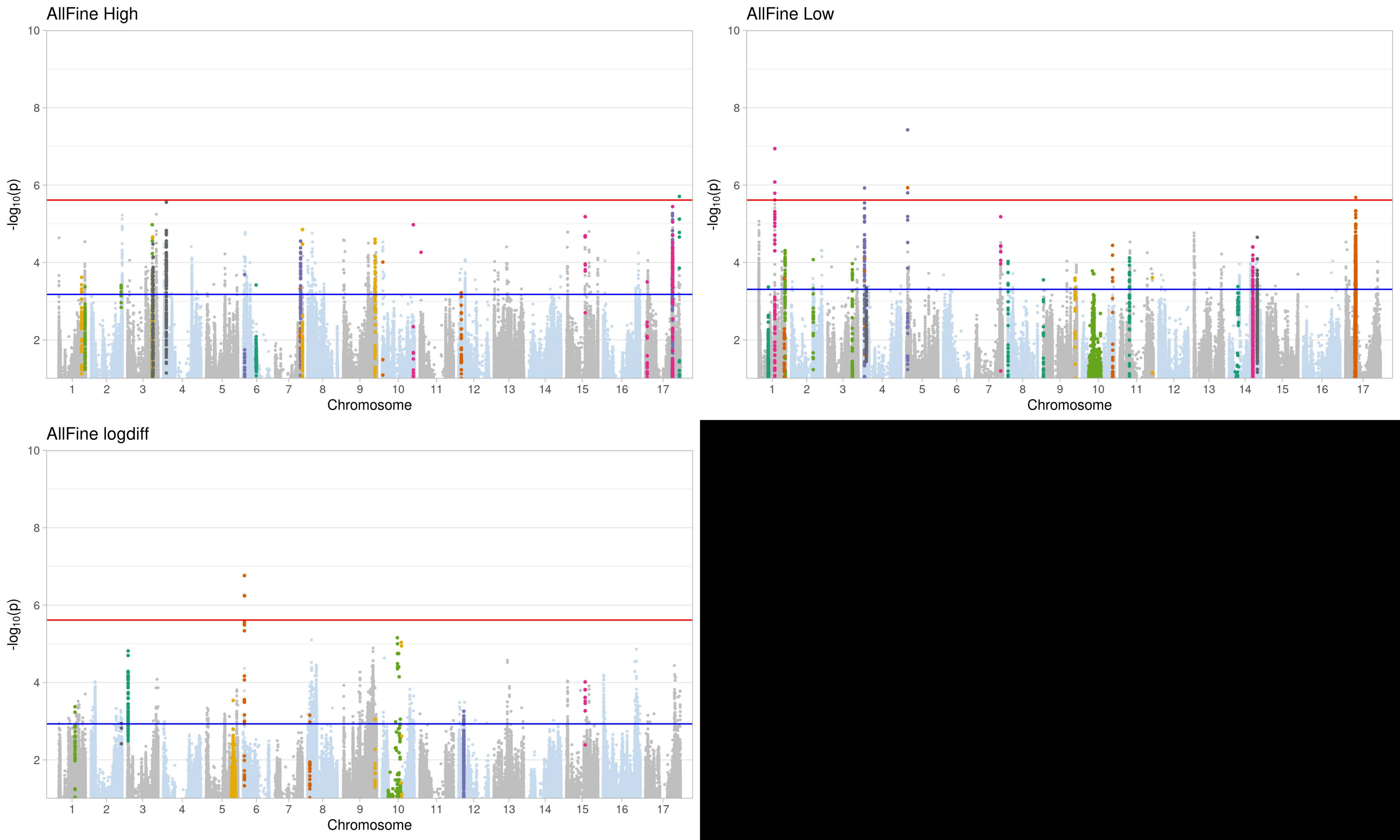

AllFine High
AllFine Low
AllFine logdiff
-log10(p)
Chromosome

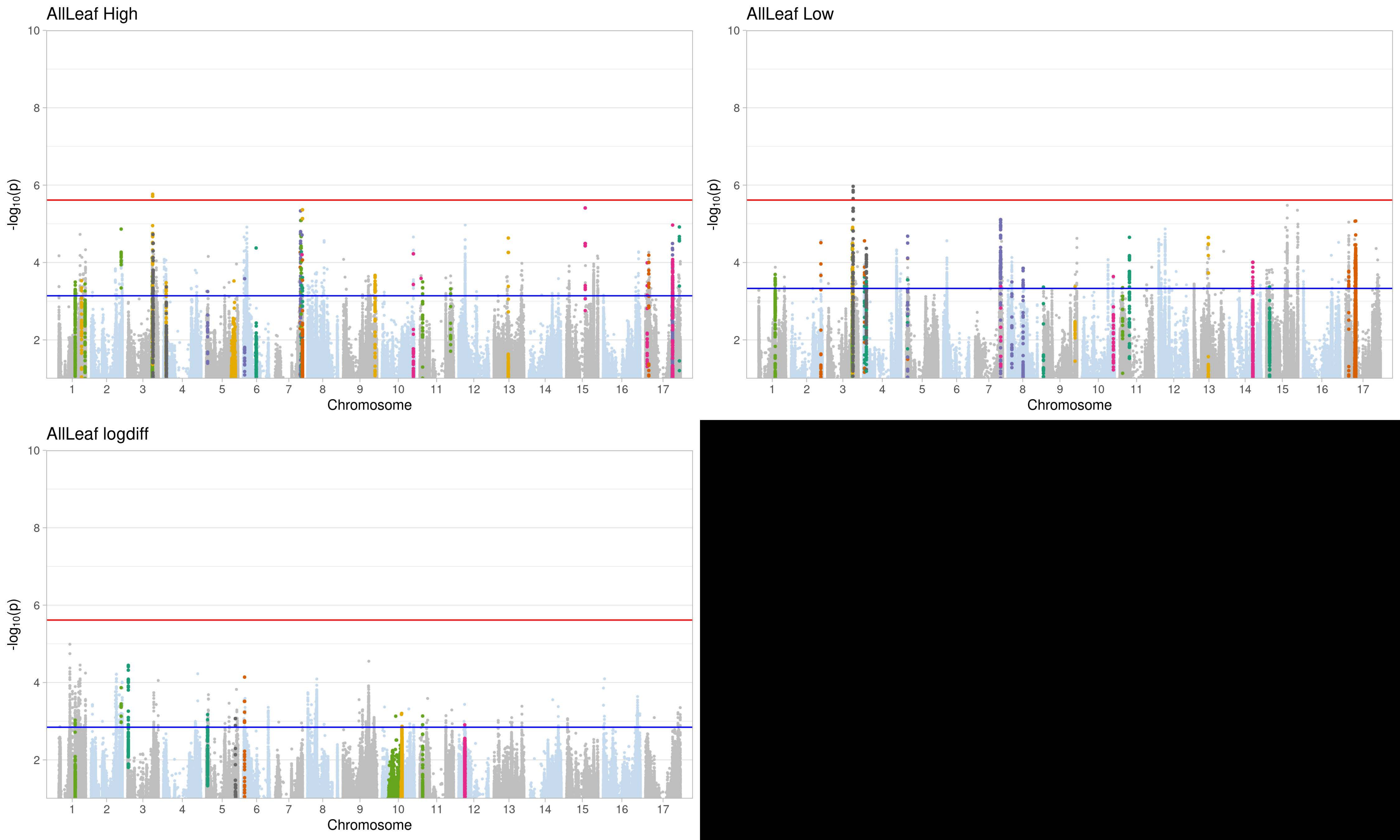
AllLeaf High
AllLeaf Low
AllLeaf logdiff
-log10(p)
Chromosome
10
8
6
4
2
1 2 3 4 5 6 7 8 9 10 11 12 13 14 15 16 17

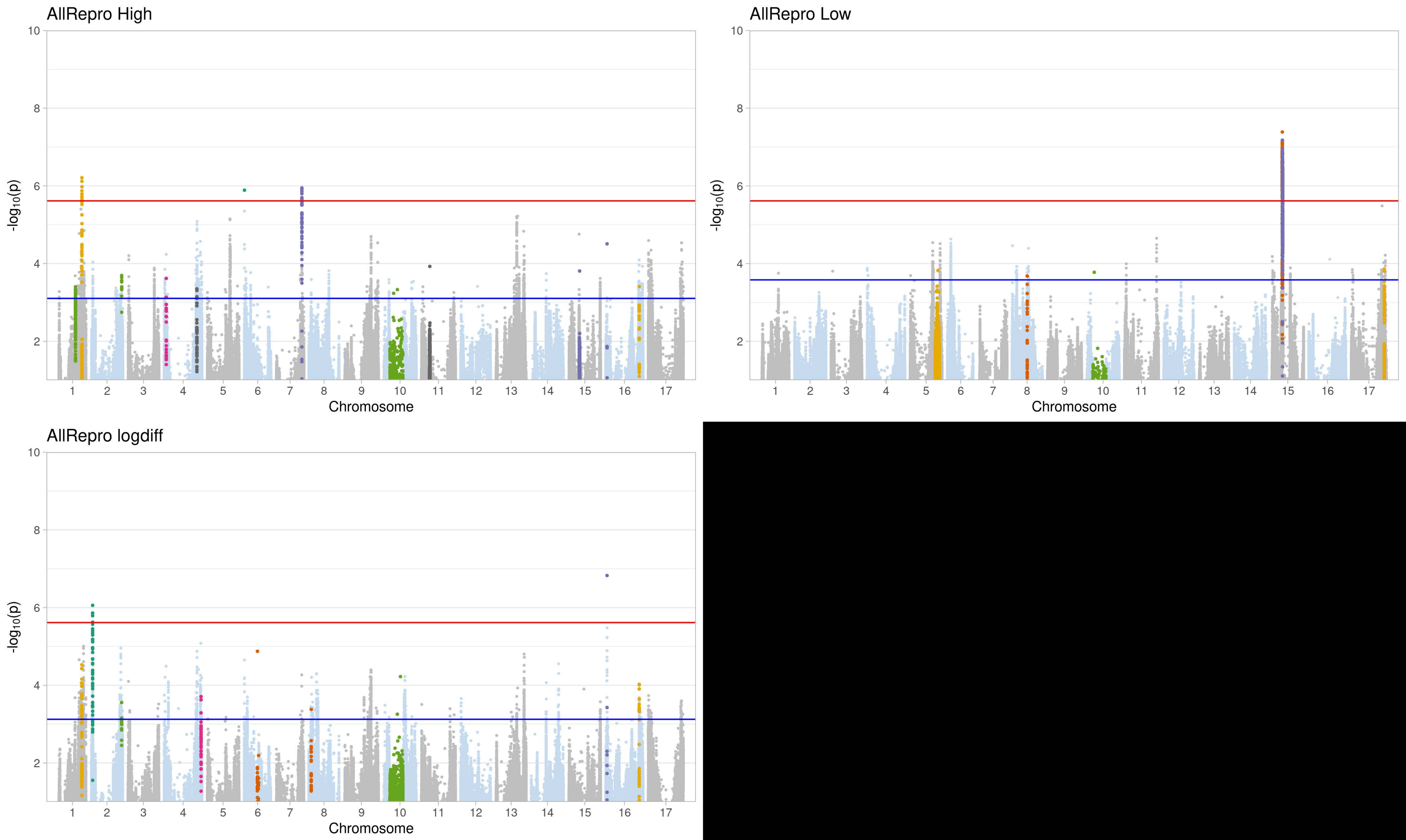
AllRepro High
AllRepro Low
AllRepro logdiff
-log10(p)
Chromosome
0 2 4 6 8 10
1 2 3 4 5 6 7 8 9 10 11 12 13 14 15 16 17

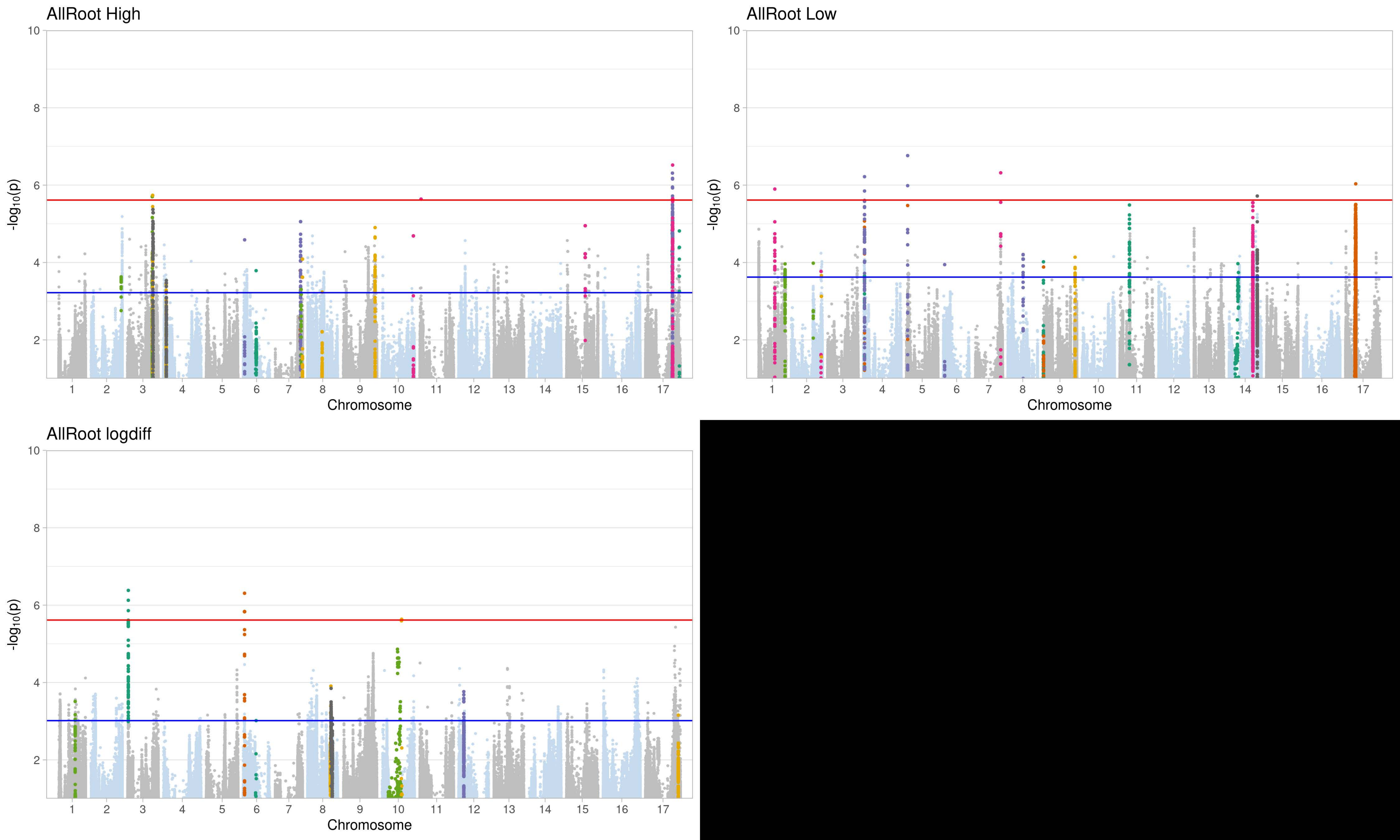
AllRoot High
AllRoot Low
AllRoot logdiff
-log10(p)
Chromosome
10
8
6
4
2
1 2 3 4 5 6 7 8 9 10 11 12 13 14 15 16 17

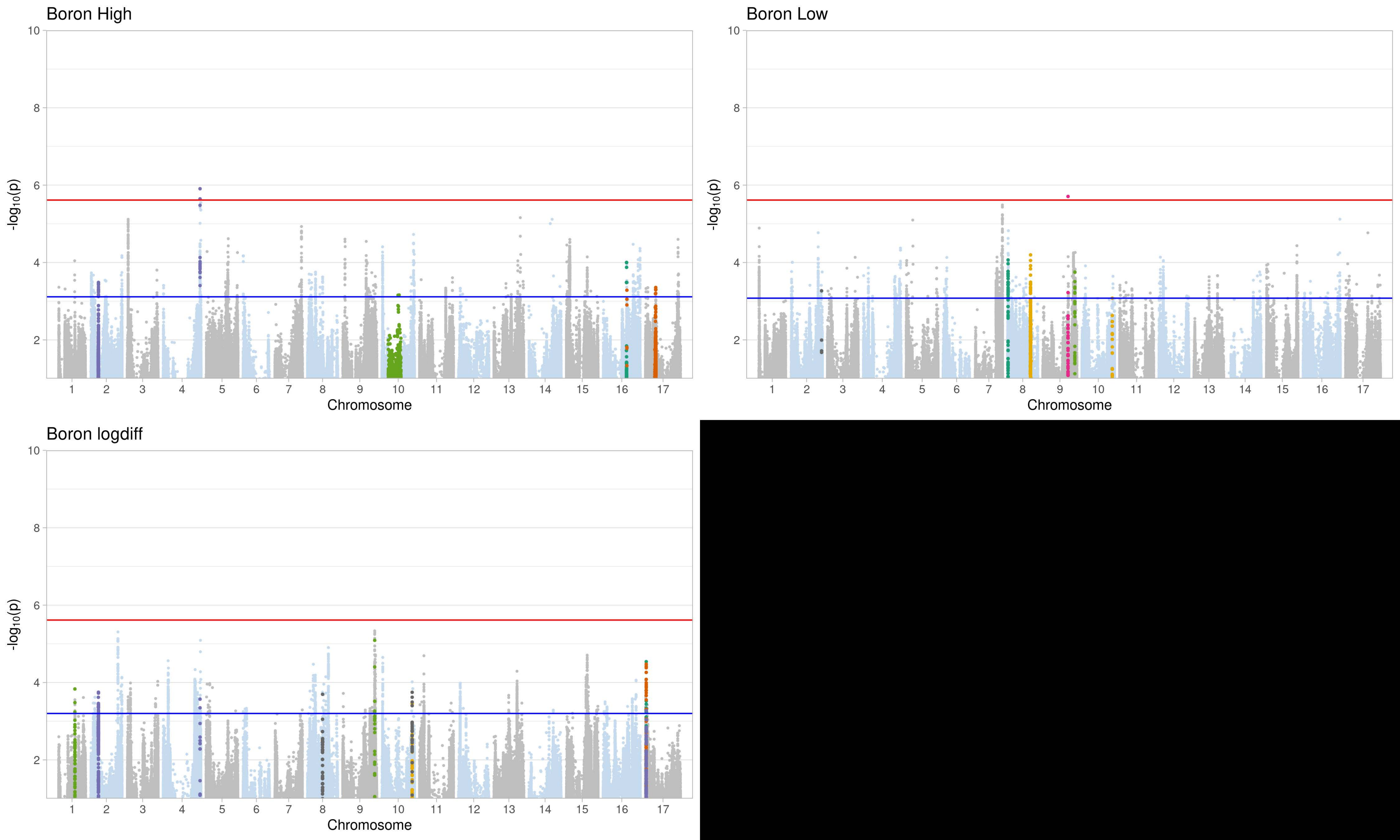

Boron High
-log10(p)
10
8
6
4
2
Chromosome
Boron Low
-log10(p)
10
8
6
4
2
Chromosome
Boron logdiff
-log10(p)
10
8
6
4
2
Chromosome

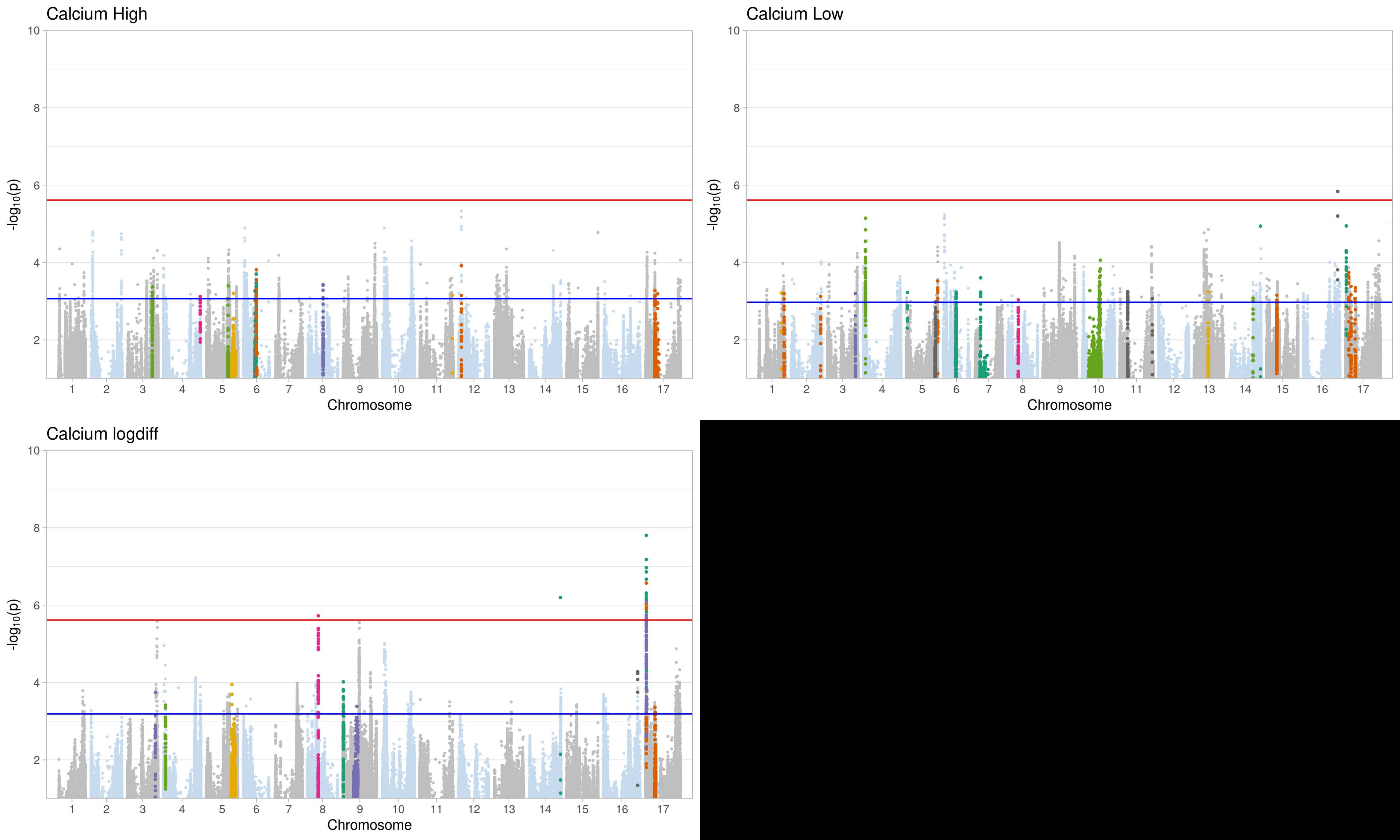
Calcium High
Calcium Low
Calcium logdiff
-log10(p)
Chromosome
1 2 3 4 5 6 7 8 9 10 11 12 13 14 15 16 17
2 4 6 8 10

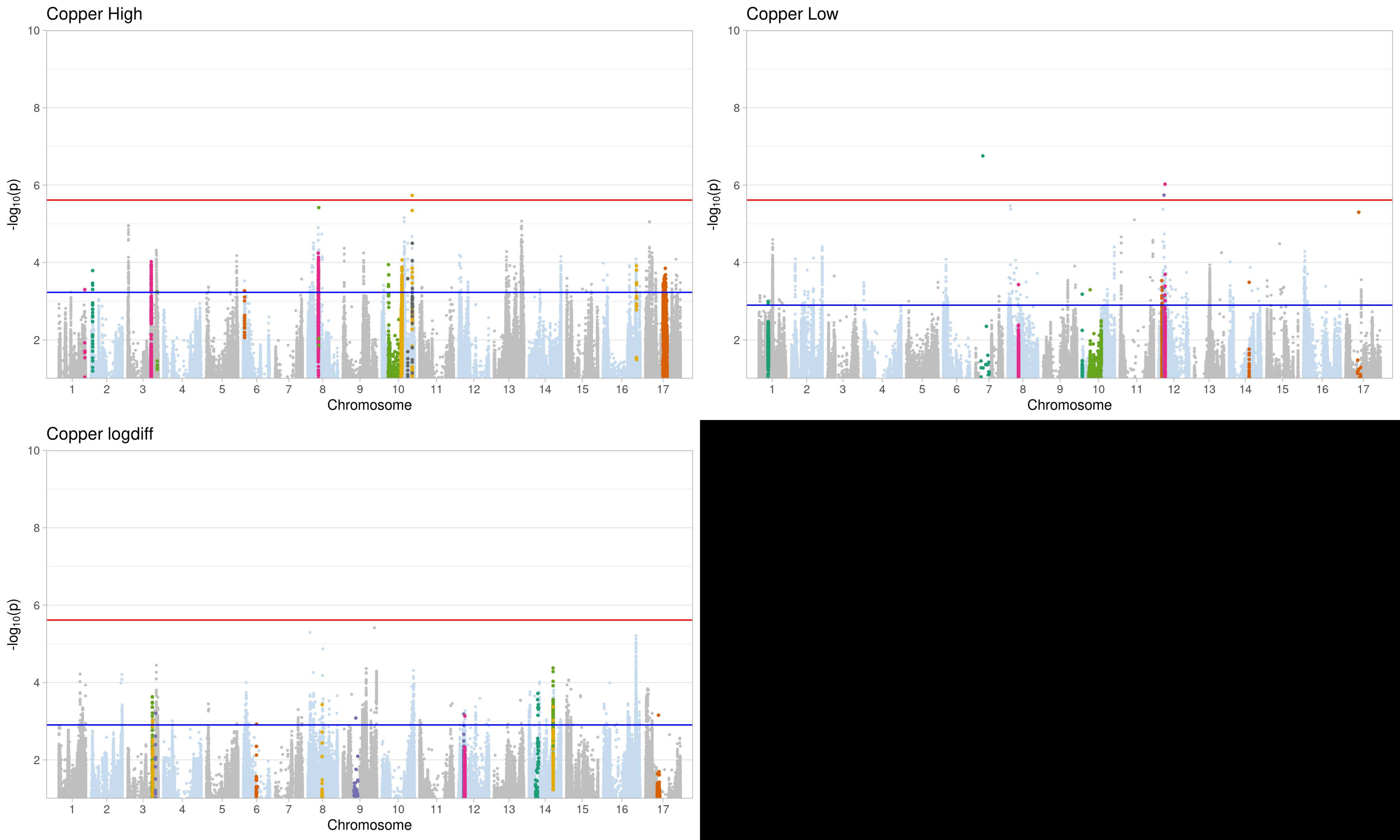
Copper High
Copper Low
Copper logdiff
-log10(p)
Chromosome
1 2 3 4 5 6 7 8 9 10 11 12 13 14 15 16 17
2 4 6 8 10

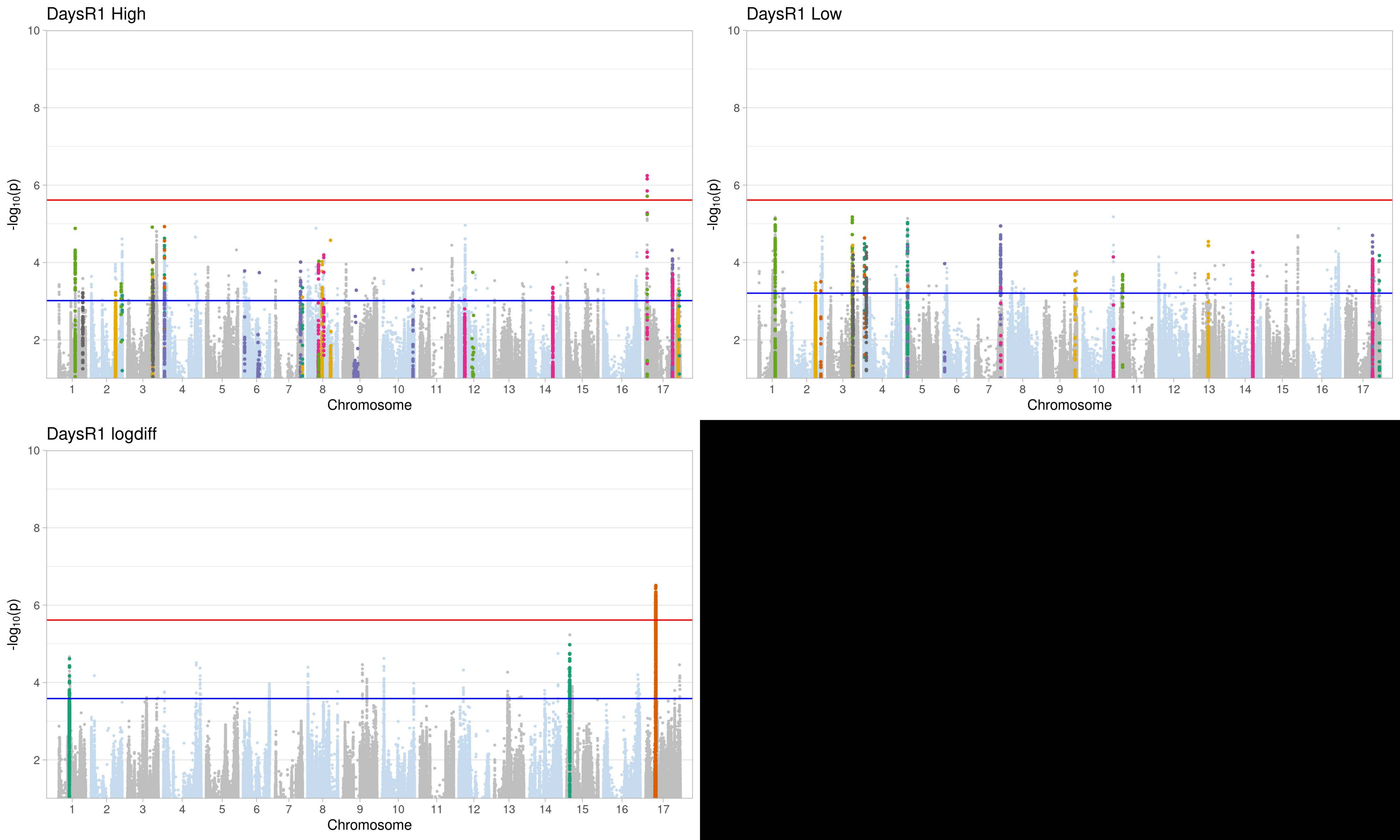
DaysR1 High
DaysR1 Low
DaysR1 logdiff
-log10(p)
Chromosome
2
4
6
8
10
1 2 3 4 5 6 7 8 9 10 11 12 13 14 15 16 17

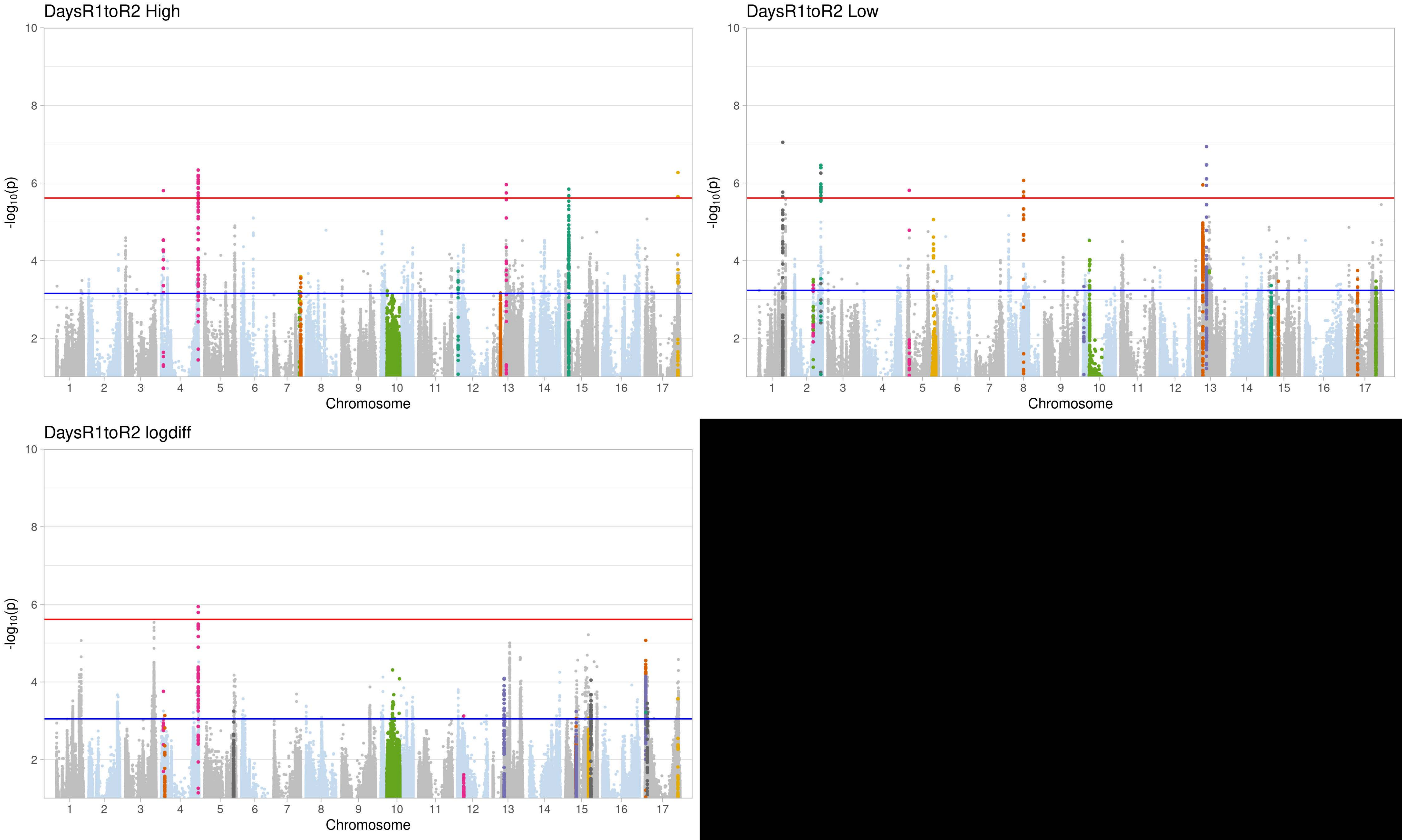
DaysR1toR2 High
DaysR1toR2 Low
DaysR1toR2 logdiff
-log10(p)
Chromosome
10
8
6
4
2
1 2 3 4 5 6 7 8 9 10 11 12 13 14 15 16 17

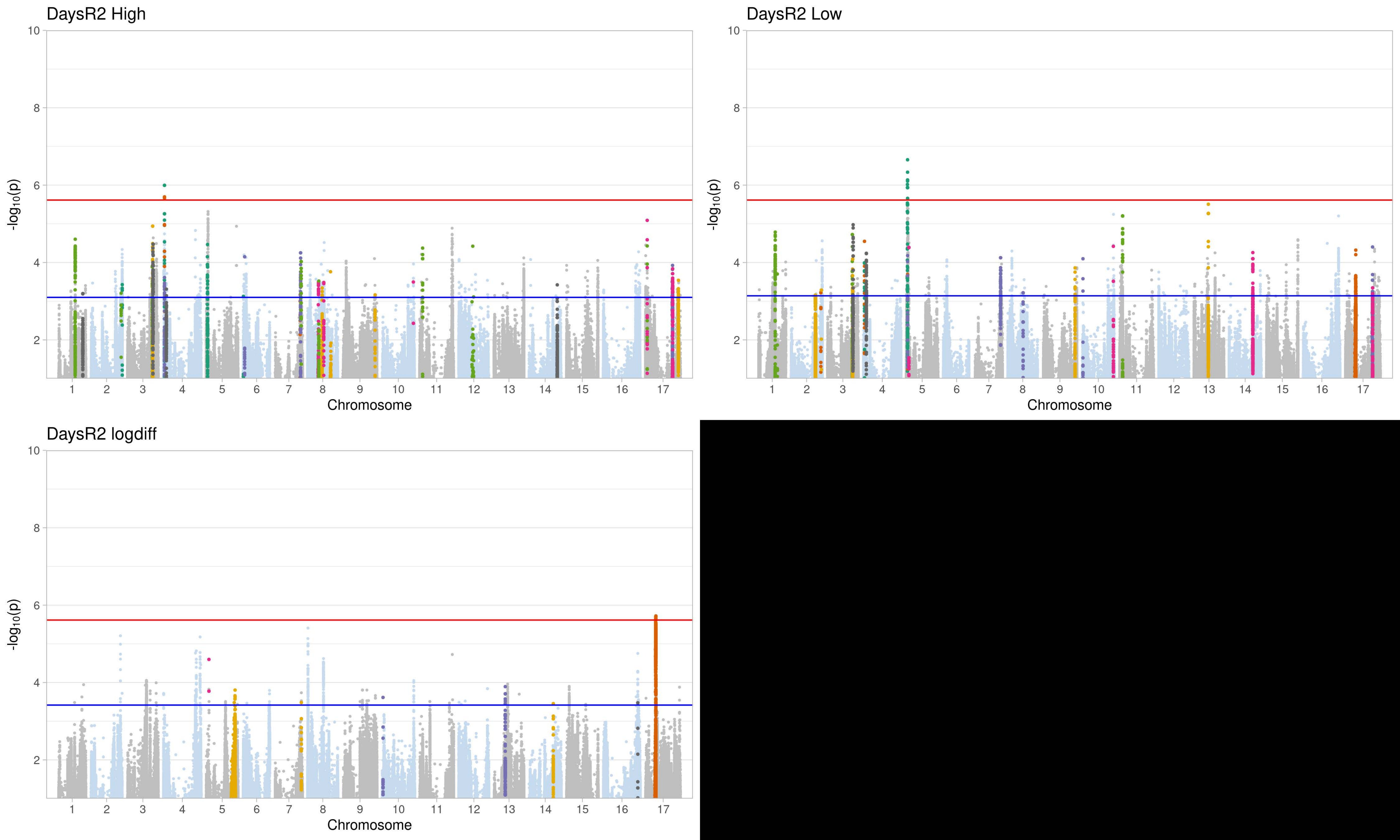
DaysR2 High
DaysR2 Low
DaysR2 logdiff
-log10(p)
Chromosome
10
8
6
4
2
1 2 3 4 5 6 7 8 9 10 11 12 13 14 15 16 17

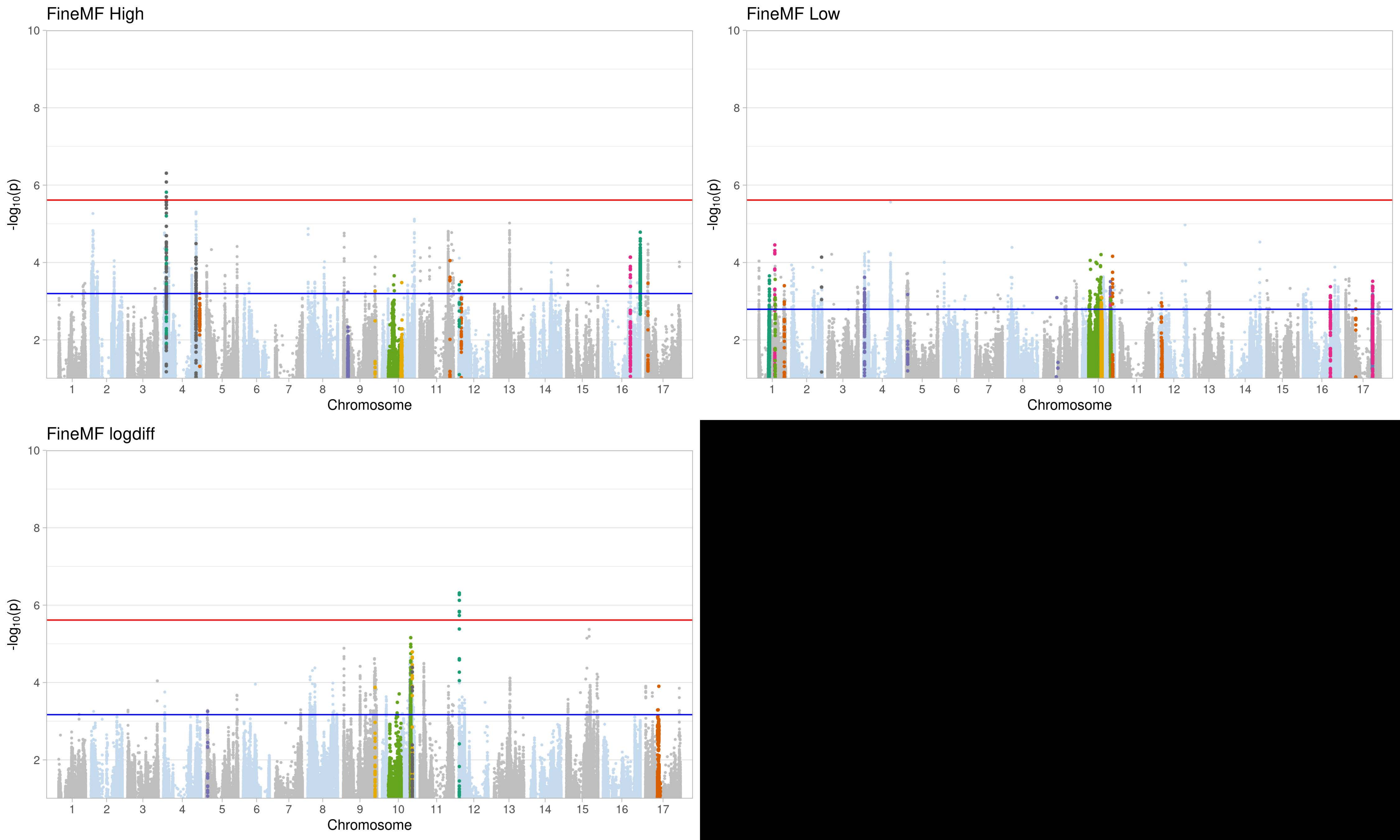
FineMF High
FineMF Low
FineMF logdiff
-log10(p)
Chromosome
10
8
6
4
2
1 2 3 4 5 6 7 8 9 10 11 12 13 14 15 16 17

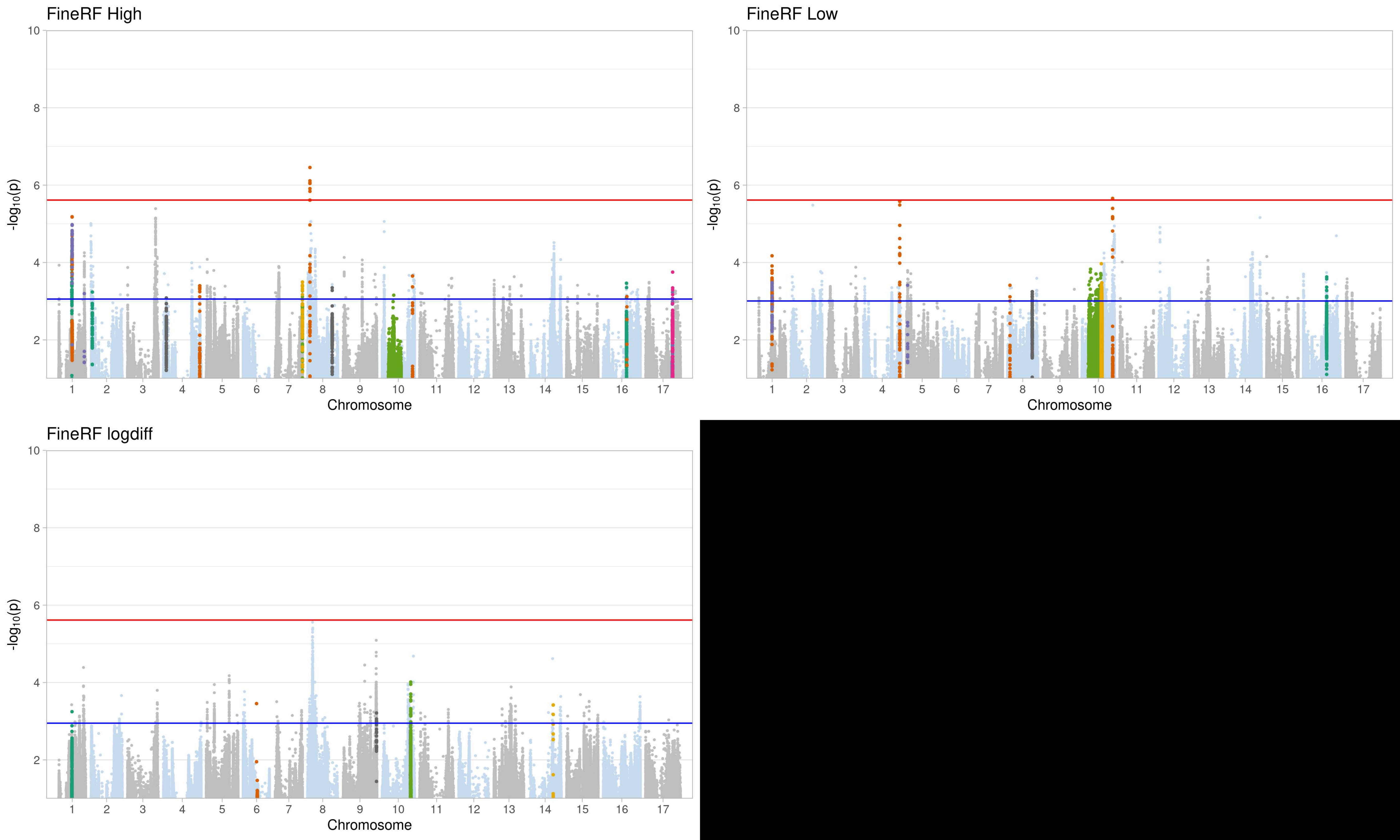
FineRF High
FineRF Low
FineRF logdiff
-log10(p)
Chromosome
10
8
6
4
2
1 2 3 4 5 6 7 8 9 10 11 12 13 14 15 16 17

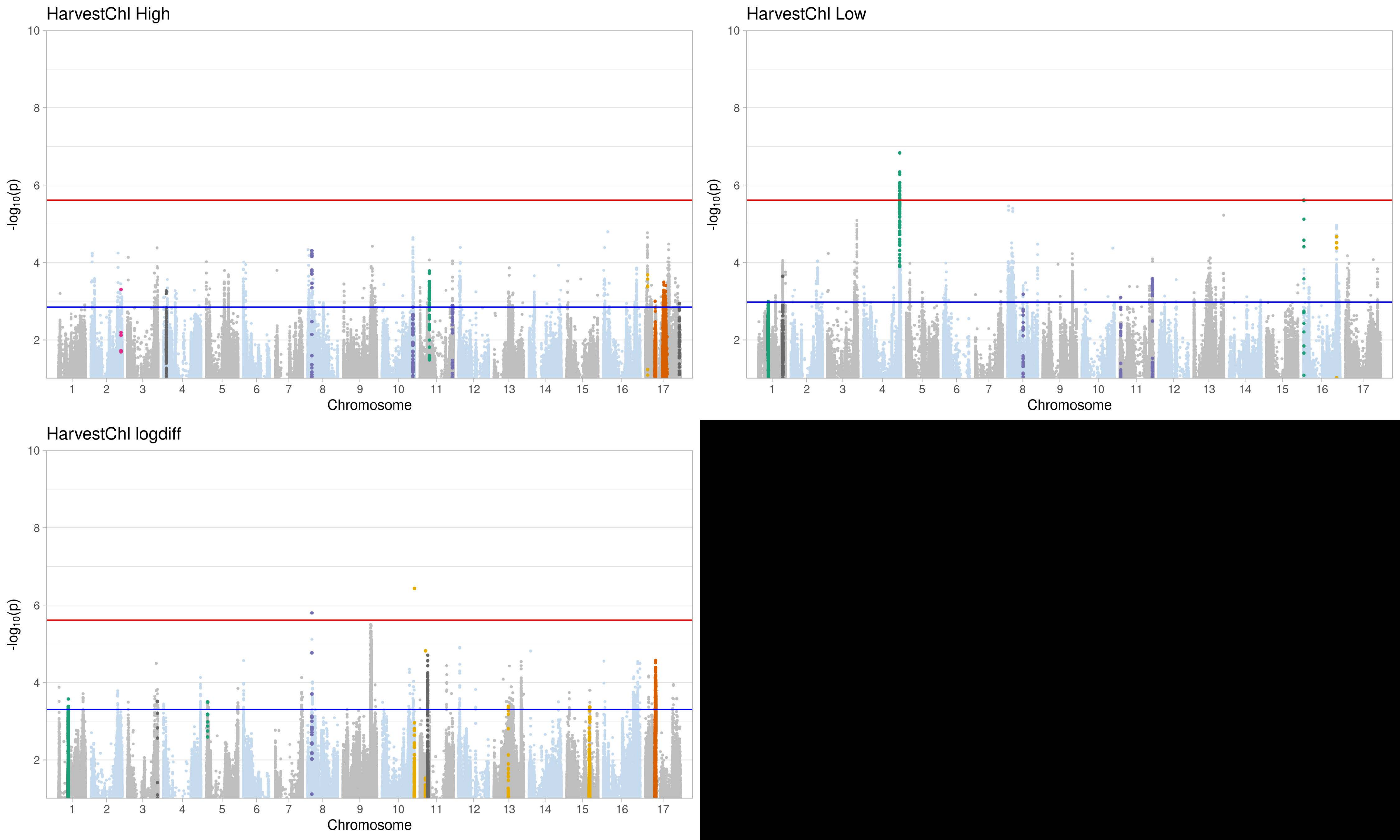
HarvestChl High
HarvestChl Low
HarvestChl logdiff
-log10(p)
Chromosome
10
8
6
4
2
1 2 3 4 5 6 7 8 9 10 11 12 13 14 15 16 17

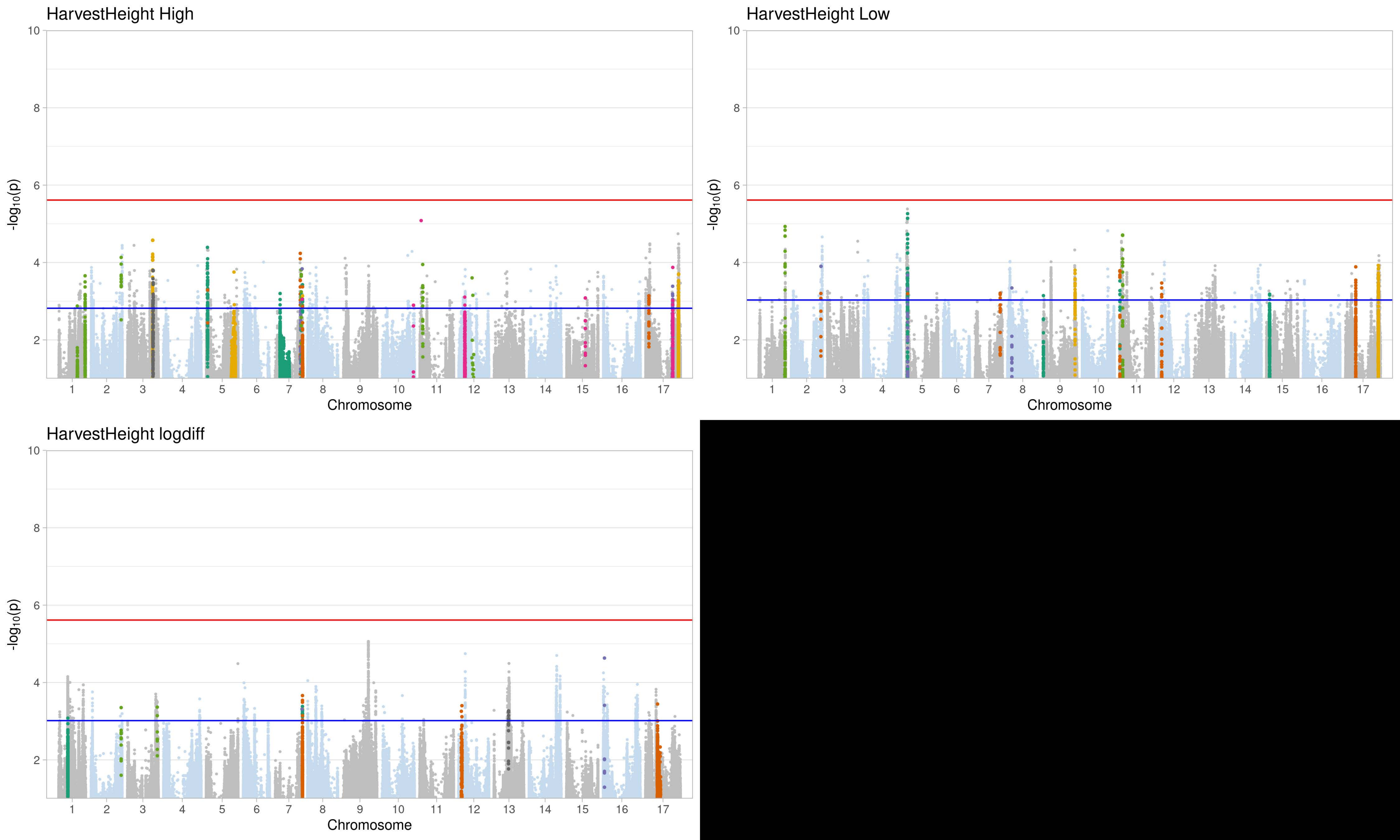
HarvestHeight High
HarvestHeight Low
HarvestHeight logdiff
-log10(p)
Chromosome
1 2 3 4 5 6 7 8 9 10 11 12 13 14 15 16 17
2 4 6 8 10

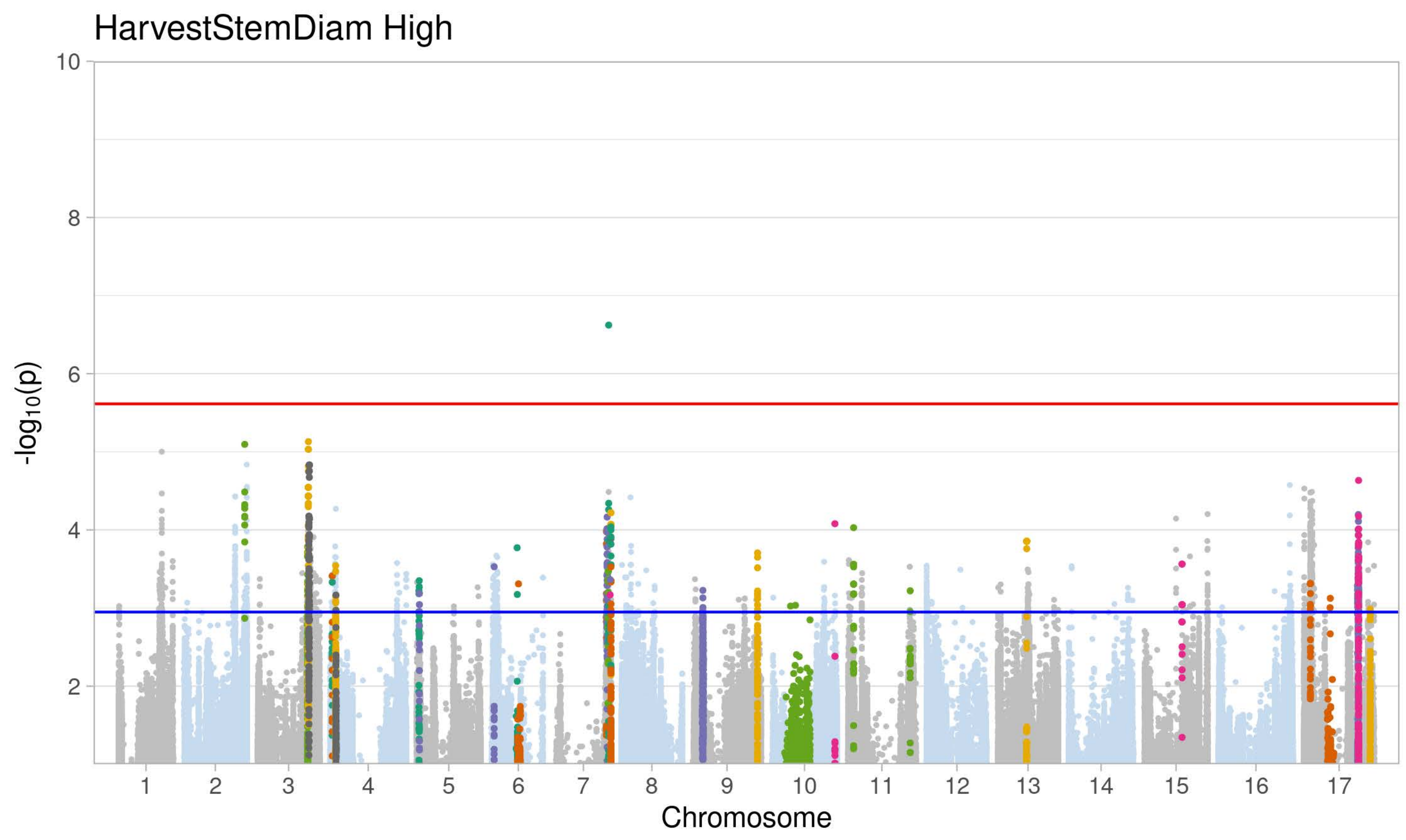
HarvestStemDiam High
-log10(p)
Chromosome

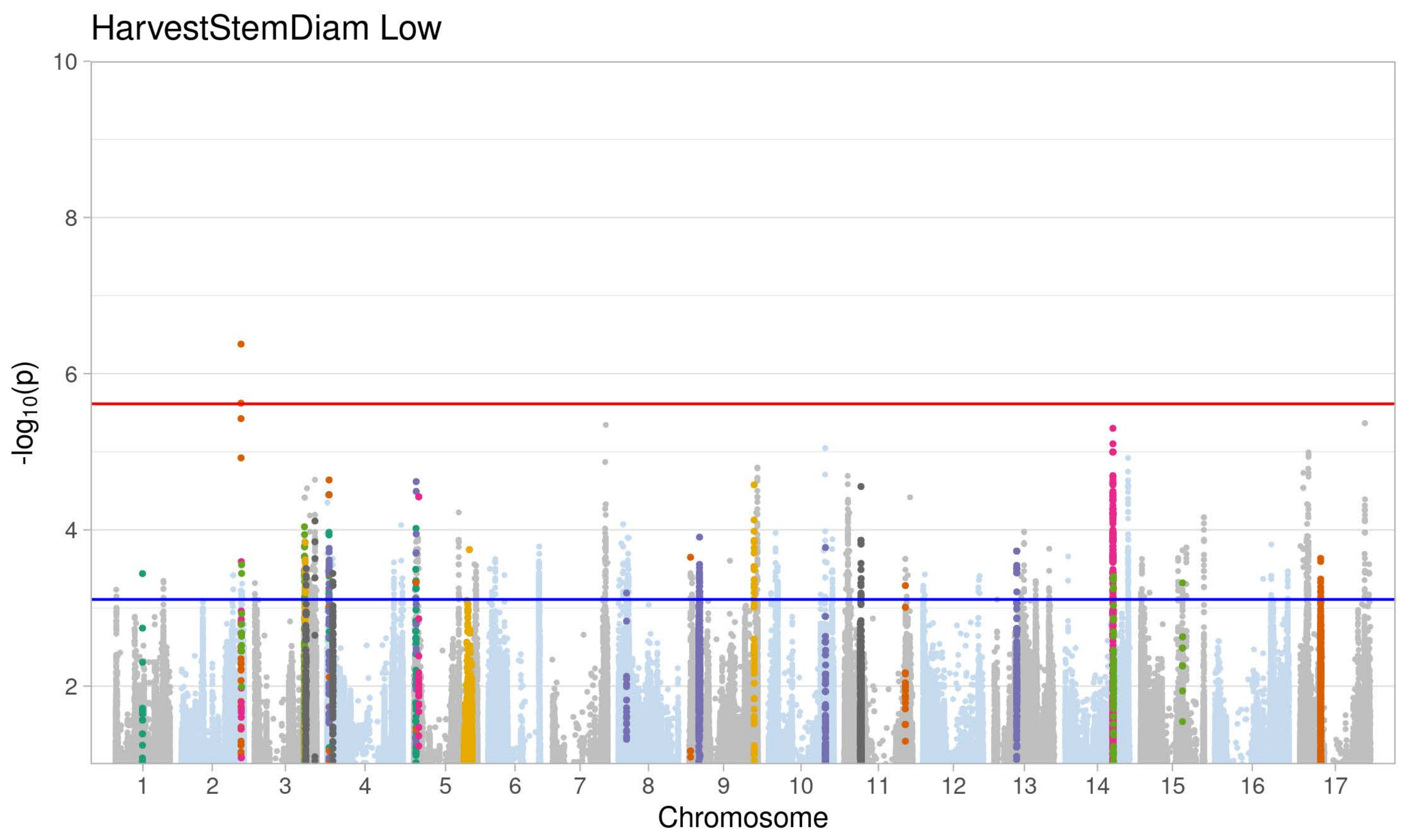
HarvestStemDiam Low
-log10(p)
Chromosome

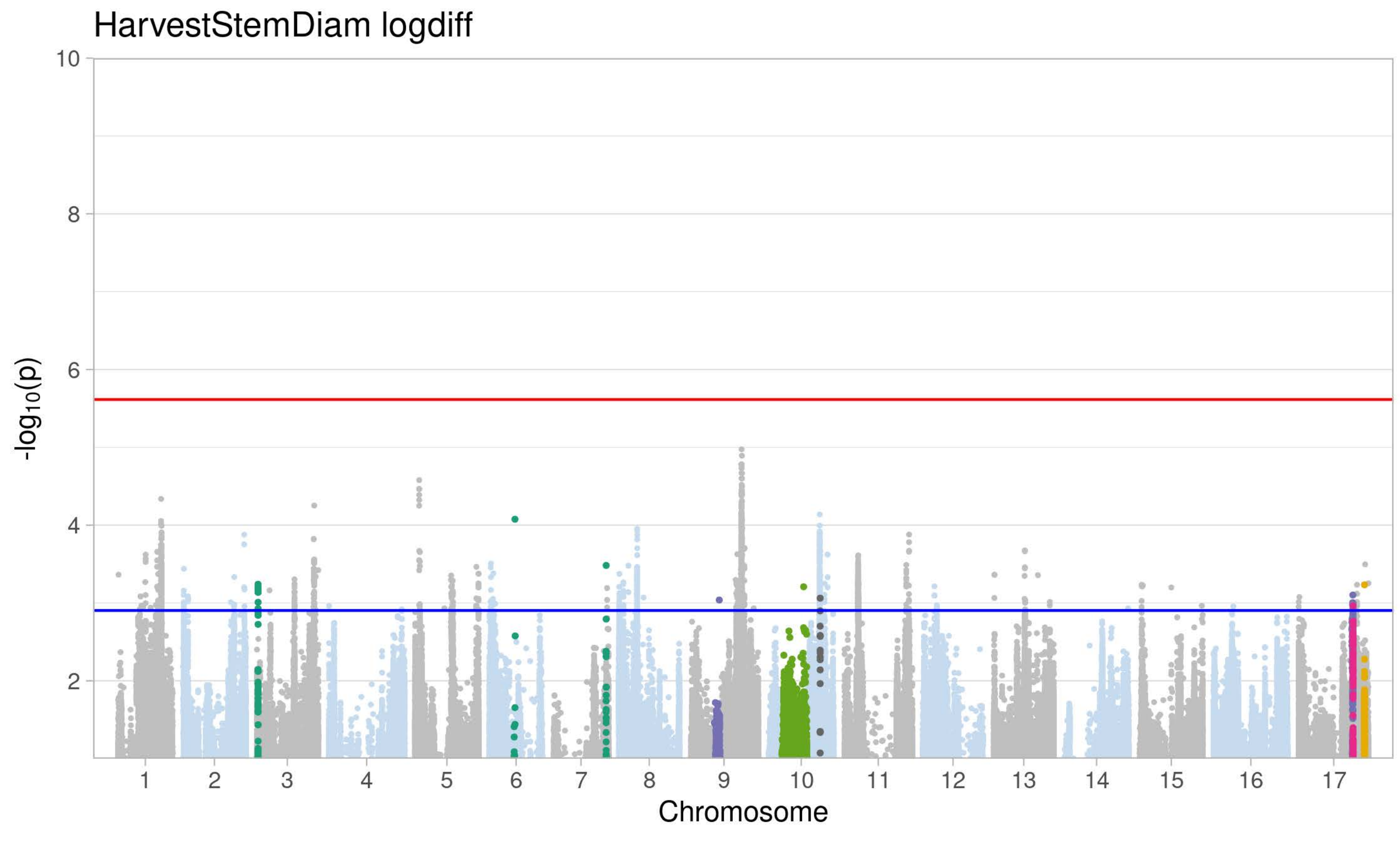
HarvestStemDiam logdiff
-log10(p)
Chromosome

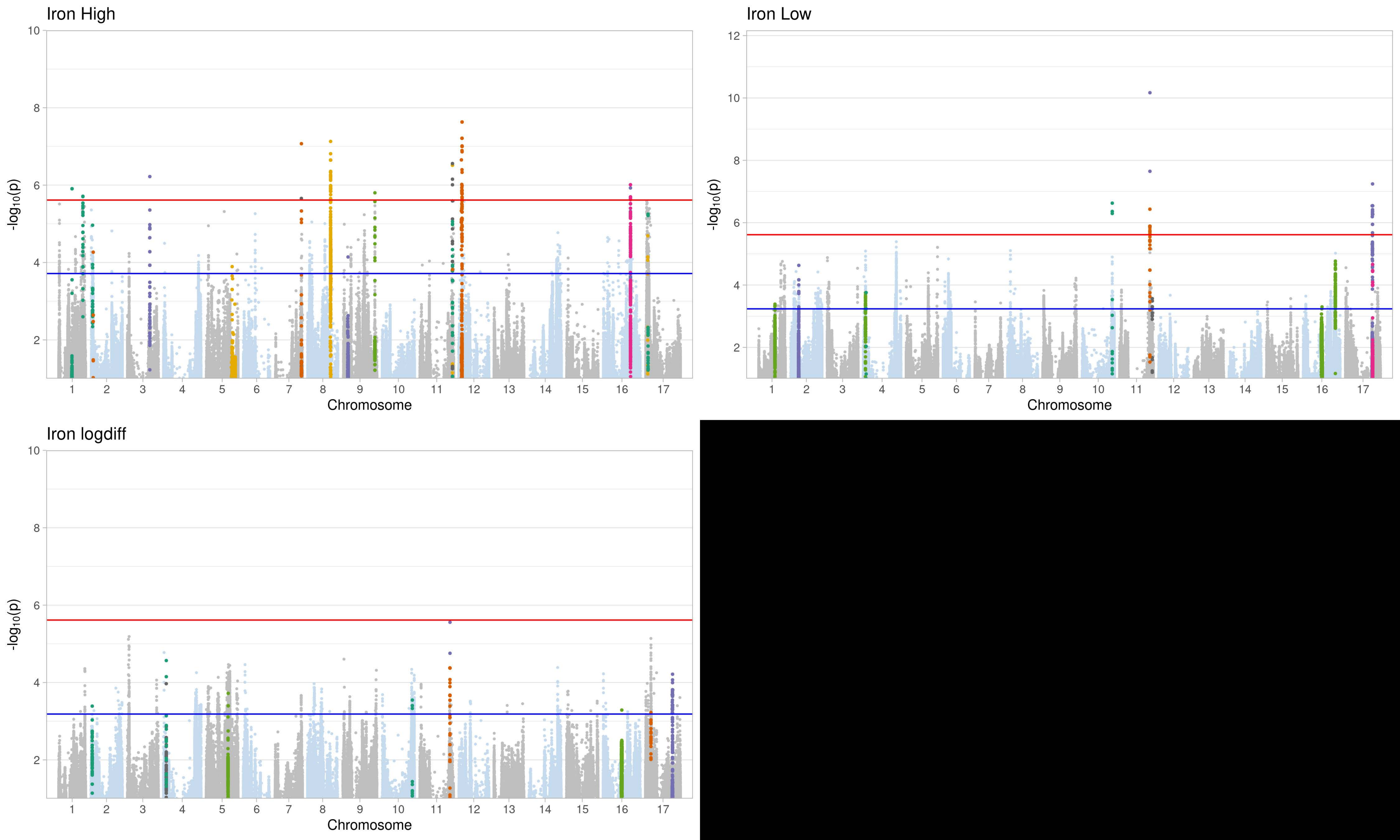
Iron High
Iron Low
Iron logdiff
-log10(p)
Chromosome
1 2 3 4 5 6 7 8 9 10 11 12 13 14 15 16 17
2 4 6 8 10 12

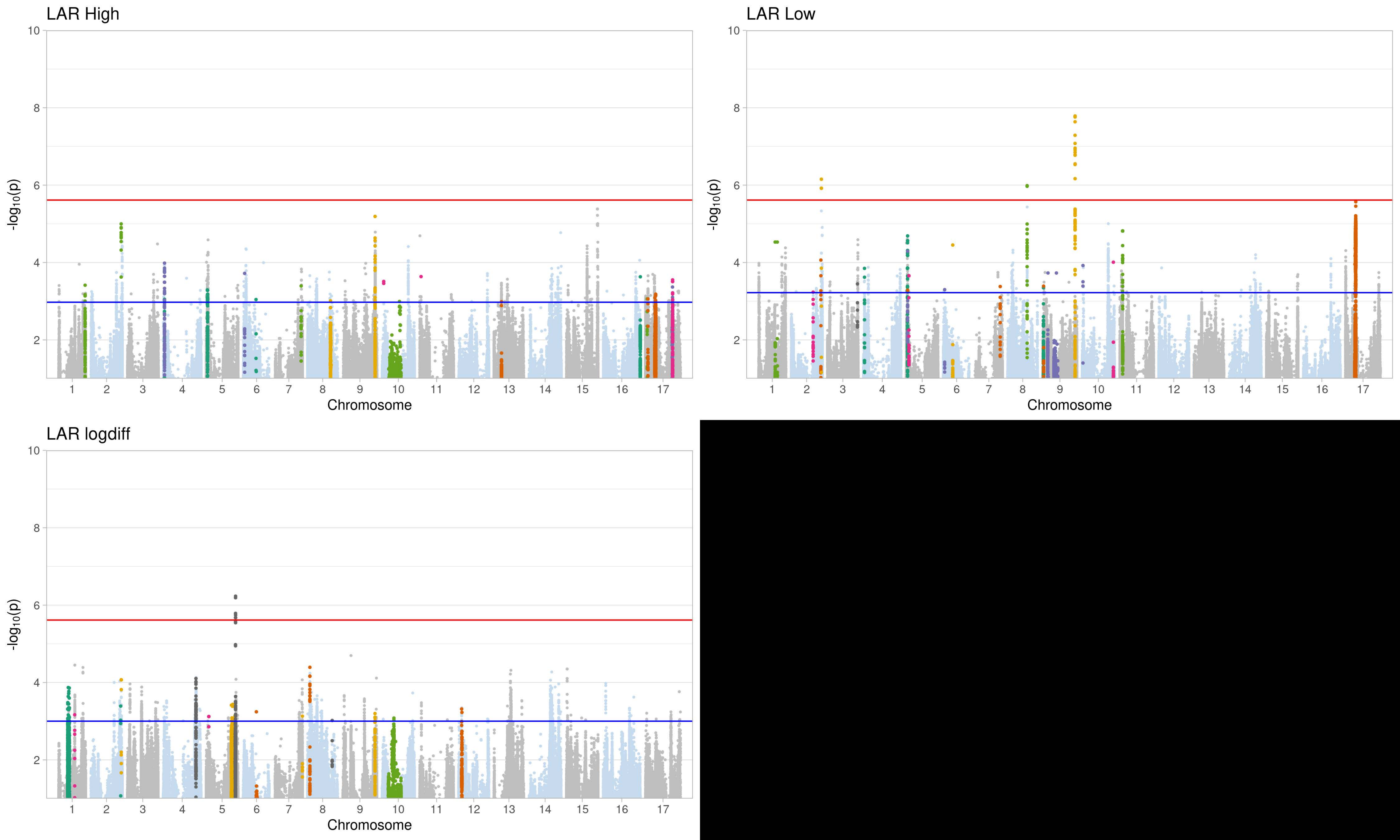

LAR High
LAR Low
LAR logdiff
-log10(p)
Chromosome
1 2 3 4 5 6 7 8 9 10 11 12 13 14 15 16 17
2 4 6 8 10

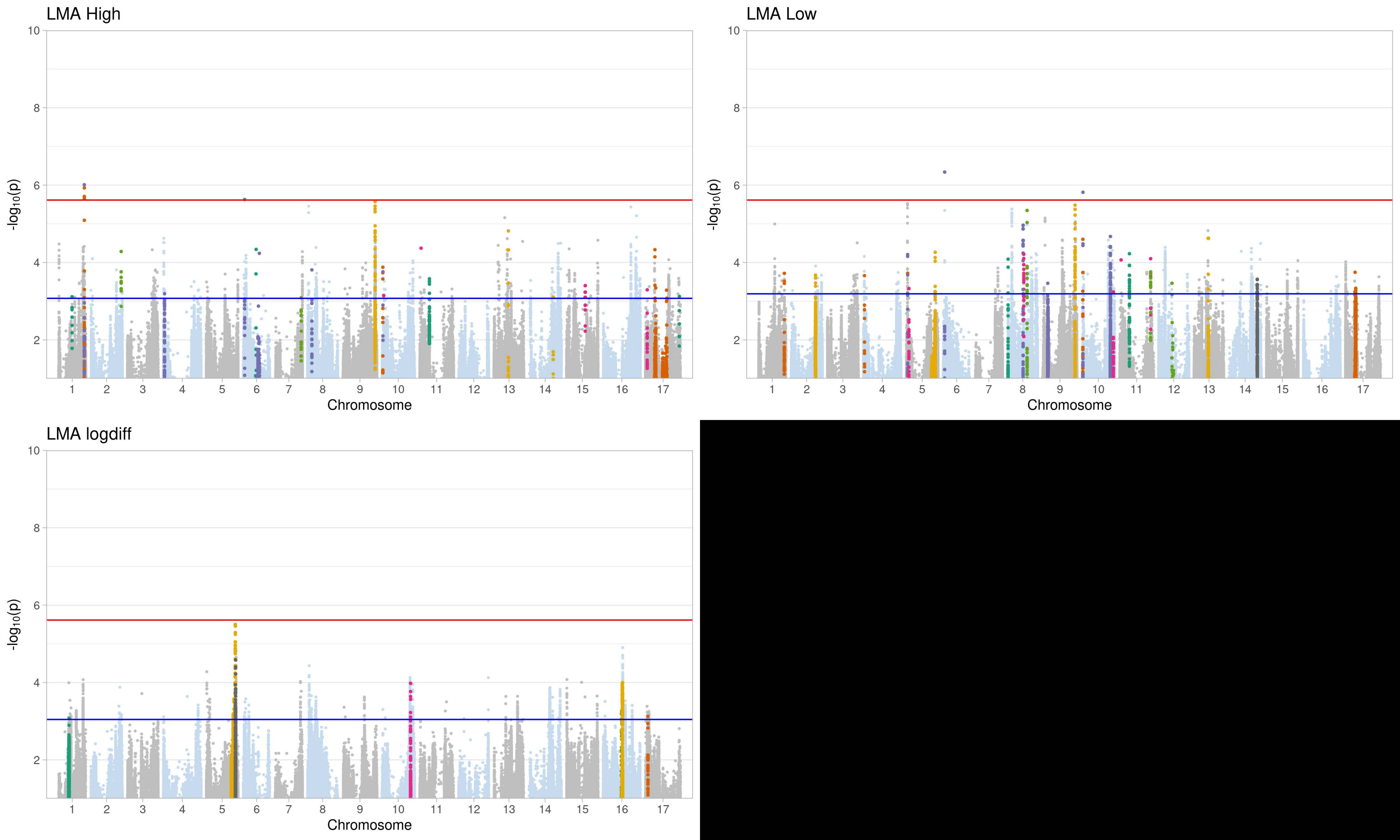
LMA High
LMA Low
LMA logdiff
-log10(p)
Chromosome
1 2 3 4 5 6 7 8 9 10 11 12 13 14 15 16 17
2 4 6 8 10

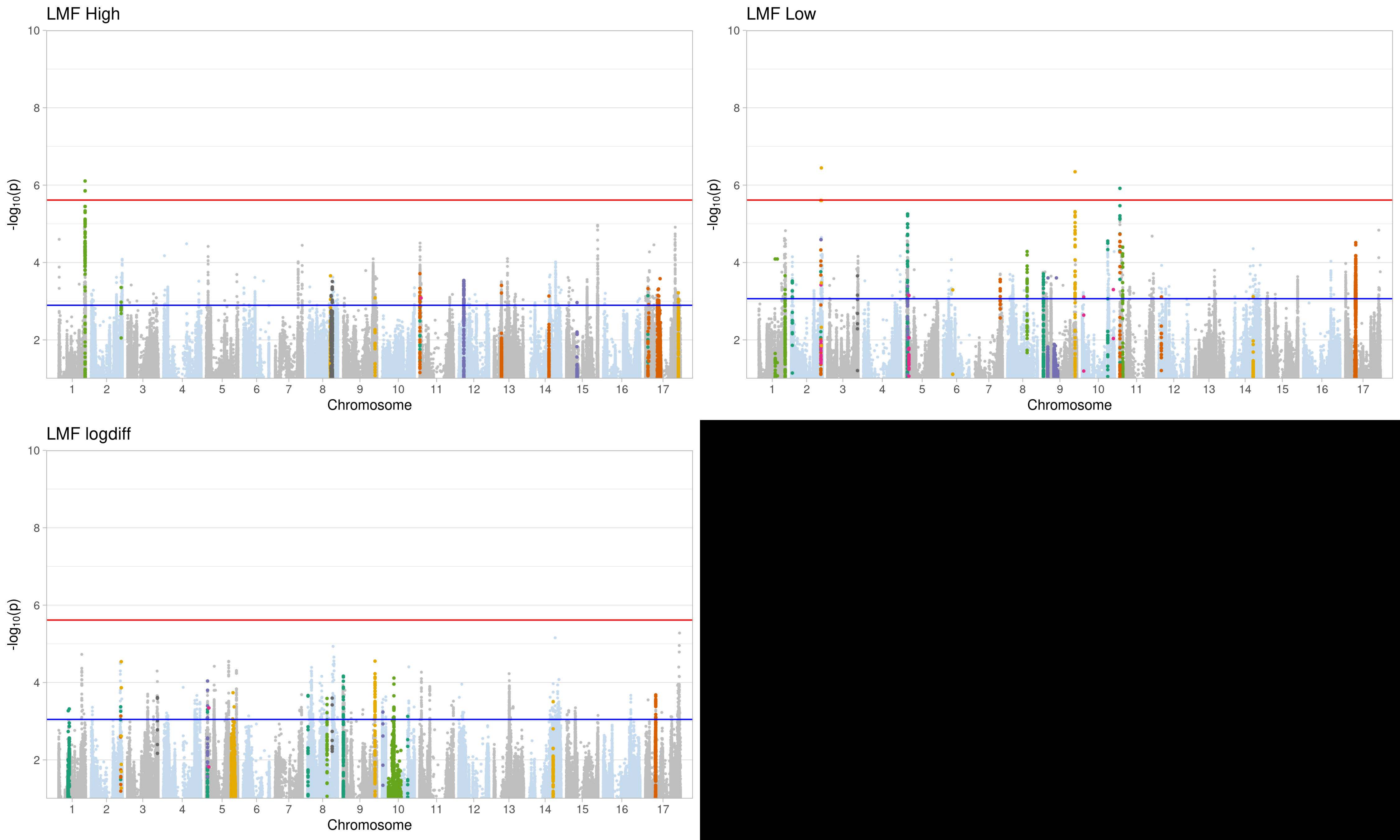
LMF High
LMF Low
LMF logdiff
-log10(p)
Chromosome
10
8
6
4
2
1 2 3 4 5 6 7 8 9 10 11 12 13 14 15 16 17

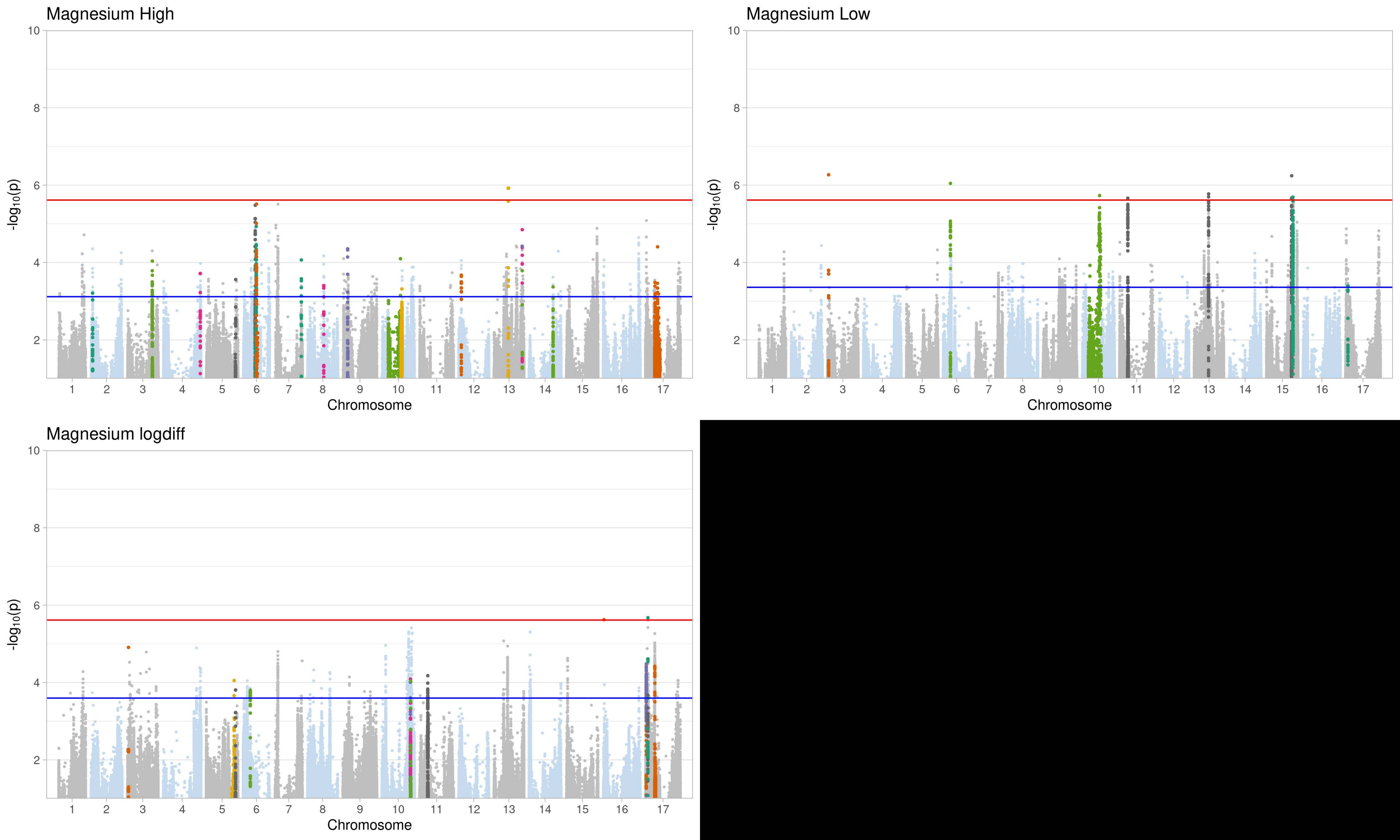
Magnesium High
Magnesium Low
Magnesium logdiff
-log10(p)
Chromosome
10
8
6
4
2
1 2 3 4 5 6 7 8 9 10 11 12 13 14 15 16 17

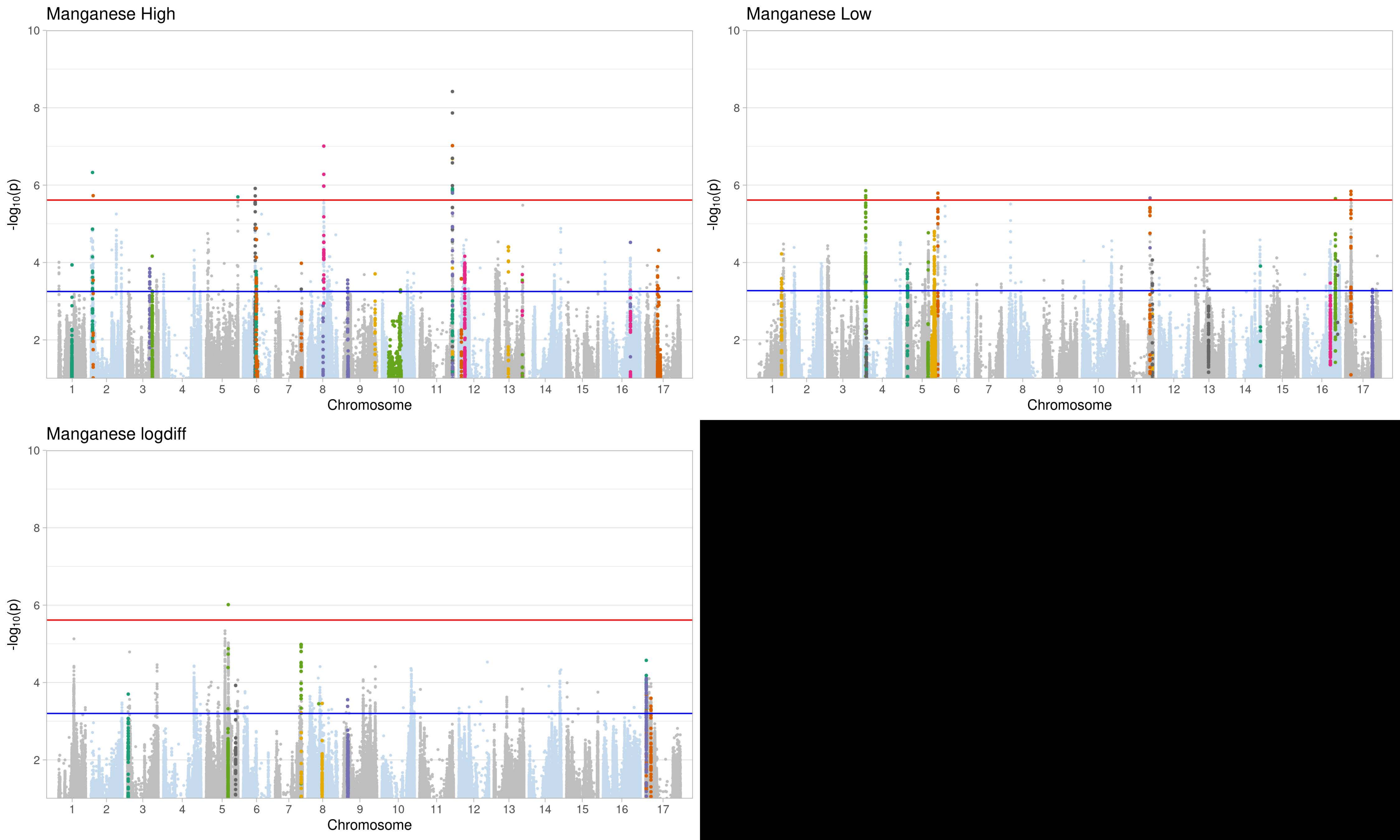
Manganese High
-log10(p)
Chromosome
1 2 3 4 5 6 7 8 9 10 11 12 13 14 15 16 17
Manganese Low
-log10(p)
Chromosome
1 2 3 4 5 6 7 8 9 10 11 12 13 14 15 16 17
Manganese logdiff
-log10(p)
Chromosome
1 2 3 4 5 6 7 8 9 10 11 12 13 14 15 16 17
10
8
6
4
2

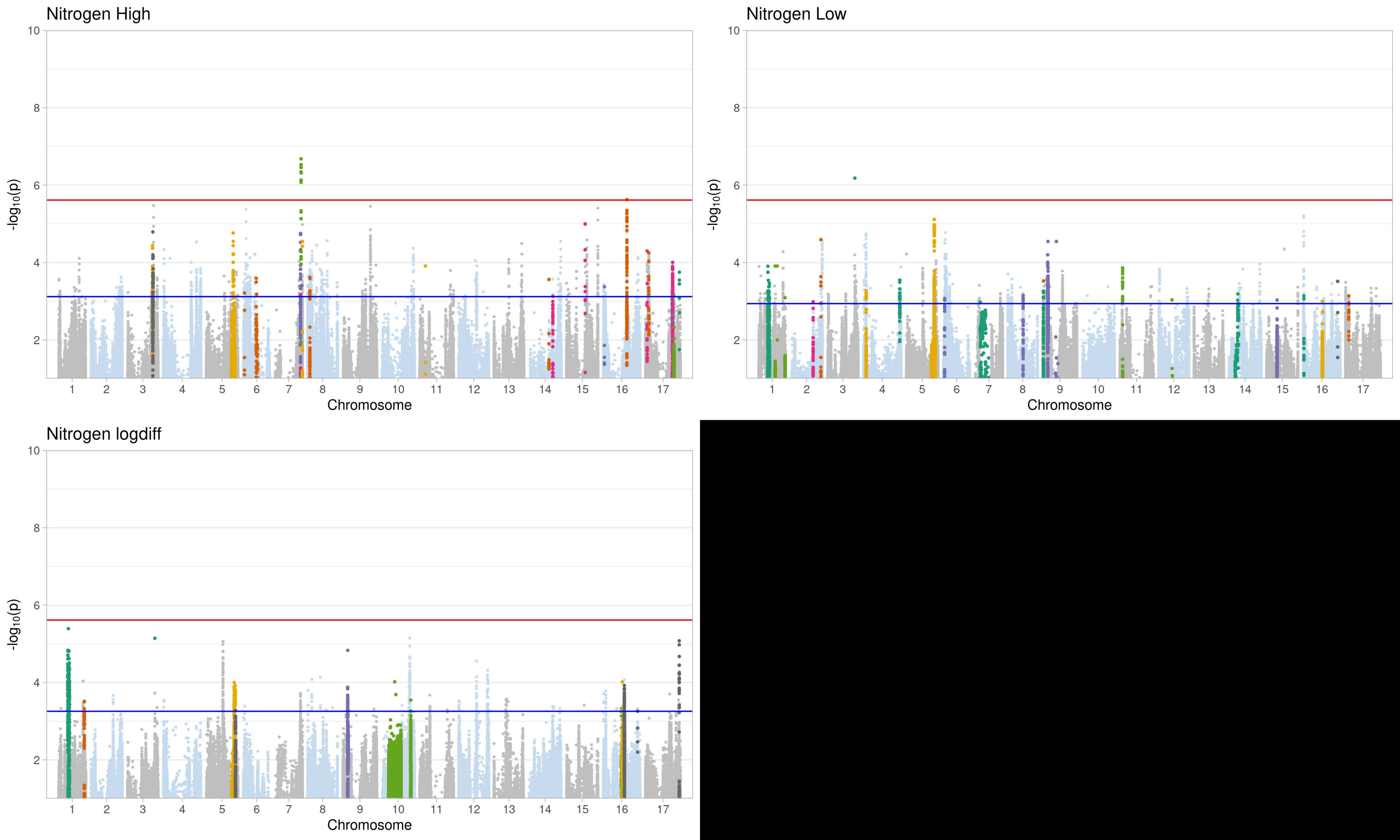

Nitrogen High
Nitrogen Low
Nitrogen logdiff
$-\log_{10}(p)$
Chromosome
0 2 4 6 8 10
1 2 3 4 5 6 7 8 9 10 11 12 13 14 15 16 17

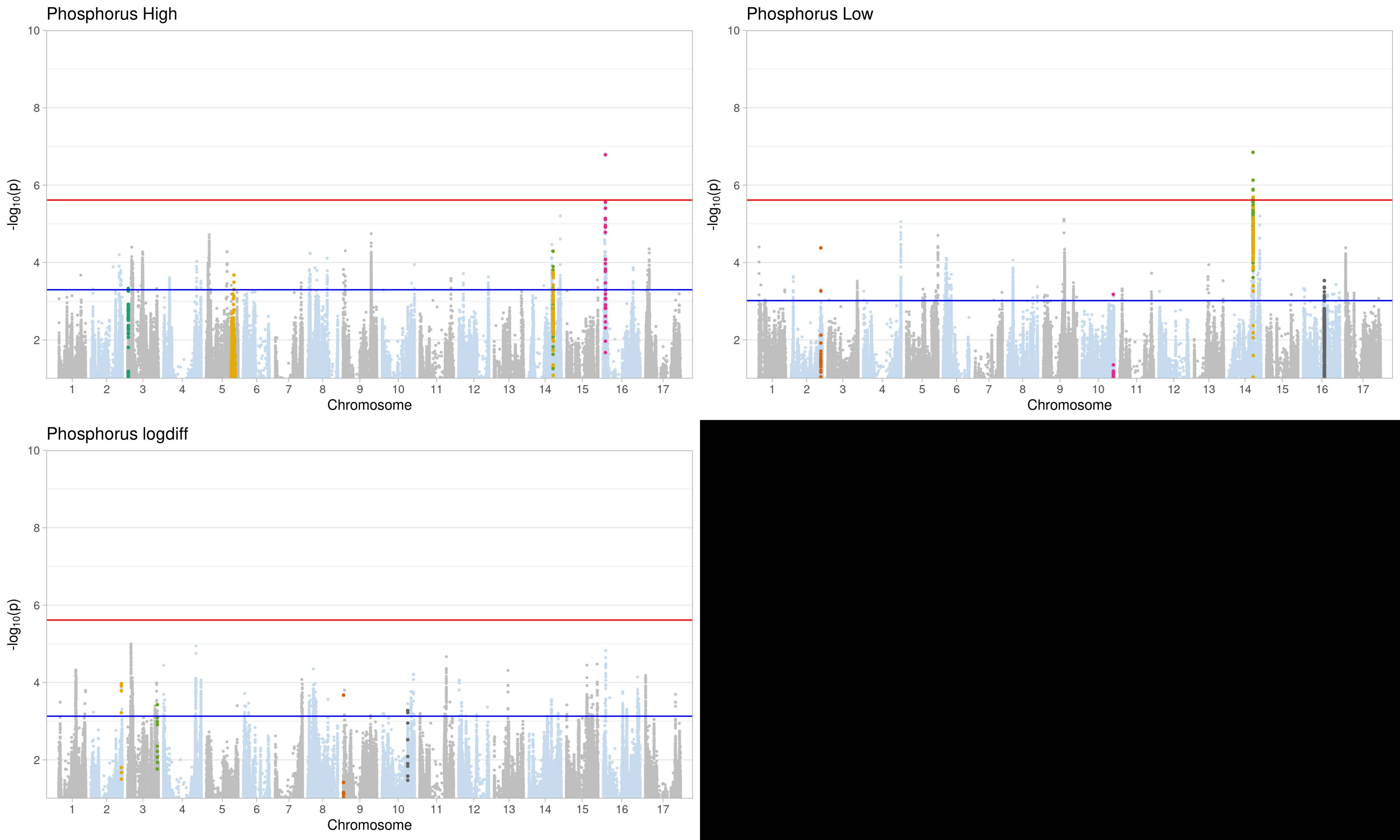

Phosphorus High
Phosphorus Low
Phosphorus logdiff
-log10(p)
Chromosome
0
2
4
6
8
10

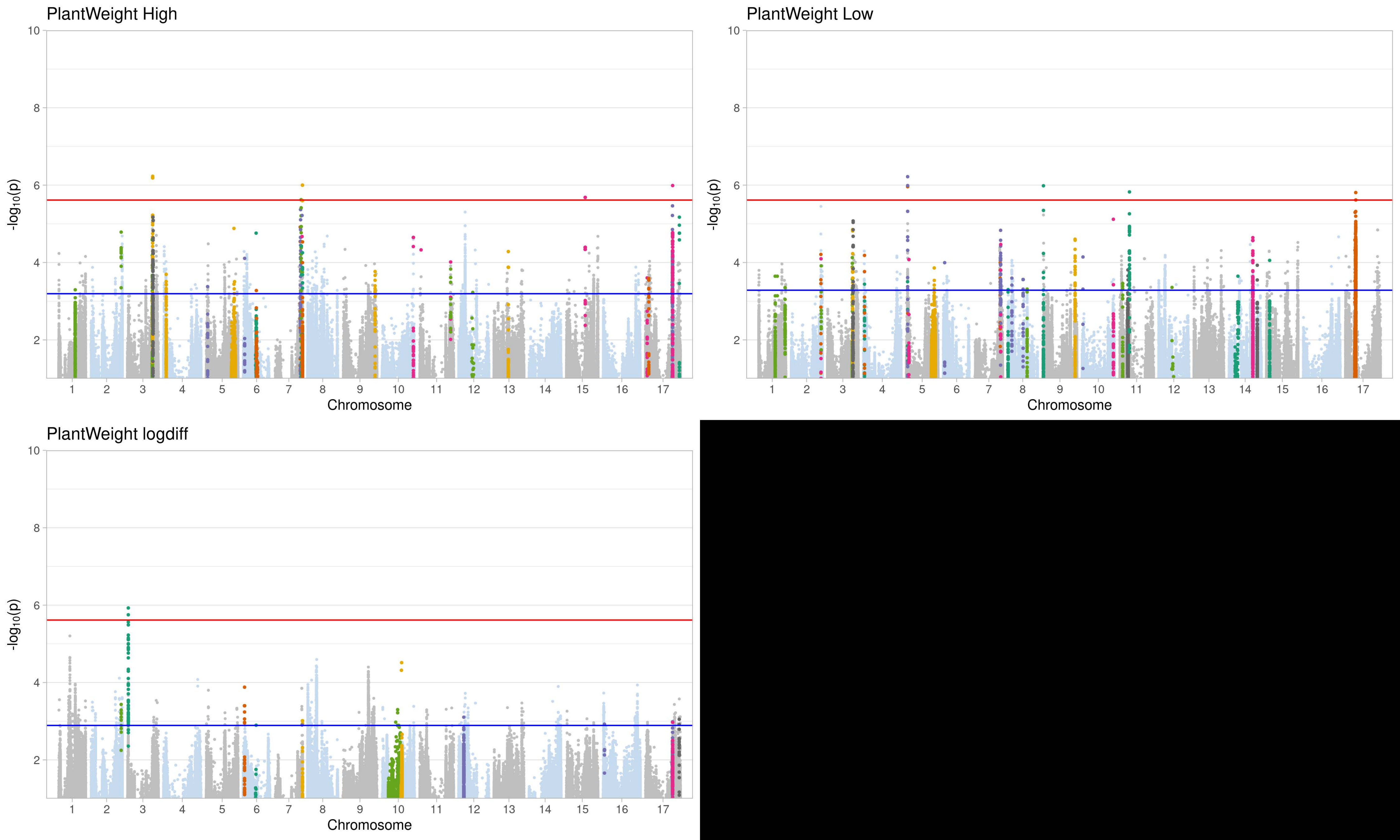
PlantWeight High
PlantWeight Low
PlantWeight logdiff
-log10(p)
Chromosome
10
8
6
4
2
1 2 3 4 5 6 7 8 9 10 11 12 13 14 15 16 17

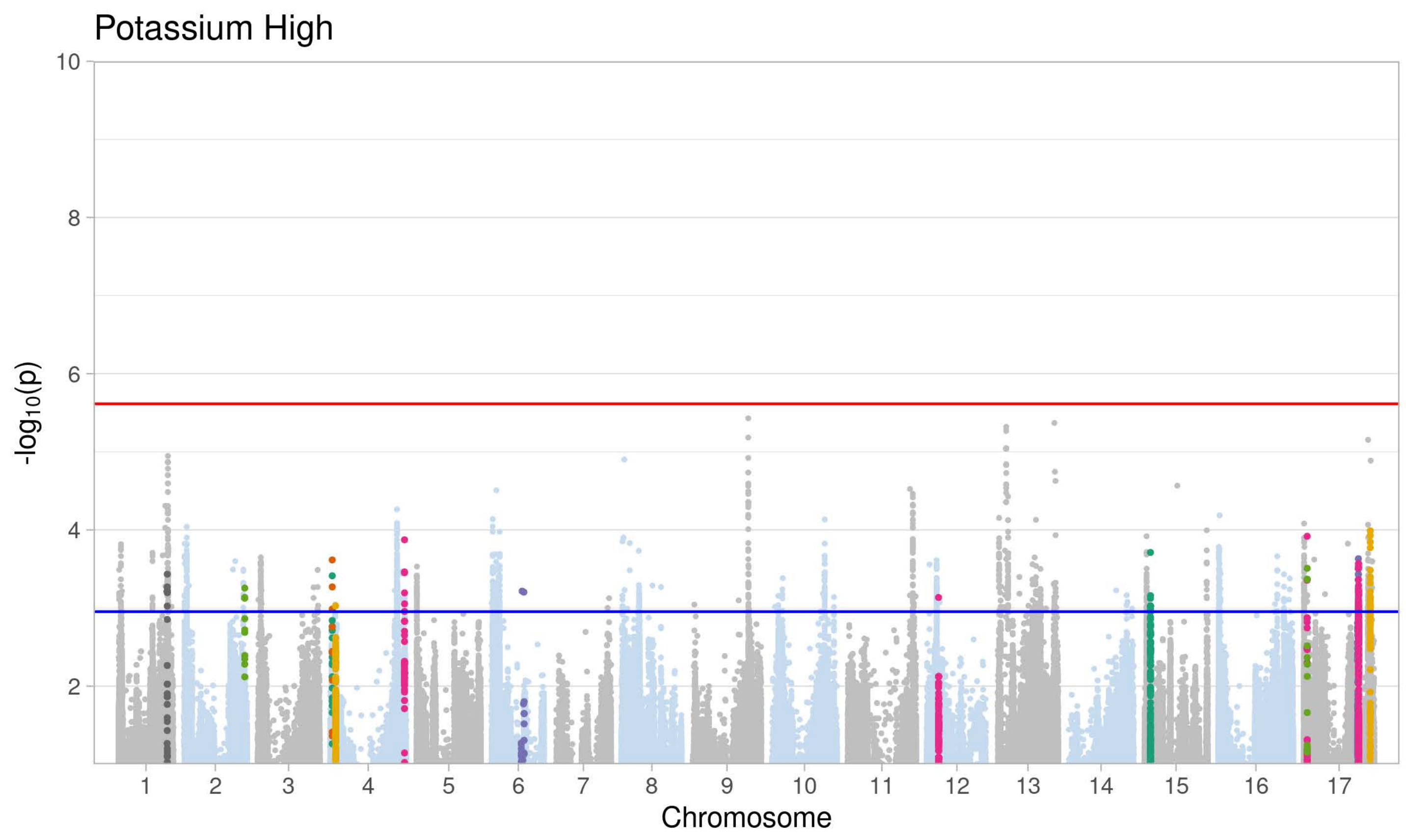
Potassium High
-log10(p)
10
8
6
4
2
1 2 3 4 5 6 7 8 9 10 11 12 13 14 15 16 17
Chromosome

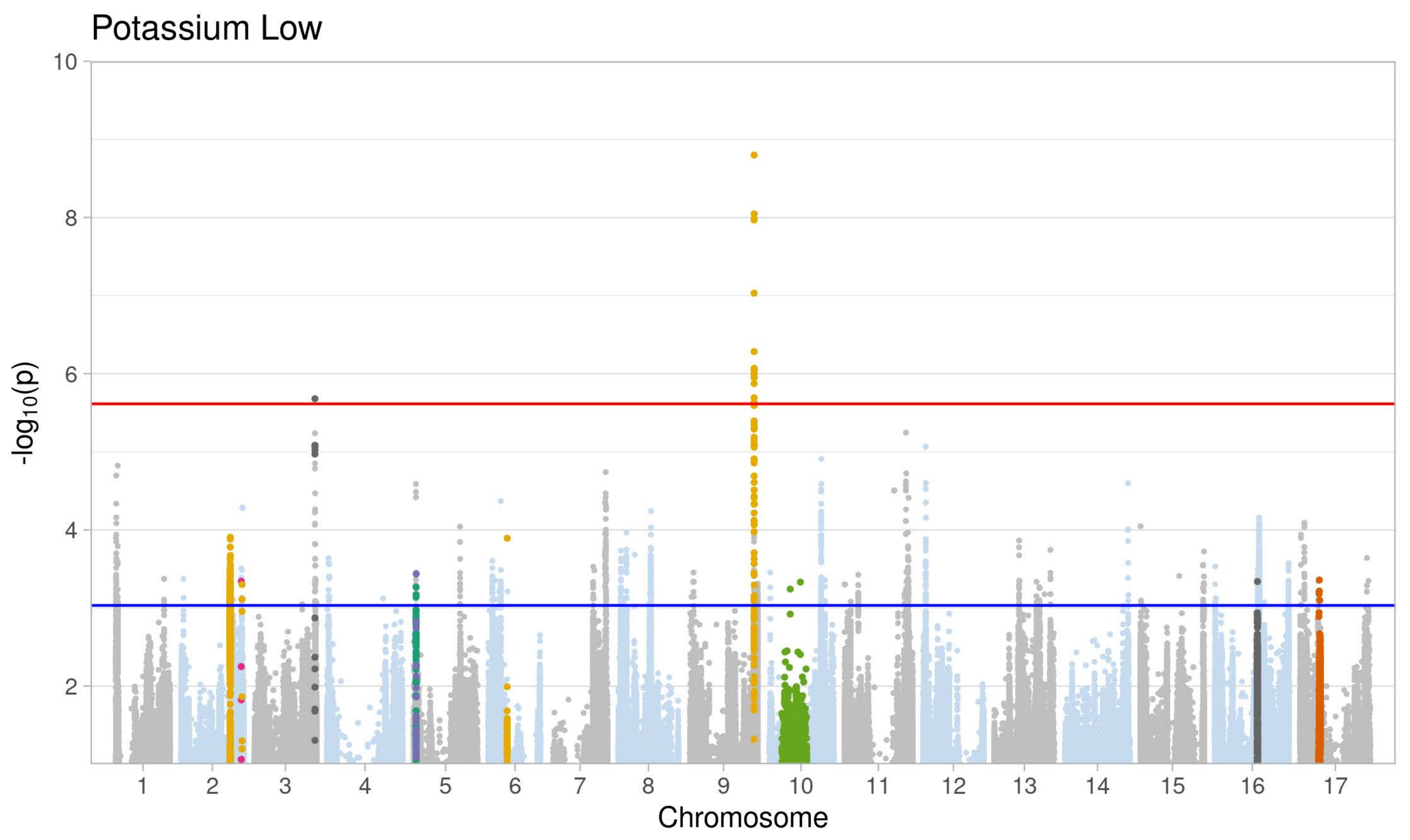
Potassium Low
-log10(p)
10
8
6
4
2
1 2 3 4 5 6 7 8 9 10 11 12 13 14 15 16 17
Chromosome

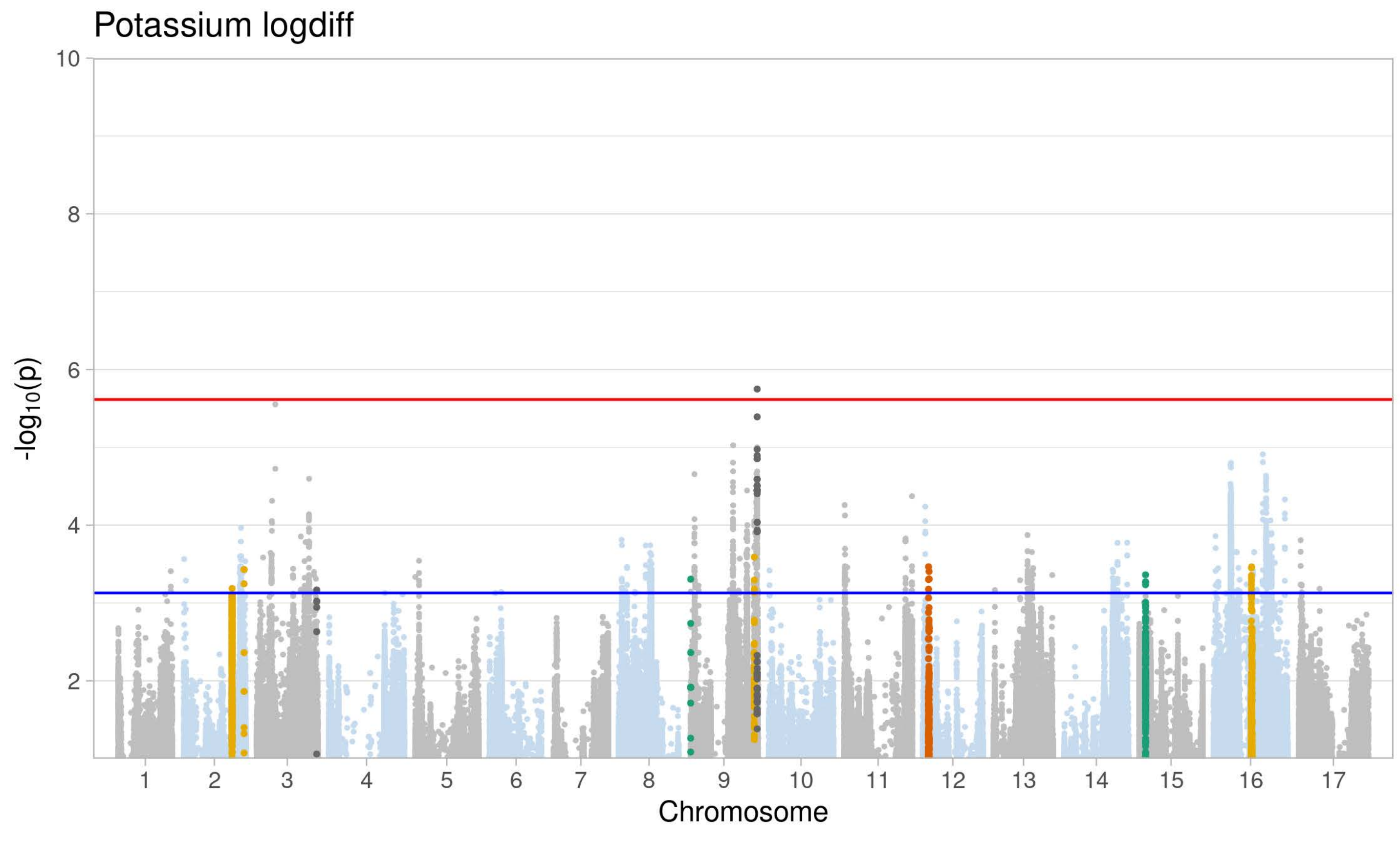
Potassium logdiff
-log10(p)
10
8
6
4
2
1 2 3 4 5 6 7 8 9 10 11 12 13 14 15 16 17
Chromosome

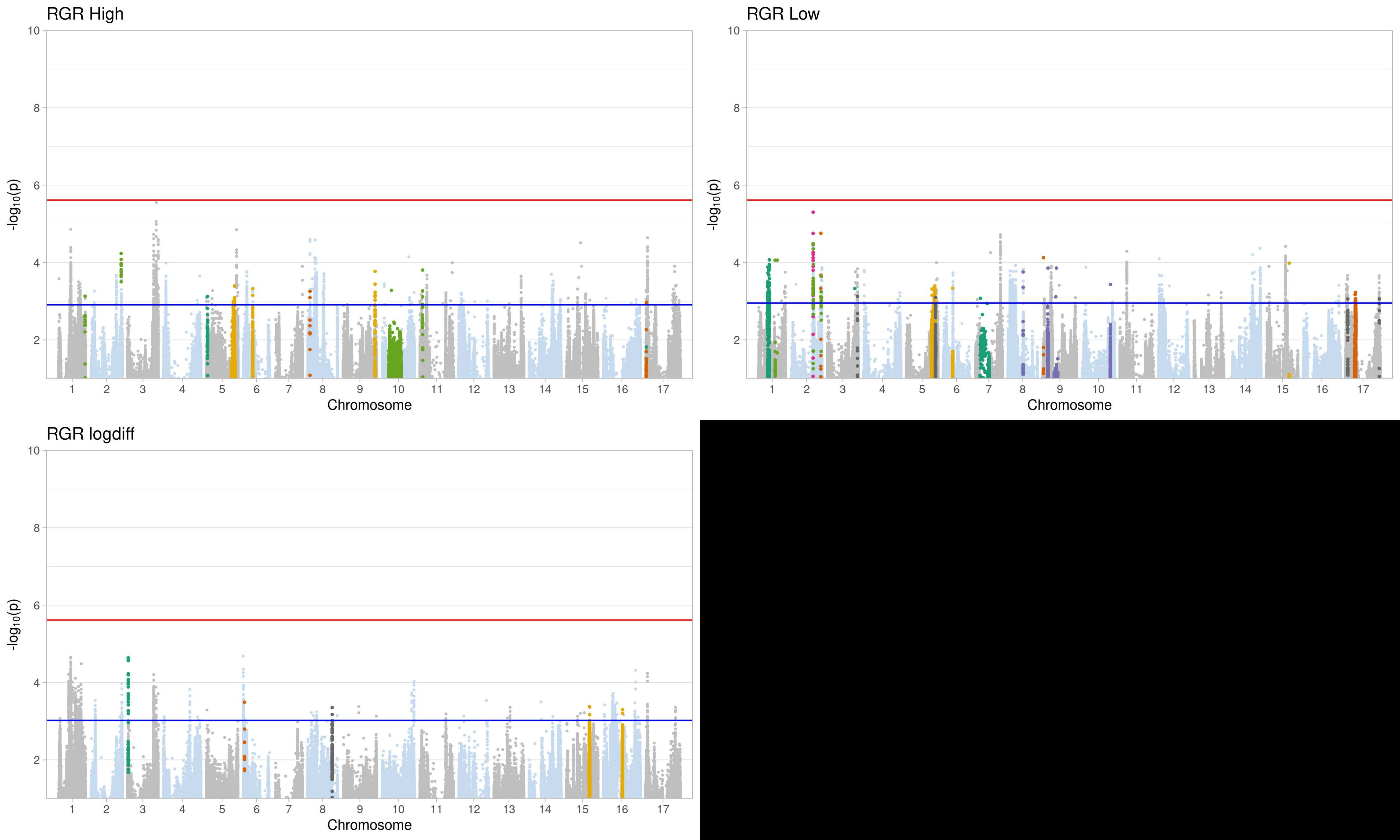
RGR High
RGR Low
RGR logdiff
-log10(p)
Chromosome
0 2 4 6 8 10
1 2 3 4 5 6 7 8 9 10 11 12 13 14 15 16 17

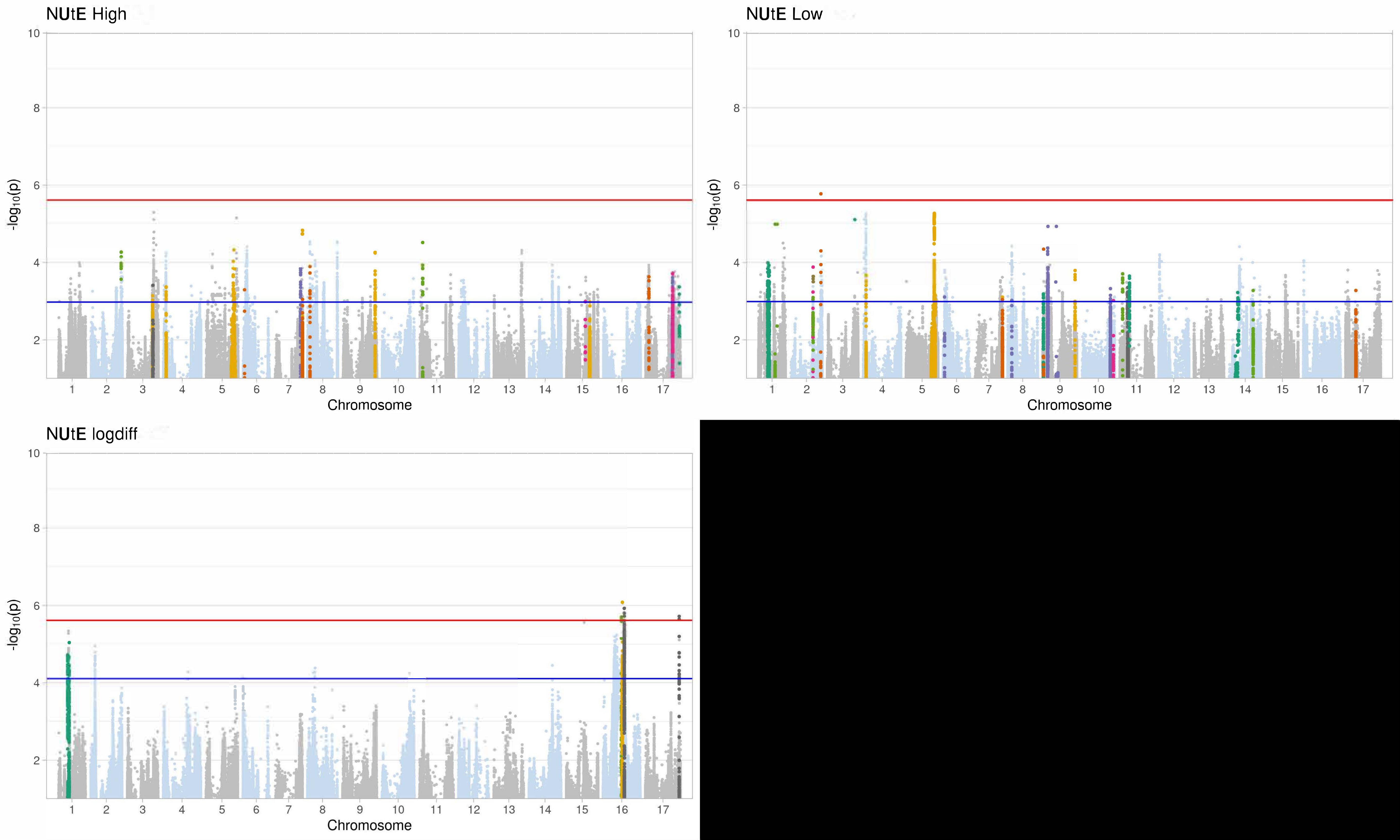

NUtE High
NUtE Low
NUtE logdiff
-log10(p)
Chromosome
1 2 3 4 5 6 7 8 9 10 11 12 13 14 15 16 17
2 4 6 8 10

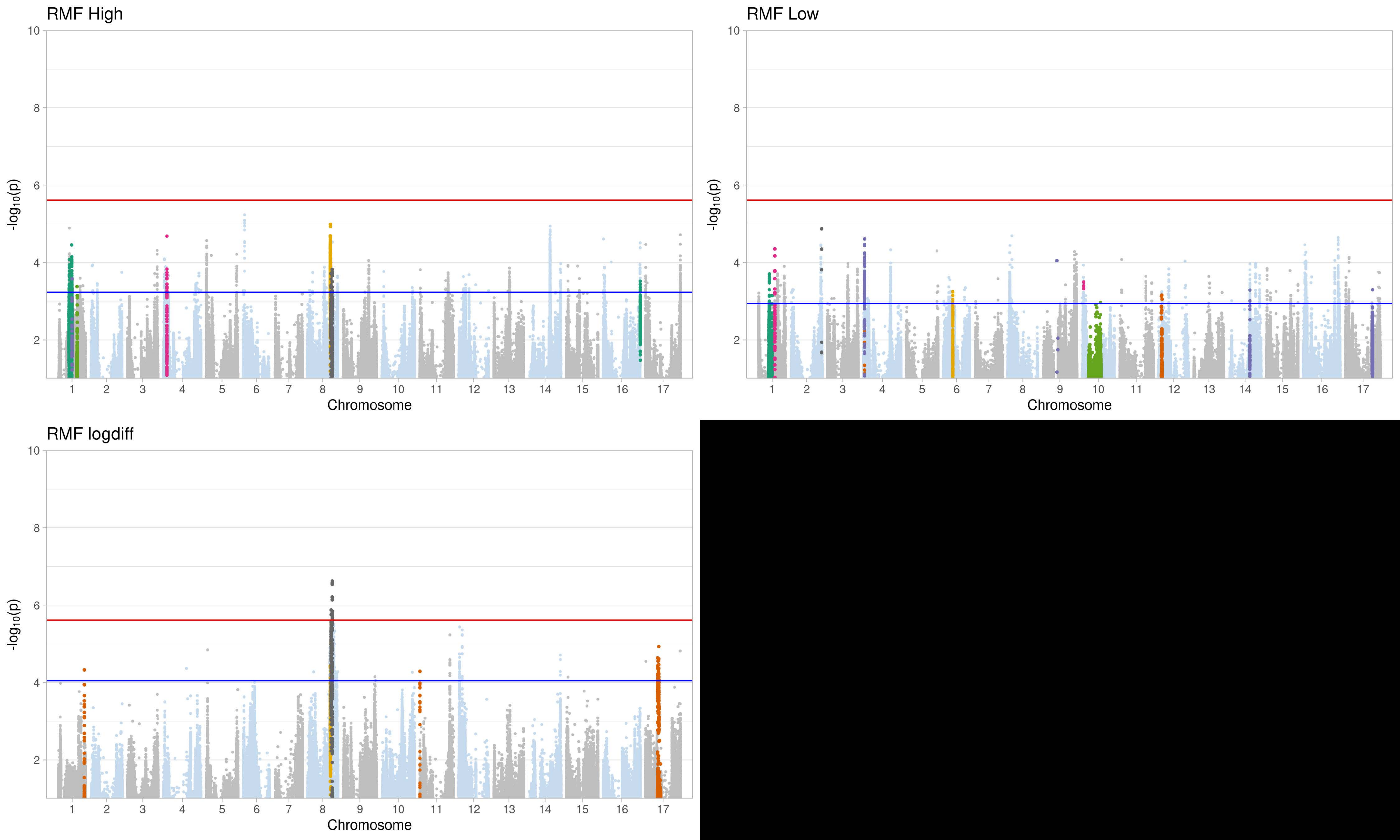
RMF High
RMF Low
RMF logdiff
-log10(p)
Chromosome
0
2
4
6
8
10
1 2 3 4 5 6 7 8 9 10 11 12 13 14 15 16 17

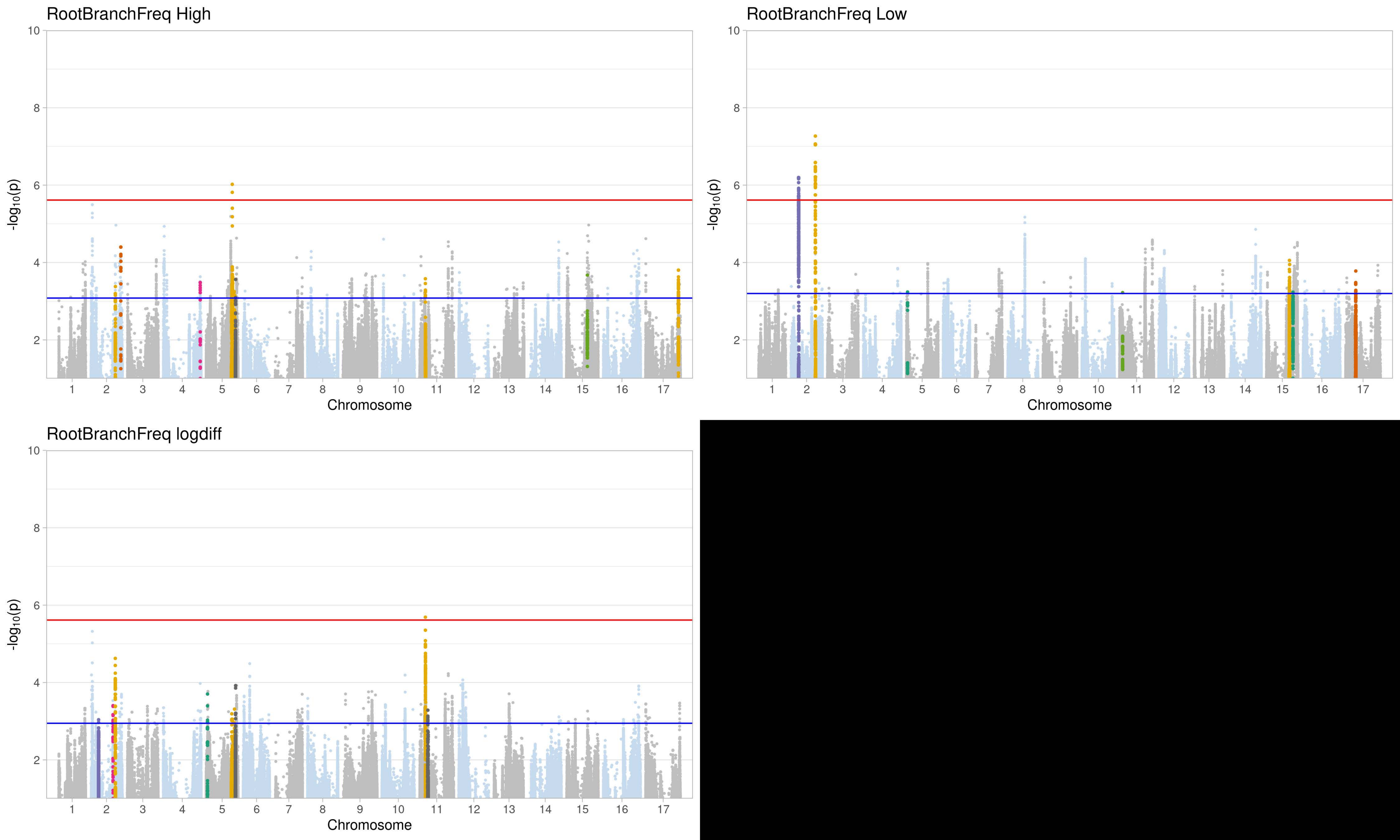

RootBranchFreq High
RootBranchFreq Low
RootBranchFreq logdiff
-log10(p)
Chromosome
10
8
6
4
2
1 2 3 4 5 6 7 8 9 10 11 12 13 14 15 16 17

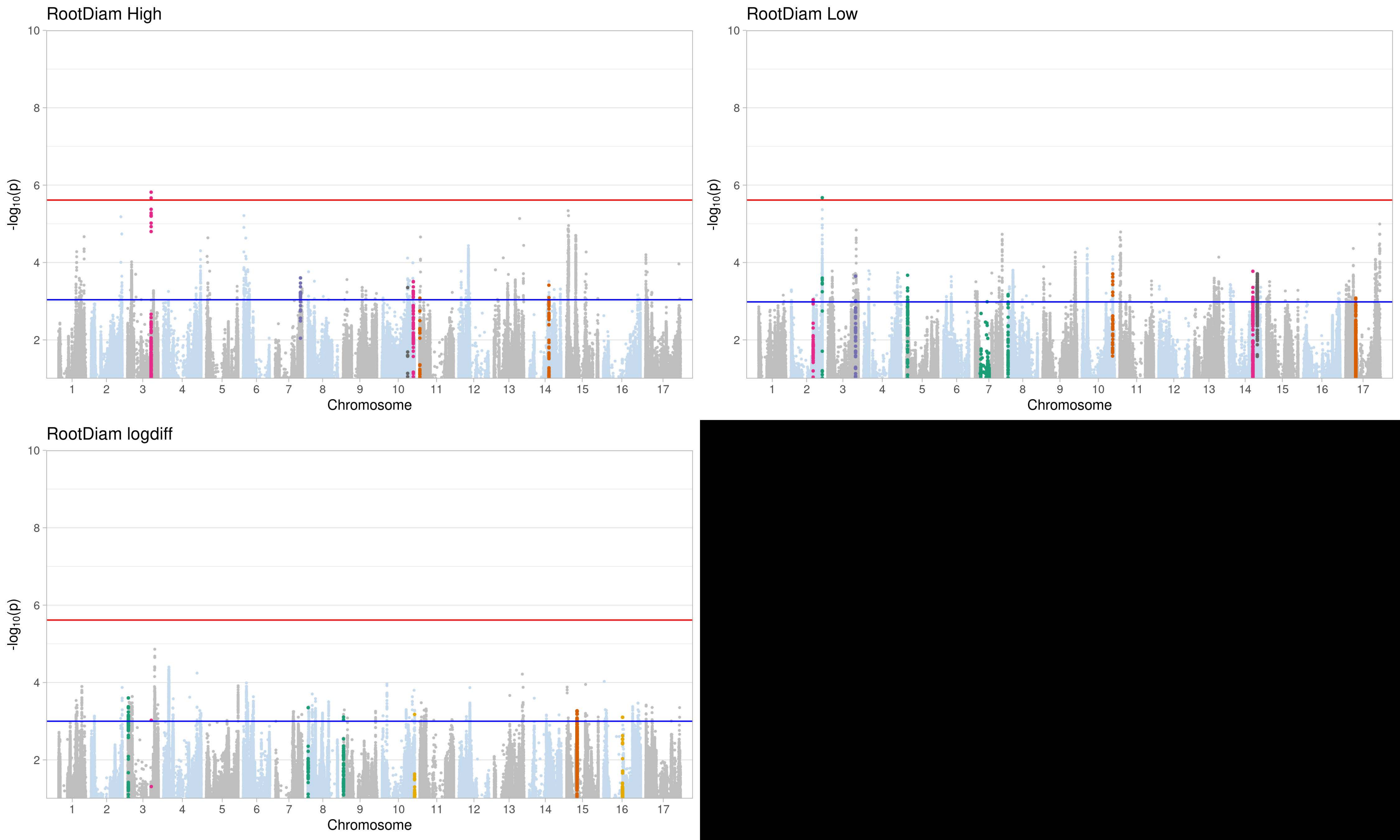
RootDiam High
RootDiam Low
RootDiam logdiff
-log10(p)
Chromosome
10
8
6
4
2
1 2 3 4 5 6 7 8 9 10 11 12 13 14 15 16 17

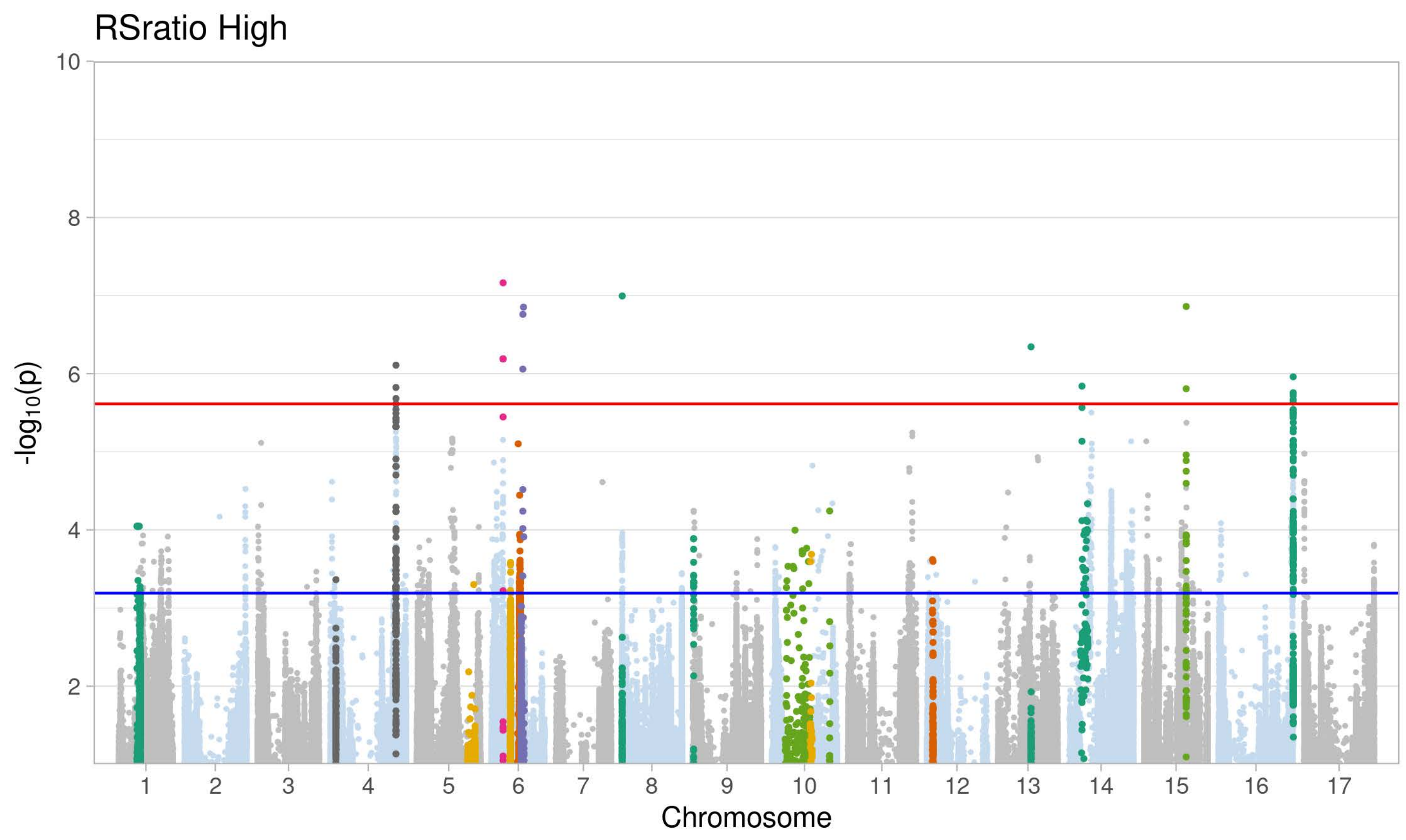
RSratio High
-log10(p)
Chromosome
1 2 3 4 5 6 7 8 9 10 11 12 13 14 15 16 17
10 8 6 4 2

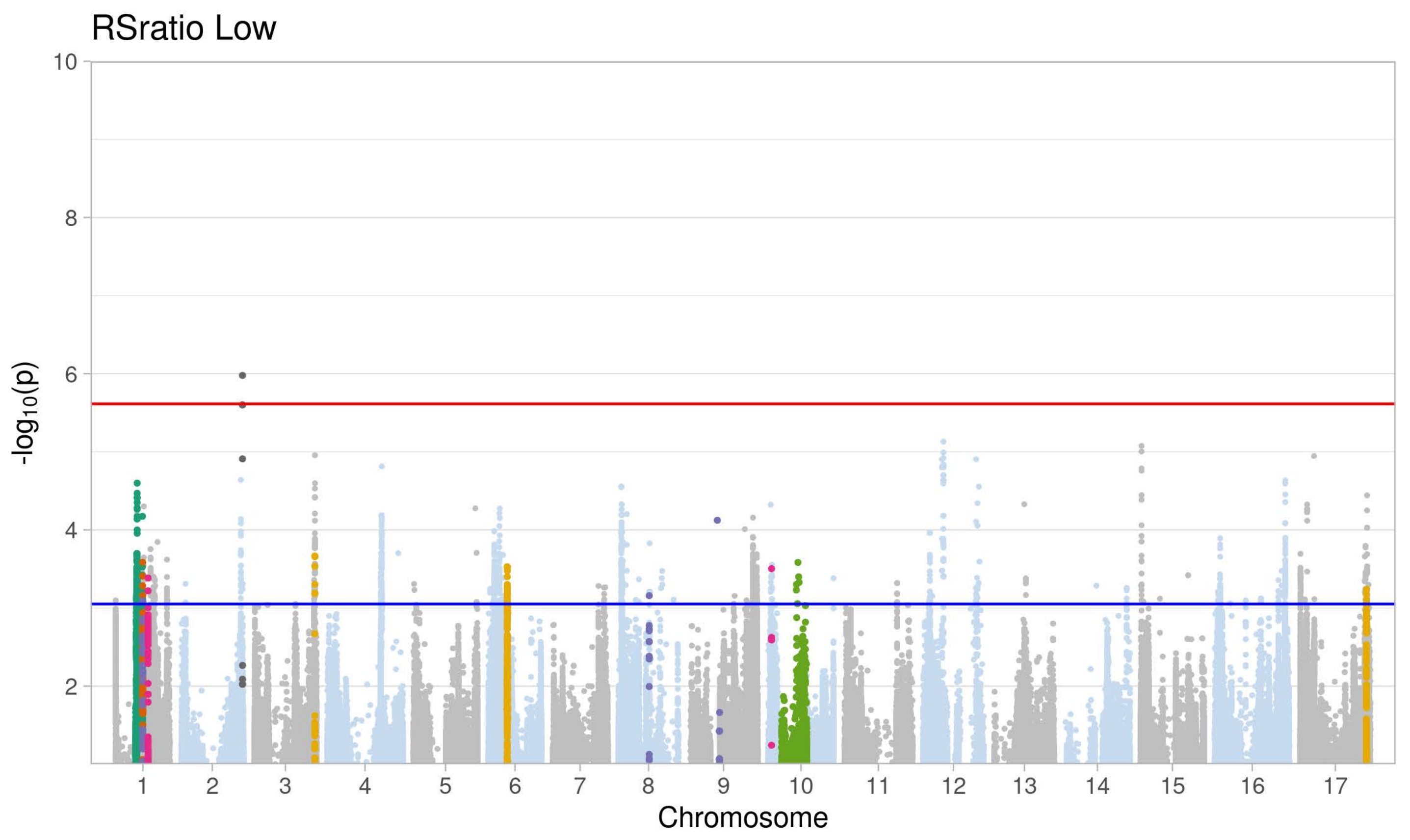
RSratio Low
-log10(p)
Chromosome
1 2 3 4 5 6 7 8 9 10 11 12 13 14 15 16 17
10 8 6 4 2

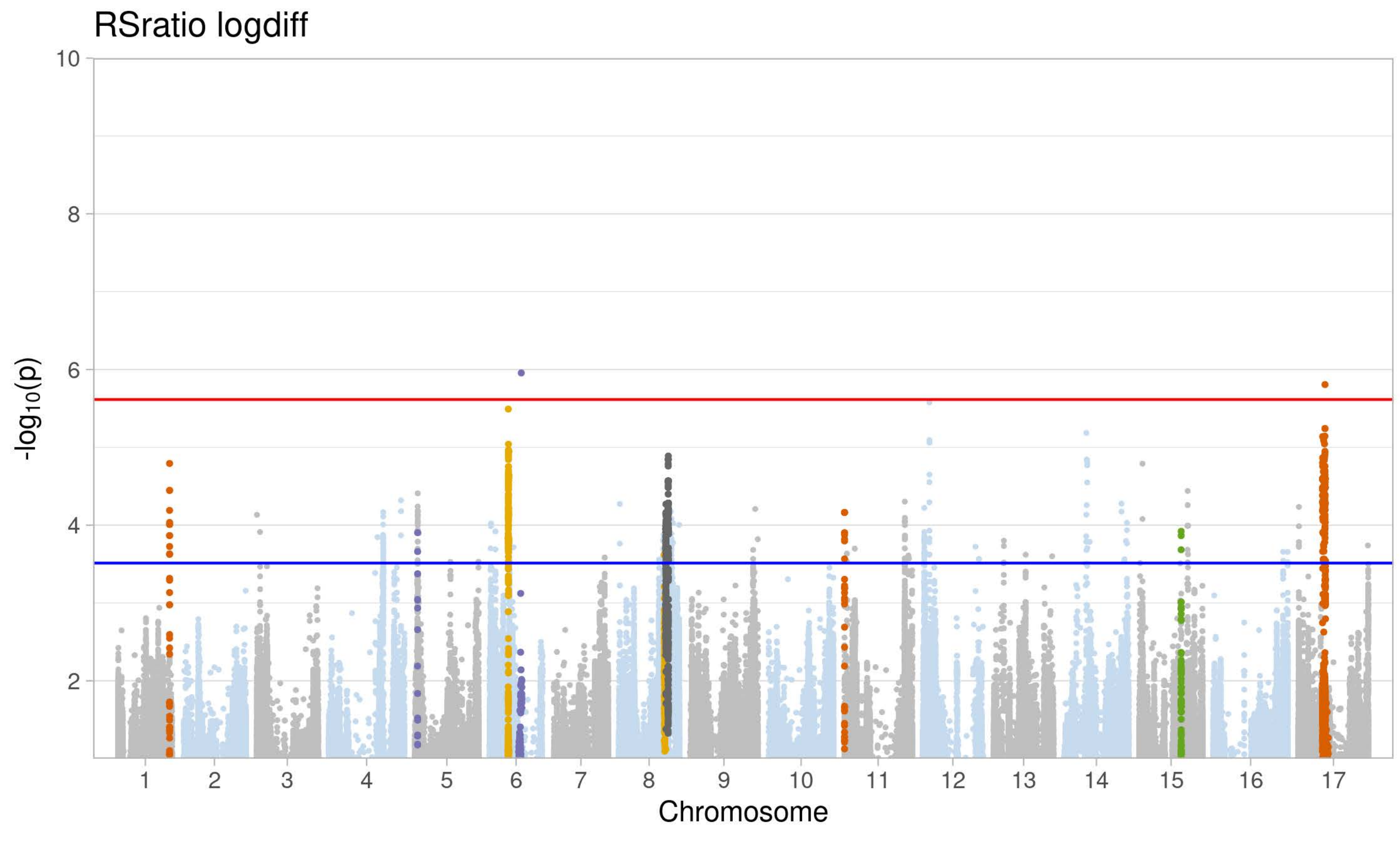
RSratio logdiff
-log10(p)
Chromosome
1 2 3 4 5 6 7 8 9 10 11 12 13 14 15 16 17
10 8 6 4 2

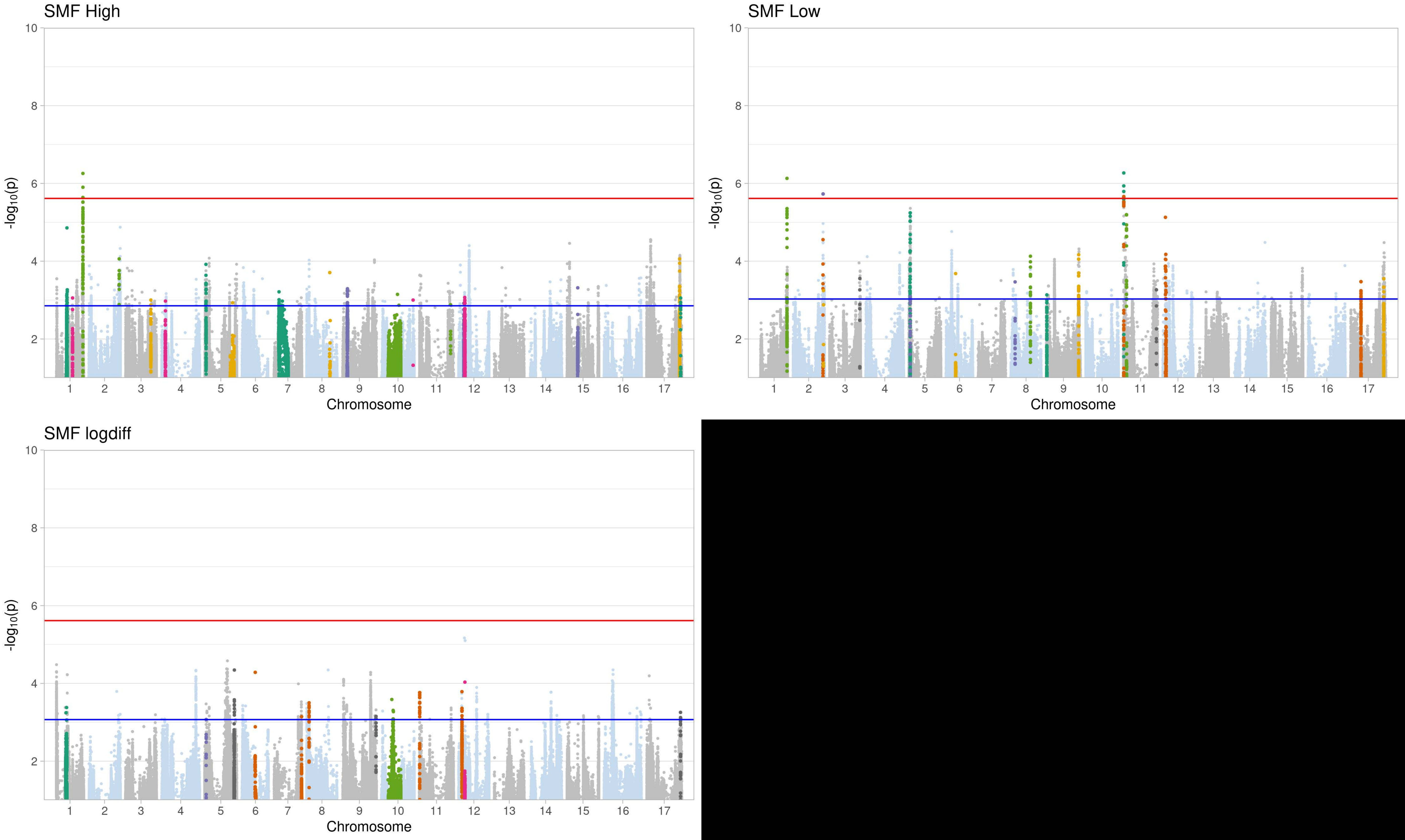
SMF High
SMF Low
SMF logdiff
-log10(p)
Chromosome
1 2 3 4 5 6 7 8 9 10 11 12 13 14 15 16 17
2 4 6 8 10

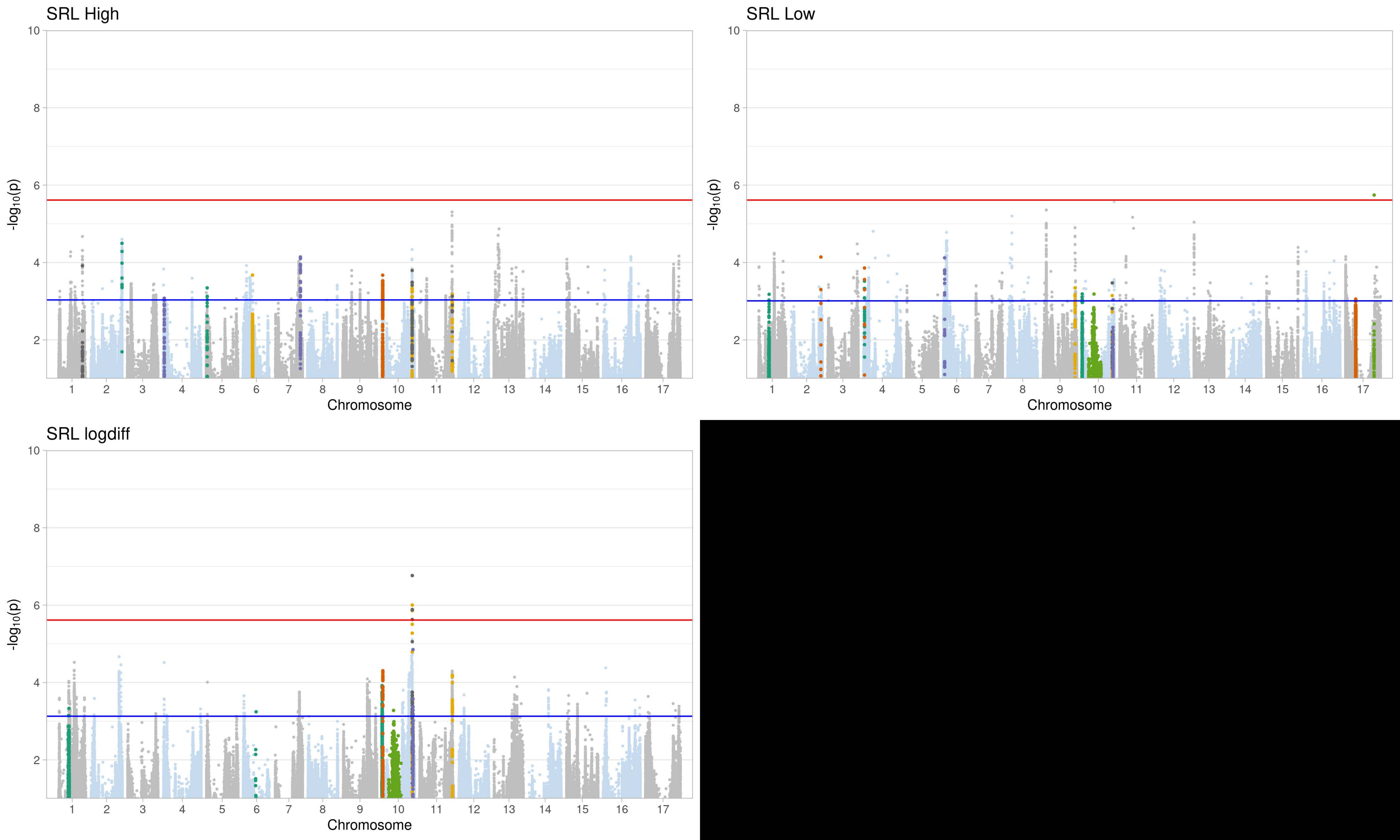
SRL High
SRL Low
SRL logdiff
-log10(p)
Chromosome
10
8
6
4
2
1 2 3 4 5 6 7 8 9 10 11 12 13 14 15 16 17

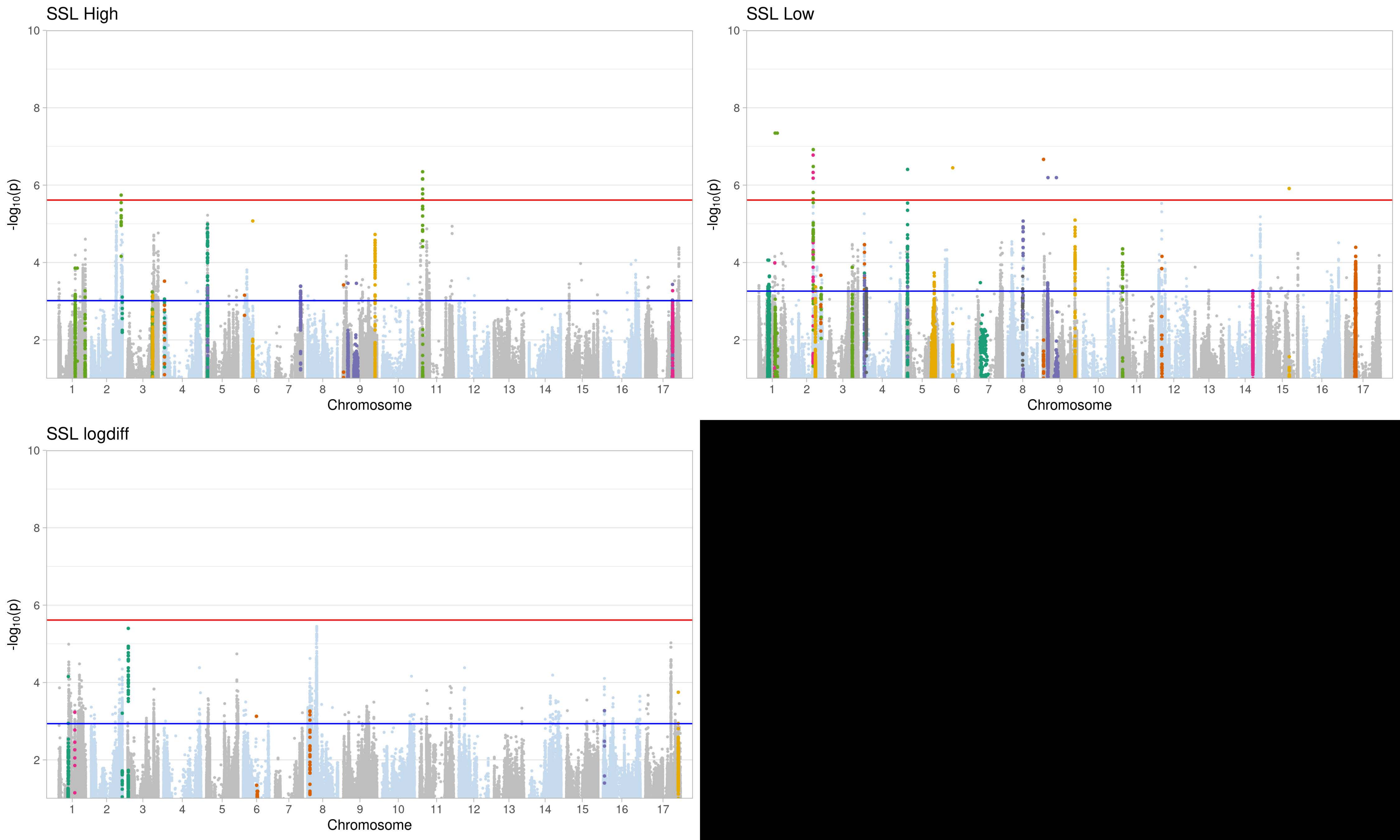
SSL High
SSL Low
SSL logdiff
-log10(p)
Chromosome
10
8
6
4
2
1 2 3 4 5 6 7 8 9 10 11 12 13 14 15 16 17

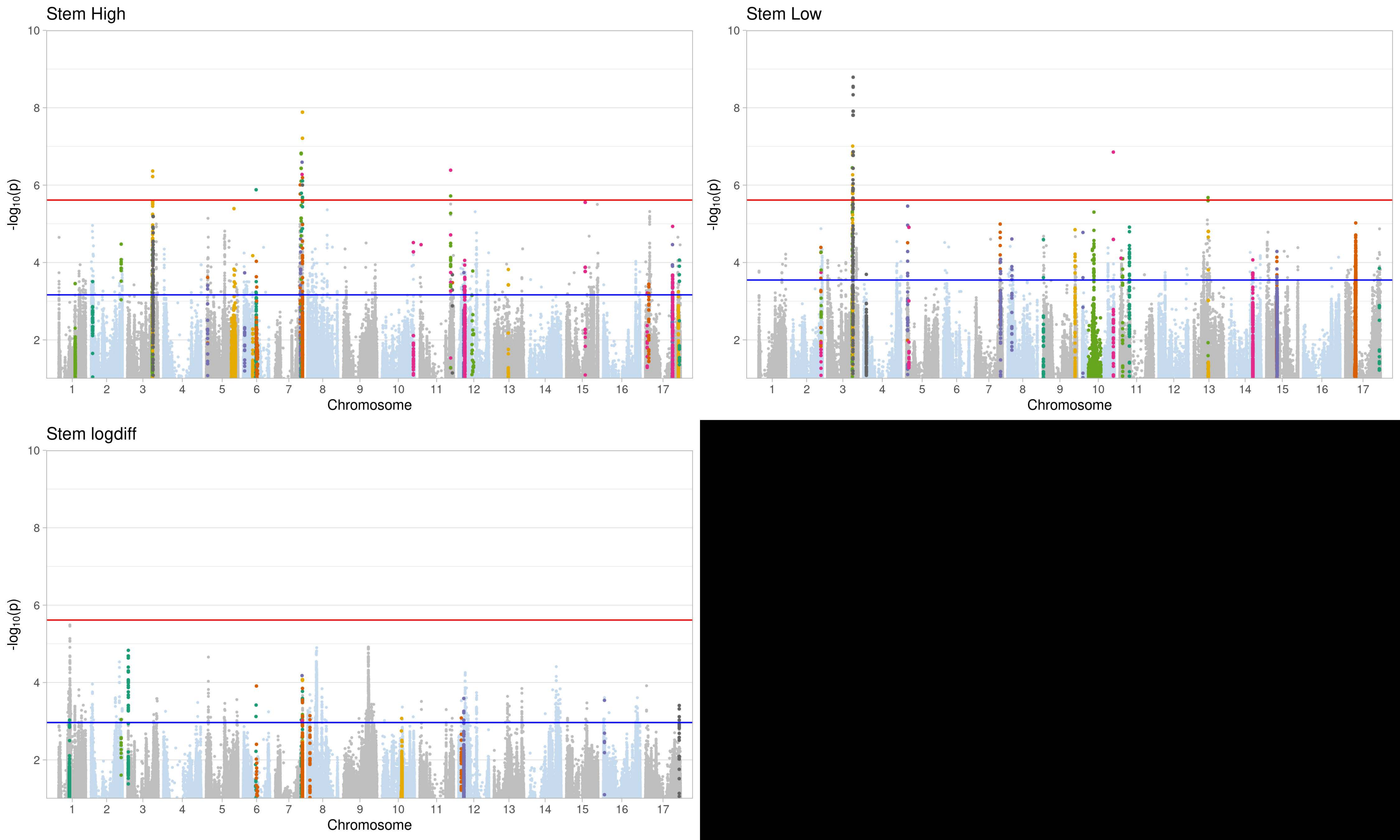
Stem High
Stem Low
Stem logdiff
-log10(p)
Chromosome
10
8
6
4
2

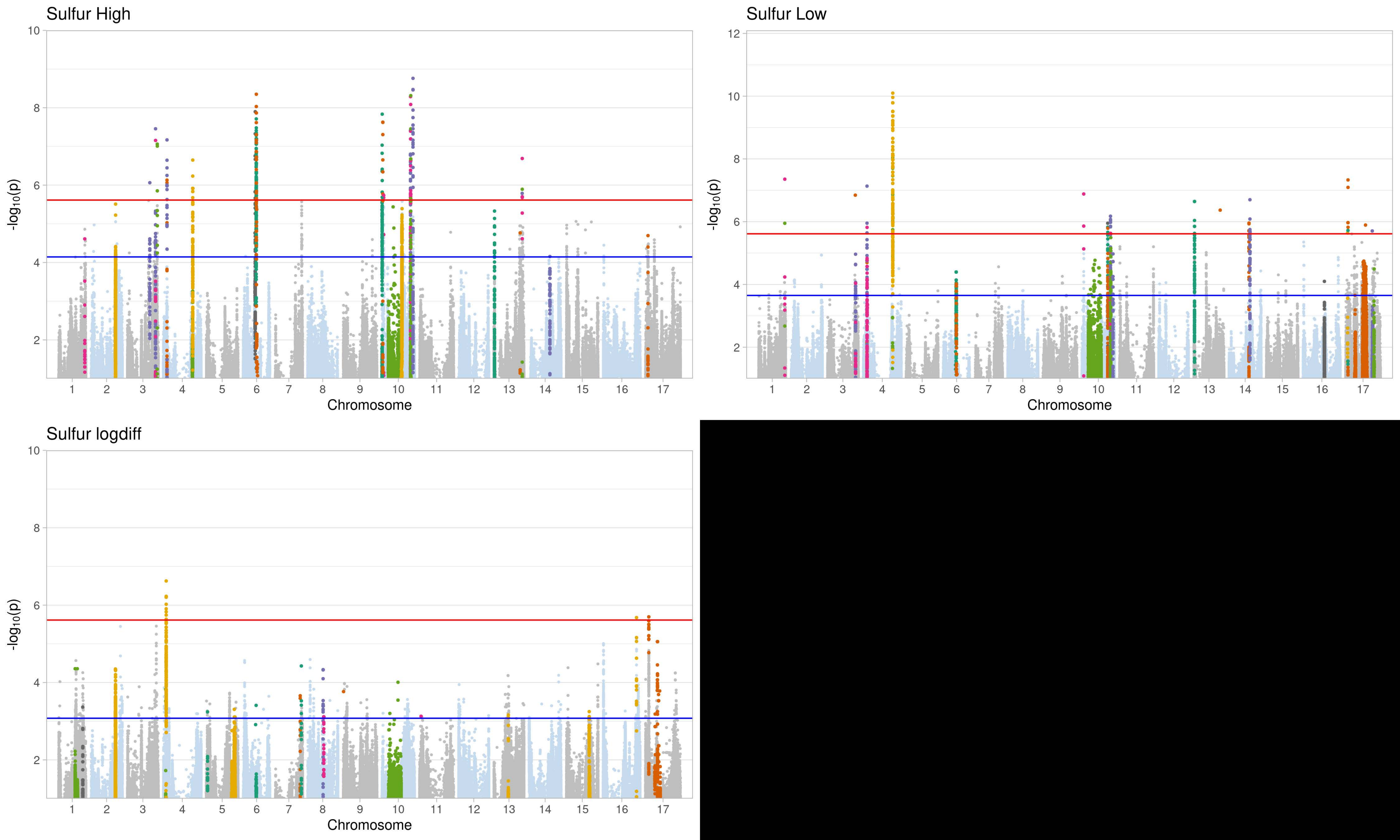
Sulfur High
-log10(p)
Chromosome
Sulfur Low
-log10(p)
Chromosome
Sulfur logdiff
-log10(p)
Chromosome

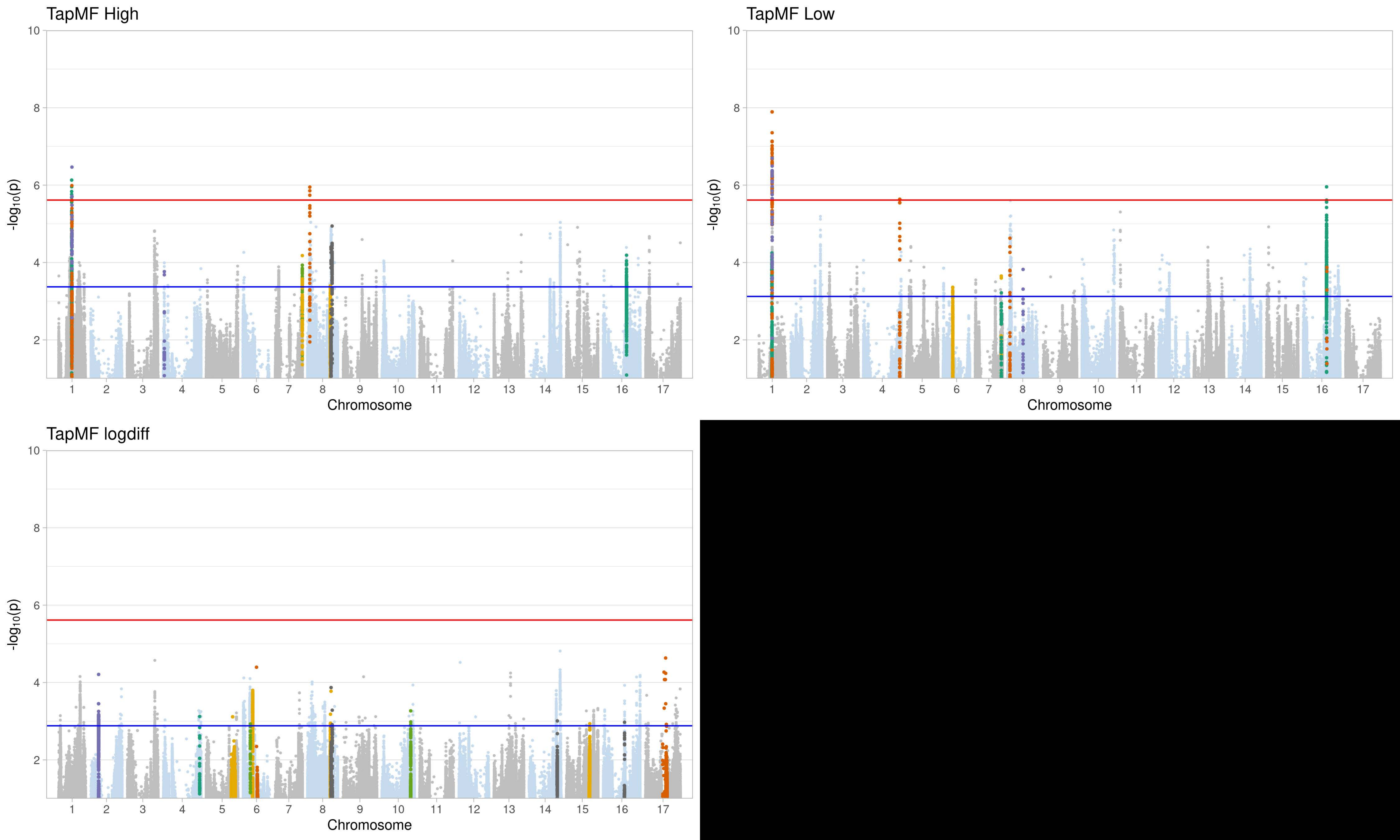
TapMF High
TapMF Low
TapMF logdiff
-log10(p)
Chromosome
1 2 3 4 5 6 7 8 9 10 11 12 13 14 15 16 17
10 8 6 4 2

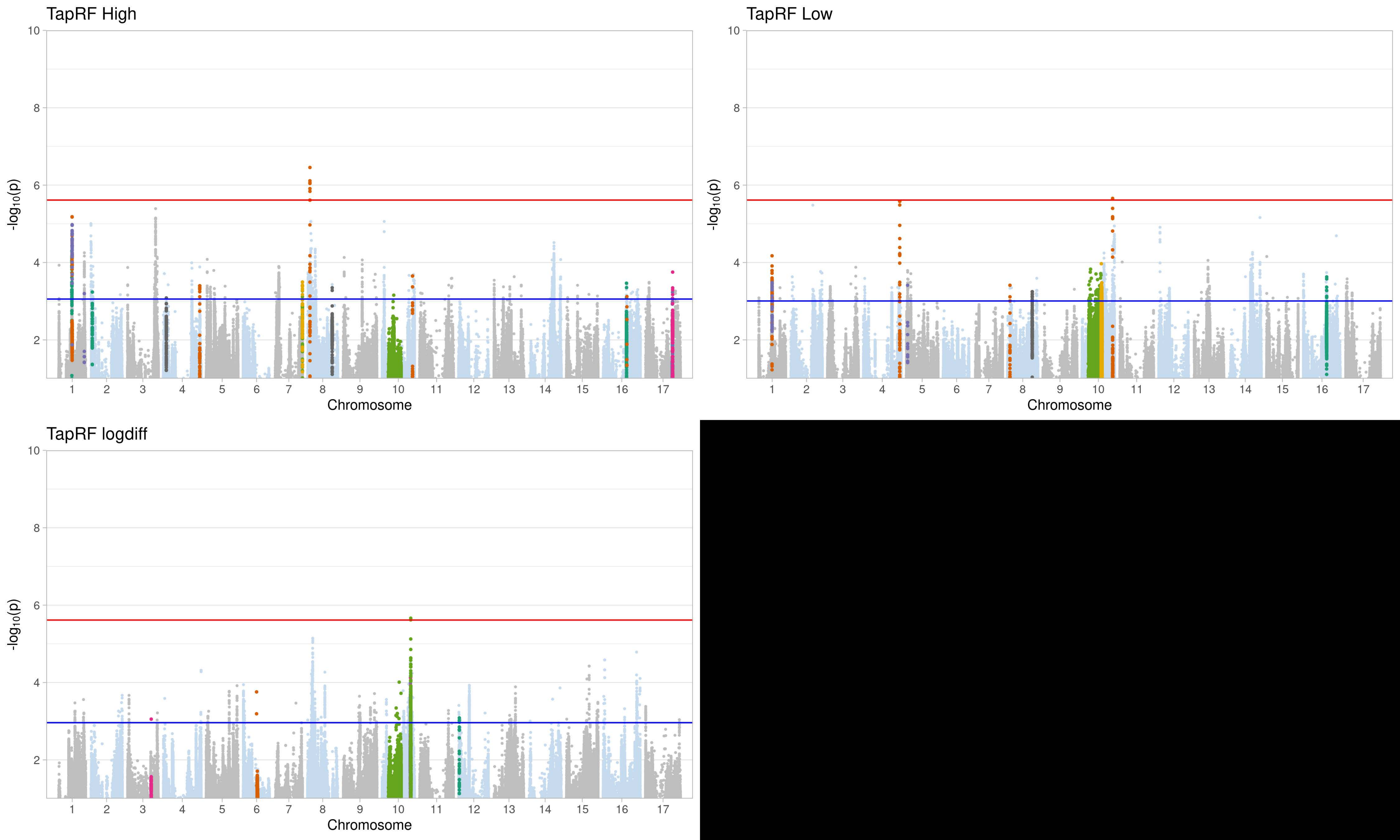
TapRF High
TapRF Low
TapRF logdiff
-log10(p)
10
8
6
4
2
Chromosome
1 2 3 4 5 6 7 8 9 10 11 12 13 14 15 16 17

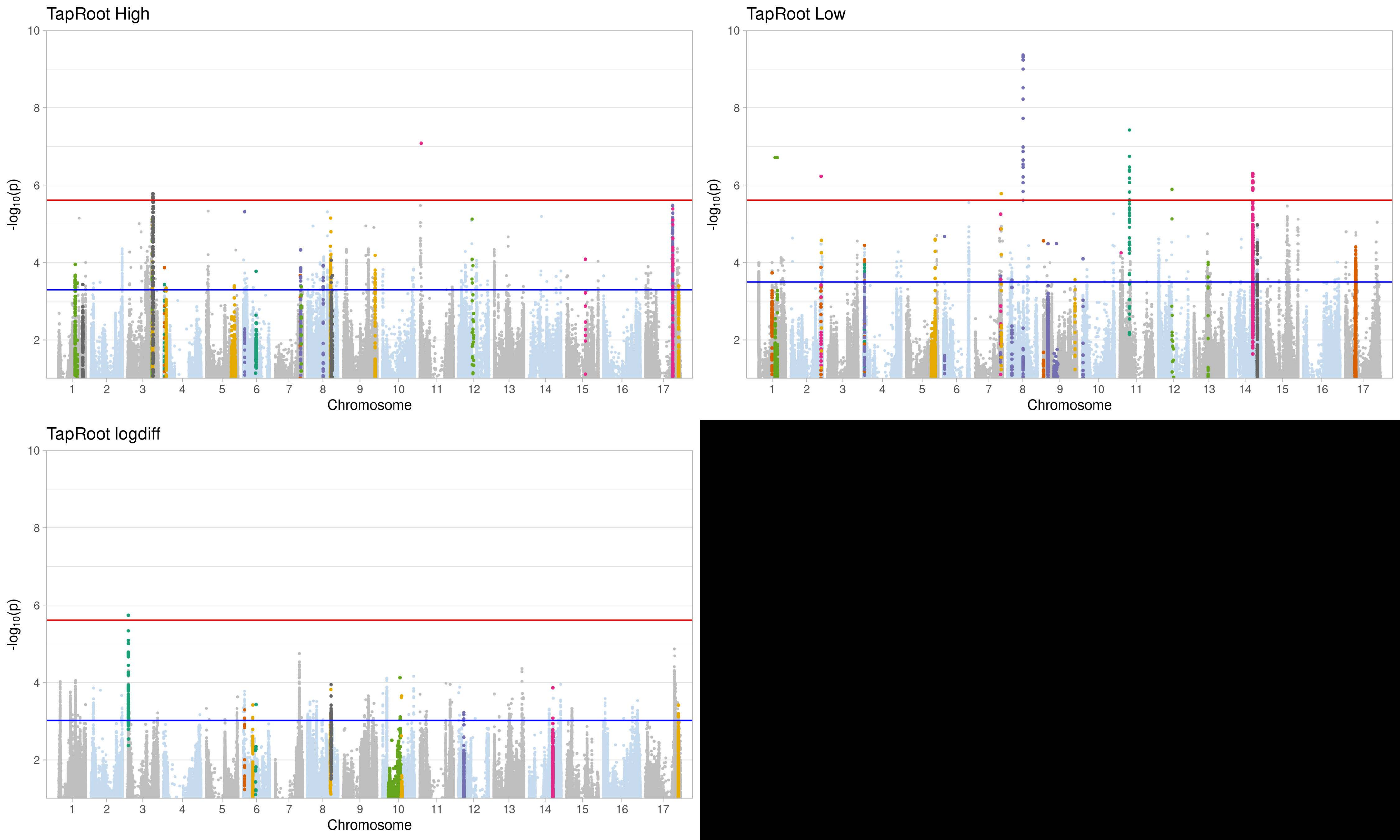
TapRoot High
TapRoot Low
TapRoot logdiff
-log10(p)
Chromosome
1 2 3 4 5 6 7 8 9 10 11 12 13 14 15 16 17
2 4 6 8 10

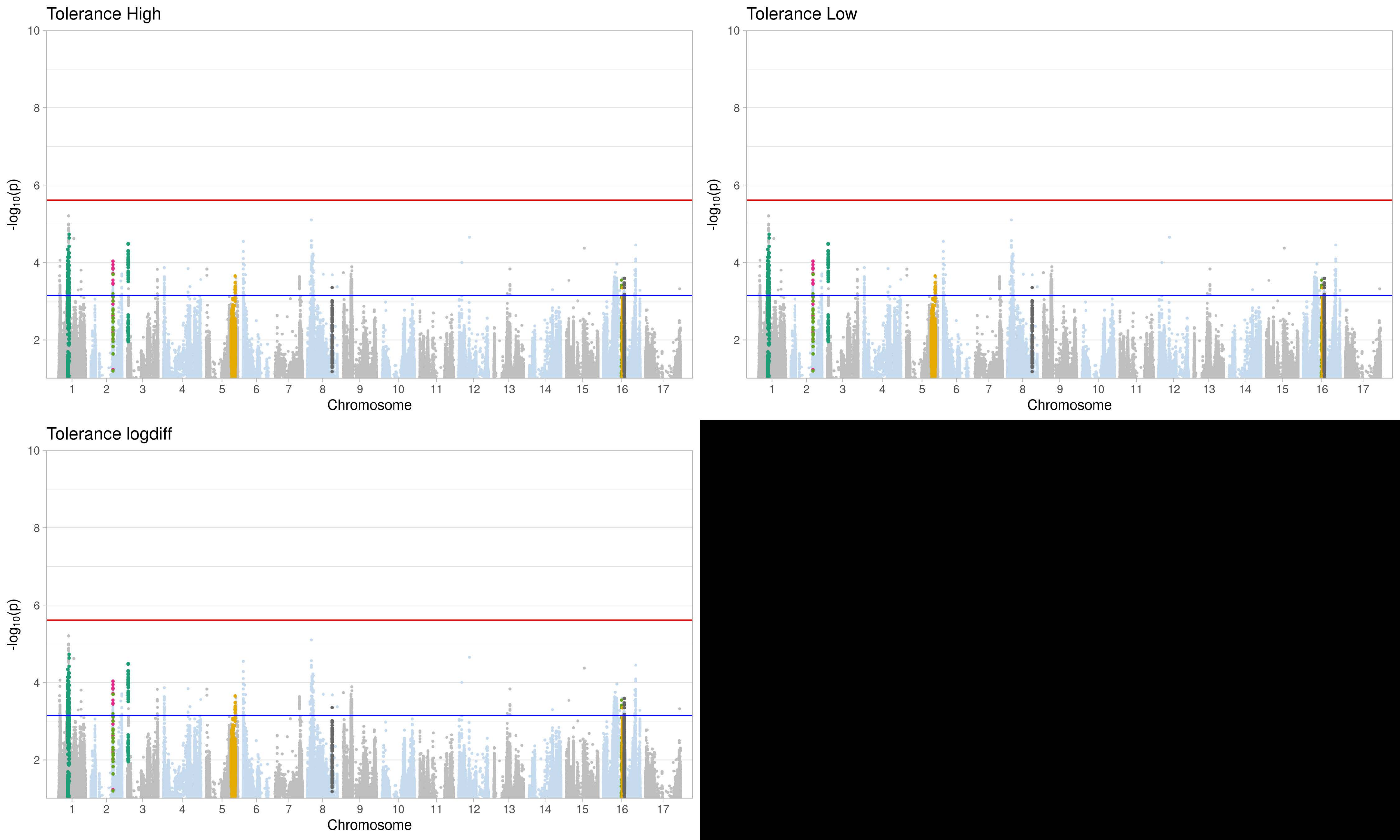
Tolerance High
Tolerance Low
Tolerance logdiff
-log10(p)
Chromosome
1 2 3 4 5 6 7 8 9 10 11 12 13 14 15 16 17
2 4 6 8 10

deltaN**U**t**E**de**v** High

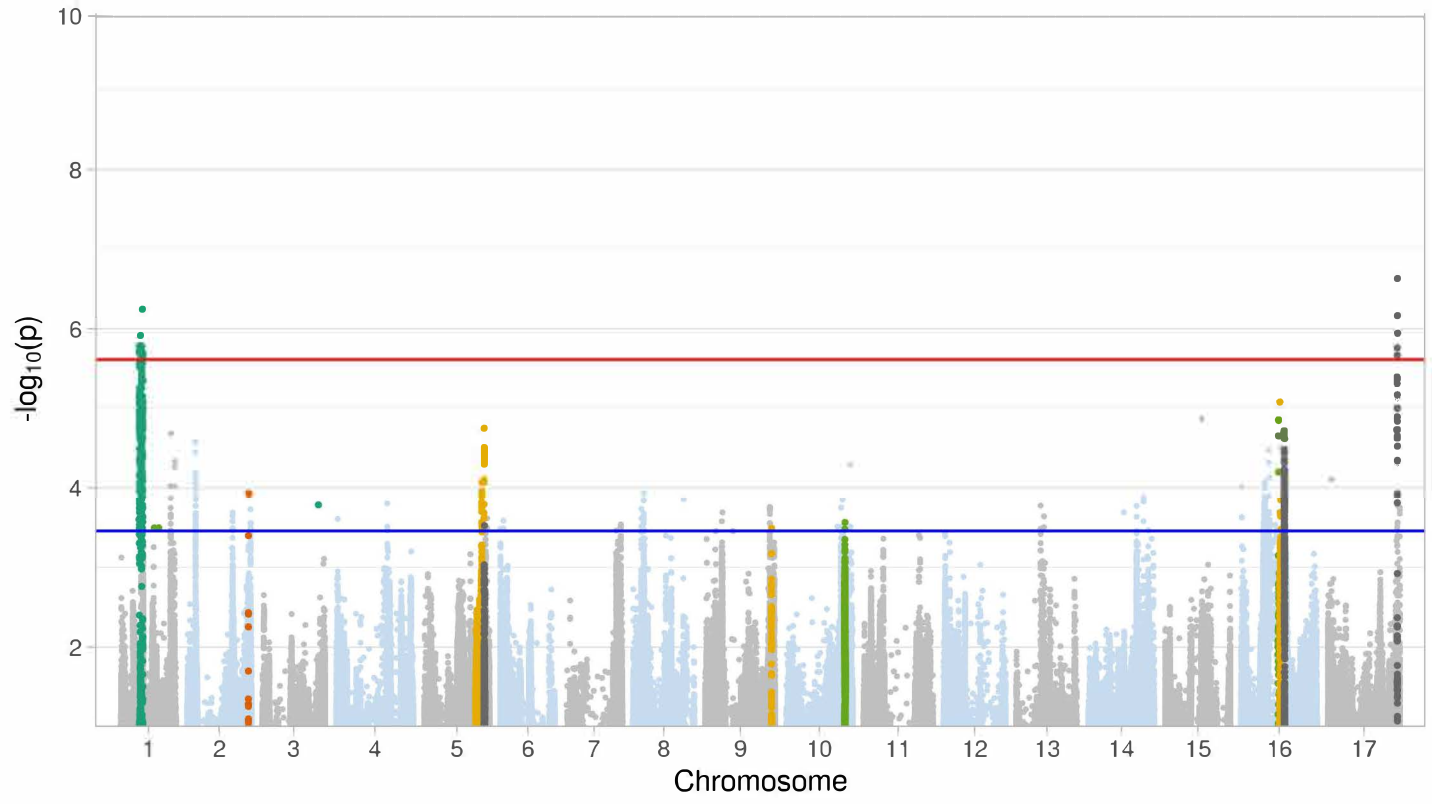

deltaN**U**t**E**de**v** Low

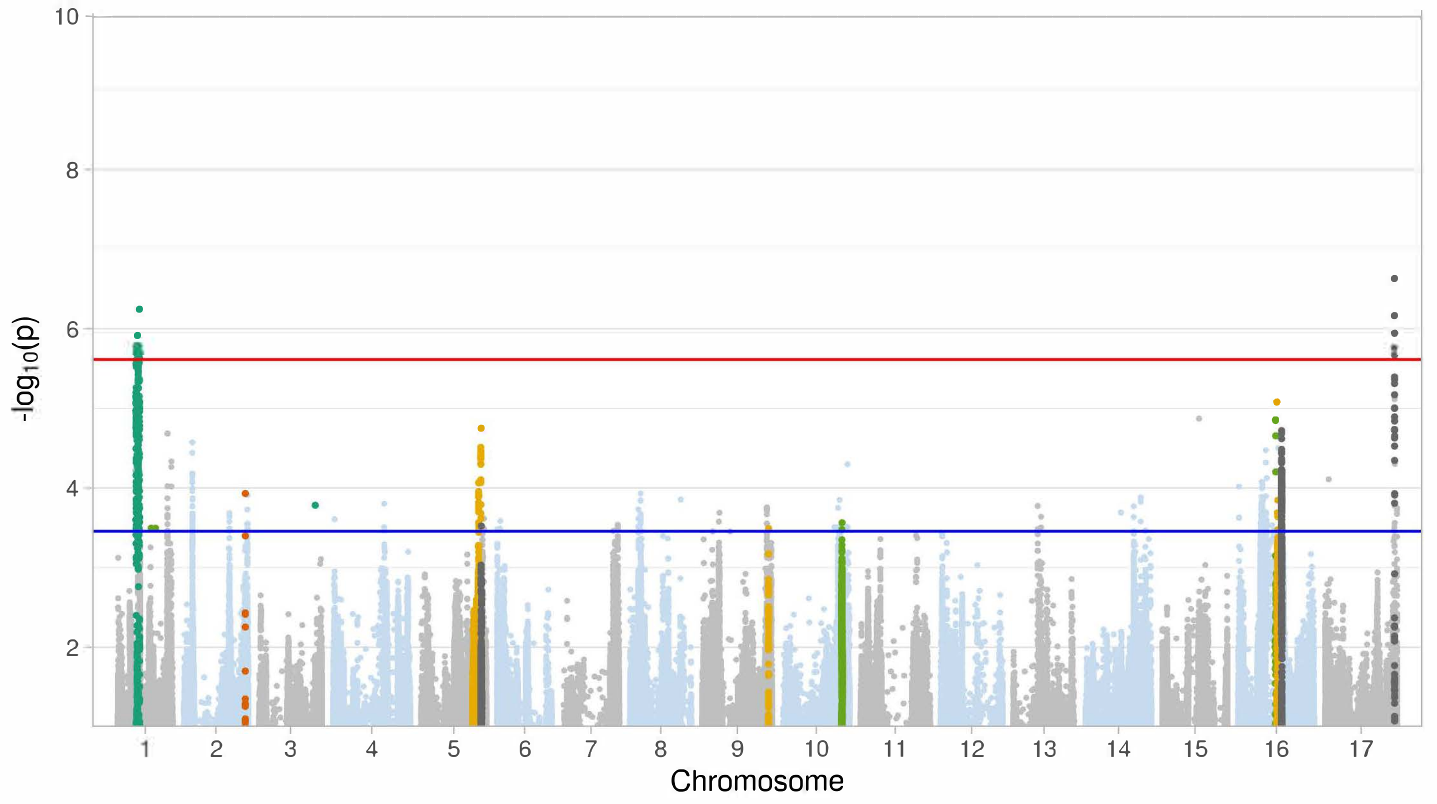

deltaN**U**t**E**de**v** logdiff

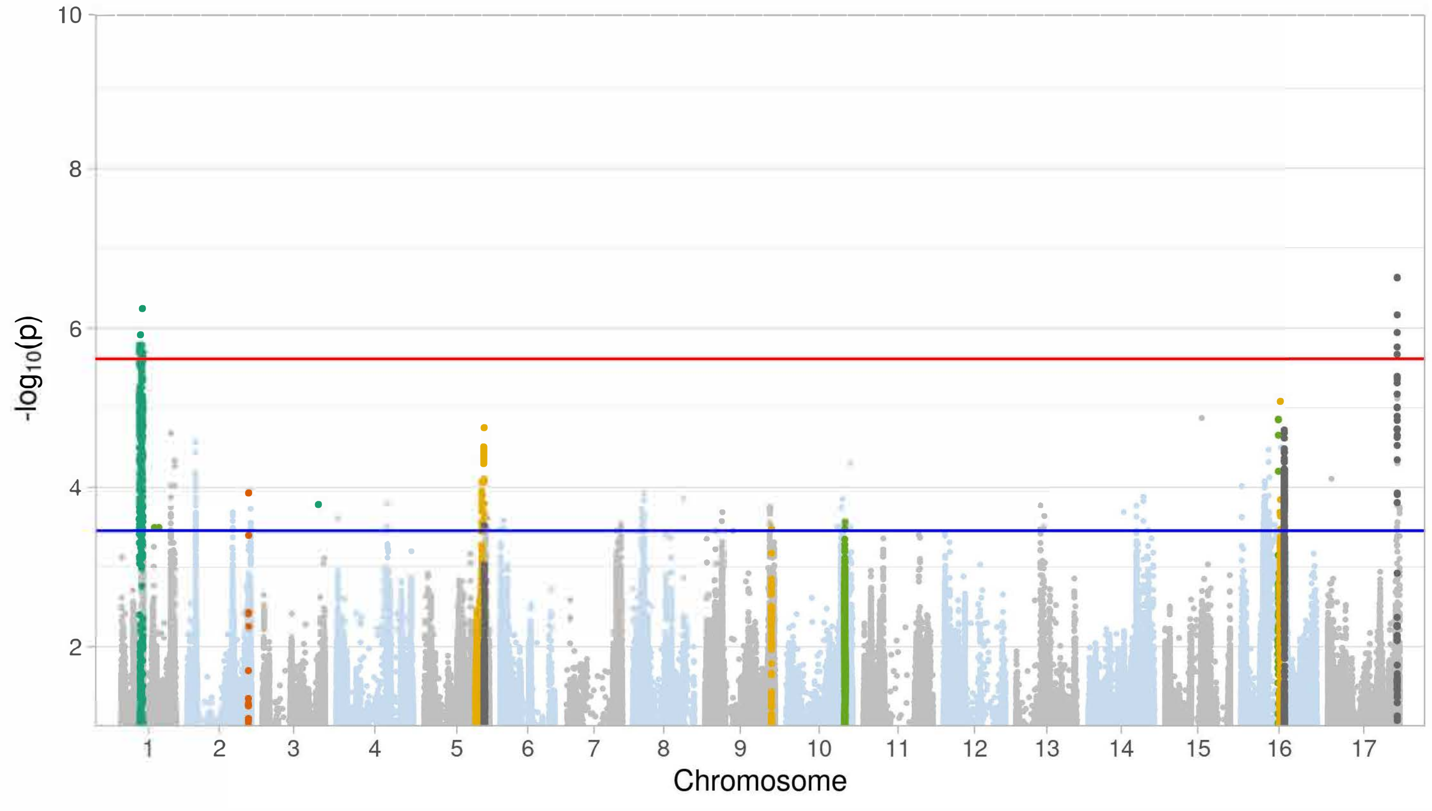

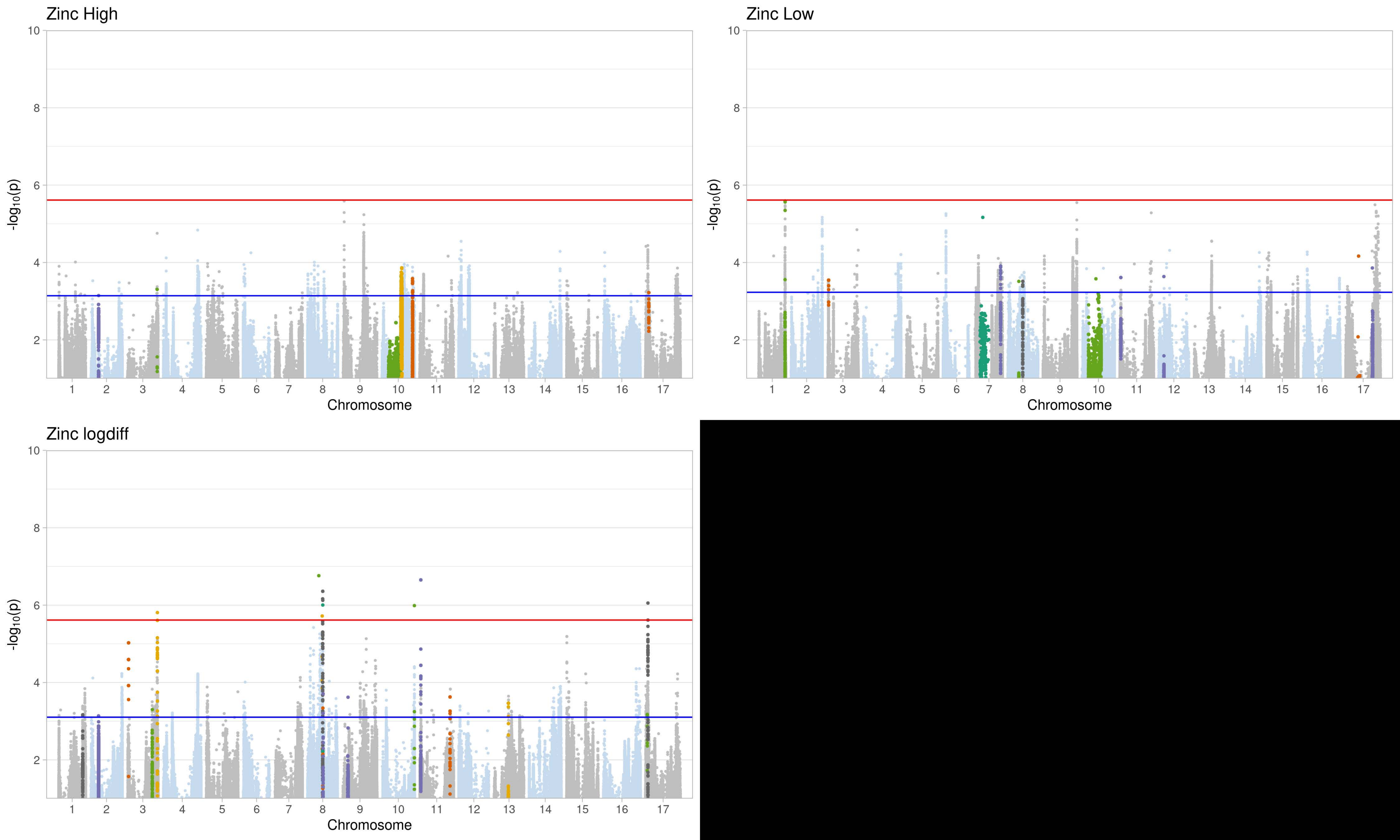

Zinc High
Zinc Low
Zinc logdiff
-log10(p)
Chromosome
10
8
6
4
2
1 2 3 4 5 6 7 8 9 10 11 12 13 14 15 16 17

Figure S5 **LD plots per chromosome** Linkage disequilibrium (estimated as $R^2$) between all SNPs that are significant for at least one trait within and between both treatments. Colored bar notes the haplotype blocks SNPs belong to based on the haplotype map presented in Fig. S7. Block membership is indicated for the calculation based on the genome-wide collection of SNPs (“genome”) and for the re-calculation based on just the significant SNPs (“significant”).

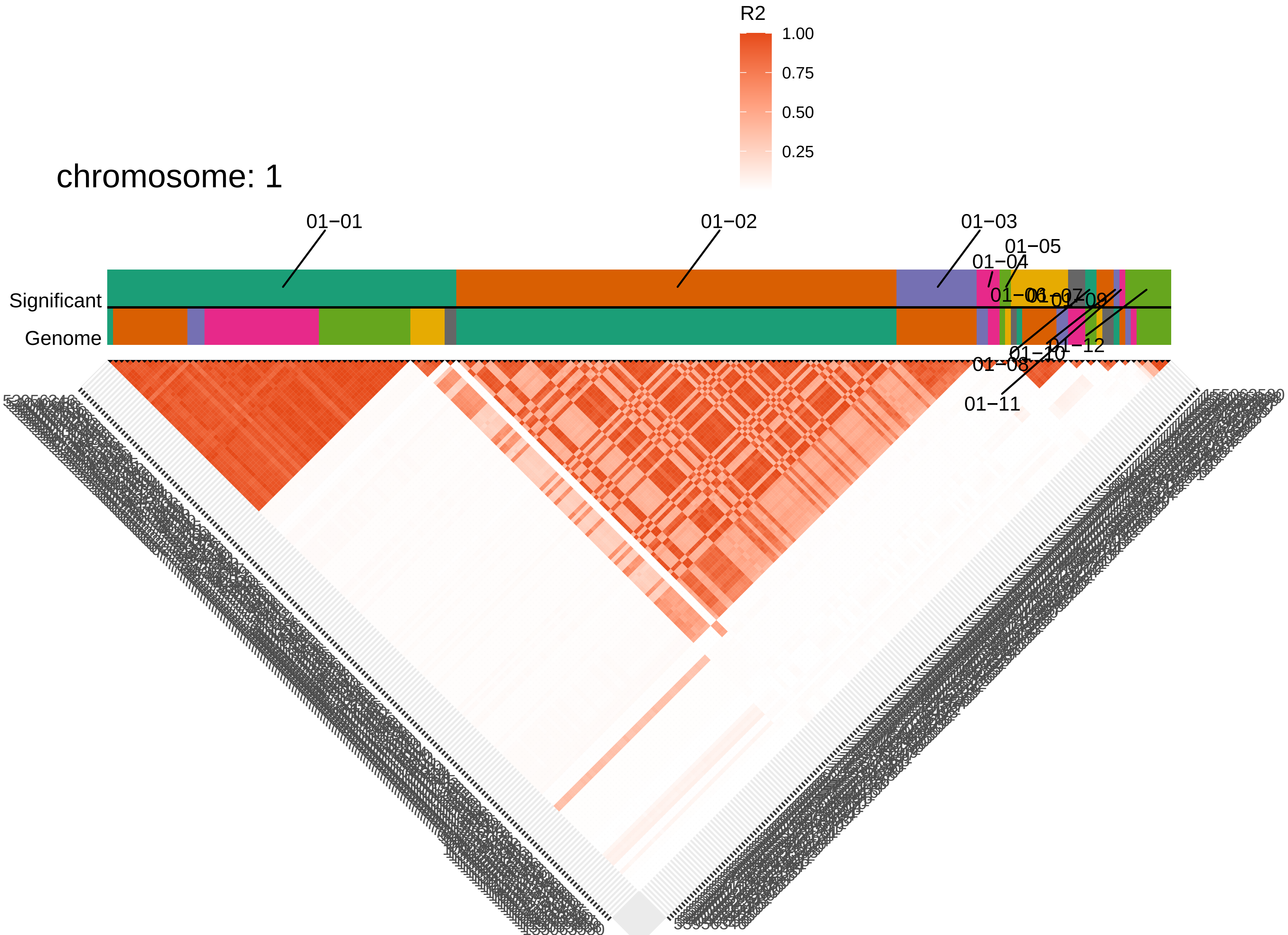
R2
1.00
0.75
0.50
0.25
chromosome: 1
01−01
01−02
01−03
01−04
01−05
01−06
01−07
01−08
01−09
01−10
01−11
01−12
Significant
Genome

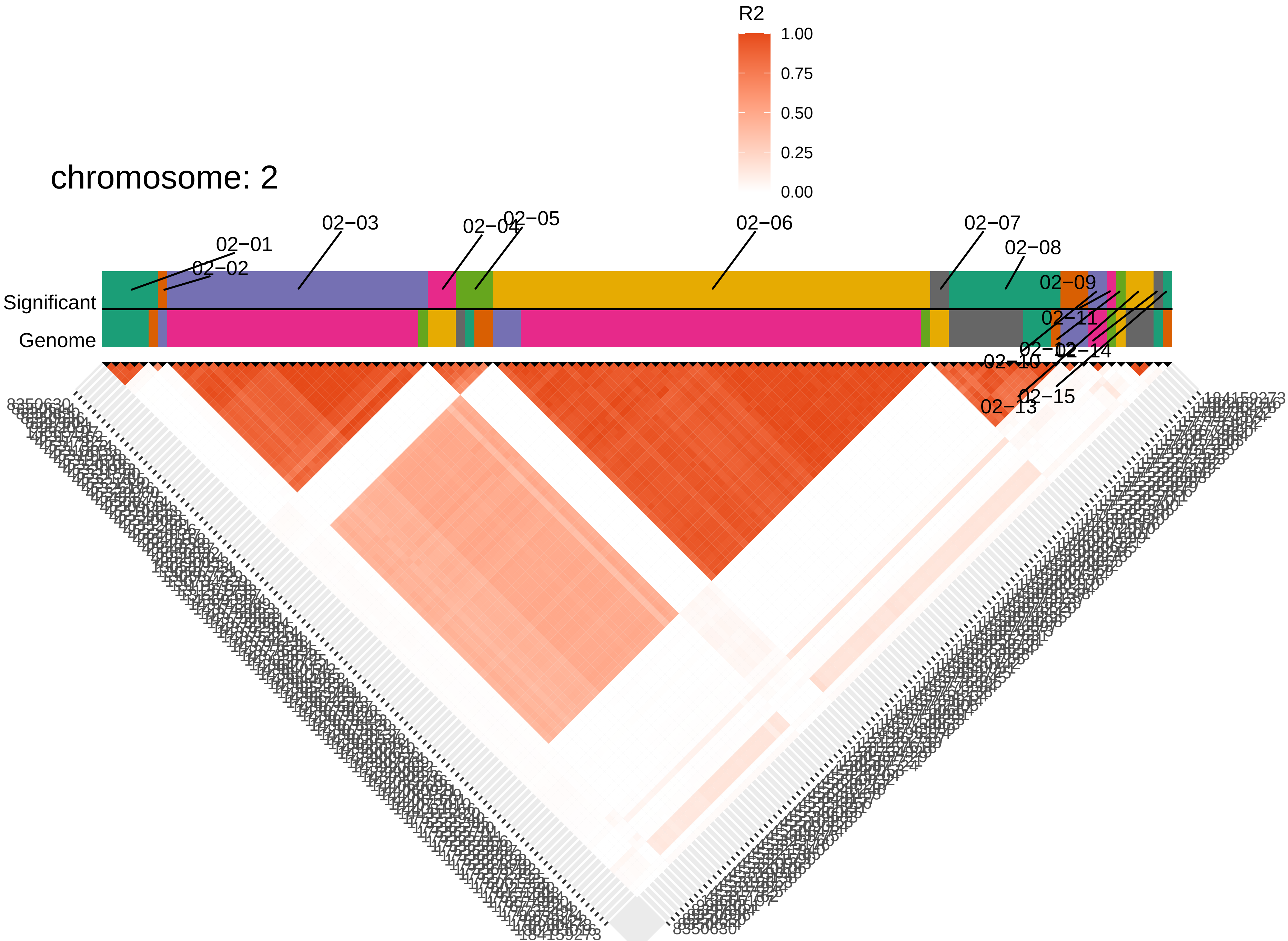
chromosome: 2
R2
1.00
0.75
0.50
0.25
0.00
02−01
02−02
02−03
02−04
02−05
02−06
02−07
02−08
02−09
02−10
02−11
02−12
02−13
02−14
02−15
Significant
Genome
83506304
184159273

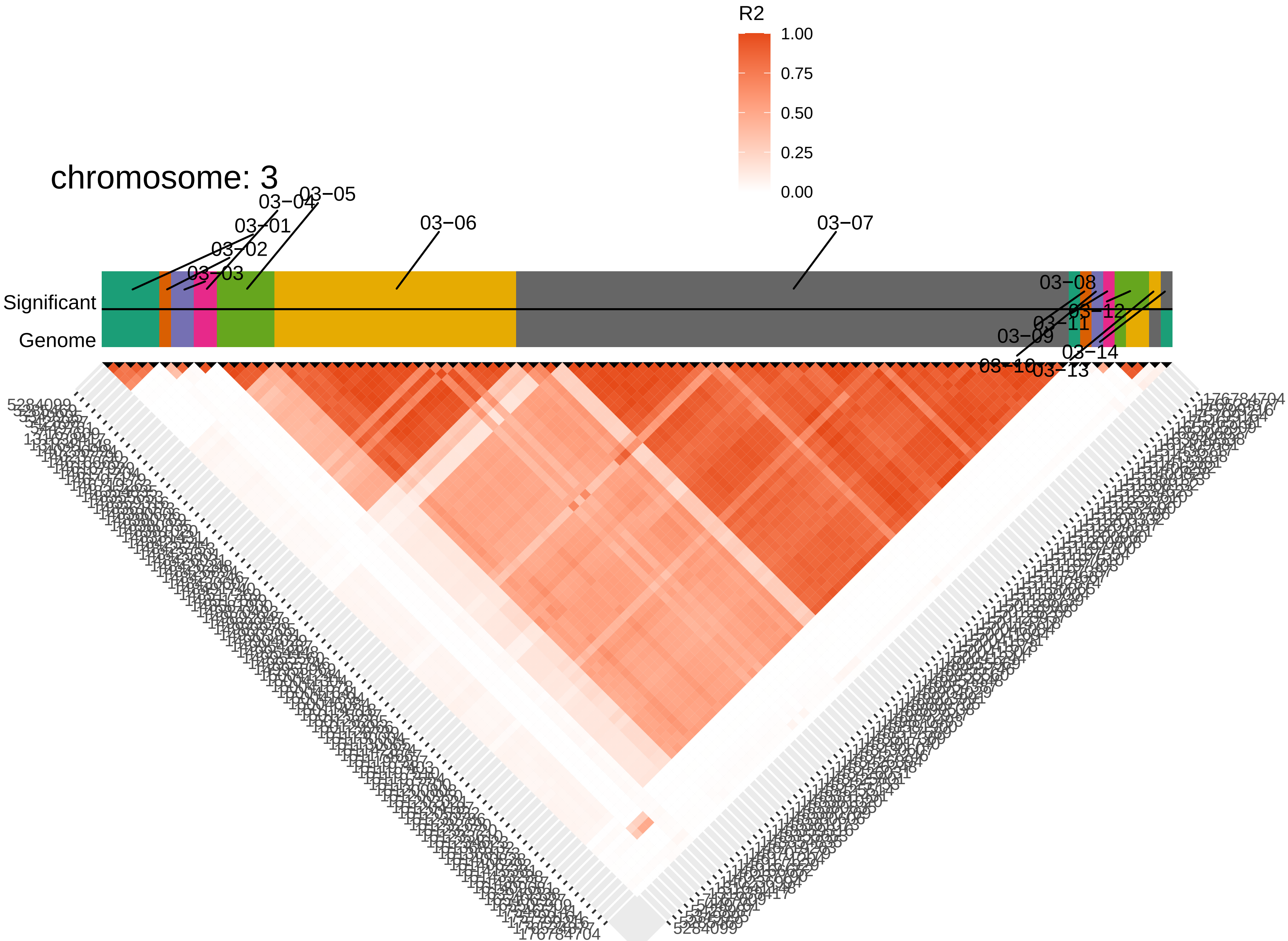

chromosome: 3
R2
1.00
0.75
0.50
0.25
0.00
03−01
03−02
03−03
03−04
03−05
03−06
03−07
03−08
03−09
03−10
03−11
03−12
03−13
03−14
Significant
Genome
5284099
176784704

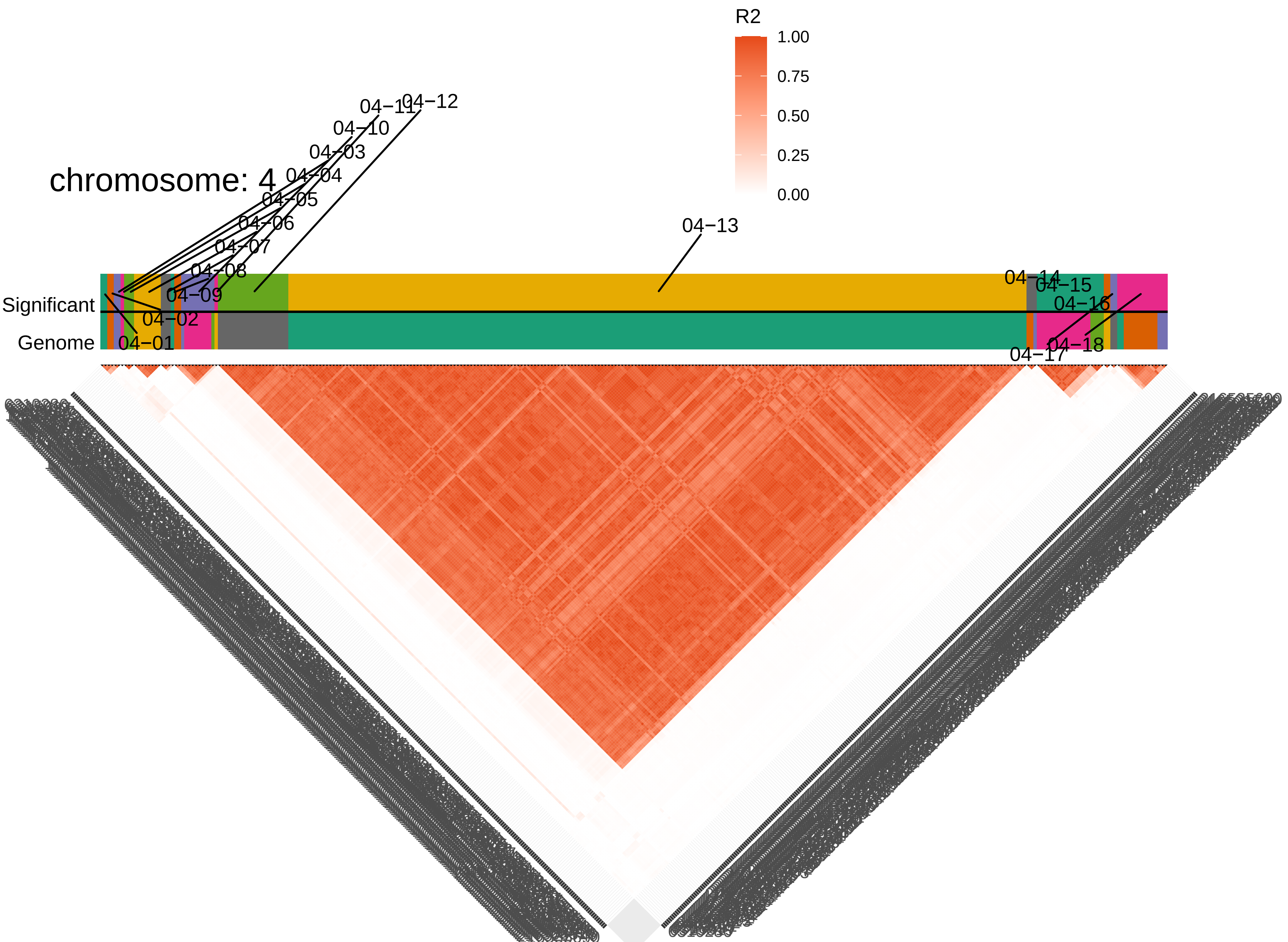
chromosome: 4
R2
1.00
0.75
0.50
0.25
0.00
Significant
Genome
04−01
04−02
04−03
04−04
04−05
04−06
04−07
04−08
04−09
04−10
04−11
04−12
04−13
04−14
04−15
04−16
04−17
04−18

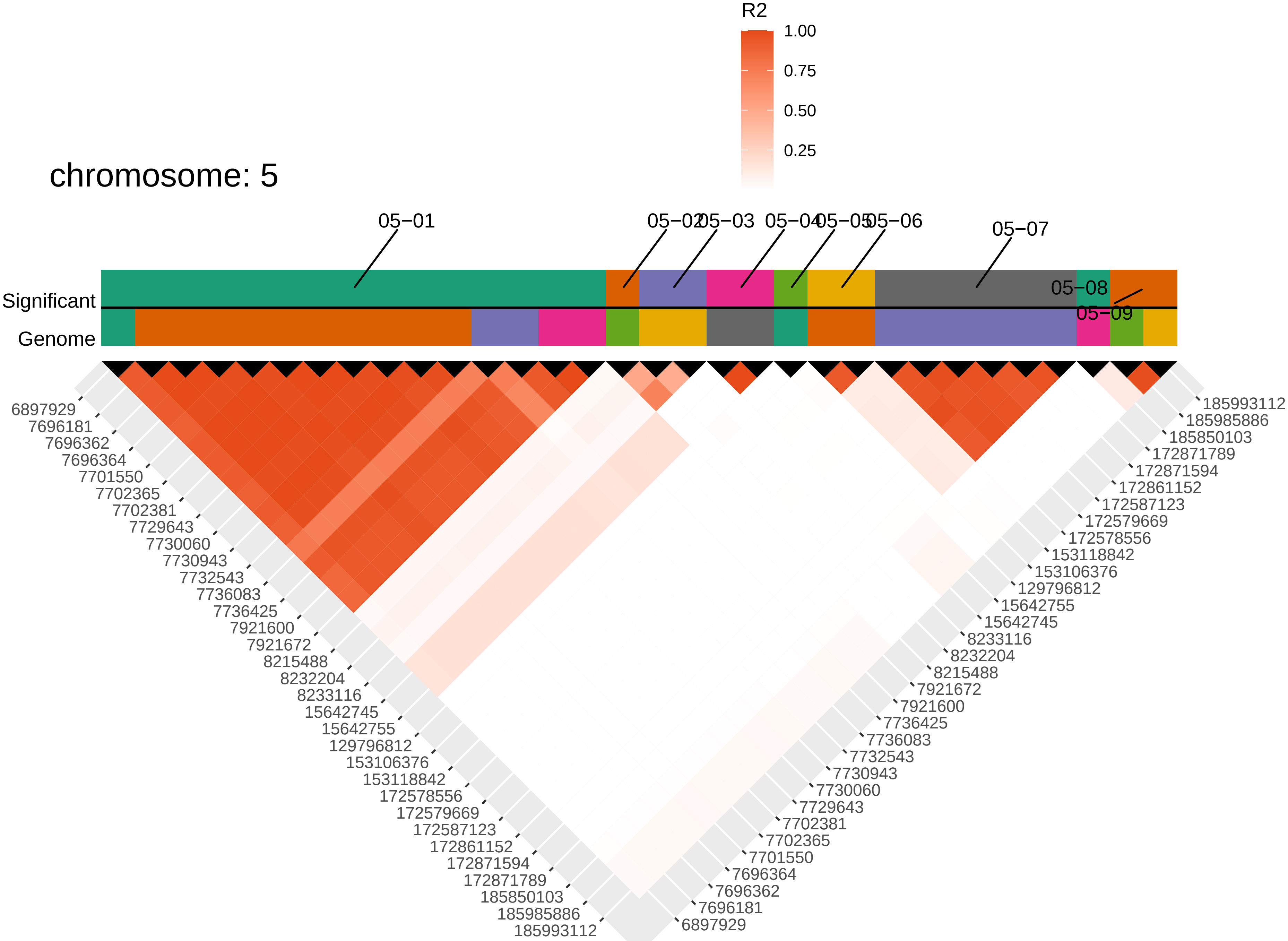

chromosome: 5
R2
1.00
0.75
0.50
0.25
05−01
05−02
05−03
05−04
05−05
05−06
05−07
05−08
05−09
Significant
Genome
6897929
7696181
7696362
7696364
7701550
7702365
7702381
7729643
7730060
7730943
7732543
7736083
7736425
7921600
7921672
8215488
8232204
8233116
15642745
15642755
129796812
153106376
153118842
172578556
172579669
172587123
172861152
172871594
172871789
185850103
185985886
185993112

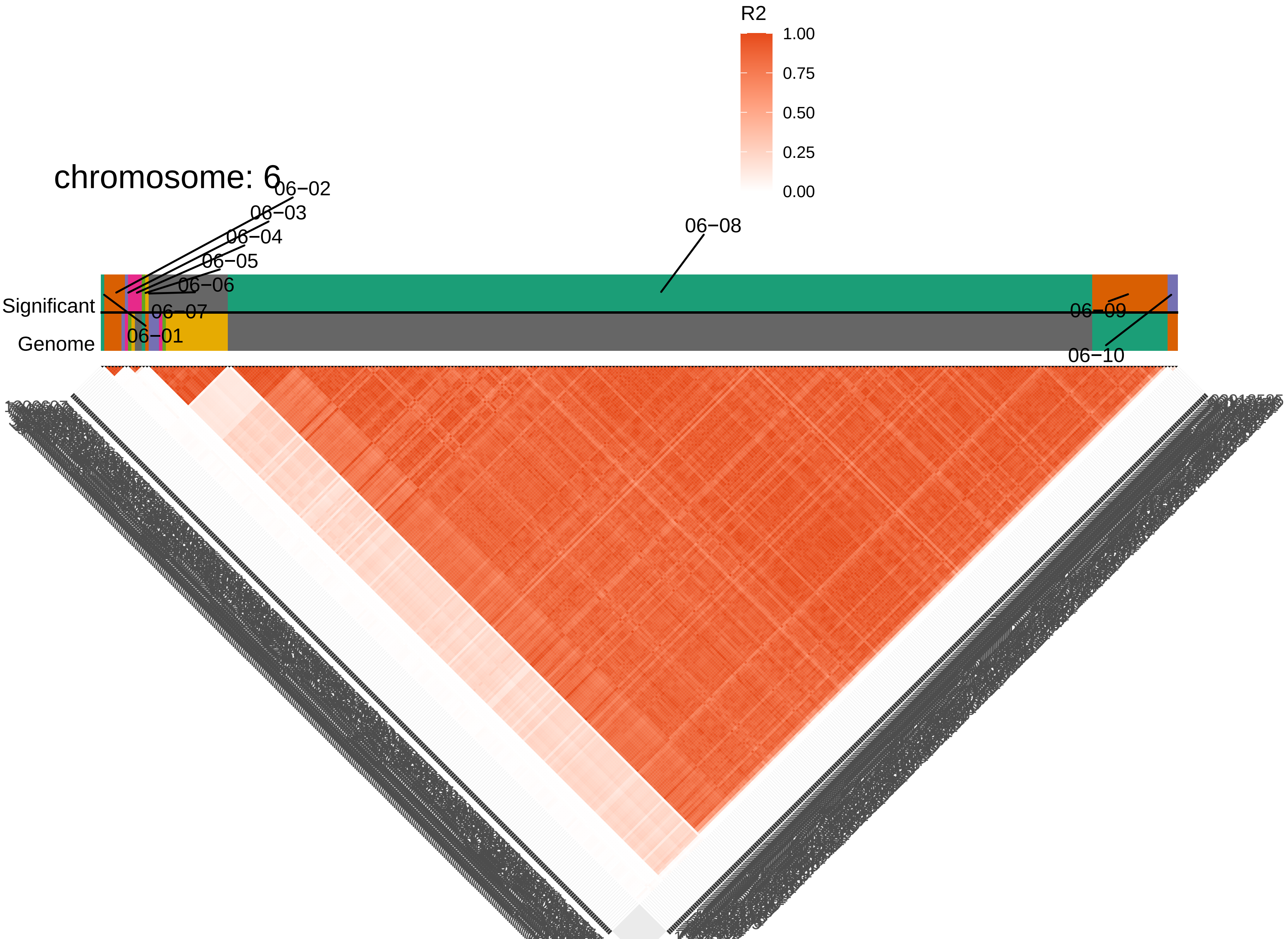

chromosome: 6
R2
1.00
0.75
0.50
0.25
0.00
Significant
Genome
06−01
06−02
06−03
06−04
06−05
06−06
06−07
06−08
06−09
06−10

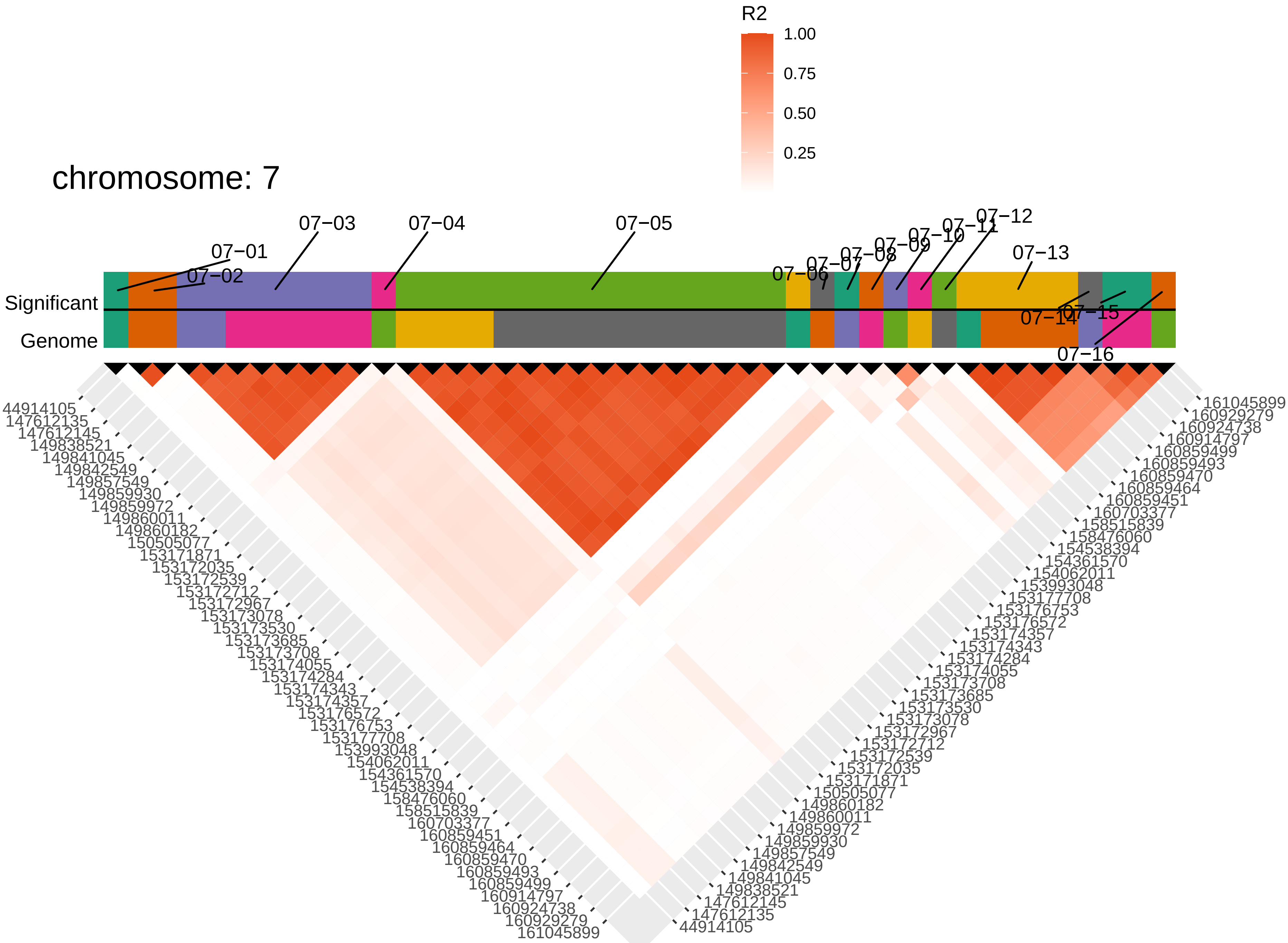
R2
1.00
0.75
0.50
0.25
chromosome: 7
07−01
07−02
07−03
07−04
07−05
07−06
07−07
07−08
07−09
07−10
07−11
07−12
07−13
07−14
07−15
07−16
Significant
Genome
44914105
147612135
147612145
149838521
149841045
149842549
149857549
149859930
149859972
149860011
149860182
150505077
153171871
153172035
153172539
153172712
153172967
153173078
153173530
153173685
153173708
153174055
153174284
153174343
153174357
153176572
153176753
153177708
153993048
154062011
154361570
154538394
158476060
158515839
160703377
160859451
160859464
160859470
160859493
160859499
160914797
160924738
160929279
161045899

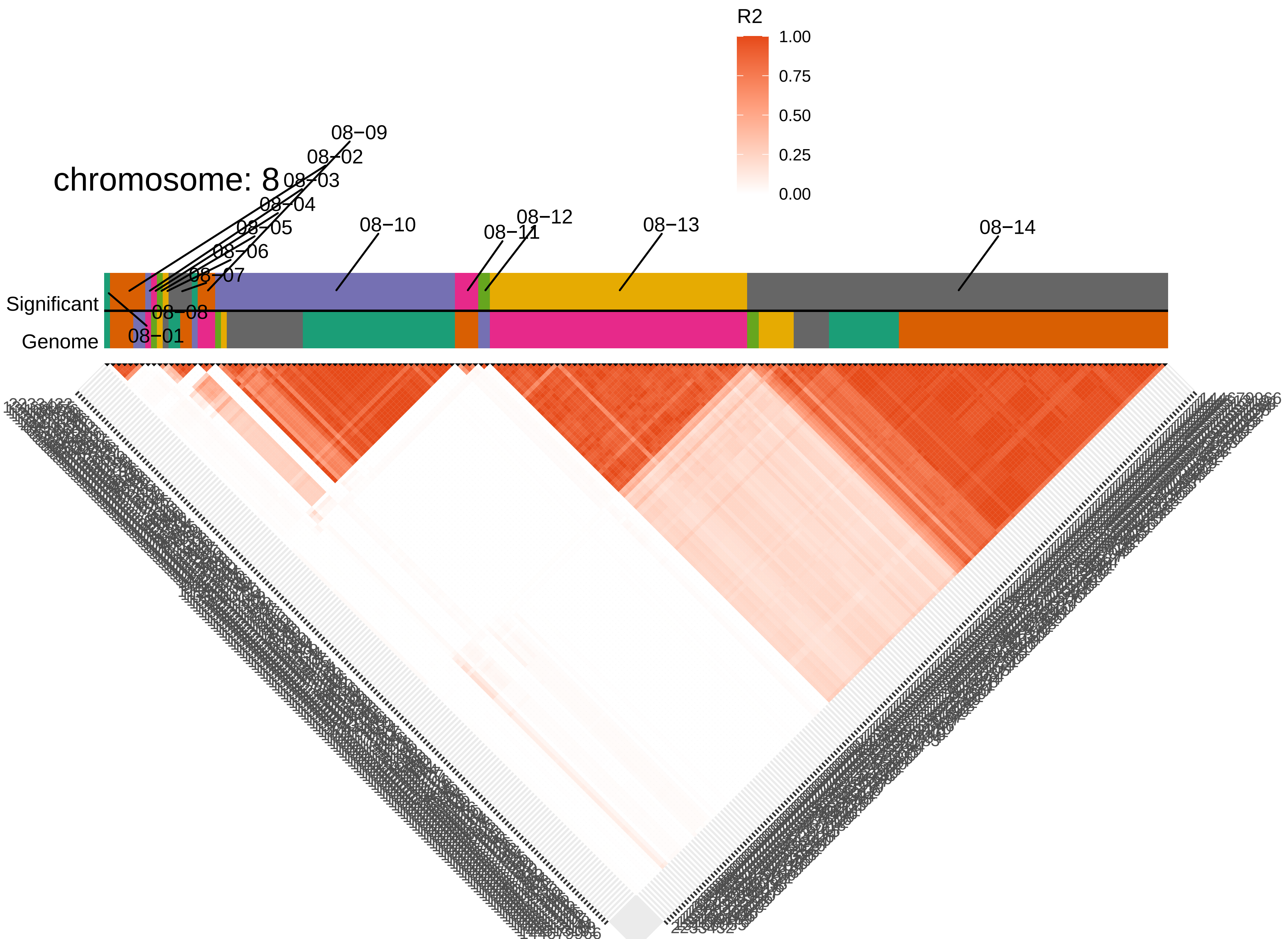

chromosome: 8
R2
1.00
0.75
0.50
0.25
0.00
08−01
08−02
08−03
08−04
08−05
08−06
08−07
08−08
08−09
08−10
08−11
08−12
08−13
08−14
Significant
Genome

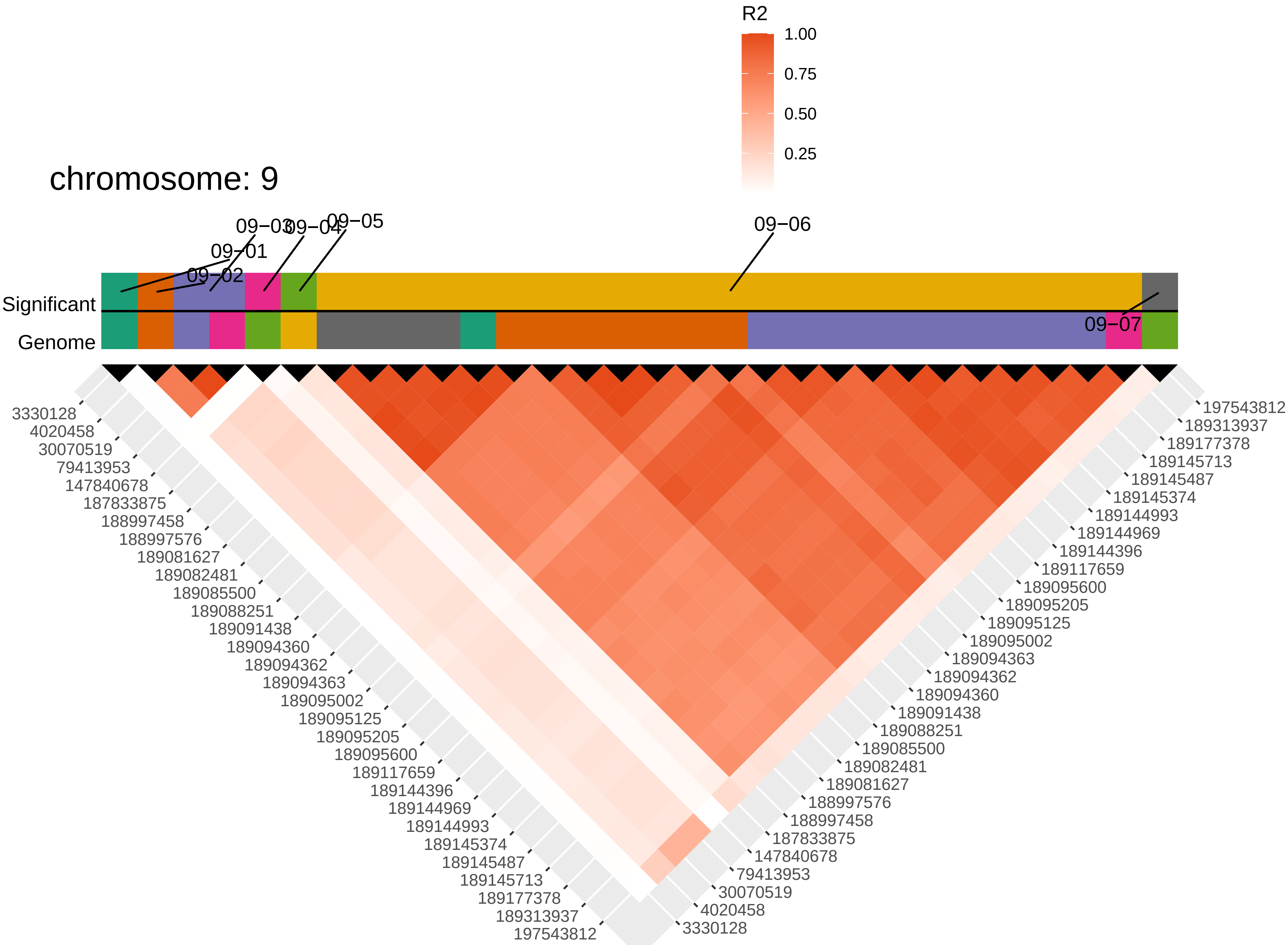

R2
1.00
0.75
0.50
0.25
chromosome: 9
09−01
09−02
09−03
09−04
09−05
09−06
09−07
Significant
Genome
3330128
4020458
30070519
79413953
147840678
187833875
188997458
188997576
189081627
189082481
189085500
189088251
189091438
189094360
189094362
189094363
189095002
189095125
189095205
189095600
189117659
189144396
189144969
189144993
189145374
189145487
189145713
189177378
189313937
197543812

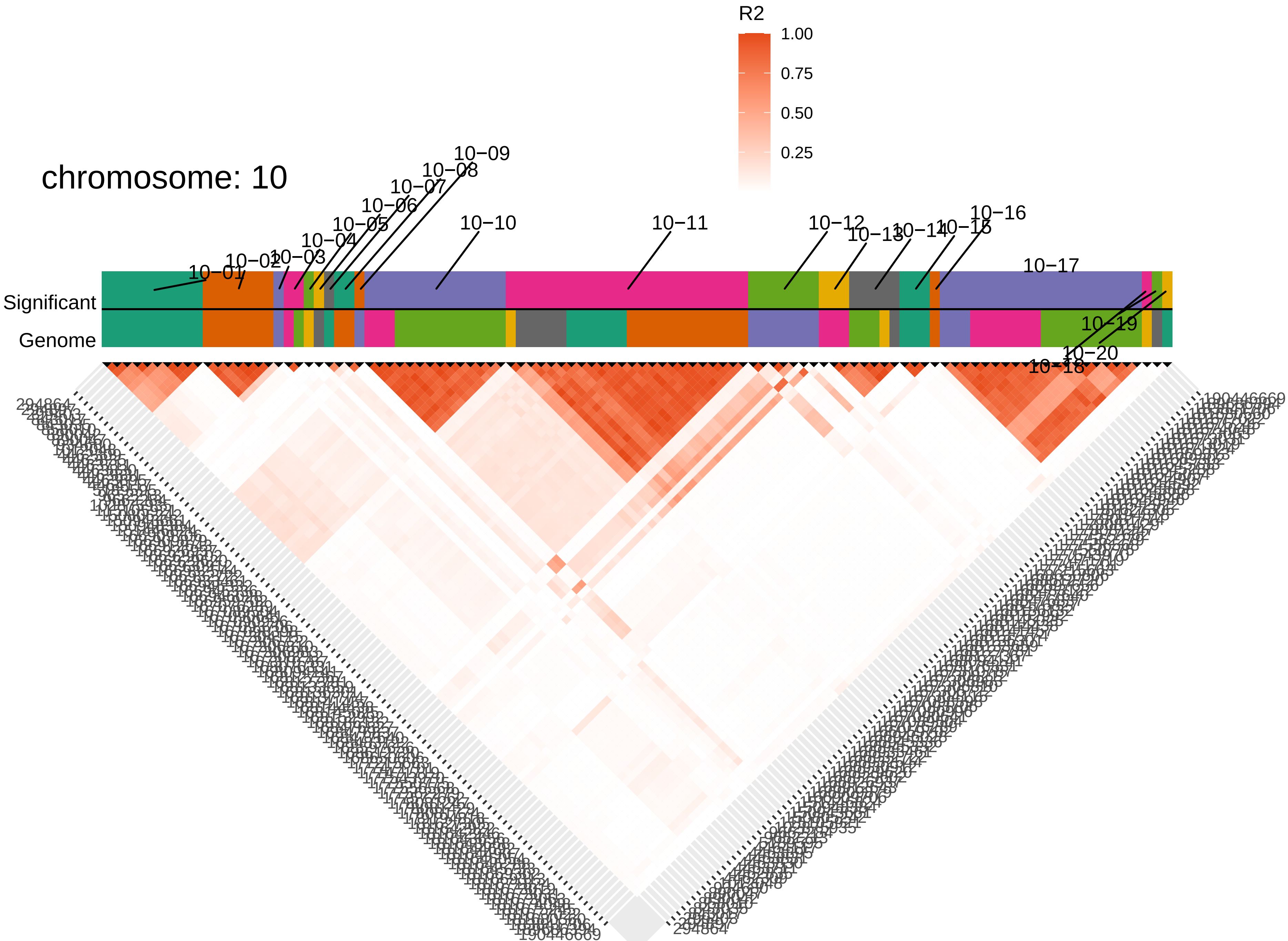
chromosome: 10
R2
1.00
0.75
0.50
0.25
Significant
Genome
10−01
10−02
10−03
10−04
10−05
10−06
10−07
10−08
10−09
10−10
10−11
10−12
10−13
10−14
10−15
10−16
10−17
10−18
10−19
10−20
294864
190446669

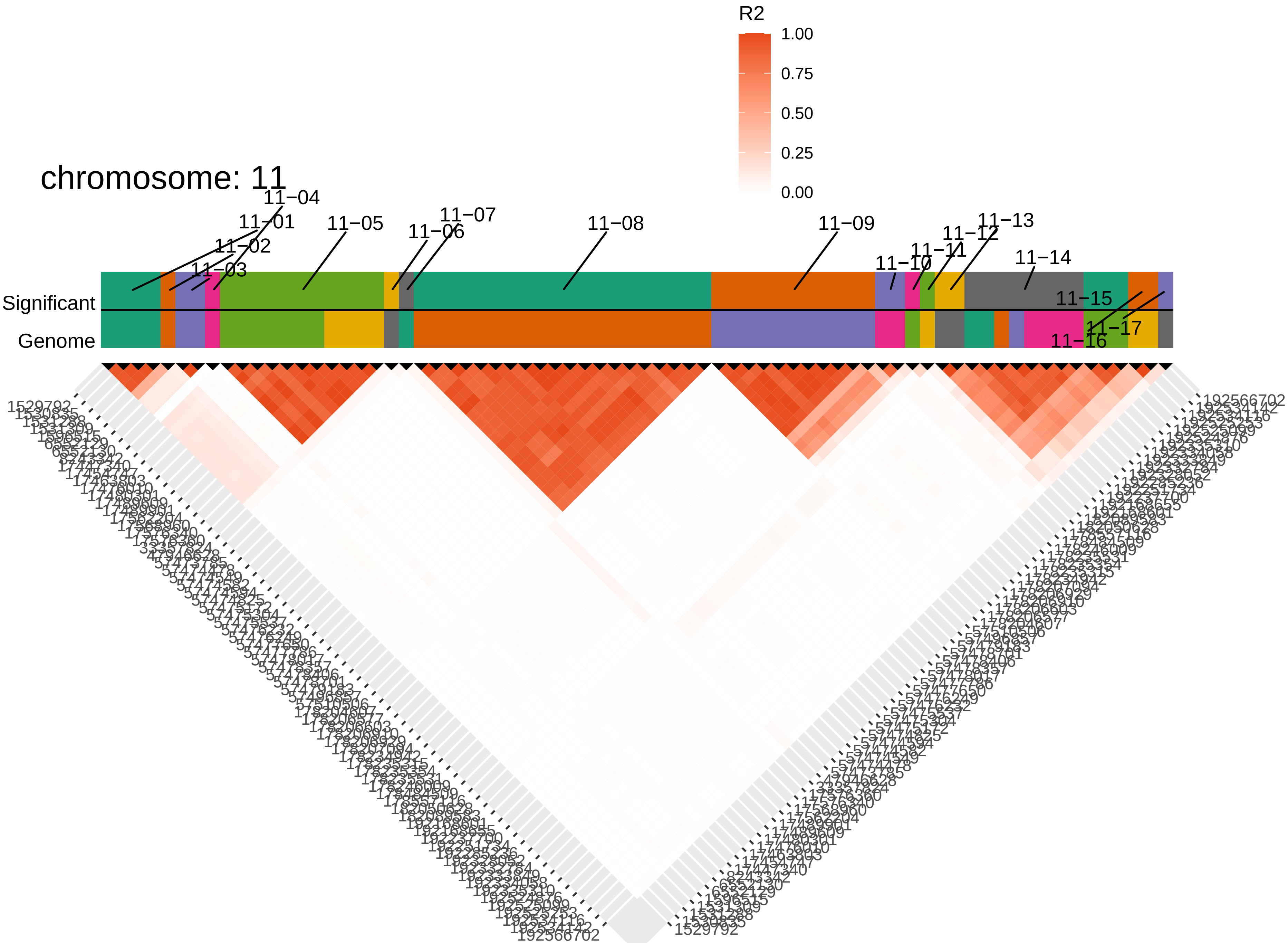

chromosome: 11
R2
1.00
0.75
0.50
0.25
0.00
Significant
Genome
11−01
11−02
11−03
11−04
11−05
11−06
11−07
11−08
11−09
11−10
11−11
11−12
11−13
11−14
11−15
11−16
11−17
1529792
192566702

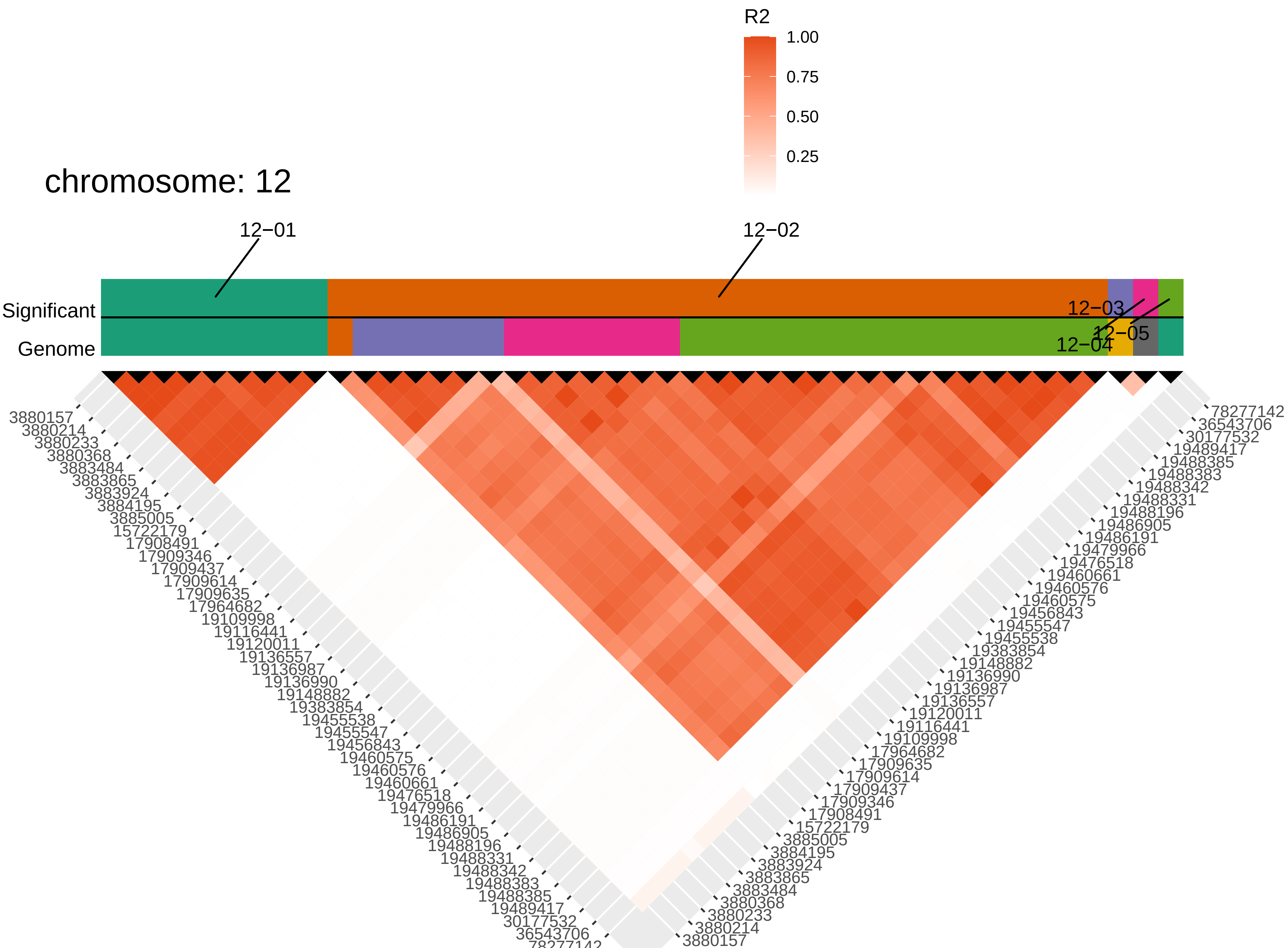

chromosome: 12
R2
1.00
0.75
0.50
0.25
12−01
12−02
12−03
12−04
12−05
Significant
Genome
3880157
3880214
3880233
3880368
3883484
3883865
3883924
3884195
3885005
15722179
17908491
17909346
17909437
17909614
17909635
17964682
19109998
19116441
19120011
19136557
19136987
19136990
19148882
19383854
19455538
19455547
19456843
19460575
19460576
19460661
19476518
19479966
19486191
19486905
19488196
19488331
19488342
19488383
19488385
19489417
30177532
36543706
78277142

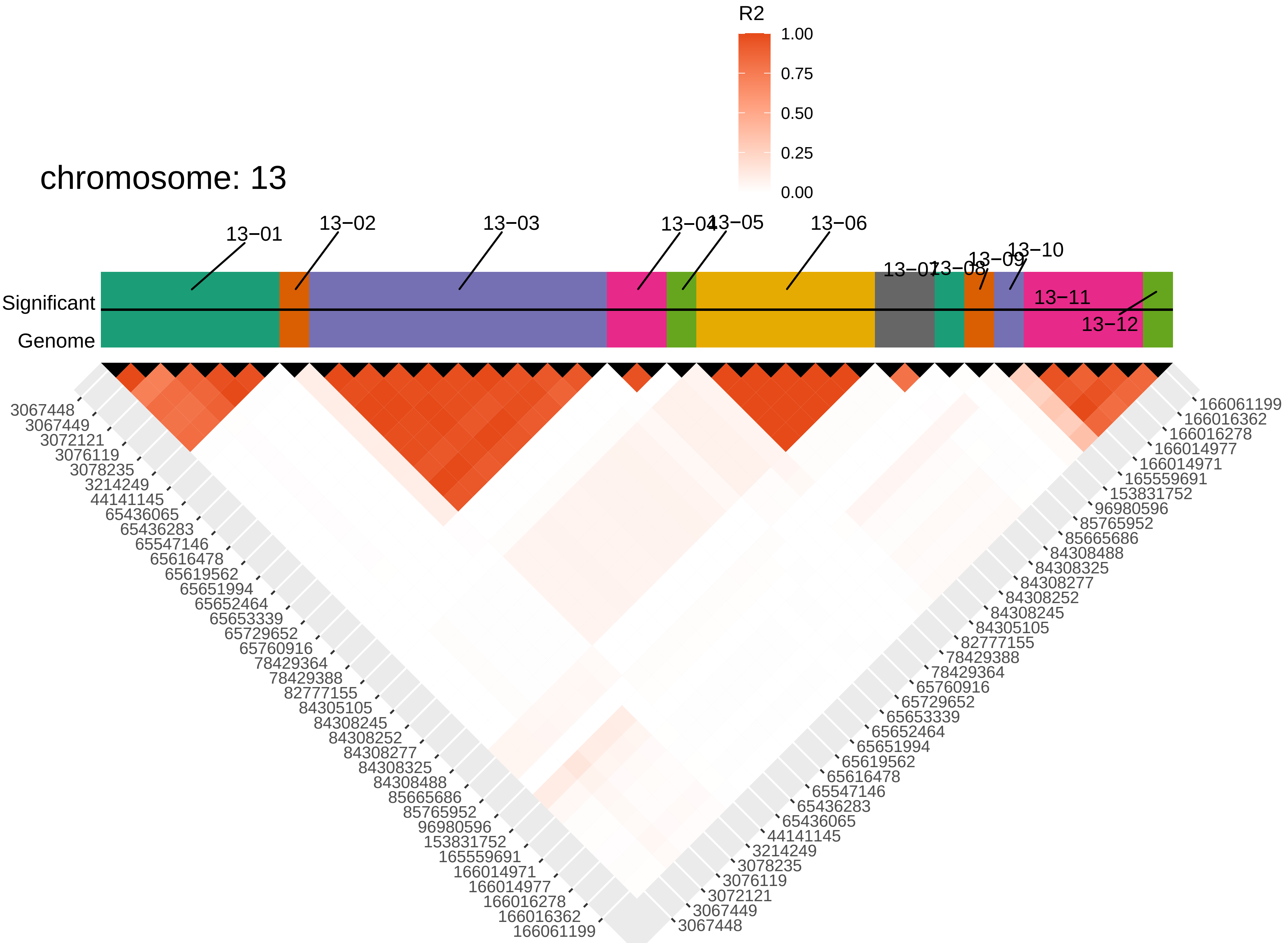

chromosome: 13
R2
1.00
0.75
0.50
0.25
0.00
13−01
13−02
13−03
13−04
13−05
13−06
13−07
13−08
13−09
13−10
13−11
13−12
Significant
Genome
3067448
3067449
3072121
3076119
3078235
3214249
44141145
65436065
65436283
65547146
65616478
65619562
65651994
65652464
65653339
65729652
65760916
78429364
78429388
82777155
84305105
84308245
84308252
84308277
84308325
84308488
85665686
85765952
96980596
153831752
165559691
166014971
166014977
166016278
166016362
166061199

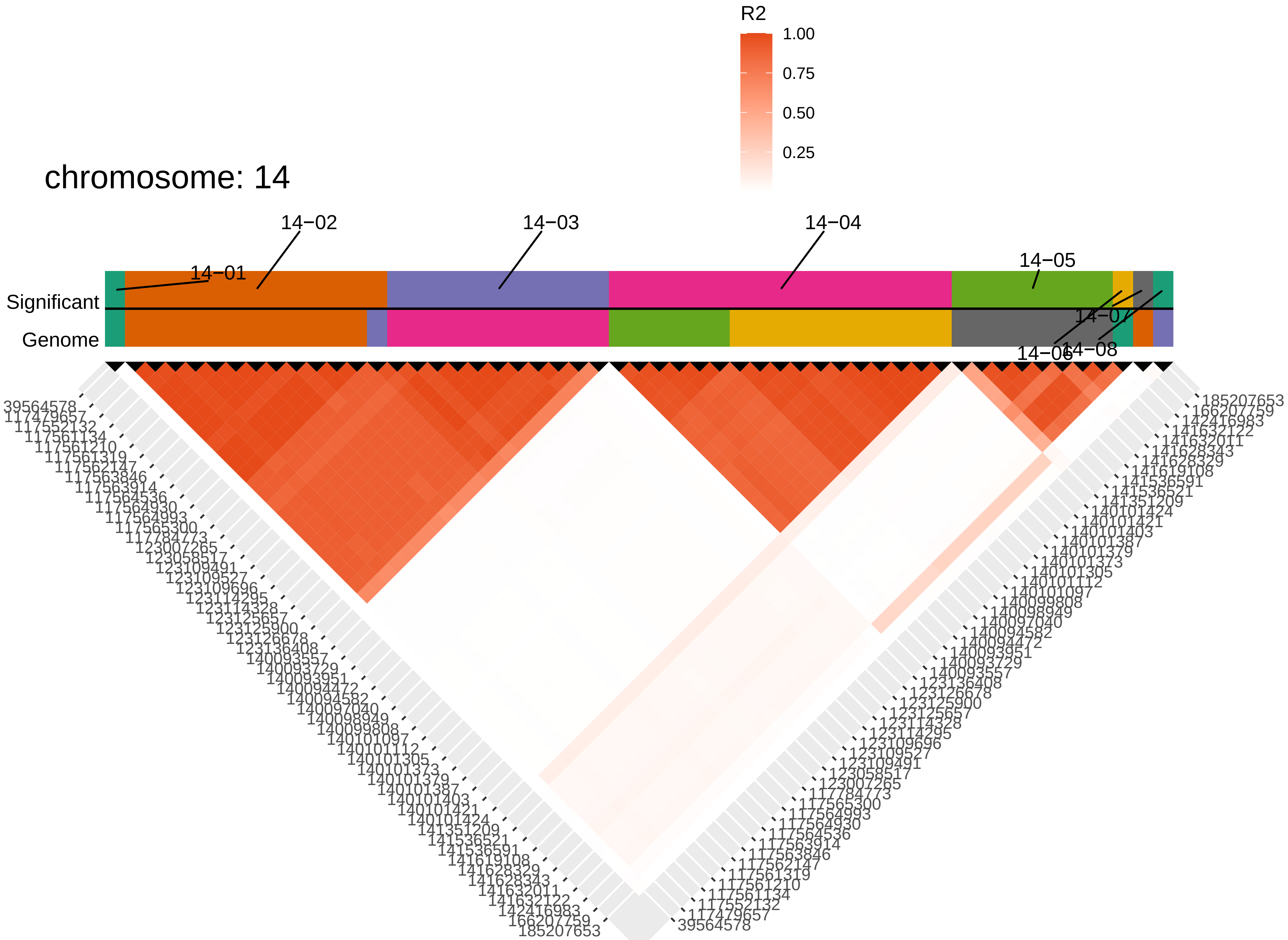

chromosome: 14
R2
1.00
0.75
0.50
0.25
14−01
14−02
14−03
14−04
14−05
14−06
14−07
14−08
Significant
Genome
39564578
117479657
117552132
117561134
117561210
117561319
117562147
117563846
117563914
117564536
117564930
117564993
117565300
117784773
123007265
123058517
123109491
123109527
123109696
123114295
123114328
123125657
123125900
123126678
123136408
140093557
140093729
140093951
140094472
140094582
140097040
140098949
140099808
140101097
140101112
140101305
140101373
140101379
140101387
140101403
140101421
140101424
141351209
141536521
141536591
141619108
141628329
141628343
141632011
141632122
142416983
166207759
185207653

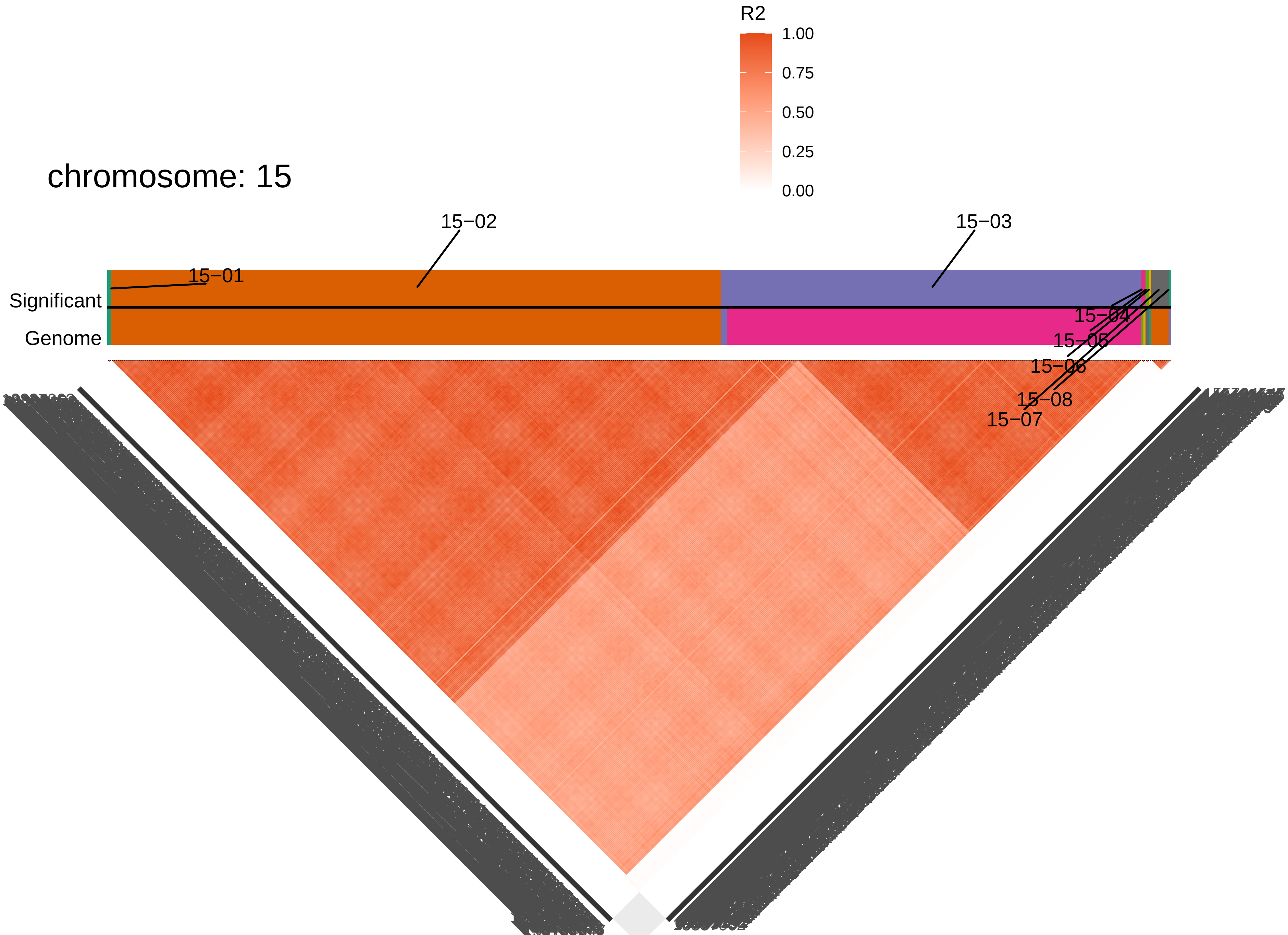
chromosome: 15
R2
1.00
0.75
0.50
0.25
0.00
15−02
15−03
15−01
Significant
Genome
15−04
15−05
15−06
15−08
15−07

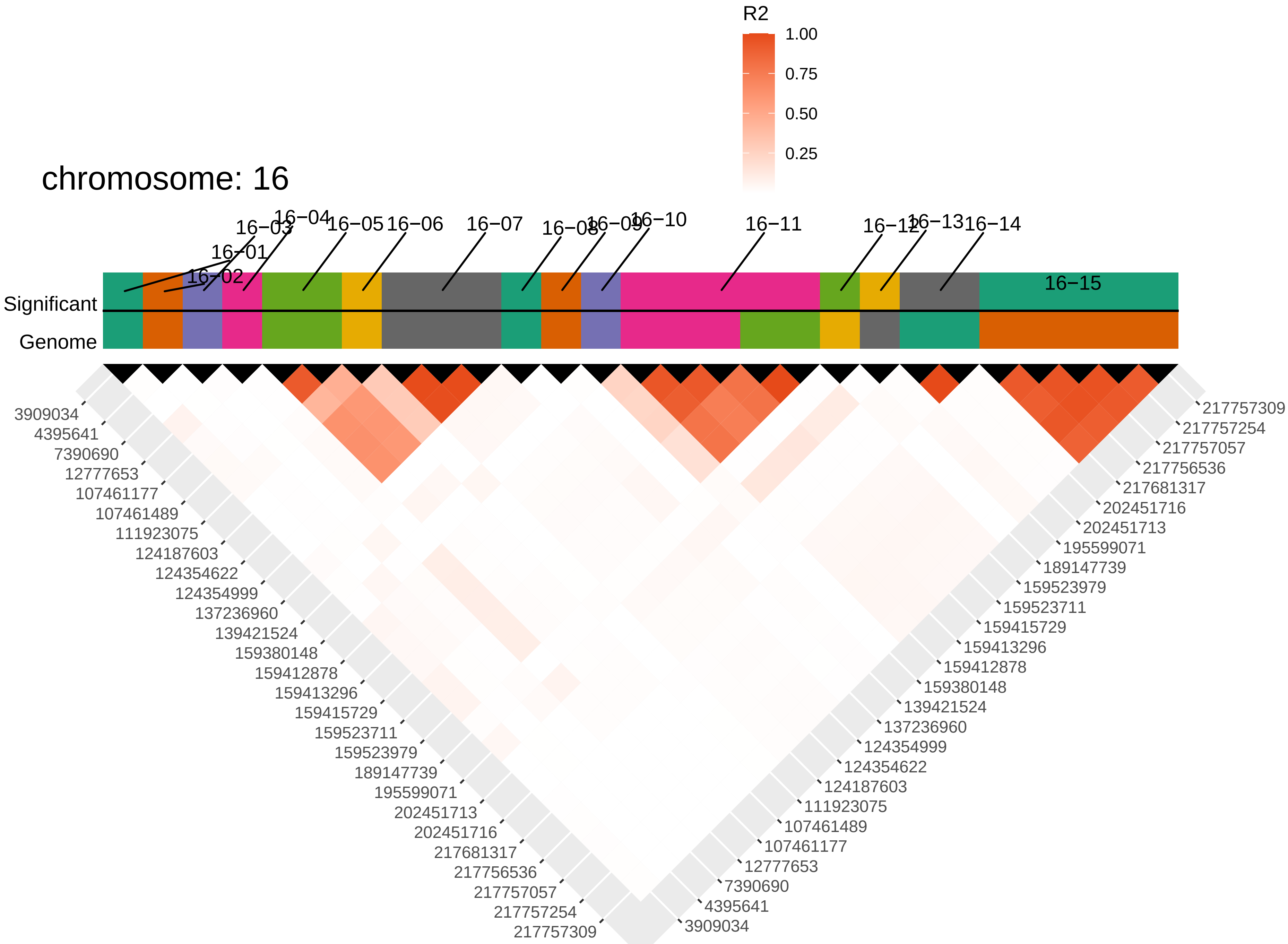

chromosome: 16
R2
1.00
0.75
0.50
0.25
16−01
16−02
16−03
16−04
16−05
16−06
16−07
16−08
16−09
16−10
16−11
16−12
16−13
16−14
16−15
Significant
Genome
3909034
4395641
7390690
12777653
107461177
107461489
111923075
124187603
124354622
124354999
137236960
139421524
159380148
159412878
159413296
159415729
159523711
159523979
189147739
195599071
202451713
202451716
217681317
217756536
217757057
217757254
217757309

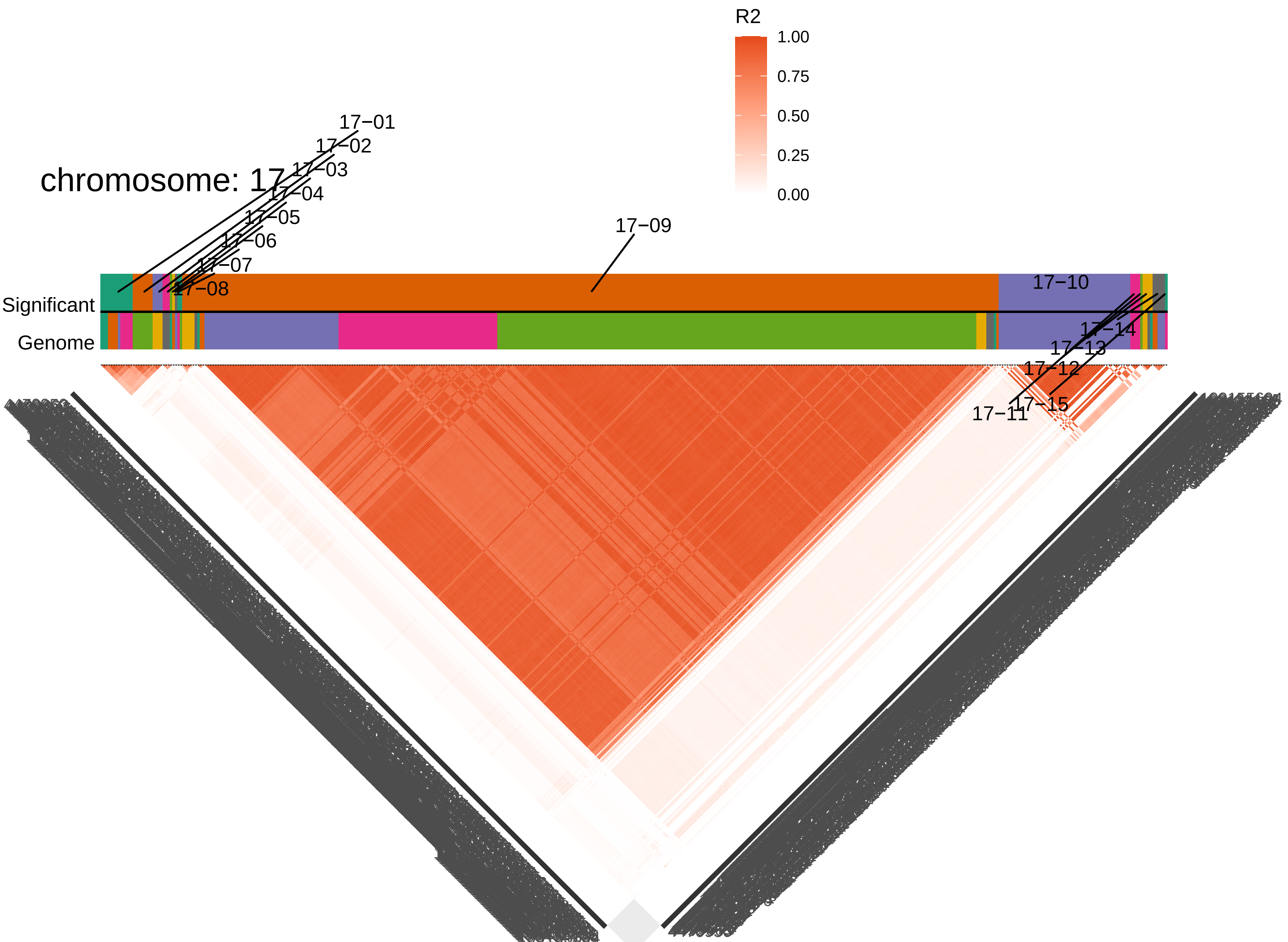

chromosome: 17
R2
1.00
0.75
0.50
0.25
0.00
17−01
17−02
17−03
17−04
17−05
17−06
17−07
17−08
17−09
17−10
17−11
17−12
17−13
17−14
17−15
Significant
Genome

Figure S6 **Trait colocalization per chromosome** Traits in (a) control , (b) low nutrient stressed, and their (c) plasticity (marked as "logdiff") are clustered based on hierarchical clustering of their bivariate (spearman) correlations. Traits grouped more closely together are more highly correlated. If a linkage block contained a significant SNP for a trait, all other traits were assessed to determine if any had a significant (green) or suggestive (i.e., top 0.1% of all SNPs; gray) in the same block . Plus/minus symbols in the tiles indicate the sign of $\beta$ (effect of minor allele on trait). Linkage blocks are ordered by chromosome with black lines separating them. Sizes of the linkage blocks vary both in physical size as well as gene number.

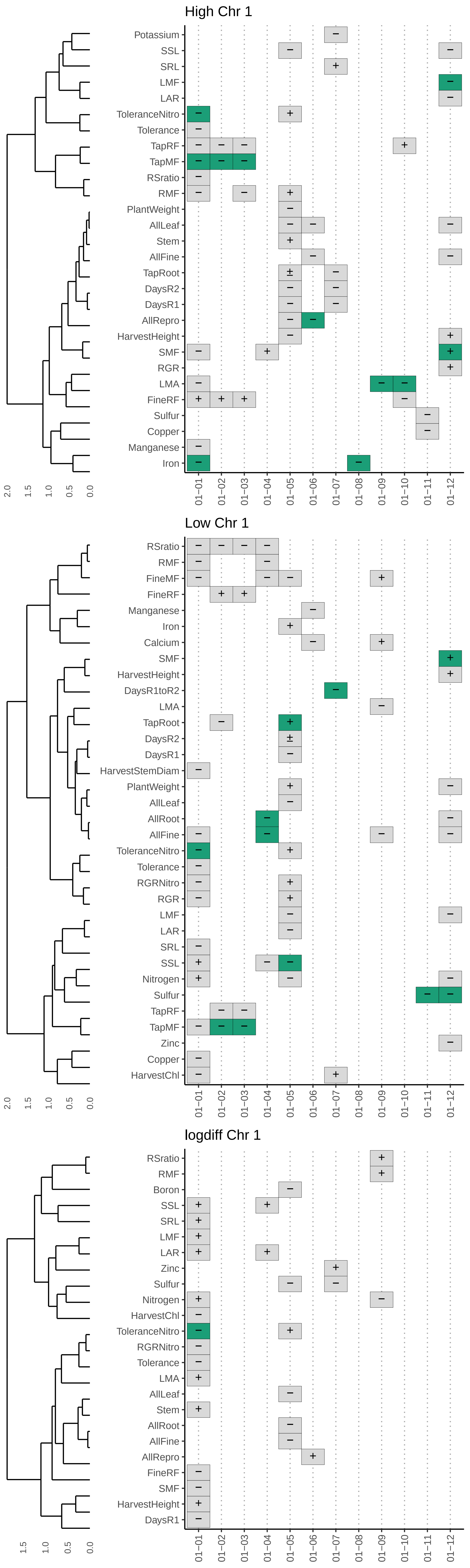
High Chr 1
Potassium
SSL
SRL
LMF
LAR
ToleranceNitro
Tolerance
TapRF
TapMF
RSratio
RMF
PlantWeight
AllLeaf
Stem
AllFine
TapRoot
DaysR2
DaysR1
AllRepro
HarvestHeight
SMF
RGR
LMA
FineRF
Sulfur
Copper
Manganese
Iron
2.0 1.5 1.0 0.5 0.0
01−01 01−02 01−03 01−04 01−05 01−06 01−07 01−08 01−09 01−10 01−11 01−12
Low Chr 1
RSratio
RMF
FineMF
FineRF
Manganese
Iron
Calcium
SMF
HarvestHeight
DaysR1toR2
LMA
TapRoot
DaysR2
DaysR1
HarvestStemDiam
PlantWeight
AllLeaf
AllRoot
AllFine
ToleranceNitro
Tolerance
RGRNitro
RGR
LMF
LAR
SRL
SSL
Nitrogen
Sulfur
TapRF
TapMF
Zinc
Copper
HarvestChl
2.0 1.5 1.0 0.5 0.0
01−01 01−02 01−03 01−04 01−05 01−06 01−07 01−08 01−09 01−10 01−11 01−12
logdiff Chr 1
RSratio
RMF
Boron
SSL
SRL
LMF
LAR
Zinc
Sulfur
Nitrogen
HarvestChl
ToleranceNitro
RGRNitro
Tolerance
LMA
AllLeaf
Stem
AllRoot
AllFine
AllRepro
FineRF
SMF
HarvestHeight
DaysR1
1.5 1.0 0.5 0.0
01−01 01−02 01−03 01−04 01−05 01−06 01−07 01−08 01−09 01−10 01−11 01−12

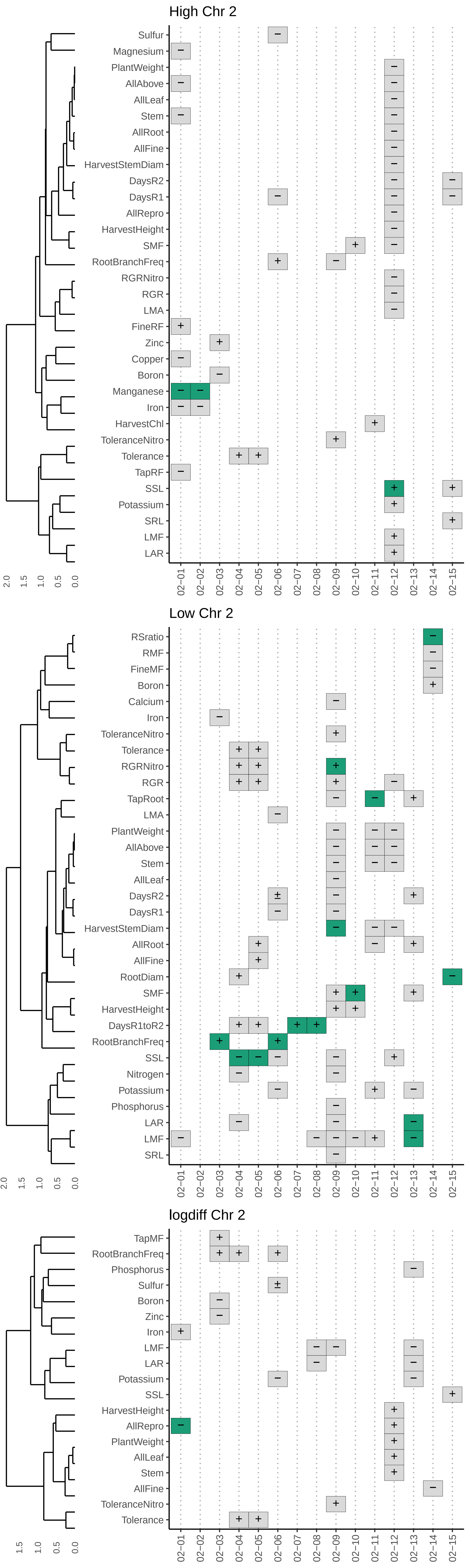

High Chr 2
Sulfur
Magnesium
PlantWeight
AllAbove
AllLeaf
Stem
AllRoot
AllFine
HarvestStemDiam
DaysR2
DaysR1
AllRepro
HarvestHeight
SMF
RootBranchFreq
RGRNitro
RGR
LMA
FineRF
Zinc
Copper
Boron
Manganese
Iron
HarvestChl
ToleranceNitro
Tolerance
TapRF
SSL
Potassium
SRL
LMF
LAR
2.0 1.5 1.0 0.5 0.0
02−01 02−02 02−03 02−04 02−05 02−06 02−07 02−08 02−09 02−10 02−11 02−12 02−13 02−14 02−15
Low Chr 2
RSratio
RMF
FineMF
Boron
Calcium
Iron
ToleranceNitro
Tolerance
RGRNitro
RGR
TapRoot
LMA
PlantWeight
AllAbove
Stem
AllLeaf
DaysR2
DaysR1
HarvestStemDiam
AllRoot
AllFine
RootDiam
SMF
HarvestHeight
DaysR1toR2
RootBranchFreq
SSL
Nitrogen
Potassium
Phosphorus
LAR
LMF
SRL
2.0 1.5 1.0 0.5 0.0
02−01 02−02 02−03 02−04 02−05 02−06 02−07 02−08 02−09 02−10 02−11 02−12 02−13 02−14 02−15
logdiff Chr 2
TapMF
RootBranchFreq
Phosphorus
Sulfur
Boron
Zinc
Iron
LMF
LAR
Potassium
SSL
HarvestHeight
AllRepro
PlantWeight
AllLeaf
Stem
AllFine
ToleranceNitro
Tolerance
1.5 1.0 0.5 0.0
02−01 02−02 02−03 02−04 02−05 02−06 02−07 02−08 02−09 02−10 02−11 02−12 02−13 02−14 02−15

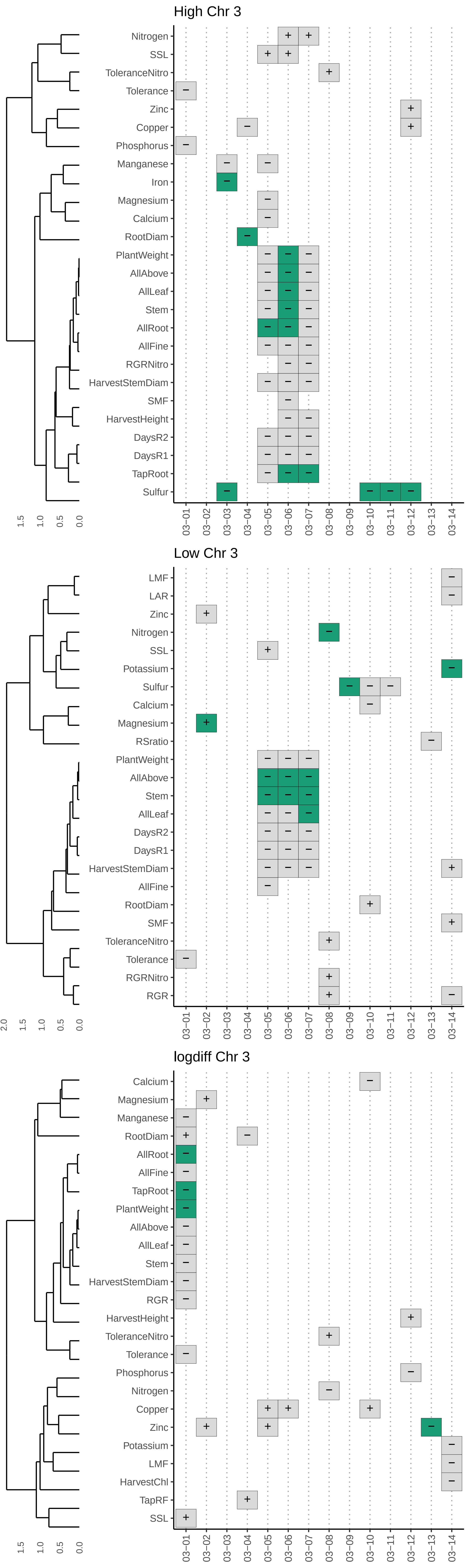
High Chr 3
Nitrogen
SSL
ToleranceNitro
Tolerance
Zinc
Copper
Phosphorus
Manganese
Iron
Magnesium
Calcium
RootDiam
PlantWeight
AllAbove
AllLeaf
Stem
AllRoot
AllFine
RGRNitro
HarvestStemDiam
SMF
HarvestHeight
DaysR2
DaysR1
TapRoot
Sulfur
1.5 1.0 0.5 0.0
03−01 03−02 03−03 03−04 03−05 03−06 03−07 03−08 03−09 03−10 03−11 03−12 03−13 03−14
Low Chr 3
LMF
LAR
Zinc
Nitrogen
SSL
Potassium
Sulfur
Calcium
Magnesium
RSratio
PlantWeight
AllAbove
Stem
AllLeaf
DaysR2
DaysR1
HarvestStemDiam
AllFine
RootDiam
SMF
ToleranceNitro
Tolerance
RGRNitro
RGR
2.0 1.5 1.0 0.5 0.0
03−01 03−02 03−03 03−04 03−05 03−06 03−07 03−08 03−09 03−10 03−11 03−12 03−13 03−14
logdiff Chr 3
Calcium
Magnesium
Manganese
RootDiam
AllRoot
AllFine
TapRoot
PlantWeight
AllAbove
AllLeaf
Stem
HarvestStemDiam
RGR
HarvestHeight
ToleranceNitro
Tolerance
Phosphorus
Nitrogen
Copper
Zinc
Potassium
LMF
HarvestChl
TapRF
SSL
1.5 1.0 0.5 0.0
03−01 03−02 03−03 03−04 03−05 03−06 03−07 03−08 03−09 03−10 03−11 03−12 03−13 03−14

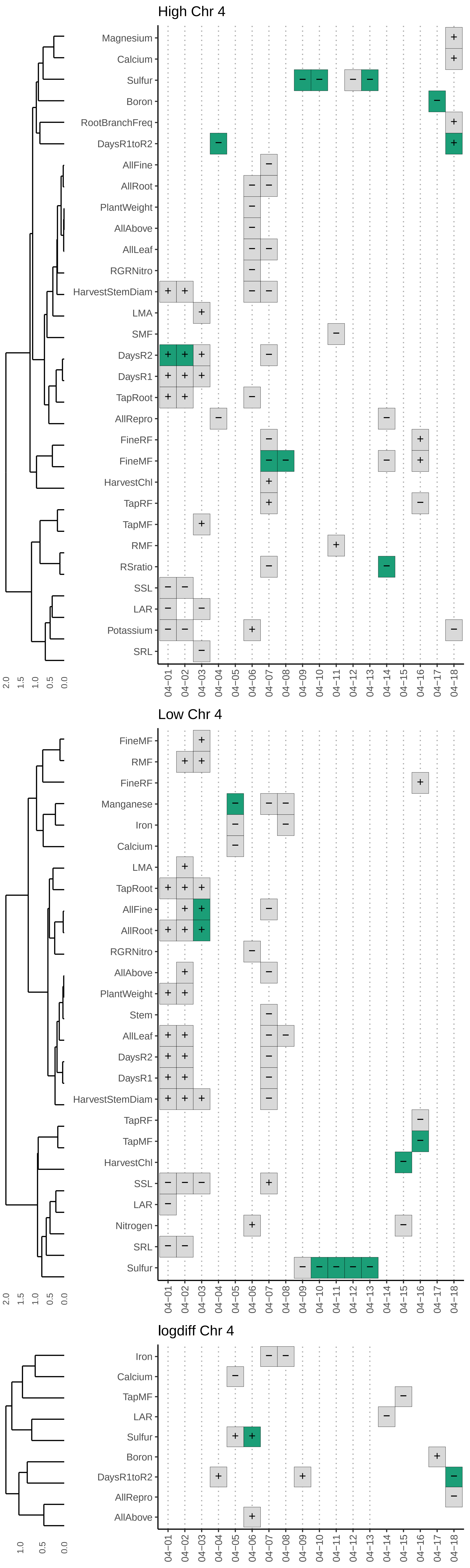
High Chr 4
Magnesium
Calcium
Sulfur
Boron
RootBranchFreq
DaysR1toR2
AllFine
AllRoot
PlantWeight
AllAbove
AllLeaf
RGRNitro
HarvestStemDiam
LMA
SMF
DaysR2
DaysR1
TapRoot
AllRepro
FineRF
FineMF
HarvestChl
TapRF
TapMF
RMF
RSratio
SSL
LAR
Potassium
SRL
2.0 1.5 1.0 0.5 0.0
04-01 04-02 04-03 04-04 04-05 04-06 04-07 04-08 04-09 04-10 04-11 04-12 04-13 04-14 04-15 04-16 04-17 04-18
Low Chr 4
FineMF
RMF
FineRF
Manganese
Iron
Calcium
LMA
TapRoot
AllFine
AllRoot
RGRNitro
AllAbove
PlantWeight
Stem
AllLeaf
DaysR2
DaysR1
HarvestStemDiam
TapRF
TapMF
HarvestChl
SSL
LAR
Nitrogen
SRL
Sulfur
2.0 1.5 1.0 0.5 0.0
04-01 04-02 04-03 04-04 04-05 04-06 04-07 04-08 04-09 04-10 04-11 04-12 04-13 04-14 04-15 04-16 04-17 04-18
logdiff Chr 4
Iron
Calcium
TapMF
LAR
Sulfur
Boron
DaysR1toR2
AllRepro
AllAbove
1.0 0.5 0.0
04-01 04-02 04-03 04-04 04-05 04-06 04-07 04-08 04-09 04-10 04-11 04-12 04-13 04-14 04-15 04-16 04-17 04-18

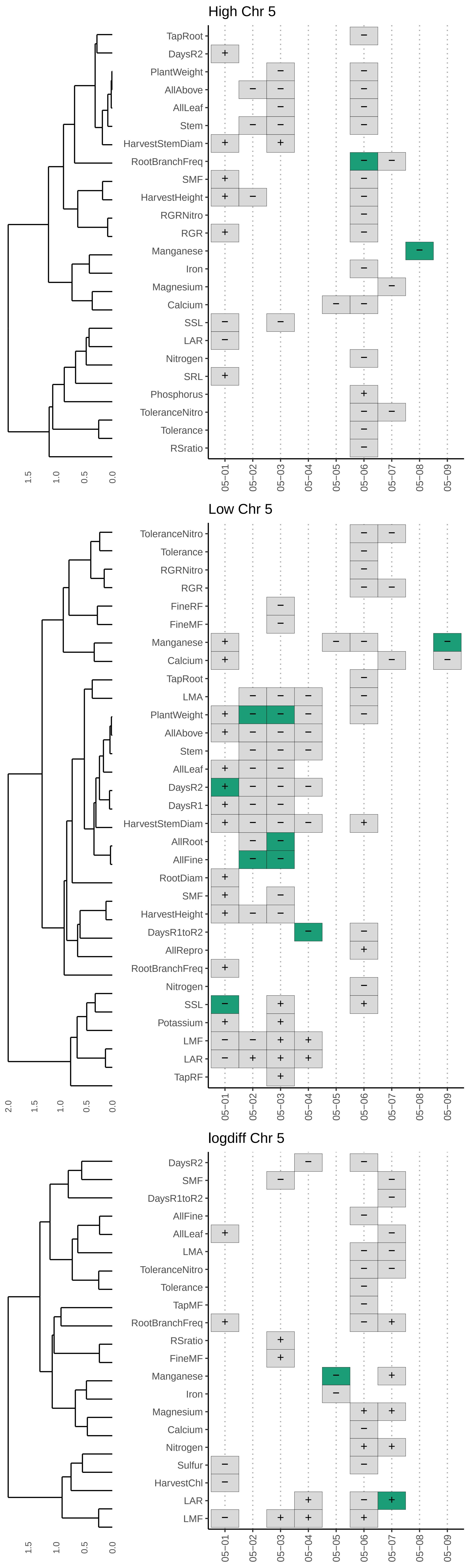
High Chr 5
TapRoot
DaysR2
PlantWeight
AllAbove
AllLeaf
Stem
HarvestStemDiam
RootBranchFreq
SMF
HarvestHeight
RGRNitro
RGR
Manganese
Iron
Magnesium
Calcium
SSL
LAR
Nitrogen
SRL
Phosphorus
ToleranceNitro
Tolerance
RSratio
1.5
1.0
0.5
0.0
05−01
05−02
05−03
05−04
05−05
05−06
05−07
05−08
05−09
Low Chr 5
ToleranceNitro
Tolerance
RGRNitro
RGR
FineRF
FineMF
Manganese
Calcium
TapRoot
LMA
PlantWeight
AllAbove
Stem
AllLeaf
DaysR2
DaysR1
HarvestStemDiam
AllRoot
AllFine
RootDiam
SMF
HarvestHeight
DaysR1toR2
AllRepro
RootBranchFreq
Nitrogen
SSL
Potassium
LMF
LAR
TapRF
2.0
1.5
1.0
0.5
0.0
05−01
05−02
05−03
05−04
05−05
05−06
05−07
05−08
05−09
logdiff Chr 5
DaysR2
SMF
DaysR1toR2
AllFine
AllLeaf
LMA
ToleranceNitro
Tolerance
TapMF
RootBranchFreq
RSratio
FineMF
Manganese
Iron
Magnesium
Calcium
Nitrogen
Sulfur
HarvestChl
LAR
LMF
1.5
1.0
0.5
0.0
05−01
05−02
05−03
05−04
05−05
05−06
05−07
05−08
05−09

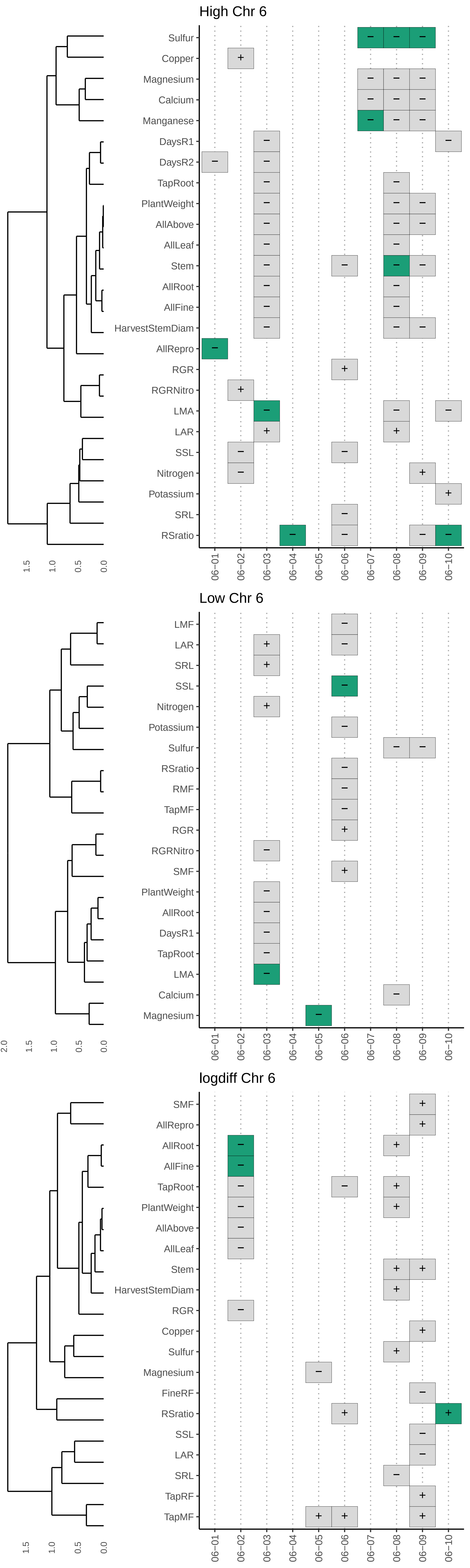
High Chr 6
Sulfur
Copper
Magnesium
Calcium
Manganese
DaysR1
DaysR2
TapRoot
PlantWeight
AllAbove
AllLeaf
Stem
AllRoot
AllFine
HarvestStemDiam
AllRepro
RGR
RGRNitro
LMA
LAR
SSL
Nitrogen
Potassium
SRL
RSratio
1.5
1.0
0.5
0.0
06−01
06−02
06−03
06−04
06−05
06−06
06−07
06−08
06−09
06−10
Low Chr 6
LMF
LAR
SRL
SSL
Nitrogen
Potassium
Sulfur
RSratio
RMF
TapMF
RGR
RGRNitro
SMF
PlantWeight
AllRoot
DaysR1
TapRoot
LMA
Calcium
Magnesium
2.0
1.5
1.0
0.5
0.0
06−01
06−02
06−03
06−04
06−05
06−06
06−07
06−08
06−09
06−10
logdiff Chr 6
SMF
AllRepro
AllRoot
AllFine
TapRoot
PlantWeight
AllAbove
AllLeaf
Stem
HarvestStemDiam
RGR
Copper
Sulfur
Magnesium
FineRF
RSratio
SSL
LAR
SRL
TapRF
TapMF
1.5
1.0
0.5
0.0
06−01
06−02
06−03
06−04
06−05
06−06
06−07
06−08
06−09
06−10

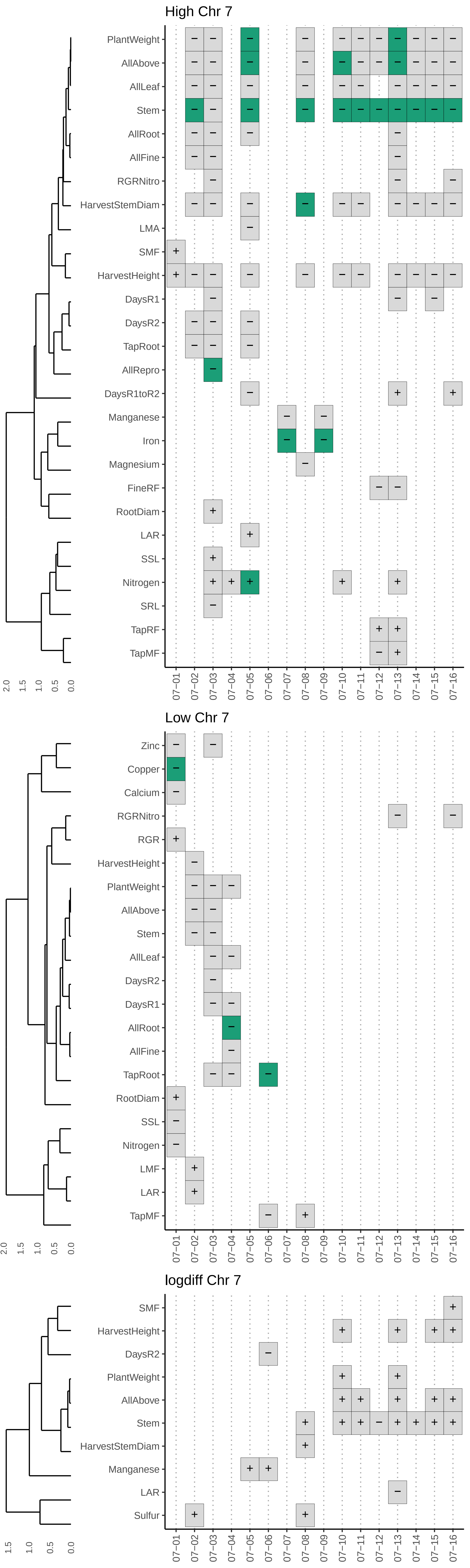

High Chr 7
PlantWeight
AllAbove
AllLeaf
Stem
AllRoot
AllFine
RGRNitro
HarvestStemDiam
LMA
SMF
HarvestHeight
DaysR1
DaysR2
TapRoot
AllRepro
DaysR1toR2
Manganese
Iron
Magnesium
FineRF
RootDiam
LAR
SSL
Nitrogen
SRL
TapRF
TapMF
2.0 1.5 1.0 0.5 0.0
07−01 07−02 07−03 07−04 07−05 07−06 07−07 07−08 07−09 07−10 07−11 07−12 07−13 07−14 07−15 07−16
Low Chr 7
Zinc
Copper
Calcium
RGRNitro
RGR
HarvestHeight
PlantWeight
AllAbove
Stem
AllLeaf
DaysR2
DaysR1
AllRoot
AllFine
TapRoot
RootDiam
SSL
Nitrogen
LMF
LAR
TapMF
2.0 1.5 1.0 0.5 0.0
07−01 07−02 07−03 07−04 07−05 07−06 07−07 07−08 07−09 07−10 07−11 07−12 07−13 07−14 07−15 07−16
logdiff Chr 7
SMF
HarvestHeight
DaysR2
PlantWeight
AllAbove
Stem
HarvestStemDiam
Manganese
LAR
Sulfur
1.5 1.0 0.5 0.0
07−01 07−02 07−03 07−04 07−05 07−06 07−07 07−08 07−09 07−10 07−11 07−12 07−13 07−14 07−15 07−16

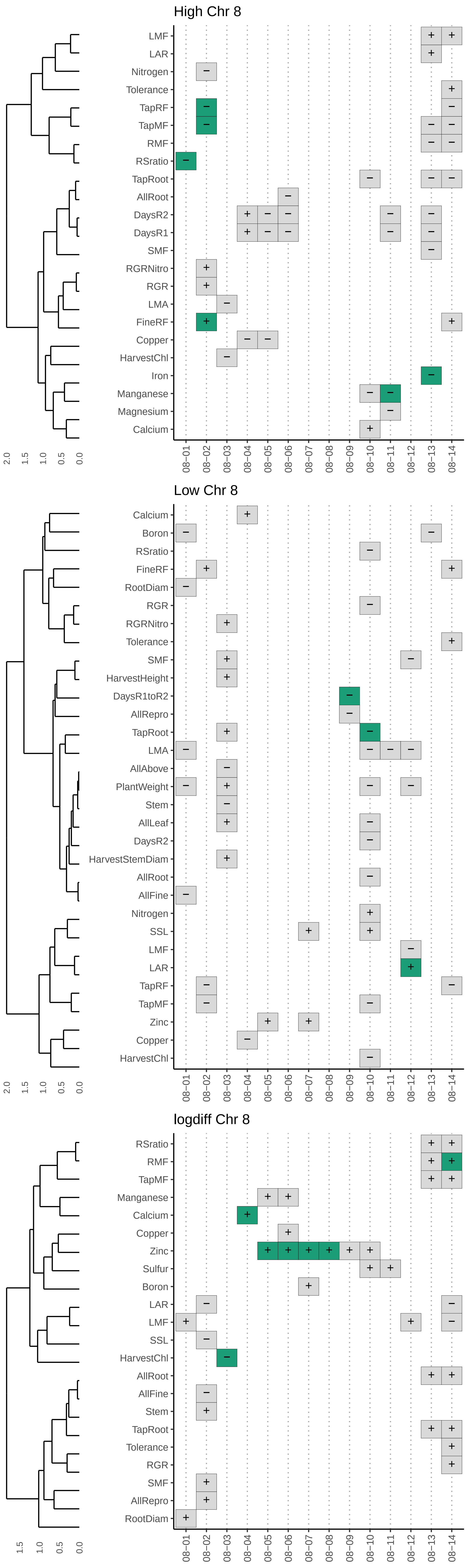
High Chr 8
LMF
LAR
Nitrogen
Tolerance
TapRF
TapMF
RMF
RSratio
TapRoot
AllRoot
DaysR2
DaysR1
SMF
RGRNitro
RGR
LMA
FineRF
Copper
HarvestChl
Iron
Manganese
Magnesium
Calcium
2.0 1.5 1.0 0.5 0.0
08−01 08−02 08−03 08−04 08−05 08−06 08−07 08−08 08−09 08−10 08−11 08−12 08−13 08−14
Low Chr 8
Calcium
Boron
RSratio
FineRF
RootDiam
RGR
RGRNitro
Tolerance
SMF
HarvestHeight
DaysR1toR2
AllRepro
TapRoot
LMA
AllAbove
PlantWeight
Stem
AllLeaf
DaysR2
HarvestStemDiam
AllRoot
AllFine
Nitrogen
SSL
LMF
LAR
TapRF
TapMF
Zinc
Copper
HarvestChl
2.0 1.5 1.0 0.5 0.0
08−01 08−02 08−03 08−04 08−05 08−06 08−07 08−08 08−09 08−10 08−11 08−12 08−13 08−14
logdiff Chr 8
RSratio
RMF
TapMF
Manganese
Calcium
Copper
Zinc
Sulfur
Boron
LAR
LMF
SSL
HarvestChl
AllRoot
AllFine
Stem
TapRoot
Tolerance
RGR
SMF
AllRepro
RootDiam
1.5 1.0 0.5 0.0
08−01 08−02 08−03 08−04 08−05 08−06 08−07 08−08 08−09 08−10 08−11 08−12 08−13 08−14

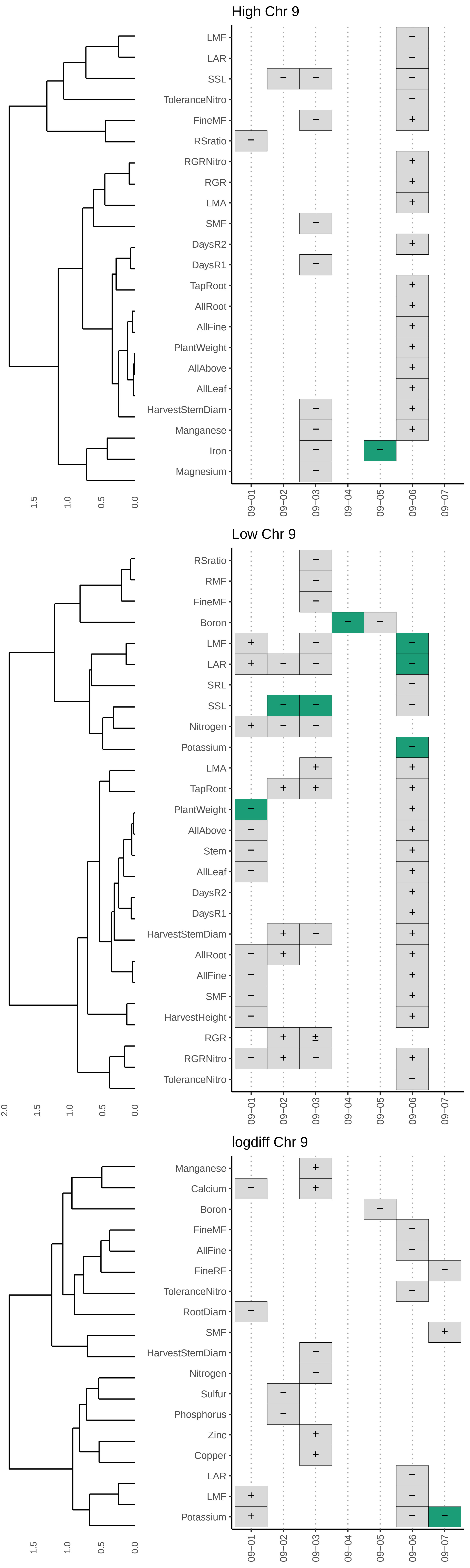
High Chr 9
LMF
LAR
SSL
ToleranceNitro
FineMF
RSratio
RGRNitro
RGR
LMA
SMF
DaysR2
DaysR1
TapRoot
AllRoot
AllFine
PlantWeight
AllAbove
AllLeaf
HarvestStemDiam
Manganese
Iron
Magnesium
Low Chr 9
RSratio
RMF
FineMF
Boron
LMF
LAR
SRL
SSL
Nitrogen
Potassium
LMA
TapRoot
PlantWeight
AllAbove
Stem
AllLeaf
DaysR2
DaysR1
HarvestStemDiam
AllRoot
AllFine
SMF
HarvestHeight
RGR
RGRNitro
ToleranceNitro
logdiff Chr 9
Manganese
Calcium
Boron
FineMF
AllFine
FineRF
ToleranceNitro
RootDiam
SMF
HarvestStemDiam
Nitrogen
Sulfur
Phosphorus
Zinc
Copper
LAR
LMF
Potassium
09-01
09-02
09-03
09-04
09-05
09-06
09-07
2.0
1.5
1.0
0.5
0.0

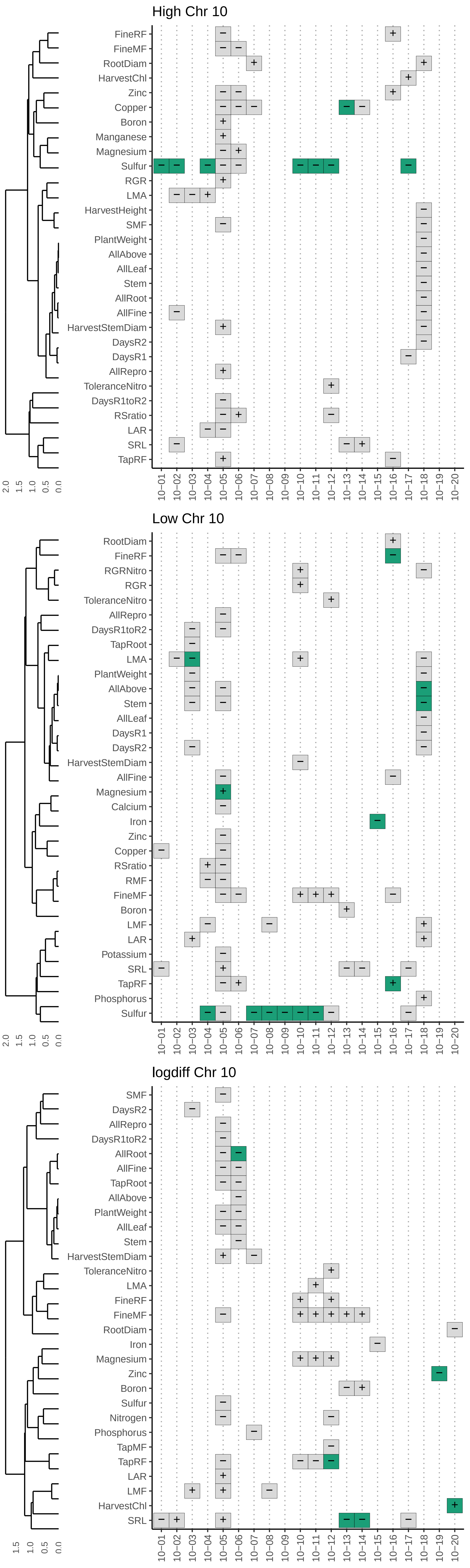
High Chr 10
FineRF
FineMF
RootDiam
HarvestChl
Zinc
Copper
Boron
Manganese
Magnesium
Sulfur
RGR
LMA
HarvestHeight
SMF
PlantWeight
AllAbove
AllLeaf
Stem
AllRoot
AllFine
HarvestStemDiam
DaysR2
DaysR1
AllRepro
ToleranceNitro
DaysR1toR2
RSratio
LAR
SRL
TapRF
2.0
1.5
1.0
0.5
0.0
10−01
10−02
10−03
10−04
10−05
10−06
10−07
10−08
10−09
10−10
10−11
10−12
10−13
10−14
10−15
10−16
10−17
10−18
10−19
10−20
Low Chr 10
RootDiam
FineRF
RGRNitro
RGR
ToleranceNitro
AllRepro
DaysR1toR2
TapRoot
LMA
PlantWeight
AllAbove
Stem
AllLeaf
DaysR1
DaysR2
HarvestStemDiam
AllFine
Magnesium
Calcium
Iron
Zinc
Copper
RSratio
RMF
FineMF
Boron
LMF
LAR
Potassium
SRL
TapRF
Phosphorus
Sulfur
2.0
1.5
1.0
0.5
0.0
10−01
10−02
10−03
10−04
10−05
10−06
10−07
10−08
10−09
10−10
10−11
10−12
10−13
10−14
10−15
10−16
10−17
10−18
10−19
10−20
logdiff Chr 10
SMF
DaysR2
AllRepro
DaysR1toR2
AllRoot
AllFine
TapRoot
AllAbove
PlantWeight
AllLeaf
Stem
HarvestStemDiam
ToleranceNitro
LMA
FineRF
FineMF
RootDiam
Iron
Magnesium
Zinc
Boron
Sulfur
Nitrogen
Phosphorus
TapMF
TapRF
LAR
LMF
HarvestChl
SRL
1.5
1.0
0.5
0.0
10−01
10−02
10−03
10−04
10−05
10−06
10−07
10−08
10−09
10−10
10−11
10−12
10−13
10−14
10−15
10−16
10−17
10−18
10−19
10−20

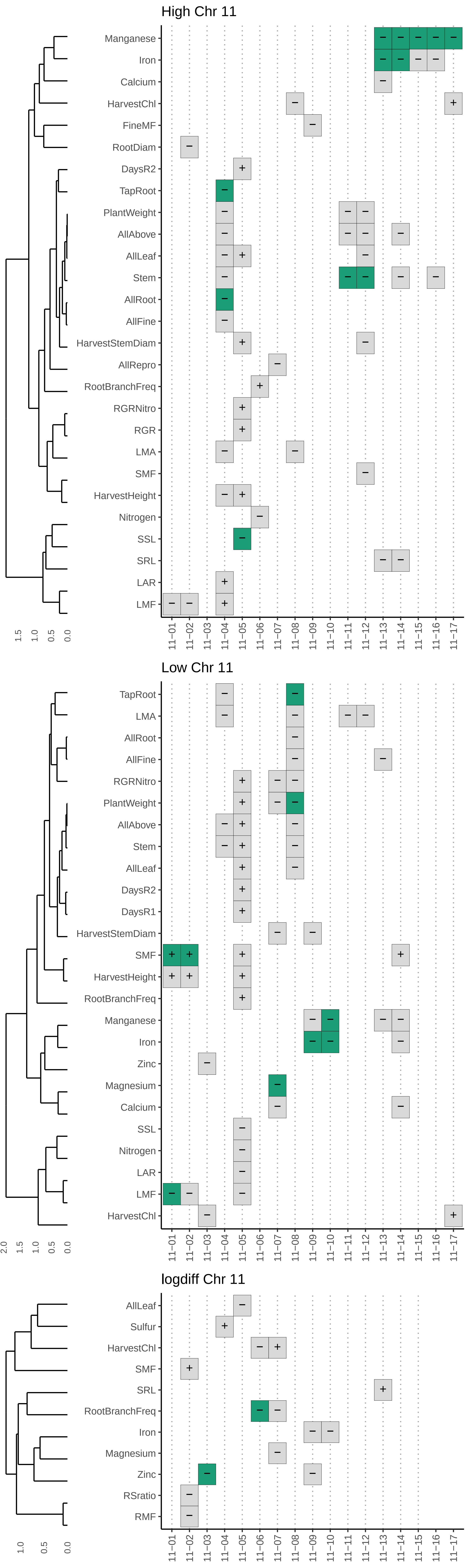

High Chr 11
Manganese
Iron
Calcium
HarvestChl
FineMF
RootDiam
DaysR2
TapRoot
PlantWeight
AllAbove
AllLeaf
Stem
AllRoot
AllFine
HarvestStemDiam
AllRepro
RootBranchFreq
RGRNitro
RGR
LMA
SMF
HarvestHeight
Nitrogen
SSL
SRL
LAR
LMF
Low Chr 11
TapRoot
LMA
AllRoot
AllFine
RGRNitro
PlantWeight
AllAbove
Stem
AllLeaf
DaysR2
DaysR1
HarvestStemDiam
SMF
HarvestHeight
RootBranchFreq
Manganese
Iron
Zinc
Magnesium
Calcium
SSL
Nitrogen
LAR
LMF
HarvestChl
logdiff Chr 11
AllLeaf
Sulfur
HarvestChl
SMF
SRL
RootBranchFreq
Iron
Magnesium
Zinc
RSratio
RMF
11-01
11-02
11-03
11-04
11-05
11-06
11-07
11-08
11-09
11-10
11-11
11-12
11-13
11-14
11-15
11-16
11-17
2.0
1.5
1.0
0.5
0.0

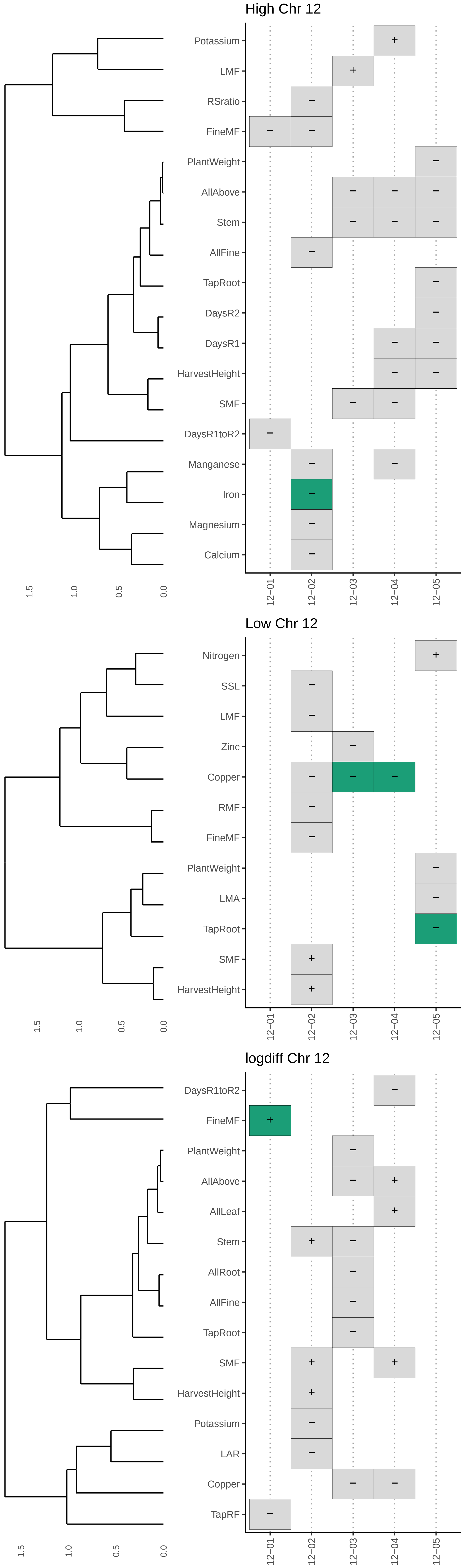
High Chr 12
Potassium
LMF
RSratio
FineMF
PlantWeight
AllAbove
Stem
AllFine
TapRoot
DaysR2
DaysR1
HarvestHeight
SMF
DaysR1toR2
Manganese
Iron
Magnesium
Calcium
1.5
1.0
0.5
0.0
12−01
12−02
12−03
12−04
12−05
Low Chr 12
Nitrogen
SSL
LMF
Zinc
Copper
RMF
FineMF
PlantWeight
LMA
TapRoot
SMF
HarvestHeight
1.5
1.0
0.5
0.0
12−01
12−02
12−03
12−04
12−05
logdiff Chr 12
DaysR1toR2
FineMF
PlantWeight
AllAbove
AllLeaf
Stem
AllRoot
AllFine
TapRoot
SMF
HarvestHeight
Potassium
LAR
Copper
TapRF
1.5
1.0
0.5
0.0
12−01
12−02
12−03
12−04
12−05

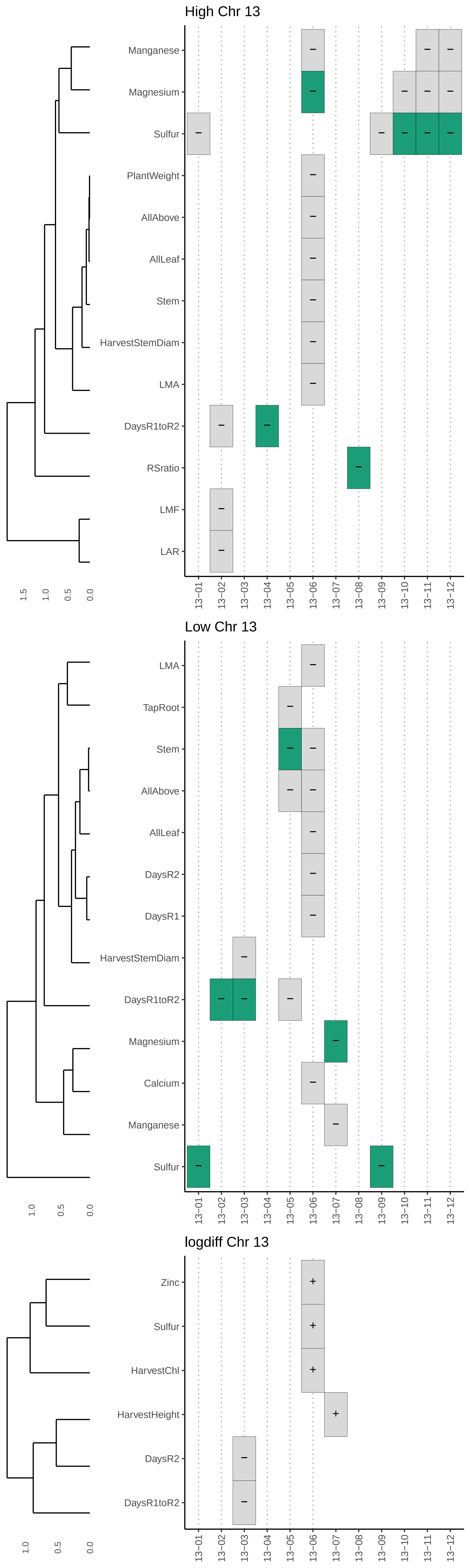
High Chr 13
Manganese
Magnesium
Sulfur
PlantWeight
AllAbove
AllLeaf
Stem
HarvestStemDiam
LMA
DaysR1toR2
RSratio
LMF
LAR
1.5
1.0
0.5
0.0
13−01 13−02 13−03 13−04 13−05 13−06 13−07 13−08 13−09 13−10 13−11 13−12
Low Chr 13
LMA
TapRoot
Stem
AllAbove
AllLeaf
DaysR2
DaysR1
HarvestStemDiam
DaysR1toR2
Magnesium
Calcium
Manganese
Sulfur
1.0
0.5
0.0
13−01 13−02 13−03 13−04 13−05 13−06 13−07 13−08 13−09 13−10 13−11 13−12
logdiff Chr 13
Zinc
Sulfur
HarvestChl
HarvestHeight
DaysR2
DaysR1toR2
1.0
0.5
0.0
13−01 13−02 13−03 13−04 13−05 13−06 13−07 13−08 13−09 13−10 13−11 13−12

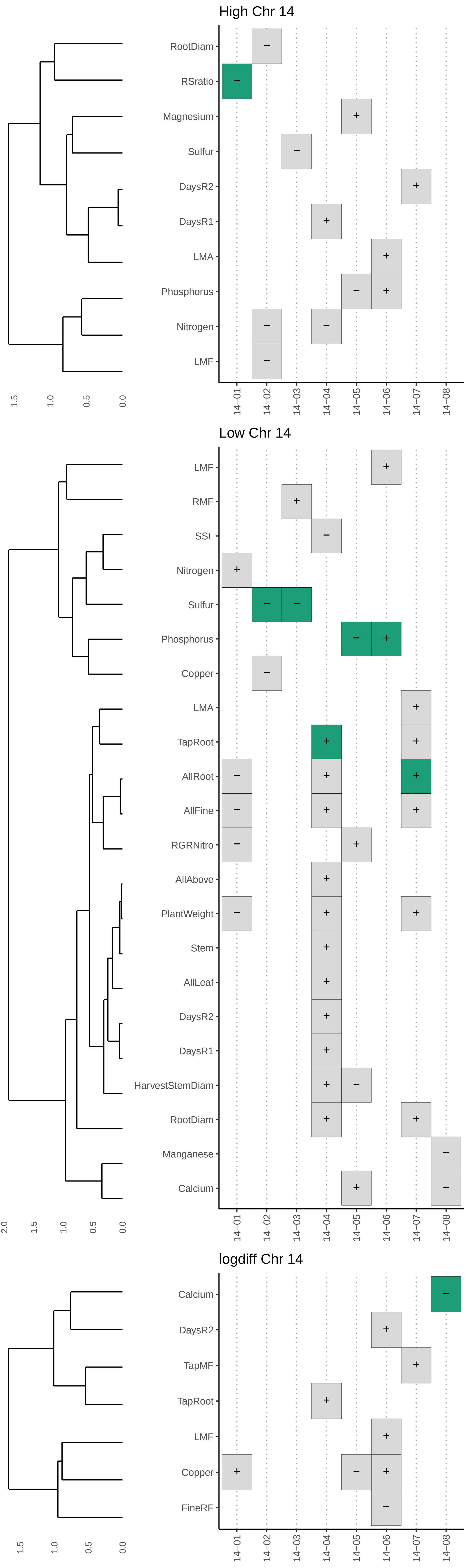
High Chr 14
RootDiam
RSratio
Magnesium
Sulfur
DaysR2
DaysR1
LMA
Phosphorus
Nitrogen
LMF
1.5
1.0
0.5
0.0
14−01
14−02
14−03
14−04
14−05
14−06
14−07
14−08
Low Chr 14
LMF
RMF
SSL
Nitrogen
Sulfur
Phosphorus
Copper
LMA
TapRoot
AllRoot
AllFine
RGRNitro
AllAbove
PlantWeight
Stem
AllLeaf
DaysR2
DaysR1
HarvestStemDiam
RootDiam
Manganese
Calcium
2.0
1.5
1.0
0.5
0.0
14−01
14−02
14−03
14−04
14−05
14−06
14−07
14−08
logdiff Chr 14
Calcium
DaysR2
TapMF
TapRoot
LMF
Copper
FineRF
1.5
1.0
0.5
0.0
14−01
14−02
14−03
14−04
14−05
14−06
14−07
14−08

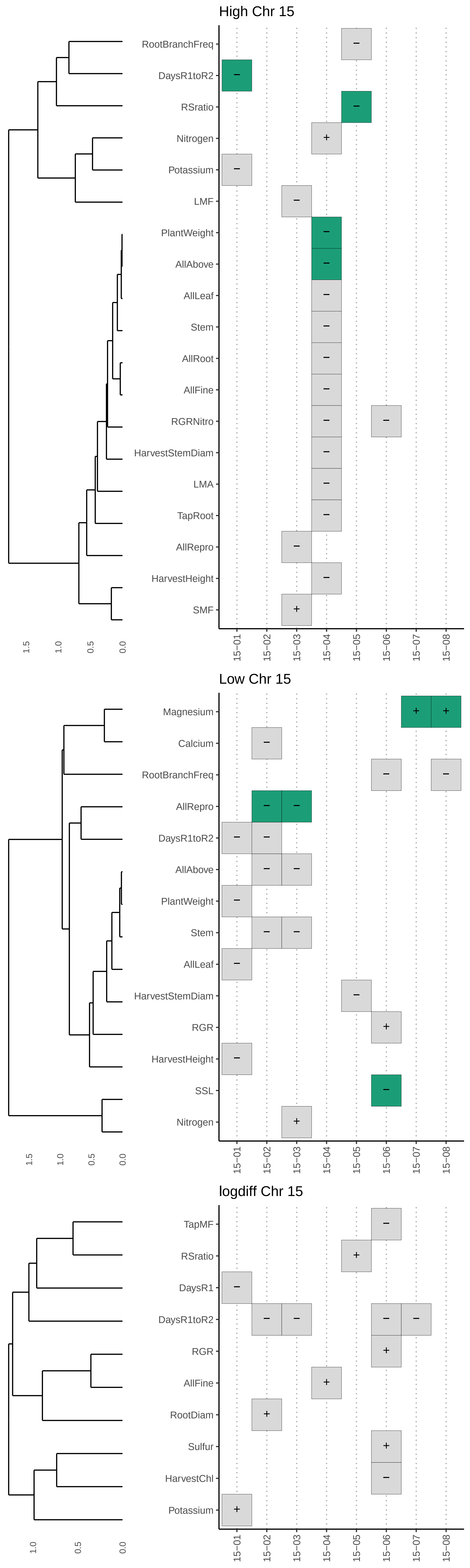
High Chr 15
RootBranchFreq
DaysR1toR2
RSratio
Nitrogen
Potassium
LMF
PlantWeight
AllAbove
AllLeaf
Stem
AllRoot
AllFine
RGRNitro
HarvestStemDiam
LMA
TapRoot
AllRepro
HarvestHeight
SMF
1.5
1.0
0.5
0.0
15−01
15−02
15−03
15−04
15−05
15−06
15−07
15−08
Low Chr 15
Magnesium
Calcium
RootBranchFreq
AllRepro
DaysR1toR2
AllAbove
PlantWeight
Stem
AllLeaf
HarvestStemDiam
RGR
HarvestHeight
SSL
Nitrogen
1.5
1.0
0.5
0.0
15−01
15−02
15−03
15−04
15−05
15−06
15−07
15−08
logdiff Chr 15
TapMF
RSratio
DaysR1
DaysR1toR2
RGR
AllFine
RootDiam
Sulfur
HarvestChl
Potassium
1.0
0.5
0.0
15−01
15−02
15−03
15−04
15−05
15−06
15−07
15−08

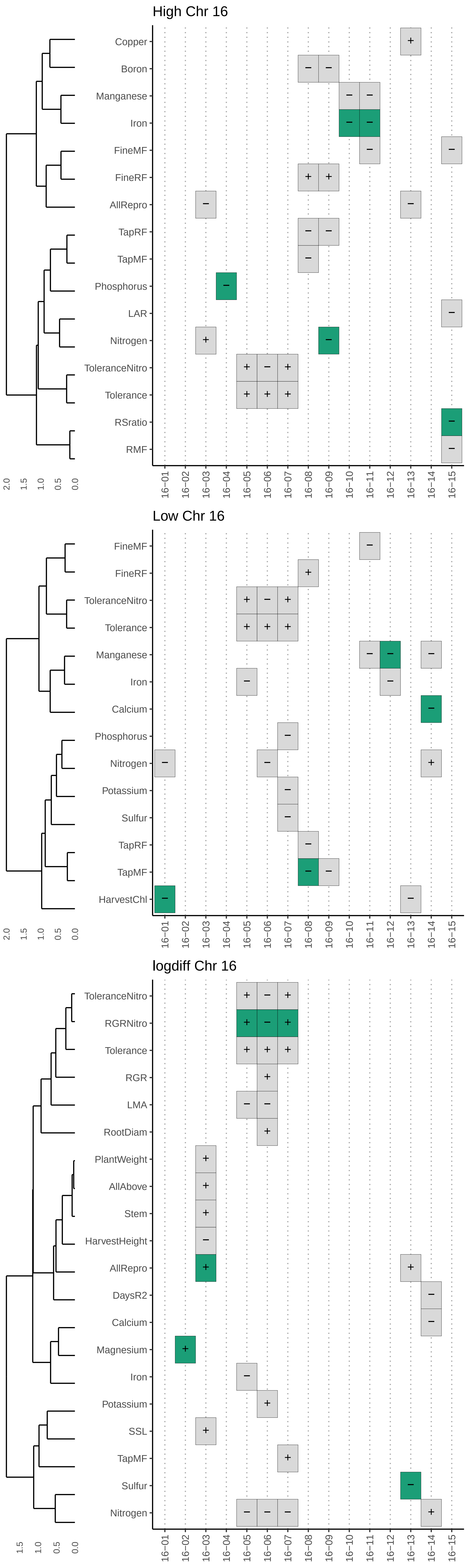
High Chr 16
Copper
Boron
Manganese
Iron
FineMF
FineRF
AllRepro
TapRF
TapMF
Phosphorus
LAR
Nitrogen
ToleranceNitro
Tolerance
RSratio
RMF
2.0 1.5 1.0 0.5 0.0
16–01 16–02 16–03 16–04 16–05 16–06 16–07 16–08 16–09 16–10 16–11 16–12 16–13 16–14 16–15
Low Chr 16
FineMF
FineRF
ToleranceNitro
Tolerance
Manganese
Iron
Calcium
Phosphorus
Nitrogen
Potassium
Sulfur
TapRF
TapMF
HarvestChl
2.0 1.5 1.0 0.5 0.0
16–01 16–02 16–03 16–04 16–05 16–06 16–07 16–08 16–09 16–10 16–11 16–12 16–13 16–14 16–15
logdiff Chr 16
ToleranceNitro
RGRNitro
Tolerance
RGR
LMA
RootDiam
PlantWeight
AllAbove
Stem
HarvestHeight
AllRepro
DaysR2
Calcium
Magnesium
Iron
Potassium
SSL
TapMF
Sulfur
Nitrogen
1.5 1.0 0.5 0.0
16–01 16–02 16–03 16–04 16–05 16–06 16–07 16–08 16–09 16–10 16–11 16–12 16–13 16–14 16–15

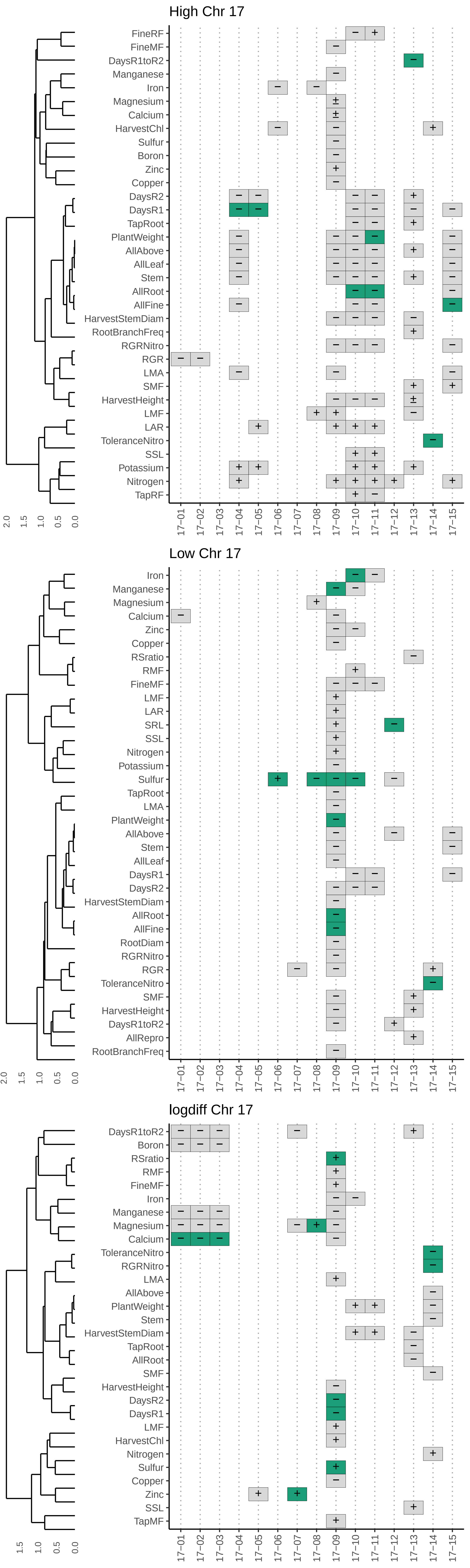
High Chr 17
FineRF
FineMF
DaysR1toR2
Manganese
Iron
Magnesium
Calcium
HarvestChl
Sulfur
Boron
Zinc
Copper
DaysR2
DaysR1
TapRoot
PlantWeight
AllAbove
AllLeaf
Stem
AllRoot
AllFine
HarvestStemDiam
RootBranchFreq
RGRNitro
RGR
LMA
SMF
HarvestHeight
LMF
LAR
ToleranceNitro
SSL
Potassium
Nitrogen
TapRF
2.0
1.5
1.0
0.5
0.0
17−01
17−02
17−03
17−04
17−05
17−06
17−07
17−08
17−09
17−10
17−11
17−12
17−13
17−14
17−15
Low Chr 17
Iron
Manganese
Magnesium
Calcium
Zinc
Copper
RSratio
RMF
FineMF
LMF
LAR
SRL
SSL
Nitrogen
Potassium
Sulfur
TapRoot
LMA
PlantWeight
AllAbove
Stem
AllLeaf
DaysR1
DaysR2
HarvestStemDiam
AllRoot
AllFine
RootDiam
RGRNitro
RGR
ToleranceNitro
SMF
HarvestHeight
DaysR1toR2
AllRepro
RootBranchFreq
2.0
1.5
1.0
0.5
0.0
17−01
17−02
17−03
17−04
17−05
17−06
17−07
17−08
17−09
17−10
17−11
17−12
17−13
17−14
17−15
logdiff Chr 17
DaysR1toR2
Boron
RSratio
RMF
FineMF
Iron
Manganese
Magnesium
Calcium
ToleranceNitro
RGRNitro
LMA
AllAbove
PlantWeight
Stem
HarvestStemDiam
TapRoot
AllRoot
SMF
HarvestHeight
DaysR2
DaysR1
LMF
HarvestChl
Nitrogen
Sulfur
Copper
Zinc
SSL
TapMF
1.5
1.0
0.5
0.0
17−01
17−02
17−03
17−04
17−05
17−06
17−07
17−08
17−09
17−10
17−11
17−12
17−13
17−14
17−15

Figure S7 **Significant regions on genome haplotype block map**
Haplotype blocks on the 17 individual sunflower genomes. Blocks are colored from the first SNP to the last SNP within each block. Adjacent blocks are colored in an alternating fashion repeating the same 4 colors. Gray marks along the x-axis note regions with significant trait associations.

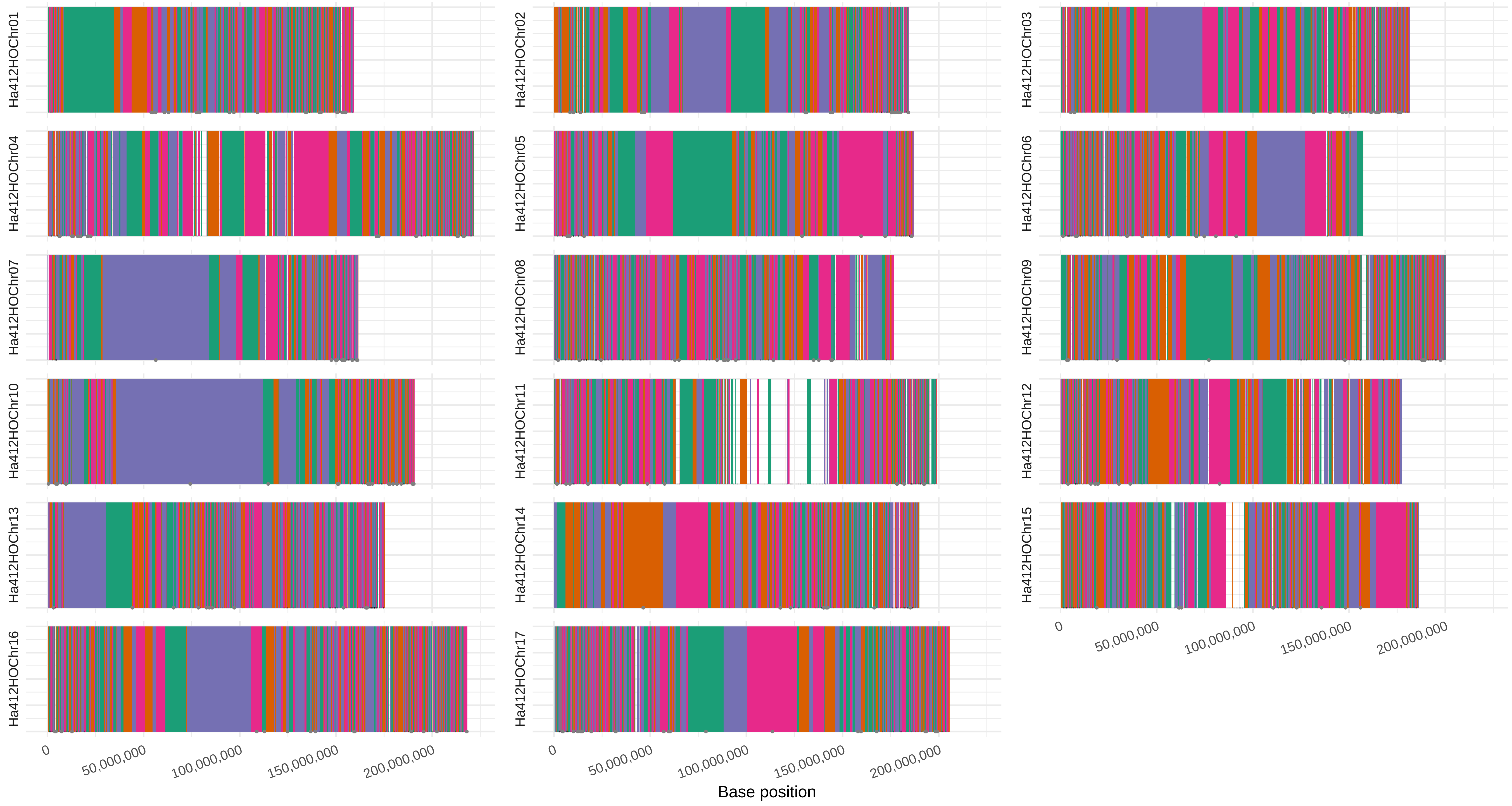
Ha412HOChr01
Ha412HOChr02
Ha412HOChr03
Ha412HOChr04
Ha412HOChr05
Ha412HOChr06
Ha412HOChr07
Ha412HOChr08
Ha412HOChr09
Ha412HOChr10
Ha412HOChr11
Ha412HOChr12
Ha412HOChr13
Ha412HOChr14
Ha412HOChr15
Ha412HOChr16
Ha412HOChr17
0
50,000,000
100,000,000
150,000,000
200,000,000
Base position
